## Supplemental Table 1 for "Rapalink-1 reveals novel mTOR-dependent genes and an agmatinergic axis-based metabolic feedback regulating mTOR activity and lifespan"

Supplementary Table 1

rapalink-1 sensitive (<0.75) and resistant (>1.25) strains

| Gene | Normalised fitness |
|---|---|
| SPAC1071.06 | 0.000732267 |
| SPAC1D4.11c | 0.001611177 |
| SPAC15E1.03 | 0.004153335 |
| SPCC126.04c | 0.005318496 |
| SPBC577.02 | 0.005466409 |
| SPAC1556.05c | 0.006331772 |
| SPBC405.07 | 0.006332439 |
| SPBC16A3.07c | 0.012706202 |
| SPAC26A3.04 | 0.014557613 |
| SPAC57A7.12 | 0.015578252 |
| SPAC1093.01 | 0.02011672 |
| SPAC328.02 | 0.020318473 |
| SPAC1039.08 | 0.024248519 |
| SPAC9G1.03c | 0.025566138 |
| SPAC4G8.13c | 0.027547084 |
| SPBC1347.02 | 0.029005551 |
| SPAC4G9.16c | 0.034326262 |
| SPAC11G7.02 | 0.035292691 |
| SPAC3F10.16c | 0.036511852 |
| SPAC1F12.10c | 0.040163548 |
| SPBC16C6.11 | 0.045932815 |
| SPAC1952.03 | 0.048751157 |
| SPAC3A12.13c | 0.051423635 |
| SPAPB17E12.05 | 0.056703796 |
| SPBC56F2.08c | 0.05740848 |
| SPAC23H4.17c | 0.058354646 |
| SPAC3F10.17 | 0.061192469 |
| SPAC22E12.18 | 0.073176441 |
| SPBC947.02 | 0.081920832 |
| SPAC10F6.13c | 0.087615775 |
| SPCPJ732.01 | 0.089020868 |
| SPBC83.13 | 0.089549435 |
| SPBC18H10.14 | 0.092907557 |
| SPBC11C11.07 | 0.111811324 |
| SPBC713.06 | 0.113445113 |
| SPBC1348.07 | 0.118964758 |
| SPBC29A3.18 | 0.120877362 |
| SPAPB1E7.12 | 0.123357761 |
| SPAC3H5.12c | 0.124434942 |
| SPBC11C11.02 | 0.125561595 |
| SPAC630.14c | 0.12639189 |
| SPBC16G5.15c | 0.127286178 |
| SPAPB1E7.02c | 0.130498213 |
| SPBPJ4664.01 | 0.132035291 |
| SPBC713.08 | 0.134519125 |

| | |
|---|---|
| SPBC13E7.04 | 0.135253406 |
| SPAC688.11 | 0.139265814 |
| SPCP1E11.06 | 0.151422554 |
| SPCC320.04c | 0.163685637 |
| SPCC162.12 | 0.170664298 |
| SPBC36.04 | 0.171028745 |
| SPBC1718.03 | 0.171514238 |
| SPBC56F2.11 | 0.171569811 |
| SPBC19C7.02 | 0.172079772 |
| SPAC3G9.03 | 0.179535837 |
| SPAC23E2.01 | 0.184799575 |
| SPBC106.17c | 0.186509801 |
| SPBC409.19c | 0.188514357 |
| SPAC1486.01 | 0.189761682 |
| SPAC8C9.10c | 0.193209541 |
| SPAC4F10.05c | 0.205500238 |
| SPCC74.05 | 0.206609854 |
| SPAC2C4.16c | 0.207695253 |
| SPAC30D11.10 | 0.208632269 |
| SPAPB18E9.01 | 0.209362 |
| SPAC23G3.03 | 0.223820271 |
| SPBC1D7.04 | 0.226669183 |
| SPAC227.05 | 0.23019376 |
| SPBC27B12.08 | 0.234482374 |
| SPBC3B9.13c | 0.235033259 |
| SPAC26F1.03 | 0.235866569 |
| SPAC4F8.03 | 0.236722307 |
| SPBC29A10.13 | 0.238002986 |
| SPAC31A2.14 | 0.239515084 |
| SPBC685.07c | 0.241400694 |
| SPCC1020.09 | 0.247229947 |
| SPBC9B6.07 | 0.251312705 |
| SPBC106.10 | 0.251316732 |
| SPCC663.04 | 0.260987654 |
| SPAC869.11 | 0.262827882 |
| SPAC4H3.07c | 0.262954468 |
| SPAC8E11.02c | 0.263429424 |
| SPCC320.12 | 0.267224552 |
| SPBC119.06 | 0.268728384 |
| SPCP1E11.09c | 0.273438712 |
| SPCC4B3.17 | 0.275426547 |
| SPBC16A3.17c | 0.278716234 |
| SPBC2G5.06c | 0.280209014 |
| SPBC776.04 | 0.280585347 |
| SPAC3H5.07 | 0.282615094 |
| SPBC16C6.01c | 0.286388719 |
| SPBC19G7.16 | 0.290698108 |
| SPBC83.12 | 0.293799669 |

| | |
|---|---|
| SPAC513.03 | 0.297071265 |
| SPBC577.04 | 0.297260297 |
| SPBC725.10 | 0.299512522 |
| SPAC22H10.11c | 0.304154706 |
| SPCC584.01c | 0.308709884 |
| SPAC22F3.09c | 0.310327052 |
| SPBC16C6.03c | 0.313996268 |
| SPBC1703.12 | 0.314088351 |
| SPCC1739.14 | 0.314127743 |
| SPAC22F3.08c | 0.314879561 |
| SPBC1709.04c | 0.317203748 |
| SPAC167.07c | 0.317253488 |
| SPAC1782.11 | 0.318435343 |
| SPBC17G9.07 | 0.320706538 |
| SPAC25G10.03 | 0.326675584 |
| SPCP1E11.10 | 0.326866694 |
| SPAC30D11.13 | 0.330948555 |
| SPAC227.17c | 0.332482125 |
| SPAC30C2.04 | 0.332764777 |
| SPAC4G8.11c | 0.334598251 |
| SPAC25G10.06 | 0.3352195 |
| SPBC2G2.13c | 0.335228819 |
| SPBC16G5.11c | 0.337637345 |
| SPBC56F2.09c | 0.33981847 |
| SPBC1652.02 | 0.340865903 |
| SPCC1183.04c | 0.34720679 |
| SPBC21B10.13c | 0.348148148 |
| SPBC1539.10 | 0.350040675 |
| SPBC2D10.07c | 0.355387289 |
| SPAC3C7.08c | 0.357124458 |
| SPAC959.08 | 0.3620089 |
| SPBC3D6.08c | 0.362839958 |
| SPBC337.13c | 0.364778226 |
| SPBC25D12.06 | 0.365089955 |
| SPAC1486.04c | 0.368153844 |
| SPCC191.01 | 0.368221398 |
| SPAC29A4.09 | 0.369121562 |
| SPAC1F5.10 | 0.371625758 |
| SPBC83.02c | 0.372605566 |
| SPCC1682.14 | 0.378219546 |
| SPBC17G9.10 | 0.38051512 |
| SPAC8E11.05c | 0.383832128 |
| SPCC1223.05c | 0.385179969 |
| SPAC23A1.07 | 0.386369461 |
| SPAC5D6.12 | 0.389266256 |
| SPBC365.06 | 0.392822457 |
| SPCC1259.01c | 0.394881786 |
| SPAC144.08 | 0.394957106 |

| | |
|---|---|
| SPBC1734.11 | 0.406396871 |
| SPBC15D4.06 | 0.407029465 |
| SPAC17H9.09c | 0.407783775 |
| SPAC6G9.10c | 0.408113122 |
| SPAC8C9.12c | 0.408612762 |
| SPBC16A3.03c | 0.40961804 |
| SPCC11E10.08 | 0.415157885 |
| SPBC27.08c | 0.419114419 |
| SPBC13E7.11 | 0.423987164 |
| SPCC895.06 | 0.424930341 |
| SPBP35G2.08c | 0.426907313 |
| SPCC1259.03 | 0.427369567 |
| SPAC6B12.12 | 0.429157077 |
| SPAC26F1.10c | 0.431485124 |
| SPAC29B12.04 | 0.431979513 |
| SPAC23H3.03c | 0.433447156 |
| SPBC577.08c | 0.436660759 |
| SPAC29A4.20 | 0.437358188 |
| SPBC1347.06c | 0.438428506 |
| SPBC19F8.06c | 0.450074405 |
| SPBC17A3.06 | 0.450194993 |
| SPBC1683.09c | 0.451227033 |
| SPAC22H12.02 | 0.451818917 |
| SPBC25D12.02c | 0.455111126 |
| SPCC736.07c | 0.455354322 |
| SPAPB17E12.04c | 0.457146441 |
| SPAC22H10.09 | 0.459100651 |
| SPAC1486.08 | 0.460403939 |
| SPBP4H10.11c | 0.461077416 |
| SPAC3H8.07c | 0.464428302 |
| SPAC9G1.12 | 0.46629179 |
| SPAP27G11.14c | 0.468580542 |
| SPBC1604.11 | 0.469078947 |
| SPCC970.05 | 0.469842276 |
| SPBC16H5.08c | 0.471698466 |
| SPBC215.02 | 0.472702475 |
| SPAC22A12.04c | 0.474310089 |
| SPBC15D4.10c | 0.474316955 |
| SPAC3H5.10 | 0.475045659 |
| SPAC3A11.07 | 0.476259957 |
| SPBC530.11c | 0.478605826 |
| SPAC521.05 | 0.481745427 |
| SPCC1753.02c | 0.485195852 |
| SPAC227.18 | 0.485870269 |
| SPAP27G11.06c | 0.487217467 |
| SPAC23G3.02c | 0.488437935 |
| SPAC14C4.06c | 0.488803476 |
| SPAC8C9.03 | 0.490812604 |

| | |
|---|---|
| SPBC725.02 | 0.49866822 |
| SPAC26F1.14c | 0.498782059 |
| SPAC3A11.13 | 0.501623681 |
| SPCC191.07 | 0.501997014 |
| SPCC4G3.04c | 0.507250295 |
| SPAC1006.03c | 0.507448356 |
| SPAC25B8.05 | 0.507588431 |
| SPCC11E10.06c | 0.508337501 |
| SPBC21B10.10 | 0.511723474 |
| SPBC14F5.09c | 0.512489112 |
| SPAC17A5.02c | 0.514075481 |
| SPCC1393.03 | 0.514590449 |
| SPAC30.02c | 0.518156653 |
| SPBC1271.05c | 0.519722936 |
| SPAC22E12.03c | 0.521639559 |
| SPBC19C2.13c | 0.530615044 |
| SPCC1739.06c | 0.531956893 |
| SPAC1805.04 | 0.538024455 |
| SPAC328.03 | 0.539533585 |
| SPAC24H6.11c | 0.541598584 |
| SPAC23G3.08c | 0.541771297 |
| SPBC354.10 | 0.548742107 |
| SPBC4F6.05c | 0.551225928 |
| SPCC777.13 | 0.553593155 |
| SPBP26C9.03c | 0.553612673 |
| SPBC30B4.06c | 0.555133269 |
| SPAC22E12.11c | 0.555405232 |
| SPAC19B12.11c | 0.555530118 |
| SPBC776.15c | 0.555651717 |
| SPBC21C3.20c | 0.558707475 |
| SPAC8F11.02c | 0.558921013 |
| SPBC16E9.07 | 0.562131051 |
| SPAC19G12.15c | 0.562365794 |
| SPAC222.08c | 0.562570644 |
| SPAC22E12.04 | 0.562739397 |
| SPCC188.07 | 0.563193617 |
| SPAC227.13c | 0.563828767 |
| SPAC823.12 | 0.564190141 |
| SPAPB8E5.06c | 0.564772727 |
| SPAC631.02 | 0.56673321 |
| SPCC4G3.19 | 0.567009591 |
| SPBC32H8.07 | 0.567371097 |
| SPBPB2B2.10c | 0.568453657 |
| SPAC17G8.06c | 0.569995127 |
| SPBC428.05c | 0.570527309 |
| SPAPYUG7.02c | 0.570696086 |
| SPCC4B3.05c | 0.574025315 |
| SPBC651.03c | 0.574515329 |

| | |
|---|---|
| SPCC1322.01 | 0.575008464 |
| SPBC24C6.05 | 0.576834951 |
| SPAC3H8.09c | 0.584200796 |
| SPBC26H8.14c | 0.586350748 |
| SPAC3A12.10 | 0.588174725 |
| SPCC1753.05 | 0.591615771 |
| SPBC3H7.10 | 0.596286633 |
| SPAC2G11.10c | 0.596785427 |
| SPCC757.09c | 0.59687787 |
| SPCC970.07c | 0.598391468 |
| SPAC4A8.03c | 0.599110962 |
| SPBC1306.02 | 0.599610255 |
| SPAC1250.02 | 0.600956697 |
| SPBC14F5.10c | 0.600964412 |
| SPBC365.16 | 0.601030063 |
| SPBP16F5.05c | 0.602352867 |
| SPAC17H9.08 | 0.602995435 |
| SPCC1442.15c | 0.603246548 |
| SPCC1840.06 | 0.607531978 |
| SPAC3G9.07c | 0.610012453 |
| SPBC1683.04 | 0.611007171 |
| SPBC1709.09 | 0.612716344 |
| SPAC1782.06c | 0.61409493 |
| SPBC3E7.05c | 0.61439003 |
| SPBC1815.01 | 0.614552213 |
| SPAC12B10.12c | 0.615278678 |
| SPCC4G3.05c | 0.616302651 |
| SPAC144.04c | 0.616415958 |
| SPCC1235.05c | 0.617307374 |
| SPAC23C4.02 | 0.619176817 |
| SPBC13G1.10c | 0.619370118 |
| SPBC215.04 | 0.623377144 |
| SPBC839.04 | 0.623679051 |
| SPCC4E9.02 | 0.625223344 |
| SPBC2D10.12 | 0.626868124 |
| SPAC16.04 | 0.627090461 |
| SPBC23G7.16 | 0.627686368 |
| SPCC777.03c | 0.62785597 |
| SPBC1347.08c | 0.628019583 |
| SPAC1071.02 | 0.628701663 |
| SPBC1604.09c | 0.628816259 |
| SPBC31E1.02c | 0.631621655 |
| SPCC11E10.07c | 0.632941365 |
| SPAC1327.01c | 0.633370734 |
| SPBC13A2.02 | 0.63352377 |
| SPBC8E4.01c | 0.635048278 |
| SPCC1259.12c | 0.636361657 |
| SPCC4B3.02c | 0.637347921 |

| | |
|---|---|
| SPAC23C11.02c | 0.639291081 |
| SPAC17A5.01 | 0.639384547 |
| SPBC30B4.08 | 0.640344011 |
| SPBPB10D8.07c | 0.640991094 |
| SPAC13G7.02c | 0.641760455 |
| SPAC1851.03 | 0.642018509 |
| SPAC4G9.02 | 0.642088403 |
| SPBC1347.12 | 0.64276733 |
| SPBC1271.11 | 0.644621995 |
| SPBC12C2.03c | 0.645860831 |
| SPAC4F10.19c | 0.646181749 |
| SPAC12B10.03 | 0.64691459 |
| SPAC17H9.19c | 0.647690607 |
| SPAC4F10.16c | 0.648090456 |
| SPCC16C4.11 | 0.648580503 |
| SPBC21D10.11c | 0.649023658 |
| SPCC132.02 | 0.649777154 |
| SPBC21C3.14c | 0.651263222 |
| SPBC1711.03 | 0.651821678 |
| SPBC336.14c | 0.6519824 |
| SPAC11G7.03 | 0.652028649 |
| SPAC222.05c | 0.653337503 |
| SPCC338.16 | 0.65373483 |
| SPAC6G9.14 | 0.654345842 |
| SPAC664.04c | 0.654572502 |
| SPBC36.07 | 0.656077004 |
| SPAC22A12.01c | 0.656175468 |
| SPBC1604.19c | 0.656742381 |
| SPAC1071.08 | 0.657179439 |
| SPBC1734.13 | 0.658174434 |
| SPAC3H8.10 | 0.658357563 |
| SPAC25H1.03 | 0.658572497 |
| SPBC1289.06c | 0.659195564 |
| SPBC21D10.10 | 0.659608274 |
| SPBC1604.08c | 0.659888156 |
| SPBC13E7.09 | 0.66050447 |
| SPBC839.13c | 0.661947342 |
| SPCC965.06 | 0.662012363 |
| SPAC1805.12c | 0.663318559 |
| SPBC800.02 | 0.663977221 |
| SPAC2H10.02c | 0.664051791 |
| SPCC777.04 | 0.66449476 |
| SPAC343.12 | 0.665160632 |
| SPAC1610.02c | 0.666567237 |
| SPBC3H7.03c | 0.66678644 |
| SPAC3C7.06c | 0.667087095 |
| SPBP35G2.13c | 0.667908958 |
| SPBC19G7.06 | 0.66877202 |

| | |
|---|---|
| SPAC167.06c | 0.671986284 |
| SPBC18E5.05c | 0.672321169 |
| SPBC1734.12c | 0.675392086 |
| SPBC25H2.03 | 0.67594154 |
| SPAC23H3.15c | 0.676354573 |
| SPBC2G5.03 | 0.676490292 |
| SPBC16A3.08c | 0.676895502 |
| SPAC17A5.04c | 0.676915428 |
| SPAC20G4.02c | 0.677773367 |
| SPAC31G5.11 | 0.677893724 |
| SPAC24H6.09 | 0.679413715 |
| SPBC1347.09 | 0.679817229 |
| SPAC23C4.08 | 0.680154856 |
| SPAC25H1.07 | 0.680155345 |
| SPAC13G7.13c | 0.6803578 |
| SPCC23B6.02c | 0.681916532 |
| SPBC16E9.12c | 0.683074963 |
| SPCPB1C11.02 | 0.68377028 |
| SPBC1347.03 | 0.684122021 |
| SPAC2C4.06c | 0.685138734 |
| SPBC557.02c | 0.685657278 |
| SPCC18.10 | 0.687693169 |
| SPCC576.04 | 0.689053508 |
| SPCC825.05c | 0.6895546 |
| SPAC1F12.06c | 0.690073511 |
| SPAC806.07 | 0.690280675 |
| SPCC569.08c | 0.690725142 |
| SPAC1F7.06 | 0.691037138 |
| SPBC30B4.02c | 0.691876597 |
| SPAC4G9.20c | 0.692270771 |
| SPAC13G7.05 | 0.692297751 |
| SPCC1442.05c | 0.692507855 |
| SPBC29A10.06c | 0.692864431 |
| SPCC4B3.11c | 0.693437114 |
| SPBC29A3.08 | 0.694048371 |
| SPAC3F10.11c | 0.694821398 |
| SPCC970.10c | 0.696090828 |
| SPCC4G3.13c | 0.696277542 |
| SPBC2G2.01c | 0.697879666 |
| SPAC186.02c | 0.70013812 |
| SPBC14C8.03 | 0.700186772 |
| SPAC1F3.09 | 0.700900602 |
| SPCC63.06 | 0.701136175 |
| SPAC21E11.04 | 0.701882027 |
| SPAC1F8.05 | 0.70231127 |
| SPAC26F1.04c | 0.703008725 |
| SPBC27.06c | 0.703306715 |
| SPCC645.14c | 0.703878179 |

| | |
|---|---|
| SPAC3H1.10 | 0.704109937 |
| SPAC637.09 | 0.704859355 |
| SPAC6F6.09 | 0.705003115 |
| SPBC1105.01 | 0.705469428 |
| SPBC2A9.05c | 0.705864589 |
| SPCC126.06 | 0.707252657 |
| SPAC27F1.03c | 0.707492555 |
| SPCC1442.02 | 0.709638932 |
| SPAC1071.11 | 0.70972973 |
| SPAC22F3.04 | 0.711876552 |
| SPAC694.02 | 0.714007102 |
| SPAC644.15 | 0.714444262 |
| SPAC5D6.07c | 0.715189424 |
| SPAC6G9.09c | 0.715678143 |
| SPBC21D10.12 | 0.717233139 |
| SPBC21C3.08c | 0.719615587 |
| SPBC25B2.07c | 0.719831367 |
| SPBP4H10.13 | 0.720685339 |
| SPAC13C5.01c | 0.720773035 |
| SPAC1A6.10 | 0.722429823 |
| SPCC1020.01c | 0.722575027 |
| SPAC23C4.09c | 0.722705451 |
| SPAC30D11.07 | 0.723243298 |
| SPBC660.11 | 0.723340804 |
| SPBC29A3.10c | 0.723780489 |
| SPCC132.04c | 0.725847124 |
| SPAC6B12.02c | 0.72608621 |
| SPBC19G7.18c | 0.726160518 |
| SPBC119.03 | 0.72623911 |
| SPBC31F10.09c | 0.72833764 |
| SPBC30D10.18c | 0.730772118 |
| SPBC21C3.15c | 0.731468009 |
| SPAC977.16c | 0.731588501 |
| SPAC750.05c | 0.731618074 |
| SPCC1795.02c | 0.732809172 |
| SPBC23G7.15c | 0.732910157 |
| SPAC29B12.08 | 0.733094663 |
| SPBC1706.01 | 0.733094938 |
| SPAC6G10.12c | 0.734495214 |
| SPAC144.01 | 0.734690569 |
| SPAC3G6.04 | 0.734723309 |
| SPAC4G8.05 | 0.735204327 |
| SPBC336.13c | 0.735477833 |
| SPCC18.06c | 0.736213066 |
| SPAC4A8.02c | 0.736523869 |
| SPBC1703.06 | 0.736806237 |
| SPAC3H1.04c | 0.73690816 |
| SPAC9E9.09c | 0.737649721 |

| | |
|---|---|
| SPBC16A3.19 | 0.737846051 |
| SPCC31H12.04c | 0.738313109 |
| SPBC1709.11c | 0.739926227 |
| SPCC622.12c | 0.740371507 |
| SPAC1565.01 | 0.740778323 |
| SPAC343.19 | 0.741571761 |
| SPCC4E9.01c | 0.741792504 |
| SPAC1002.19 | 0.742054656 |
| SPAC664.14 | 0.74231559 |
| SPBC1773.13 | 0.742357083 |
| SPBC12D12.07c | 0.742760188 |
| SPCC1494.03 | 0.742946658 |
| SPCC285.17 | 0.742973615 |
| SPBC18H10.09 | 0.743053744 |
| SPAPB2B4.03 | 0.743457434 |
| SPBC1271.15c | 0.744128515 |
| SPAC23C4.12 | 0.744807439 |
| SPCC777.10c | 0.746224667 |
| SPAC17G8.11c | 0.746523423 |
| SPCC1259.05c | 0.746743035 |
| SPAC2C4.10c | 0.747215306 |
| SPAC23C11.01 | 0.747321429 |
| SPBC15D4.15 | 0.748015099 |
| SPAC977.05c | 0.74818321 |
| SPCC11E10.03 | 0.748699578 |
| SPAC15A10.06 | 1.2500047 |
| SPAPJ760.02c | 1.251731226 |
| SPAC6B12.05c | 1.252160047 |
| SPAC3G6.01 | 1.252740858 |
| SPAC15A10.08 | 1.253953687 |
| SPAC323.03c | 1.254007511 |
| SPBC25H2.10c | 1.254235953 |
| SPAC11D3.15 | 1.254793546 |
| SPAC1952.11c | 1.255135574 |
| SPAC5H10.07 | 1.255452194 |
| SPBC16G5.06 | 1.255578027 |
| SPCC613.06 | 1.255631122 |
| SPBC646.13 | 1.256779835 |
| SPBC354.03 | 1.256967262 |
| SPBC29A3.11c | 1.258860471 |
| SPCC4F11.02 | 1.258916459 |
| SPAC3H5.05c | 1.260315544 |
| SPAC18G6.09c | 1.261073312 |
| SPAC9.06c | 1.261334713 |
| SPBC12C2.12c | 1.261568493 |
| SPAC7D4.06c | 1.261622888 |
| SPAC2F3.12c | 1.262341691 |
| SPBC725.05c | 1.262860943 |

| | |
|---|---|
| SPBC4B4.04 | 1.263111551 |
| SPAC694.04c | 1.263131873 |
| SPBC1861.07 | 1.263331777 |
| SPBC19C2.06c | 1.26335598 |
| SPBC36.03c | 1.263477875 |
| SPAC3F10.04 | 1.263517082 |
| SPCC613.02 | 1.264431347 |
| SPAC4A8.10 | 1.264924347 |
| SPBP8B7.05c | 1.265556191 |
| SPBP22H7.08 | 1.265579702 |
| SPAC23G3.12c | 1.265897407 |
| SPBP35G2.07 | 1.266377444 |
| SPAC9G1.02 | 1.268965405 |
| SPAC926.03 | 1.270554103 |
| SPBC3H7.11 | 1.27294806 |
| SPBC106.04 | 1.273001458 |
| SPBC21B10.05c | 1.273101532 |
| SPAP8A3.12c | 1.273611373 |
| SPBC1778.02 | 1.274040392 |
| SPBC13G1.08c | 1.275261771 |
| SPAC1834.04 | 1.276118421 |
| SPAC17A5.09c | 1.276461968 |
| SPAP8A3.07c | 1.277332665 |
| SPBC29A10.16c | 1.277557079 |
| SPCC737.09c | 1.278936456 |
| SPBC1289.15 | 1.279809884 |
| SPAC30D11.11 | 1.280109445 |
| SPBC4F6.08c | 1.282387327 |
| SPAC644.06c | 1.282448126 |
| SPBC12C2.08 | 1.283293635 |
| SPBC21C3.19 | 1.283400232 |
| SPAC1B3.03c | 1.283522727 |
| SPAC1805.16c | 1.283756875 |
| SPAC1805.11c | 1.284570617 |
| SPBC27B12.03c | 1.286887089 |
| SPBC336.05c | 1.289061322 |
| SPAC1071.04c | 1.290535445 |
| SPCC1739.13 | 1.291380097 |
| SPBC3H7.05c | 1.291853534 |
| SPBC106.08c | 1.292759289 |
| SPAC11E3.11c | 1.294872204 |
| SPCC126.03 | 1.296626942 |
| SPBC1271.12 | 1.297265716 |
| SPAC1527.01 | 1.297447399 |
| SPBC342.03 | 1.298487275 |
| SPAC869.01 | 1.299285695 |
| SPAC19G12.13c | 1.300358852 |
| SPAC105.02c | 1.301138531 |

| | |
|---|---|
| SPBC2D10.13 | 1.301899414 |
| SPAC105.01c | 1.302125308 |
| SPAC1639.02c | 1.302159688 |
| SPBC4B4.06 | 1.302814179 |
| SPAC683.02c | 1.303006869 |
| SPBC29A10.02 | 1.304223715 |
| SPBC21C3.01c | 1.304524218 |
| SPCC63.02c | 1.305489322 |
| SPBC2G2.03c | 1.306610639 |
| SPAC25G10.09c | 1.307102744 |
| SPCP1E11.03 | 1.307627664 |
| SPBC1347.01c | 1.307649214 |
| SPBC3D6.13c | 1.311363707 |
| SPAC13G6.09 | 1.31183827 |
| SPBC336.01 | 1.312681788 |
| SPCC18.09c | 1.312803997 |
| SPAC15A10.11 | 1.31280438 |
| SPCC63.08c | 1.313441909 |
| SPAC17G6.08 | 1.315816336 |
| SPCC1919.07 | 1.31867591 |
| SPAC1F5.05c | 1.319039448 |
| SPAC29A4.19c | 1.322154676 |
| SPAC6C3.08 | 1.322534664 |
| SPBC17D1.02 | 1.322558979 |
| SPAC4A8.04 | 1.323064099 |
| SPAC5H10.01 | 1.323271323 |
| SPBC1347.07 | 1.3236998 |
| SPCC23B6.05c | 1.325614777 |
| SPBC1604.18c | 1.325999784 |
| SPAC607.09c | 1.326071253 |
| SPAC1B3.10c | 1.32612071 |
| SPBC1921.04c | 1.329016172 |
| SPCP1E11.02 | 1.329620492 |
| SPAC227.01c | 1.329927643 |
| SPCC1682.12c | 1.331237363 |
| SPCC285.16c | 1.333704813 |
| SPAC3H8.04 | 1.335677918 |
| SPBC16E9.19 | 1.336514939 |
| SPBC28F2.03 | 1.337224477 |
| SPAC19A8.05c | 1.33812294 |
| SPAC9E9.03 | 1.339191862 |
| SPBC3H7.14 | 1.340393976 |
| SPAC57A7.08 | 1.344449296 |
| SPBC36.10 | 1.344665038 |
| SPAC13C5.02 | 1.345268476 |
| SPCC965.09 | 1.348235997 |
| SPAC6F6.12 | 1.350042906 |
| SPCC188.02 | 1.353312147 |

| | |
|---|---|
| SPAP32A8.03c | 1.353748491 |
| SPCC1322.05c | 1.35602267 |
| SPAC890.03 | 1.356046608 |
| SPAC589.02c | 1.356563957 |
| SPBC1604.02c | 1.357513027 |
| SPAC13G6.02c | 1.35961437 |
| SPAC644.09 | 1.360903412 |
| SPBC3B8.02 | 1.361527431 |
| SPBC3B9.09 | 1.361651577 |
| SPAC24H6.03 | 1.361972886 |
| SPCC1494.07 | 1.362261695 |
| SPBC1711.04 | 1.363781122 |
| SPAC17A5.16 | 1.365847503 |
| SPBC1734.15 | 1.366950698 |
| SPCC1494.08c | 1.368325285 |
| SPAC1B3.07c | 1.371835858 |
| SPAC4D7.11 | 1.373339108 |
| SPCC4B3.07 | 1.374058823 |
| SPAC19B12.10 | 1.374280268 |
| SPBC146.13c | 1.375150442 |
| SPAC29B12.06c | 1.375658954 |
| SPCC74.03c | 1.376409534 |
| SPAC144.06 | 1.377556137 |
| SPAC13C5.07 | 1.379980755 |
| SPCC569.06 | 1.382365212 |
| SPCC4B3.03c | 1.387446884 |
| SPBC1861.05 | 1.38956557 |
| SPAC16A10.03c | 1.391100927 |
| SPAC2C4.15c | 1.392838518 |
| SPBC947.15c | 1.394545455 |
| SPBC31F10.17c | 1.395906433 |
| SPCC553.01c | 1.396793322 |
| SPBC24C6.10c | 1.398996693 |
| SPBC887.04c | 1.399135625 |
| SPBC12D12.06 | 1.403434678 |
| SPBC887.10 | 1.404121925 |
| SPAC9.13c | 1.406329879 |
| SPAC23H4.12 | 1.407144019 |
| SPBC839.17c | 1.407658828 |
| SPAC25B8.17 | 1.407924754 |
| SPAC16E8.06c | 1.408551311 |
| SPCC1259.07 | 1.410400503 |
| SPBC8D2.17 | 1.415974939 |
| SPAC664.03 | 1.417260933 |
| SPBC530.13 | 1.418029278 |
| SPCC1223.11 | 1.420886407 |
| SPCC794.03 | 1.42106526 |
| SPAC23H3.05c | 1.421640065 |

| | |
|---|---|
| SPAC6F6.03c | 1.422877347 |
| SPAC3H5.09c | 1.425174961 |
| SPBC3H7.15 | 1.429785823 |
| SPCC297.05 | 1.430677012 |
| SPBC18H10.02 | 1.431367256 |
| SPCC320.06 | 1.433822522 |
| SPAC11H11.01 | 1.434784639 |
| SPAC140.01 | 1.435115886 |
| SPAC26A3.06 | 1.440340465 |
| SPBC3E7.01 | 1.440849379 |
| SPCC18.15 | 1.442623414 |
| SPAC24H6.08 | 1.444128044 |
| SPAC16E8.12c | 1.450380666 |
| SPAC2G11.03c | 1.452765322 |
| SPAC222.04c | 1.456400929 |
| SPCC188.08c | 1.45933734 |
| SPCC1672.06c | 1.466886282 |
| SPBC1778.06c | 1.471708683 |
| SPAC458.05 | 1.473446473 |
| SPAC1556.01c | 1.475988722 |
| SPAC17C9.07 | 1.477400584 |
| SPBC25B2.11 | 1.478618913 |
| SPCC895.09c | 1.481649738 |
| SPCC1919.10c | 1.486608087 |
| SPCC594.04c | 1.492641199 |
| SPAC2E1P3.05c | 1.492927302 |
| SPCC16C4.01 | 1.500722543 |
| SPBC2F12.11c | 1.503009259 |
| SPAC11E3.04c | 1.504643776 |
| SPAC25H1.05 | 1.514413902 |
| SPBC16H5.12c | 1.514923916 |
| SPCP31B10.03c | 1.517750553 |
| SPBC83.09c | 1.519293318 |
| SPCC594.05c | 1.535047729 |
| SPAC140.04 | 1.538004308 |
| SPBC646.02 | 1.53990783 |
| SPAC17G6.04c | 1.539991472 |
| SPAC2F7.04 | 1.5474124 |
| SPBC20F10.06 | 1.554548027 |
| SPCC1620.02 | 1.557343844 |
| SPCC1393.13 | 1.558802642 |
| SPAC2E1P5.01c | 1.563623224 |
| SPBC577.12 | 1.566480864 |
| SPAC17A5.08 | 1.568921164 |
| SPCC4G3.11 | 1.571951718 |
| SPAC328.07c | 1.597982125 |
| SPBC32F12.01c | 1.600661063 |
| SPAC1D4.06c | 1.611233715 |

| | |
|---|---|
| SPAC14C4.13 | 1.612390756 |
| SPAC630.05 | 1.615355202 |
| SPBC18E5.04 | 1.628726537 |
| SPBC18H10.13 | 1.630300025 |
| SPAC1851.04c | 1.631670652 |
| SPAC11G7.06c | 1.633932186 |
| SPCC1739.03 | 1.636636842 |
| SPBC29A3.07c | 1.648197786 |
| SPBP22H7.06 | 1.666583543 |
| SPCC1183.10 | 1.672643554 |
| SPBC25B2.01 | 1.681229488 |
| SPBC543.07 | 1.684555991 |
| SPAC17G6.05c | 1.691370772 |
| SPBC11G11.02c | 1.691503846 |
| SPAC688.13 | 1.70015019 |
| SPAC3H1.09c | 1.709087013 |
| SPAC4F10.20 | 1.718391575 |
| SPAC2G11.06 | 1.724330664 |
| SPCC1393.02c | 1.738358247 |
| SPCC830.06 | 1.740336263 |
| SPCC16A11.10c | 1.742236718 |
| SPBC1734.08 | 1.748896209 |
| SPBC13G1.12 | 1.762585452 |
| SPBP8B7.22 | 1.768784085 |
| SPBC1A4.04 | 1.778815489 |
| SPAC30D11.12 | 1.796662404 |
| SPAC20G4.05c | 1.817405084 |
| SPAC31G5.03 | 1.829536949 |
| SPCC1529.01 | 1.852432947 |
| SPCC1919.03c | 1.853544974 |
| SPAC5H10.08c | 1.902697346 |
| SPAC16E8.05c | 1.908099174 |
| SPBC24C6.08c | 1.929744083 |
| SPAC1F8.04c | 1.955352132 |
| SPAC3A11.05c | 1.998438714 |
| SPBC3H7.09 | 2.053909091 |
| SPCC794.08 | 2.061465489 |
| SPBCPT2R1.03 | 2.097858067 |
| SPBC1A4.03c | 2.13353458 |
| SPAC31A2.02 | 2.144574635 |
| SPAC4D7.10c | 2.167724664 |
| SPBC776.09 | 2.241729852 |
| SPBC17D11.08 | 2.309075696 |
| SPAC16A10.05c | 2.338798648 |
| SPBPB2B2.19c | 2.902699838 |
