## Supplemental Table 2 for "Rapalink-1 reveals novel mTOR-dependent genes and an agmatinergic axis-based metabolic feedback regulating mTOR activity and lifespan"

**Supplementary Table 2**

**Rapamycin upregulated genes**

| gene_id | log2FoldChange | pvalue | padj | gene_name |
|---|---|---|---|---|
| SPBPB21E7.07 | 1.327525153 | 2.73149E-61 | 1.28298E-57 | aes1 |
| SPCC70.12c | 1.282382668 | 2.89012E-42 | 6.78745E-39 | ecl1 |
| SPBC26H8.08c | 0.784254782 | 2.48525E-29 | 3.76814E-26 | grn1 |
| SPAC16.05c | 1.00717996 | 3.20898E-29 | 3.76814E-26 | sfp1 |
| SPAC19D5.01 | 1.054885335 | 1.51644E-27 | 1.42454E-24 | pyp2 |
| SPCC737.04 | 1.001584819 | 1.94349E-27 | 1.52143E-24 | SPCC737.04 |
| SPAC19E9.03 | 0.583324554 | 4.00467E-27 | 2.68713E-24 | pas1 |
| SPBC1271.08c | 1.656407901 | 6.11105E-25 | 3.58795E-22 | SPBC1271.08c |
| SPBC725.03 | 0.824593932 | 9.07428E-24 | 4.26219E-21 | SPBC725.03 |
| SPAC4G8.13c | 0.471094719 | 8.78721E-20 | 3.75214E-17 | prz1 |
| SPAC637.03 | 0.543597884 | 9.3465E-19 | 3.65838E-16 | SPAC637.03 |
| SPBC32H8.07 | 0.693232086 | 1.10805E-17 | 4.00346E-15 | git5 |
| SPNCRNA.130 | 2.534904852 | 2.28553E-17 | 7.66794E-15 | omt3 |
| SPAC2E1P5.05 | 0.72123448 | 4.7187E-17 | 1.47758E-14 | rrp9 |
| SPCC794.04c | 0.95048128 | 1.20152E-16 | 3.52723E-14 | SPCC794.04c |
| SPBC1105.14 | 1.526464168 | 1.49781E-16 | 4.13835E-14 | rsv2 |
| SPCPB1C11.02 | 0.679991008 | 3.01186E-16 | 7.85929E-14 | SPCPB1C11.02 |
| SPCC1393.08 | 0.422668241 | 7.10379E-16 | 1.75613E-13 | SPCC1393.08 |
| SPAC140.02 | 0.618487225 | 1.44692E-15 | 3.39809E-13 | gar2 |
| SPBC16D10.08c | 0.50177127 | 5.17449E-15 | 1.15736E-12 | hsp104 |
| SPBC21C3.19 | 0.417427084 | 9.06803E-15 | 1.93602E-12 | SPBC21C3.19 |
| SPCPB16A4.07 | 0.609578607 | 1.20634E-14 | 2.46356E-12 | SPCPB16A4.07 |
| SPAPB1A10.08 | 0.589215115 | 1.5549E-14 | 3.04307E-12 | SPAPB1A10.08 |
| SPBC1711.07 | 0.600833626 | 1.90686E-14 | 3.58262E-12 | rrb1 |
| SPCC757.11c | 0.618880774 | 1.09845E-13 | 1.9844E-11 | SPCC757.11c |
| SPRRNA.45 | 0.694105606 | 1.42703E-13 | 2.4825E-11 | SPRRNA.45 |
| SPCC1235.01 | 0.426535834 | 1.58125E-13 | 2.65255E-11 | SPCC1235.01 |
| SPCC4G3.03 | 0.876996391 | 8.16656E-13 | 1.3227E-10 | SPCC4G3.03 |
| SPCC16A11.15c | 0.639247505 | 1.61074E-12 | 2.52188E-10 | SPCC16A11.15c |
| SPBC21C3.11 | 0.611821287 | 2.49209E-12 | 3.77591E-10 | ubx4 |
| SPAC4F8.04 | 0.75162002 | 2.77935E-12 | 4.07956E-10 | SPAC4F8.04 |

| | | | | |
|---|---|---|---|---|
| SPBP35G2.13c | 0.669163344 | 3.70418E-12 | 5.27229E-10 | swc2 |
| SPBC17D1.06 | 0.504184343 | 4.53067E-12 | 6.25898E-10 | dbp3 |
| SPAC2F3.08 | 0.642033706 | 6.08492E-12 | 8.16596E-10 | sut1 |
| SPBC1347.02 | 0.459216178 | 7.82184E-12 | 1.0044E-09 | fkbp39 |
| SPBC460.04c | 0.749651407 | 1.01863E-11 | 1.25908E-09 | SPBC460.04c |
| SPCC330.03c | 0.839784717 | 1.2424E-11 | 1.4963E-09 | SPCC330.03c |
| SPBC651.01c | 0.417899841 | 2.10345E-11 | 2.46997E-09 | nog1 |
| SPAC869.02c | 0.525883986 | 4.74775E-11 | 5.43907E-09 | SPAC869.02c |
| SPBC56F2.09c | 0.369154355 | 7.92926E-11 | 8.66134E-09 | arg5 |
| SPAPJ691.02 | 0.845740846 | 1.75139E-10 | 1.86961E-08 | SPAPJ691.02 |
| SPCC191.01 | 0.427165361 | 1.91661E-10 | 2.00052E-08 | SPCC191.01 |
| SPAC57A7.05 | 0.493866309 | 2.12754E-10 | 2.1724E-08 | SPAC57A7.05 |
| SPBC13G1.09 | 0.53648369 | 2.50305E-10 | 2.50145E-08 | enp1 |
| SPBC1685.13 | 0.452987604 | 3.46139E-10 | 3.38711E-08 | fhn1 |
| SPBC660.07 | 0.339378578 | 3.90026E-10 | 3.73868E-08 | ntp1 |
| SPAC8C9.03 | 0.666635228 | 4.29719E-10 | 4.03678E-08 | cgs1 |
| SPAC2C4.11c | 0.575272062 | 7.89847E-10 | 7.13444E-08 | rbp28 |
| SPAP8A3.04c | 0.393940865 | 8.84332E-10 | 7.69205E-08 | hsp9 |
| SPAC17H9.04c | 0.395111032 | 1.16104E-09 | 9.91525E-08 | nrp1 |
| SPCC4F11.02 | 0.375731799 | 2.5891E-09 | 2.06119E-07 | ptc1 |
| SPCC132.04c | 0.551973711 | 2.95969E-09 | 2.27896E-07 | gdh2 |
| SPAC22F8.05 | 0.320544038 | 3.11698E-09 | 2.34154E-07 | SPAC22F8.05 |
| SPAC19G12.09 | 0.359899884 | 3.14066E-09 | 2.34154E-07 | SPAC19G12.09 |
| SPBC11C11.06c | 0.345777975 | 3.51614E-09 | 2.58052E-07 | SPBC11C11.06c |
| SPCC18B5.01c | 0.404885454 | 3.65784E-09 | 2.64321E-07 | bfr1 |
| SPAC6C3.04 | 0.350199927 | 4.0918E-09 | 2.912E-07 | cit1 |
| SPBP35G2.16c | 0.419685589 | 5.07932E-09 | 3.56083E-07 | ecl2 |
| SPAC926.08c | 0.478873475 | 7.24074E-09 | 5.00144E-07 | rpf2 |
| SPCC16A11.12c | 0.463719419 | 9.20889E-09 | 6.26872E-07 | ubp1 |
| SPAC22H10.13 | 0.518821852 | 1.26825E-08 | 8.27358E-07 | zym1 |
| SPAC27E2.08 | 0.64175203 | 2.60524E-08 | 1.63157E-06 | Tf2-6 |
| SPBP35G2.05c | 0.379586329 | 2.67012E-08 | 1.6502E-06 | cki2 |
| SPCC965.07c | 0.336895072 | 2.92024E-08 | 1.78135E-06 | gst2 |

| | | | | |
|---|---|---|---|---|
| SPCC306.11 | 0.322387871 | 3.00181E-08 | 1.80763E-06 | SPCC306.11 |
| SPAC821.10c | 0.293080643 | 3.07105E-08 | 1.82591E-06 | sod1 |
| SPBPB21E7.04c | 1.554025534 | 3.1852E-08 | 1.87011E-06 | SPBPB21E7.04c |
| SPAC11D3.08c | 0.431019314 | 3.33032E-08 | 1.93117E-06 | SPAC11D3.08c |
| SPCC70.10 | 0.536379203 | 3.98199E-08 | 2.25342E-06 | SPCC70.10 |
| SPBPB21E7.09 | 0.494440071 | 4.28575E-08 | 2.36825E-06 | SPBPB21E7.09 |
| SPCPB16A4.06c | 0.847141126 | 5.00029E-08 | 2.73097E-06 | SPCPB16A4.06c |
| SPAC10F6.16 | 0.371854748 | 5.26841E-08 | 2.84434E-06 | mug134 |
| SPAC16E8.06c | 0.545626884 | 6.31302E-08 | 3.34103E-06 | nop12 |
| SPBC651.08c | 0.417560743 | 6.33068E-08 | 3.34103E-06 | rpc1 |
| SPCC16A11.01 | 0.347550129 | 7.02611E-08 | 3.58714E-06 | sfk1 |
| SPCC320.11c | 0.476811173 | 7.97413E-08 | 4.02737E-06 | nip7 |
| SPAC1486.09 | 0.475793433 | 1.09414E-07 | 5.46723E-06 | nob1 |
| SPAC26H5.09c | 0.355131732 | 1.23518E-07 | 6.107E-06 | SPAC26H5.09c |
| SPAC32A11.02c | 0.351655759 | 1.2878E-07 | 6.23587E-06 | SPAC32A11.02c |
| SPAC26F1.10c | 0.380283438 | 1.38546E-07 | 6.64033E-06 | pyp1 |
| SPAC13G7.05 | 0.390341296 | 1.5501E-07 | 7.35436E-06 | are1 |
| SPAC2C4.17c | 0.39863063 | 1.68676E-07 | 7.92269E-06 | msy2 |
| SPAC222.06 | 0.504231379 | 1.84609E-07 | 8.58521E-06 | mak16 |
| SPAC1D4.11c | 0.431500254 | 1.9074E-07 | 8.78337E-06 | lkh1 |
| SPBPB10D8.06c | 0.43519602 | 2.36881E-07 | 1.08022E-05 | SPBPB10D8.06c |
| SPAC3H8.04 | 0.413804653 | 2.80166E-07 | 1.24895E-05 | SPAC3H8.04 |
| SPBC8E4.04 | 0.347823514 | 2.81857E-07 | 1.24895E-05 | SPBC8E4.04 |
| SPBC31E1.06 | 0.323214977 | 2.96525E-07 | 1.30166E-05 | bms1 |
| SPBC1703.05 | 0.466144478 | 3.05976E-07 | 1.33071E-05 | rio2 |
| SPBC83.15 | 0.479899616 | 4.13961E-07 | 1.76761E-05 | SPBC83.15 |
| SPBC19C2.11c | 0.350823977 | 4.24696E-07 | 1.79712E-05 | mdm34 |
| SPBC1685.14c | 0.430473991 | 4.66258E-07 | 1.95537E-05 | SPBC1685.14c |
| SPBC56F2.15 | 0.426082823 | 5.31142E-07 | 2.1884E-05 | tam13 |
| SPAC1F7.02c | 0.418301878 | 6.51653E-07 | 2.6286E-05 | has1 |
| SPBC336.02 | 0.587000816 | 6.67809E-07 | 2.65822E-05 | SPBC336.02 |
| SPAC1751.01c | 0.423598016 | 7.52821E-07 | 2.97143E-05 | gti1 |
| SPBC14C8.02 | 0.399468823 | 9.15414E-07 | 3.52434E-05 | tim44 |

| | | | | |
|---|---|---|---|---|
| SPBC32H8.05 | 0.389799263 | 9.97103E-07 | 3.77693E-05 | SPBC32H8.05 |
| SPCC24B10.18 | 0.592706426 | 1.03798E-06 | 3.86935E-05 | SPCC24B10.18 |
| SPBC18H10.20c | 0.33463371 | 1.06021E-06 | 3.92111E-05 | any1 |
| SPCC4B3.12 | 0.35065386 | 1.18963E-06 | 4.29822E-05 | set9 |
| SPACUNK4.16c | 0.285933962 | 1.22693E-06 | 4.39915E-05 | SPACUNK4.16c |
| SPAC20G8.09c | 0.413871673 | 1.33501E-06 | 4.75042E-05 | nat10 |
| SPAC821.04c | 0.354368003 | 1.44807E-06 | 5.04956E-05 | cid13 |
| SPBC16E9.16c | 0.383569267 | 1.55994E-06 | 5.30944E-05 | lsd90 |
| SPAC24C9.08 | 0.277774686 | 1.63227E-06 | 5.47824E-05 | SPAC24C9.08 |
| SPAC24B11.09 | 0.38863381 | 1.63286E-06 | 5.47824E-05 | mpc2 |
| SPBC947.07 | 0.595039782 | 1.87205E-06 | 6.14897E-05 | SPBC947.07 |
| SPAC977.10 | 0.334090744 | 1.89699E-06 | 6.18762E-05 | sod2 |
| SPNCRNA.1115 | 0.332434568 | 2.64038E-06 | 8.26792E-05 | SPNCRNA.1115 |
| SPCC338.12 | 0.371879217 | 2.9326E-06 | 9.12213E-05 | pbi2 |
| SPBC660.06 | 0.309551875 | 3.3634E-06 | 0.000103933 | SPBC660.06 |
| SPAPB18E9.02c | 0.349727481 | 3.51916E-06 | 0.000108036 | ppk18 |
| SPCC737.03c | 0.320357575 | 4.52823E-06 | 0.00013703 | ima1 |
| SPBC1683.07 | 0.568907478 | 4.54341E-06 | 0.00013703 | mal1 |
| SPBC8E4.03 | 0.353443546 | 4.5738E-06 | 0.00013703 | SPBC8E4.03 |
| SPAC688.04c | 0.34826854 | 4.5803E-06 | 0.00013703 | gst3 |
| SPCC320.08 | 0.46266315 | 4.95235E-06 | 0.000147223 | SPCC320.08 |
| SPAC4F10.09c | 0.338788728 | 5.31642E-06 | 0.000157052 | noc1 |
| SPCC2H8.02 | 0.317914026 | 6.47451E-06 | 0.000187721 | SPCC2H8.02 |
| SPBC359.06 | 0.77050885 | 6.91372E-06 | 0.000199225 | mug14 |
| SPCC645.03c | 0.283380692 | 7.2354E-06 | 0.000207129 | isa1 |
| SPCC1827.01c | 0.420649555 | 7.2762E-06 | 0.000207129 | utp25 |
| SPCC895.06 | 0.369212847 | 7.88214E-06 | 0.000223027 | elp2 |
| SPAC24B11.05 | 0.368400463 | 8.24204E-06 | 0.000231813 | SPAC24B11.05 |
| SPAC328.09 | 0.318740749 | 8.8676E-06 | 0.000246456 | SPAC328.09 |
| SPCC1442.04c | 0.350969658 | 1.00545E-05 | 0.000276177 | SPCC1442.04c |
| SPBC12C2.12c | 0.27215759 | 1.03347E-05 | 0.000282221 | glo1 |
| SPCC569.05c | 0.289095448 | 1.06299E-05 | 0.000288605 | SPCC569.05c |
| SPBC106.02c | 0.906169362 | 1.08151E-05 | 0.000291673 | srx1 |

| | | | | |
|---|---|---|---|---|
| SPBC1539.10 | 0.488054444 | 1.08671E-05 | 0.000291673 | nop16 |
| SPAC1687.14c | 0.466444035 | 1.10825E-05 | 0.000295765 | SPAC1687.14c |
| SPBC1A4.07c | 0.398942003 | 1.17321E-05 | 0.000307852 | sof1 |
| SPBC15C4.06c | 0.425759107 | 1.21146E-05 | 0.000316123 | SPBC15C4.06c |
| SPCC1393.14 | 0.977085707 | 1.23428E-05 | 0.0003203 | ten1 |
| SPBC1289.06c | 0.275428859 | 1.42777E-05 | 0.000364469 | ppr8 |
| SPNCRNA.1560 | 0.677812338 | 1.48738E-05 | 0.000377634 | SPNCRNA.1560 |
| SPBC1347.11 | 0.431657205 | 1.55147E-05 | 0.000391787 | sro1 |
| SPBC119.03 | 0.305885641 | 1.62447E-05 | 0.00040803 | SPBC119.03 |
| SPAC1B3.06c | 0.313712966 | 1.65617E-05 | 0.000413779 | SPAC1B3.06c |
| SPAC29A4.17c | 0.606758121 | 1.75139E-05 | 0.000435253 | SPAC29A4.17c |
| SPBC17D11.08 | 0.293106759 | 1.96868E-05 | 0.00048161 | SPBC17D11.08 |
| SPBC2G2.04c | 0.289471083 | 2.06104E-05 | 0.000496446 | mmf1 |
| SPBC19F5.05c | 0.347599446 | 2.08003E-05 | 0.000498465 | ppp1 |
| SPAC1687.16c | 0.305873417 | 2.17486E-05 | 0.000518265 | erg31 |
| SPAPB8E5.07c | 0.265906086 | 2.41549E-05 | 0.000569596 | rrp12 |
| SPAC22H10.07 | 0.377089425 | 2.64742E-05 | 0.000618654 | scd2 |
| SPAC16C9.03 | 0.361340828 | 2.80094E-05 | 0.000649461 | nmd3 |
| SPAC27D7.03c | 0.829107477 | 2.80691E-05 | 0.000649461 | mei2 |
| SPCC330.09 | 0.367852193 | 2.92236E-05 | 0.000667586 | enp2 |
| SPAC26F1.11 | 0.839625587 | 2.94168E-05 | 0.000667586 | SPAC26F1.11 |
| SPCC1183.07 | 0.271030543 | 2.95816E-05 | 0.000668003 | rrp5 |
| SPBC21D10.09c | 0.383278868 | 3.06197E-05 | 0.000684861 | rkr1 |
| SPBPJ4664.02 | 0.356548793 | 3.63286E-05 | 0.000797362 | SPBPJ4664.02 |
| SPCC576.04 | 0.304111995 | 3.73653E-05 | 0.000815853 | bxi1 |
| SPNCRNA.111 | 0.307828279 | 3.83023E-05 | 0.000825414 | SPNCRNA.111 |
| SPAC8F11.04 | 0.317916442 | 4.23991E-05 | 0.000893042 | SPAC8F11.04 |
| SPCC290.02 | 0.334565452 | 4.47282E-05 | 0.000929594 | rpc34 |
| SPAC3A11.10c | 0.286143681 | 4.9898E-05 | 0.001016105 | SPAC3A11.10c |
| SPAC869.08 | 1.239773557 | 5.16846E-05 | 0.001046391 | pcm2 |
| SPBP22H7.02c | 0.368269939 | 5.26825E-05 | 0.001062016 | mrd1 |
| SPBC19C7.04c | 1.272428258 | 5.30228E-05 | 0.001064308 | SPBC19C7.04c |
| SPCP1E11.08 | 0.311630314 | 5.44993E-05 | 0.001080098 | nsa2 |

| | | | | |
|---|---|---|---|---|
| SPBC21B10.15 | 0.449390392 | 5.82186E-05 | 0.001144154 | SPBC21B10.15 |
| SPBC365.12c | 0.29692363 | 6.29251E-05 | 0.001218036 | ish1 |
| SPBPB2B2.06c | 0.995180041 | 6.33015E-05 | 0.001218555 | SPBPB2B2.06c |
| SPAC23H4.15 | 0.300467788 | 6.92039E-05 | 0.001310689 | tsr1 |
| SPBP4G3.02 | 0.594690897 | 7.5662E-05 | 0.001410256 | pho1 |
| SPAC20G8.10c | 0.448252188 | 7.72951E-05 | 0.001429916 | atg6 |
| SPAC22E12.13c | 0.269923554 | 7.73257E-05 | 0.001429916 | rlp24 |
| SPAC4H3.03c | 0.379204422 | 8.10257E-05 | 0.001475107 | SPAC4H3.03c |
| SPAC19A8.07c | 0.4385688 | 8.15525E-05 | 0.001478966 | imp4 |
| SPAC20G4.03c | 0.328292992 | 9.12246E-05 | 0.001621003 | hri1 |
| SPCC1020.10 | 0.287766885 | 9.21456E-05 | 0.001621003 | oca2 |
| SPNCRNA.304 | 0.555027064 | 0.000105292 | 0.001772597 | SPNCRNA.304 |
| SPBPB10D8.05c | 0.607279874 | 0.000122161 | 0.002006262 | SPBPB10D8.05c |
| SPBC20F10.03 | 0.283897233 | 0.000126221 | 0.002058546 | SPBC20F10.03 |
| SPCC1753.02c | 0.325040544 | 0.000135749 | 0.002206268 | git3 |
| SPAC3C7.05c | 0.313771034 | 0.000139256 | 0.002247721 | mug191 |
| SPAC1039.11c | 0.323851064 | 0.000146158 | 0.002335045 | gto1 |
| SPNCRNA.975 | 0.412159987 | 0.000150717 | 0.002376514 | SPNCRNA.975 |
| SPBC106.14c | 0.968202861 | 0.000157149 | 0.002452255 | sda1 |
| SPCC663.17 | 0.549808229 | 0.000171393 | 0.002630821 | wtf15 |
| SPBC244.02c | 0.330670985 | 0.000180283 | 0.002740414 | utp6 |
| SPAC823.08c | 0.312026249 | 0.000183378 | 0.002778478 | rrp3 |
| SPAC664.08c | 0.378390494 | 0.000193125 | 0.002892719 | bfr2 |
| SPAC4G8.06c | 0.283075766 | 0.000211447 | 0.003103646 | trm12 |
| SPBC29A3.06 | 0.315117826 | 0.000219986 | 0.003208926 | utp18 |
| SPNCRNA.103 | 1.266076958 | 0.000225767 | 0.003261971 | sme2 |
| SPAC18G6.02c | 0.330041303 | 0.0002264 | 0.003261971 | chp1 |
| SPBC29A3.08 | 0.44985286 | 0.000230331 | 0.003298374 | pof4 |
| SPAC513.03 | 0.455060707 | 0.000238137 | 0.003365583 | mfm2 |
| SPAC1039.06 | 0.308347279 | 0.000245266 | 0.003418447 | SPAC1039.06 |
| SPAPB24D3.04c | 0.704169575 | 0.0002537 | 0.003515136 | mag1 |
| SPBC26H8.13c | 0.347458007 | 0.000390291 | 0.005135 | SPBC26H8.13c |
| SPAC823.16c | 0.364045688 | 0.000427924 | 0.005521871 | atg1802 |

| | | | | |
|---|---|---|---|---|
| SPNCRNA.863 | 0.823396361 | 0.00044088 | 0.005627208 | SPNCRNA.863 |
| SPBC19C2.05 | 0.368488905 | 0.000448106 | 0.005673193 | pat1 |
| SPNCRNA.1157 | 0.611973745 | 0.000455177 | 0.005747222 | SPNCRNA.1157 |
| SPBC32C12.02 | 0.763949222 | 0.000460719 | 0.005801599 | ste11 |
| SPBC1706.01 | 0.272265684 | 0.000465734 | 0.005833479 | tea4 |
| SPCC553.12c | 0.277606075 | 0.000473863 | 0.005870491 | SPCC553.12c |
| SPBC19G7.13 | 0.33464974 | 0.000473952 | 0.005870491 | tbf1 |
| SPCC285.09c | 0.286108699 | 0.000474939 | 0.005870491 | cgs2 |
| SPBC3F6.04c | 0.271675952 | 0.000556345 | 0.006683253 | nop14 |
| SPAC27D7.09c | 0.26592486 | 0.000561849 | 0.006732157 | SPAC27D7.09c |
| SPAC3C7.13c | 0.291083549 | 0.000564639 | 0.006748374 | SPAC3C7.13c |
| SPAC27F1.06c | 0.278626764 | 0.000611131 | 0.007158312 | SPAC27F1.06c |
| SPAC212.06c | 1.354057675 | 0.000619553 | 0.007203065 | SPAC212.06c |
| SPBCPT2R1.04c | 0.508746883 | 0.000628827 | 0.007274873 | SPBCPT2R1.04c |
| SPBPB10D8.07c | 0.378609836 | 0.000673816 | 0.007626299 | SPBPB10D8.07c |
| SPAC30D11.03 | 0.30524997 | 0.000871881 | 0.009436004 | ddx27 |
| SPBC21H7.04 | 0.368032644 | 0.000876185 | 0.009460779 | dbp7 |
| SPAC32A11.01 | 0.442672628 | 0.000900671 | 0.009571153 | mug8 |
| SPCC320.05 | 0.274985661 | 0.000907223 | 0.009604961 | SPCC320.05 |
| SPAC19A8.05c | 0.264263536 | 0.000907942 | 0.009604961 | sst4 |
| SPAC31G5.04 | 0.300455685 | 0.000920059 | 0.00964624 | lys12 |
| SPAC9.04 | 0.510564068 | 0.001223558 | 0.012253842 | Tf2-1 |
| SPBP4H10.12 | 0.27290429 | 0.001252744 | 0.012440039 | SPBP4H10.12 |
| SPAC25A8.02 | 0.285028941 | 0.001269746 | 0.012582276 | atg14 |
| SPBC16C6.12c | 0.352470786 | 0.001321714 | 0.012975023 | las1 |
| SPBC1348.12 | 0.524358752 | 0.00140702 | 0.013711144 | SPBC1348.12 |
| SPAC4G9.22 | 0.755582366 | 0.001426558 | 0.013844094 | SPAC4G9.22 |
| SPAC22G7.08 | 0.313290827 | 0.001460241 | 0.014083682 | ppk8 |
| SPCC1906.04 | 0.706001132 | 0.001580309 | 0.015031451 | wtf20 |
| SPBC24C6.02 | 0.319740617 | 0.0016407 | 0.015568423 | SPBC24C6.02 |
| SPAC22F8.02c | 0.294387927 | 0.00167064 | 0.015757017 | pvg5 |
| SPNCRNA.58 | 0.427111507 | 0.001708731 | 0.015987868 | prl58 |
| SPBC947.13 | 0.354115971 | 0.001907272 | 0.017369846 | rba50 |

| | | | | |
|---|---|---|---|---|
| SPBC19G7.06 | 0.341399839 | 0.001956565 | 0.017615641 | mbx1 |
| SPCC1739.08c | 0.852754721 | 0.00215714 | 0.019189556 | SPCC1739.08c |
| SPBPB21E7.11 | 1.273857949 | 0.00216905 | 0.019259033 | SPBPB21E7.11 |
| SPAC890.05 | 0.608610466 | 0.00227193 | 0.019835044 | SPAC890.05 |
| SPCC364.01 | 0.333376014 | 0.002368014 | 0.020437278 | cif1 |
| SPCC126.07c | 0.29275227 | 0.002380941 | 0.020438224 | pbr1 |
| SPAC750.01 | 0.516156569 | 0.002412475 | 0.020581951 | SPAC750.01 |
| SPAC1687.07 | 0.300085783 | 0.002430583 | 0.02064457 | SPAC1687.07 |
| SPAC9E9.09c | 0.629085148 | 0.002466333 | 0.020872728 | atd1 |
| SPAC11D3.16c | 0.297926645 | 0.002618461 | 0.02176798 | SPAC11D3.16c |
| SPNCRNA.99 | 0.779660966 | 0.002729433 | 0.022491487 | SPNCRNA.99 |
| SPBC19C2.04c | 0.330462656 | 0.002769394 | 0.022740982 | ubp11 |
| SPAC212.04c | 0.450336684 | 0.00284057 | 0.023203748 | SPAC212.04c |
| SPBC21.07c | 0.382901623 | 0.002942254 | 0.023745306 | ppk24 |
| SPBC2D10.19c | 0.449972618 | 0.003133943 | 0.024865084 | alb1 |
| SPAC11D3.01c | 0.433038744 | 0.003380038 | 0.026460066 | SPAC11D3.01c |
| SPBC1289.15 | 0.376246932 | 0.004030973 | 0.030439678 | pfl5 |
| SPCC550.05 | 0.369508978 | 0.004190264 | 0.031505341 | nse1 |
| SPNCRNA.1133 | 0.425703374 | 0.004239901 | 0.031762067 | SPNCRNA.1133 |
| SPBC460.05 | 0.289754855 | 0.004275537 | 0.031876505 | SPBC460.05 |
| SPAC15E1.02c | 0.324618057 | 0.004299236 | 0.031962439 | SPAC15E1.02c |
| SPAPB24D3.10c | 0.302874998 | 0.004307478 | 0.031962439 | agl1 |
| SPAC31G5.12c | 0.309948631 | 0.004354271 | 0.032157251 | maf1 |
| SPBC18H10.05 | 0.270715187 | 0.004424621 | 0.032574361 | SPBC18H10.05 |
| SPAC1565.04c | 0.392627927 | 0.004781445 | 0.034711665 | ste4 |
| SPCC1620.02 | 0.552141245 | 0.004860926 | 0.034964421 | wtf23 |
| SPNCRNA.1095 | 0.530075307 | 0.004942852 | 0.035399763 | SPNCRNA.1095 |
| SPAC5H10.07 | 0.533725281 | 0.005059148 | 0.036004273 | SPAC5H10.07 |
| SPNCRNA.608 | 0.292656898 | 0.00519873 | 0.036830221 | SPNCRNA.608 |
| SPBC1271.09 | 0.438342971 | 0.005244897 | 0.037101324 | tgp1 |
| SPNCRNA.1668 | 1.192609951 | 0.005266353 | 0.037197079 | mug147-antisense-1 |
| SPCC970.11c | 0.411888336 | 0.00534322 | 0.037514358 | wtf9 |
| SPAC11D3.05 | 0.347933231 | 0.005369606 | 0.037643343 | mfs2 |

| | | | | |
|---|---|---|---|---|
| SPBC24C6.06 | 0.321218303 | 0.005447182 | 0.037960556 | gpa1 |
| SPNCRNA.1350 | 0.914217483 | 0.005543728 | 0.038519066 | SPNCRNA.1350 |
| SPBC1773.17c | 0.277803621 | 0.005661004 | 0.038974522 | SPBC1773.17c |
| SPBC660.05 | 0.459912368 | 0.00571255 | 0.039170581 | SPBC660.05 |
| SPAC19B12.08 | 0.27781102 | 0.00586234 | 0.040080655 | atg4 |
| SPBC215.08c | 0.873592286 | 0.005881766 | 0.040155019 | arg4 |
| SPBC1348.14c | 0.855005403 | 0.006048546 | 0.041054945 | ght7 |
| SPAC18B11.06 | 0.282855655 | 0.006190139 | 0.041830562 | lcp5 |
| SPCC553.05c | 0.601096855 | 0.006303788 | 0.04235893 | wtf6 |
| SPAC1556.01c | 0.352920717 | 0.006323077 | 0.042427844 | rad50 |
| SPNCRNA.1691 | 1.021218345 | 0.00681654 | 0.044905032 | SPNCRNA.1691 |
| SPAC167.08 | 1.007997116 | 0.007314324 | 0.047126721 | Tf2-2 |
| SPAC2H10.01 | 0.377872005 | 0.007566458 | 0.048419145 | SPAC2H10.01 |
| SPBC1683.08 | 0.266430404 | 0.007619783 | 0.048694039 | ght4 |
| SPAC4G9.19 | 0.312899492 | 0.007699872 | 0.04893951 | SPAC4G9.19 |
