## Supplemental Table 3 for "Rapalink-1 reveals novel mTOR-dependent genes and an agmatinergic axis-based metabolic feedback regulating mTOR activity and lifespan"

**Supplementary Table 3**
**Rapamycin downregulated genes**

| gene_id | log2FoldChange | pvalue | padj | gene_name |
|---|---|---|---|---|
| SPAC25B8.13c | -1.195695258 | 0.004628733 | 0.033812068 | isp7 |
| SPNCRNA.228 | -1.029389361 | 0.007149565 | 0.046255522 | SPNCRNA.228 |
| SPAC1399.04c | -0.837399055 | 0.000503856 | 0.006163057 | SPAC1399.04c |
| SPBC13A2.04c | -0.680948636 | 4.62409E-05 | 0.0009568 | ptr2 |
| SPNCRNA.630 | -0.572233289 | 0.00041732 | 0.005429785 | SPNCRNA.630 |
| SPAC23H4.06 | -0.553422289 | 2.33162E-24 | 1.21685E-21 | gln1 |
| SPBC36.01c | -0.482113128 | 1.12863E-08 | 7.46647E-07 | SPBC36.01c |
| SPAC13G7.13c | -0.439566924 | 1.25471E-07 | 6.13894E-06 | msa1 |
| SPAC821.08c | -0.438519212 | 5.58508E-06 | 0.000162939 | slp1 |
| SPBC23E6.10c | -0.415932674 | 1.94401E-08 | 1.25082E-06 | mri1 |
| SPBC36.03c | -0.415622587 | 8.56674E-10 | 7.59207E-08 | mfs3 |
| SPBP22H7.07 | -0.415265194 | 2.92453E-09 | 2.27896E-07 | prp5 |
| SPBP8B7.05c | -0.41237306 | 2.05769E-09 | 1.66638E-07 | nce103 |
| SPCC584.13 | -0.400966931 | 8.81629E-05 | 0.001590847 | SPCC584.13 |
| SPAC17A2.01 | -0.399538156 | 2.53317E-07 | 1.14407E-05 | bsu1 |
| SPAC23G3.04 | -0.399358688 | 0.00121374 | 0.012195543 | ies4 |
| SPAPB2B4.03 | -0.392999679 | 0.000913048 | 0.009615662 | cig2 |
| SPBPB7E8.01 | -0.38164611 | 7.91202E-12 | 1.0044E-09 | SPBPB7E8.01 |
| SPBC1198.02 | -0.37948153 | 5.43204E-10 | 5.0028E-08 | dea2 |
| SPCC1739.13 | -0.365677711 | 5.97756E-11 | 6.68491E-09 | ssa2 |
| SPBC1685.09 | -0.353681161 | 0.000104818 | 0.001770978 | rps29 |
| SPAC2F7.14c | -0.350991993 | 3.91177E-05 | 0.000835163 | rrp4 |
| SPAC14C4.12c | -0.3496561 | 0.005596634 | 0.038829233 | laf1 |
| SPAC23G3.12c | -0.348564697 | 1.23086E-09 | 1.03238E-07 | SPAC23G3.12c |
| SPAC513.07 | -0.343032126 | 2.30251E-06 | 7.40745E-05 | SPAC513.07 |
| SPBC8D2.16c | -0.342408566 | 0.001247575 | 0.01241496 | SPBC8D2.16c |
| SPBC29B5.02c | -0.341846178 | 0.000812057 | 0.008870309 | isp4 |
| SPBC13E7.09 | -0.337664927 | 0.004824552 | 0.034955047 | vrp1 |
| SPCC4G3.17 | -0.335814622 | 1.38105E-06 | 4.87728E-05 | hdd1 |
| SPBC359.03c | -0.334838625 | 1.98968E-05 | 0.000481728 | aat1 |
| SPCC1223.05c | -0.334043289 | 1.12364E-05 | 0.000298176 | rpl3702 |
| SPAC110.01 | -0.333917008 | 3.3985E-05 | 0.000756529 | ppk1 |
| SPAP7G5.06 | -0.333356837 | 0.000883922 | 0.009478958 | per1 |
| SPBC2F12.08c | -0.332734334 | 0.002920594 | 0.023651777 | ceg1 |
| SPBC2D10.12 | -0.328188431 | 4.15704E-08 | 2.32448E-06 | rhp23 |
| SPCC330.06c | -0.327290159 | 9.78797E-07 | 3.73773E-05 | pmp20 |
| SPCC306.03c | -0.325153211 | 9.49322E-05 | 0.001647974 | cnd2 |
| SPAP11E10.01 | -0.322098471 | 0.000150777 | 0.002376514 | SPAP11E10.01 |
| SPCC1259.04 | -0.320621827 | 0.000640486 | 0.007324917 | iec3 |
| SPBC23G7.08c | -0.317444217 | 3.53775E-05 | 0.000780133 | rga7 |
| SPAC9E9.13 | -0.314379276 | 1.4535E-09 | 1.19773E-07 | wos2 |
| SPAC13F5.05 | -0.314230758 | 0.002845602 | 0.0232045 | mpd1 |
| SPCC338.07c | -0.309776448 | 6.60293E-08 | 3.40813E-06 | naa15 |
| SPAC3G9.09c | -0.304673168 | 8.97614E-07 | 3.48592E-05 | tif211 |
| SPBC1A4.03c | -0.304482154 | 1.25817E-05 | 0.000324706 | top2 |
| SPBC2D10.04 | -0.303787705 | 2.37549E-06 | 7.54802E-05 | SPBC2D10.04 |
| SPCC162.11c | -0.30367296 | 6.50266E-05 | 0.001236559 | SPCC162.11c |

| | | | | |
|---|---|---|---|---|
| SPAC1039.01 | -0.303015736 | 0.000796428 | 0.008760704 | SPAC1039.01 |
| SPAC1834.05 | -0.30081086 | 6.37678E-07 | 2.6045E-05 | alg9 |
| SPBC651.11c | -0.299321549 | 0.000200916 | 0.002976985 | apm3 |
| SPBC1734.02c | -0.297999082 | 0.002298495 | 0.019955694 | cdc27 |
| SPCC338.17c | -0.295516251 | 6.20886E-05 | 0.001210084 | rad21 |
| SPBC543.08 | -0.295255792 | 1.78077E-05 | 0.000440225 | SPBC543.08 |
| SPAC5H10.12c | -0.292065166 | 0.006545728 | 0.043548563 | SPAC5H10.12c |
| SPBP22H7.06 | -0.291075214 | 0.000241899 | 0.0034018 | SPBP22H7.06 |
| SPAC3F10.18c | -0.28999642 | 0.002396499 | 0.020503378 | rpl4102 |
| SPBP4H10.11c | -0.289678189 | 2.4866E-06 | 7.83863E-05 | lcf2 |
| SPAC18G6.07c | -0.287491972 | 1.45133E-06 | 5.04956E-05 | mra1 |
| SPCC553.09c | -0.287396918 | 0.000147626 | 0.002349644 | spb70 |
| SPAC15E1.08 | -0.287314705 | 8.88916E-05 | 0.001593603 | naa10 |
| SPCC645.08c | -0.287279716 | 6.40395E-08 | 3.34215E-06 | snd1 |
| SPCC1494.03 | -0.285464183 | 0.007112948 | 0.046209565 | arz1 |
| SPAC1002.02 | -0.284248906 | 0.000462366 | 0.005806769 | pom34 |
| SPAC1093.01 | -0.281170106 | 2.42536E-05 | 0.000569596 | ppr5 |
| SPAC12B10.03 | -0.280472028 | 0.001696812 | 0.015908035 | bun62 |
| SPAC22F8.10c | -0.27937113 | 0.000250026 | 0.003474471 | sap145 |
| SPAC4G9.02 | -0.277194134 | 0.007766772 | 0.049182347 | rnh201 |
| SPCC4G3.14 | -0.276636579 | 0.000143322 | 0.002297554 | mdj1 |
| SPBC83.17 | -0.276475267 | 1.1858E-06 | 4.29822E-05 | SPBC83.17 |
| SPCC1281.06c | -0.276328509 | 1.03724E-08 | 6.95986E-07 | SPCC1281.06c |
| SPCC830.03 | -0.275177121 | 0.000671019 | 0.00761299 | grc3 |
| SPBC1604.03c | -0.274734223 | 0.005850059 | 0.040054994 | SPBC1604.03c |
| SPCC18.07 | -0.274694119 | 0.000287619 | 0.003927171 | rpc53 |
| SPBC25B2.06c | -0.272094839 | 0.000185208 | 0.002797179 | btb2 |
| SPAC4D7.04c | -0.272020241 | 0.000197545 | 0.002936287 | rer2 |
| SPBC30B4.05 | -0.271507789 | 1.87033E-05 | 0.000459945 | kap109 |
| SPAC926.04c | -0.271415984 | 3.73104E-08 | 2.13716E-06 | hsp90 |
| SPAC25G10.06 | -0.270383635 | 0.000162903 | 0.002533633 | rps2801 |
| SPBC646.04 | -0.269713041 | 9.28269E-05 | 0.001622979 | pla1 |
| SPAC110.04c | -0.268951685 | 2.33681E-08 | 1.48324E-06 | pss1 |
| SPBC1773.07c | -0.26844288 | 6.54772E-07 | 2.6286E-05 | sbp1 |
| SPBC1A4.05 | -0.268035825 | 0.000613928 | 0.007173179 | blt1 |
| SPCC1919.09 | -0.267759399 | 1.48547E-06 | 5.13032E-05 | tif6 |
| SPCC576.11 | -0.265592848 | 7.42108E-05 | 0.001388718 | rpl15 |
| SPBP35G2.08c | -0.263984241 | 0.00334294 | 0.026213338 | air1 |
| SPCC74.04 | -0.263936882 | 0.007216672 | 0.046561414 | SPCC74.04 |
| SPAC5D6.12 | -0.263910796 | 0.000148072 | 0.002349644 | mtf2 |
