## Supplemental Table 4 for "Rapalink-1 reveals novel mTOR-dependent genes and an agmatinergic axis-based metabolic feedback regulating mTOR activity and lifespan"

**Supplementary Table 4**

**Rapalink-1 upregulated genes**

| gene_id | log2FoldChange | pvalue | padj | gene_name |
|---|---|---|---|---|
| SPAC869.04 | 8.003174332 | 9.32166E-33 | 4.49169E-31 | SPAC869.04 |
| SPNCRNA.935 | 5.480580142 | 5.21351E-60 | 5.92254E-58 | SPNCRNA.935 |
| SPNCRNA.130 | 5.301437738 | 2.3814E-119 | 8.116E-117 | omt3 |
| SPAC869.03c | 5.252526189 | 3.62263E-34 | 1.90916E-32 | SPAC869.03c |
| SPAC1039.09 | 4.845567221 | 1.0344E-161 | 1.0576E-158 | isp5 |
| SPBPB2B2.01 | 4.601872136 | 7.12405E-57 | 7.43228E-55 | SPBPB2B2.01 |
| SPNCRNA.603 | 4.192743094 | 1.20715E-05 | 7.91148E-05 | SPNCRNA.603 |
| SPCC576.01c | 4.174447772 | 7.61301E-78 | 1.34199E-75 | xan1 |
| SPAC1039.10 | 4.123775353 | 3.4596E-152 | 2.2107E-149 | mmf2 |
| SPAPB8E5.05 | 4.078239603 | 5.4263E-58 | 6.03027E-56 | mfm1 |
| SPNCRNA.577 | 3.989423602 | 0.000549768 | 0.002590244 | SPBC1652.02-antisense-1 |
| SPAC922.03 | 3.967738811 | 0 | 0 | SPAC922.03 |
| SPBC29B5.02c | 3.863519185 | 3.74952E-85 | 8.71252E-83 | isp4 |
| SPAC1002.17c | 3.674416033 | 5.79155E-41 | 4.22949E-39 | urg2 |
| SPAC1002.19 | 3.655343929 | 2.34911E-23 | 8.05949E-22 | urg1 |
| SPAC27D7.03c | 3.573325442 | 9.5298E-150 | 5.4129E-147 | mei2 |
| SPNCRNA.860 | 3.273698336 | 2.55248E-05 | 0.000155337 | SPNCRNA.860 |
| SPNCRNA.103 | 3.021400134 | 1.23242E-28 | 5.08074E-27 | sme2 |
| SPAC1039.02 | 3.001842803 | 3.71505E-40 | 2.60155E-38 | SPAC1039.02 |
| SPAC1039.07c | 2.994096063 | 5.19922E-90 | 1.26564E-87 | SPAC1039.07c |
| SPAC1039.01 | 2.959317961 | 1.05983E-54 | 1.06232E-52 | SPAC1039.01 |
| SPCC777.03c | 2.80929804 | 6.10525E-08 | 5.61332E-07 | SPCC777.03c |
| SPRRNA.31 | 2.778134637 | 0.001603031 | 0.006684423 | SPRRNA.31 |
| SPBPB2B2.05 | 2.769898416 | 1.40404E-30 | 6.296E-29 | SPBPB2B2.05 |
| SPAC1002.18 | 2.756371375 | 7.96513E-47 | 7.0203E-45 | urg3 |
| SPCC1739.08c | 2.752832726 | 4.42723E-38 | 2.86481E-36 | SPCC1739.08c |
| SPAC1399.04c | 2.74563147 | 1.14608E-19 | 3.06742E-18 | SPAC1399.04c |
| SPAC521.03 | 2.622062562 | 8.13266E-71 | 1.25982E-68 | SPAC521.03 |
| SPCC285.05 | 2.599258715 | 8.6957E-128 | 4.0411E-125 | SPCC285.05 |
| SPAC513.03 | 2.548104694 | 1.0061E-152 | 7.3474E-150 | mfm2 |
| SPAC6B12.03c | 2.542033041 | 1.11712E-08 | 1.1286E-07 | bit2 |

| | | | | |
|---|---|---|---|---|
| SPAC869.10c | 2.49112136 | 4.54882E-35 | 2.55533E-33 | put4 |
| SPNCRNA.993 | 2.461453295 | 1.09716E-06 | 8.6284E-06 | SPNCRNA.993 |
| SPCC965.08c | 2.392211901 | 1.42367E-56 | 1.45556E-54 | alr1 |
| SPNCRNA.178 | 2.388101827 | 3.16997E-48 | 2.84297E-46 | SPNCRNA.178 |
| SPAC922.07c | 2.381085083 | 2.87954E-60 | 3.42331E-58 | atd2 |
| SPBP4G3.03 | 2.37360431 | 0.000140364 | 0.000753717 | fub2 |
| SPAC3H1.06c | 2.370444977 | 4.50731E-33 | 2.21552E-31 | SPAC3H1.06c |
| SPBPB21E7.11 | 2.362176772 | 3.6691E-10 | 4.42369E-09 | SPBPB21E7.11 |
| SPBPB2B2.18 | 2.34109016 | 0.002142962 | 0.008625844 | SPBPB2B2.18 |
| SPBC800.11 | 2.261787749 | 3.4551E-107 | 1.039E-104 | SPBC800.11 |
| SPBC1105.14 | 2.260423337 | 1.0314E-142 | 5.2727E-140 | rsv2 |
| SPBPB21E7.01c | 2.245642603 | 9.53933E-07 | 7.57222E-06 | eno102 |
| SPAC1399.01c | 2.242588338 | 1.62247E-24 | 5.84089E-23 | SPAC1399.01c |
| SPCC794.04c | 2.237831453 | 2.11621E-96 | 5.69371E-94 | SPCC794.04c |
| SPBC8E4.03 | 2.19410243 | 1.1367E-188 | 2.9053E-185 | SPBC8E4.03 |
| SPNCRNA.1122 | 2.142436114 | 0.000150227 | 0.000802465 | SPNCRNA.1122 |
| SPBC725.10 | 2.131704997 | 1.5144E-06 | 1.15713E-05 | SPBC725.10 |
| SPNCRNA.311 | 2.131073708 | 0.001179429 | 0.005053849 | SPNCRNA.311 |
| SPAC977.13c | 2.12877114 | 7.51738E-16 | 1.54955E-14 | SPAC977.13c |
| SPAC13F5.03c | 2.074526098 | 3.8237E-11 | 5.07708E-10 | gld1 |
| SPBPB2B2.08 | 2.048696759 | 0.000220927 | 0.001146575 | SPBPB2B2.08 |
| SPAC1F7.12 | 2.03352379 | 4.9562E-158 | 4.2227E-155 | yak3 |
| SPAC1039.06 | 2.026789207 | 1.2163E-127 | 5.1814E-125 | SPAC1039.06 |
| SPCC1223.09 | 1.977745426 | 1.16438E-70 | 1.75068E-68 | SPCC1223.09 |
| SPCC338.12 | 1.973645573 | 1.0593E-119 | 3.8679E-117 | pbi2 |
| SPCC737.04 | 1.96936275 | 1.3972E-116 | 4.4641E-114 | SPCC737.04 |
| SPAC869.07c | 1.968427601 | 9.41525E-06 | 6.29984E-05 | mel1 |
| SPBC359.02 | 1.95887583 | 7.45327E-11 | 9.57315E-10 | alr2 |
| SPAC27E2.11c | 1.939025071 | 0.006089799 | 0.0217852 | SPAC27E2.11c |
| SPCC965.06 | 1.936514058 | 1.2315E-172 | 1.5739E-169 | osr2 |
| SPAC19D5.01 | 1.935538694 | 2.01482E-95 | 5.14988E-93 | pyp2 |
| SPBC1271.09 | 1.93149985 | 5.22362E-43 | 4.17236E-41 | tgp1 |
| SPAC806.05 | 1.926129771 | 0.014192684 | 0.044333514 | SPAC806.05 |

| | | | | |
|---|---|---|---|---|
| SPNCRNA.32 | 1.895865385 | 0.00119438 | 0.005105078 | prl32 |
| SPNCRNA.1668 | 1.870674352 | 3.8335E-06 | 2.73317E-05 | mug147-antisense-1 |
| SPNCRNA.12 | 1.868362676 | 0.001151686 | 0.004959915 | prl12 |
| SPCC70.04c | 1.866487937 | 8.4212E-21 | 2.40709E-19 | SPCC70.04c |
| SPCC965.11c | 1.843659166 | 5.2693E-19 | 1.3536E-17 | agp3 |
| SPAC922.09 | 1.843255533 | 7.37882E-08 | 6.6762E-07 | SPAC922.09 |
| SPBPB2B2.06c | 1.824035572 | 1.33171E-12 | 2.06921E-11 | SPBPB2B2.06c |
| SPNCRNA.1527 | 1.770357563 | 0.005161523 | 0.018833479 | SPNCRNA.1527 |
| SPAC869.01 | 1.751953067 | 2.50835E-05 | 0.000153565 | SPAC869.01 |
| SPNCRNA.1696 | 1.75009664 | 1.34507E-21 | 4.11736E-20 | SPNCRNA.1696 |
| SPNCRNA.1015 | 1.739892659 | 0.01022421 | 0.033546959 | SPNCRNA.1015 |
| SPCPB16A4.06c | 1.727400327 | 9.40162E-33 | 4.49169E-31 | SPCPB16A4.06c |
| SPBP4G3.02 | 1.719236935 | 6.52485E-20 | 1.7742E-18 | pho1 |
| SPNCRNA.1216 | 1.714752668 | 1.60525E-07 | 1.40514E-06 | SPNCRNA.1216 |
| SPAC1399.03 | 1.712219694 | 2.0773E-21 | 6.32093E-20 | fur4 |
| SPBC1271.08c | 1.70107412 | 4.79574E-22 | 1.50404E-20 | SPBC1271.08c |
| SPNCRNA.987 | 1.697569698 | 0.001015977 | 0.004465757 | SPAC9E9.17c-antisense-1 |
| SPAC17G6.03 | 1.692595249 | 2.5616E-05 | 0.000155706 | SPAC17G6.03 |
| SPAC1F7.09c | 1.675857932 | 2.47044E-74 | 4.07384E-72 | SPAC1F7.09c |
| SPBC19C7.04c | 1.664389779 | 3.35685E-07 | 2.8411E-06 | SPBC19C7.04c |
| SPNCRNA.282 | 1.656014834 | 1.62088E-05 | 0.000103964 | SPNCRNA.282 |
| SPBC1773.03c | 1.64270025 | 1.3144E-68 | 1.86645E-66 | SPBC1773.03c |
| SPBC16E9.16c | 1.641655378 | 4.3631E-78 | 7.96578E-76 | lsd90 |
| SPBC1271.07c | 1.632085552 | 0.01572153 | 0.048240374 | SPBC1271.07c |
| SPAC13G7.02c | 1.60214143 | 1.0572E-173 | 1.8015E-170 | ssa1 |
| SPNCRNA.1691 | 1.590178873 | 2.4323E-05 | 0.000149987 | SPNCRNA.1691 |
| SPAC17C9.16c | 1.581688218 | 9.42295E-85 | 2.09435E-82 | mfs1 |
| SPAC869.08 | 1.540135075 | 2.01302E-06 | 1.50008E-05 | pcm2 |
| SPNCRNA.1306 | 1.536662507 | 0.011931929 | 0.03845903 | SPNCRNA.1306 |
| SPCC757.13 | 1.532948145 | 9.11409E-21 | 2.5884E-19 | SPCC757.13 |
| SPAC22H10.13 | 1.528603496 | 2.31659E-74 | 3.94747E-72 | zym1 |
| SPBC1683.05 | 1.524534364 | 1.00179E-24 | 3.68427E-23 | SPBC1683.05 |
| SPBPB2B2.12c | 1.508708239 | 0.000251294 | 0.00128719 | gal10 |

| | | | | |
|---|---|---|---|---|
| SPCC794.01c | 1.508587436 | 6.33493E-40 | 4.26895E-38 | zwf2 |
| SPAC3G6.07 | 1.506055389 | 0.000227367 | 0.001177609 | SPAC3G6.07 |
| SPAC11D3.09 | 1.504328814 | 1.51658E-06 | 1.15713E-05 | SPAC11D3.09 |
| SPNCRNA.925 | 1.486670919 | 0.00625178 | 0.02225564 | SPAC8C9.04-antisense-1 |
| SPBC16D10.08c | 1.479063116 | 5.7707E-127 | 2.2692E-124 | hsp104 |
| SPAC513.02 | 1.466278043 | 7.80681E-21 | 2.25471E-19 | SPAC513.02 |
| SPAPJ691.02 | 1.450772833 | 2.81024E-27 | 1.12234E-25 | SPAPJ691.02 |
| SPAC31G5.09c | 1.450245923 | 5.48413E-07 | 4.47842E-06 | spk1 |
| SPBC1773.17c | 1.44862167 | 4.03156E-45 | 3.43489E-43 | SPBC1773.17c |
| SPNCRNA.1459 | 1.443766949 | 2.81714E-15 | 5.51772E-14 | SPNCRNA.1459 |
| SPNCRNA.1489 | 1.426868025 | 3.68409E-12 | 5.49068E-11 | aim27-antisense-1 |
| SPNCRNA.908 | 1.425313495 | 0.0030257 | 0.011629608 | rpt5-antisense-1 |
| SPNCRNA.451 | 1.423380837 | 0.002556858 | 0.010124445 | SPNCRNA.451 |
| SPBP26C9.02c | 1.422837418 | 2.3263E-45 | 2.0156E-43 | car1 |
| SPAC26F1.11 | 1.420265237 | 2.01163E-13 | 3.3606E-12 | SPAC26F1.11 |
| SPBC32C12.02 | 1.405256153 | 4.69464E-50 | 4.36346E-48 | ste11 |
| SPAPB24D3.03 | 1.401268923 | 3.03291E-54 | 2.92533E-52 | SPAPB24D3.03 |
| SPBPB21E7.04c | 1.396481704 | 1.55035E-23 | 5.35498E-22 | SPBPB21E7.04c |
| SPNCRNA.1278 | 1.393964216 | 1.24916E-13 | 2.12856E-12 | SPNCRNA.1278 |
| SPBC3E7.02c | 1.39227683 | 2.11071E-09 | 2.36104E-08 | hsp16 |
| SPCC11E10.01 | 1.370592891 | 8.65752E-79 | 1.63916E-76 | SPCC11E10.01 |
| SPCC965.07c | 1.368723961 | 4.85234E-98 | 1.37806E-95 | gst2 |
| SPCC1494.01 | 1.360763403 | 2.42798E-19 | 6.39786E-18 | SPCC1494.01 |
| SPNCRNA.1316 | 1.342206498 | 2.07294E-24 | 7.41038E-23 | SPNCRNA.1316 |
| SPCC1450.07c | 1.341308364 | 2.8034E-43 | 2.27476E-41 | dao1 |
| SPAP7G5.06 | 1.32192467 | 5.99437E-14 | 1.04584E-12 | per1 |
| SPAC1565.04c | 1.311140149 | 2.56149E-23 | 8.72954E-22 | ste4 |
| SPBPB21E7.02c | 1.294170826 | 0.015495006 | 0.047752651 | SPBPB21E7.02c |
| SPAC5H10.02c | 1.285890307 | 0.009760558 | 0.032170194 | hsp3102 |
| SPCC4G3.03 | 1.276976695 | 1.47322E-24 | 5.37937E-23 | SPCC4G3.03 |
| SPBC1683.02 | 1.273630255 | 6.84995E-21 | 2.0241E-19 | SPBC1683.02 |
| SPAC20H4.11c | 1.273555584 | 2.45962E-13 | 4.06911E-12 | rho5 |
| SPBC1861.05 | 1.270691911 | 8.38721E-11 | 1.07189E-09 | SPBC1861.05 |

| | | | | |
|---|---|---|---|---|
| SPCC132.04c | 1.265581739 | 2.35913E-44 | 1.97703E-42 | gdh2 |
| SPAC4G9.22 | 1.251628691 | 2.01483E-36 | 1.18388E-34 | SPAC4G9.22 |
| SPCC191.06 | 1.248736976 | 6.64893E-07 | 5.3659E-06 | SPCC191.06 |
| SPCC13B11.04c | 1.247045317 | 1.9292E-37 | 1.20269E-35 | fmd3 |
| SPNCRNA.1472 | 1.236741464 | 0.009242516 | 0.030921297 | SPNCRNA.1472 |
| SPCC16A11.15c | 1.235174708 | 7.77676E-43 | 6.02345E-41 | SPCC16A11.15c |
| SPBC359.06 | 1.233165864 | 6.10927E-13 | 9.85193E-12 | mug14 |
| SPNCRNA.834 | 1.225580105 | 3.71958E-06 | 2.66672E-05 | SPNCRNA.834 |
| SPBC1773.02c | 1.225099325 | 6.05507E-61 | 7.36988E-59 | bcp1 |
| SPAC1F8.01 | 1.218552837 | 0.000277247 | 0.001406038 | ght3 |
| SPAC2E1P3.04 | 1.216669525 | 1.27043E-68 | 1.85556E-66 | cao1 |
| SPBC56F2.06 | 1.215605374 | 6.89822E-06 | 4.7081E-05 | mug147 |
| SPAC11E3.06 | 1.207102064 | 4.46172E-06 | 3.13874E-05 | map1 |
| SPAC25B8.13c | 1.191989927 | 0.000848192 | 0.003816864 | isp7 |
| SPAC9E9.09c | 1.191306355 | 2.80681E-16 | 6.02874E-15 | atd1 |
| SPAC323.07c | 1.186295253 | 6.8325E-50 | 6.23709E-48 | SPAC323.07c |
| SPAC4A8.04 | 1.186096189 | 6.62053E-63 | 8.90635E-61 | isp6 |
| SPNCRNA.897 | 1.184404436 | 0.009530873 | 0.031658103 | ace2-antisense-1 |
| SPAC869.05c | 1.181372471 | 5.51642E-33 | 2.68571E-31 | SPAC869.05c |
| SPNCRNA.1366 | 1.17170078 | 5.62397E-09 | 5.92778E-08 | SPNCRNA.1366 |
| SPBC16G5.02c | 1.157228581 | 7.76866E-43 | 6.02345E-41 | rbk1 |
| SPAC31G5.14 | 1.15708918 | 6.27363E-32 | 2.91553E-30 | gcv1 |
| SPAC24C9.08 | 1.155019209 | 3.36041E-82 | 7.15767E-80 | SPAC24C9.08 |
| SPBC16D10.10 | 1.152826729 | 3.58994E-39 | 2.35279E-37 | tad2 |
| SPAC637.03 | 1.140736289 | 8.6909E-81 | 1.77712E-78 | SPAC637.03 |
| SPNCRNA.1327 | 1.140573109 | 0.014927074 | 0.04624679 | SPNCRNA.1327 |
| SPNCRNA.920 | 1.138853401 | 0.002365631 | 0.009410977 | SPNCRNA.920 |
| SPAC212.06c | 1.134749652 | 0.013010034 | 0.041308877 | SPAC212.06c |
| SPAC1805.16c | 1.132463213 | 2.68396E-39 | 1.78187E-37 | SPAC1805.16c |
| SPBC359.01 | 1.120609158 | 4.46349E-42 | 3.40557E-40 | SPBC359.01 |
| SPBC1198.07c | 1.11773288 | 0.004218905 | 0.015677366 | SPBC1198.07c |
| SPAC29B12.14c | 1.117427788 | 3.14955E-34 | 1.67714E-32 | SPAC29B12.14c |
| SPBPB21E7.07 | 1.117081573 | 1.69235E-36 | 1.00596E-34 | aes1 |

| | | | | |
|---|---|---|---|---|
| SPNCRNA.279 | 1.085361015 | 0.003596586 | 0.013610893 | SPNCRNA.279 |
| SPCC70.03c | 1.084669359 | 3.06851E-72 | 4.90195E-70 | SPCC70.03c |
| SPBC24C6.06 | 1.080871268 | 2.44495E-21 | 7.39561E-20 | gpa1 |
| SPAC57A7.05 | 1.076961601 | 1.24812E-43 | 1.0291E-41 | SPAC57A7.05 |
| SPCC794.02 | 1.076891104 | 6.31007E-17 | 1.41479E-15 | wtf5 |
| SPNCRNA.1350 | 1.07671218 | 0.002105518 | 0.008501901 | SPNCRNA.1350 |
| SPAC19G12.09 | 1.076335671 | 1.81706E-63 | 2.51049E-61 | SPAC19G12.09 |
| SPAC8C9.03 | 1.071484982 | 3.03435E-22 | 9.63452E-21 | cgs1 |
| SPAC26H5.09c | 1.070070481 | 2.59953E-62 | 3.40739E-60 | SPAC26H5.09c |
| SPCC1393.14 | 1.062958284 | 1.10747E-05 | 7.29559E-05 | ten1 |
| SPAC15E1.02c | 1.061556451 | 9.06436E-22 | 2.79139E-20 | SPAC15E1.02c |
| SPAC23E2.03c | 1.060717693 | 2.09997E-14 | 3.79331E-13 | ste7 |
| SPCC1442.04c | 1.05050489 | 1.01841E-33 | 5.2587E-32 | SPCC1442.04c |
| SPAC24B11.05 | 1.046740245 | 1.54497E-32 | 7.31288E-31 | SPAC24B11.05 |
| SPBC725.03 | 1.04398143 | 1.73466E-33 | 8.86759E-32 | SPBC725.03 |
| SPBC23G7.10c | 1.03490937 | 4.85778E-25 | 1.81263E-23 | SPBC23G7.10c |
| SPCC1753.02c | 1.028714805 | 8.95882E-42 | 6.73492E-40 | git3 |
| SPAC222.15 | 1.023236232 | 0.009659253 | 0.031959937 | meu13 |
| SPCC548.07c | 1.022617804 | 8.0687E-10 | 9.41716E-09 | ght1 |
| SPNCRNA.1503 | 1.020224815 | 0.000590795 | 0.002753093 | SPNCRNA.1503 |
| SPAC4F10.20 | 1.017351736 | 3.4753E-62 | 4.44143E-60 | grx1 |
| SPAC212.12 | 1.013526154 | 0.003208463 | 0.012267513 | SPAC212.12 |
| SPBC119.03 | 1.010369526 | 1.44561E-57 | 1.57233E-55 | SPBC119.03 |
| SPBC30D10.14 | 1.010337617 | 4.67972E-37 | 2.84794E-35 | SPBC30D10.14 |
| SPCC1393.12 | 1.007628229 | 1.04564E-28 | 4.3814E-27 | SPCC1393.12 |
| SPAC2F3.05c | 1.00375978 | 1.27639E-36 | 7.67635E-35 | SPAC2F3.05c |
| SPBC2G2.04c | 1.002634905 | 1.58493E-79 | 3.11621E-77 | mmf1 |
| SPBC1348.14c | 0.99659047 | 0.003473071 | 0.013200254 | ght7 |
| SPAC32A11.02c | 0.99511735 | 4.40656E-57 | 4.69299E-55 | SPAC32A11.02c |
| SPCC1620.05 | 0.984569562 | 0.000162404 | 0.000861214 | SPCC1620.05 |
| SPNCRNA.1601 | 0.979834586 | 0.000387529 | 0.001886715 | SPBC2G2.17c-antisense-1 |
| SPCPB16A4.07 | 0.976598021 | 1.26874E-37 | 8.00717E-36 | SPCPB16A4.07 |
| SPAC1F7.10 | 0.965032079 | 1.11245E-18 | 2.78767E-17 | SPAC1F7.10 |

| | | | | |
|---|---|---|---|---|
| SPAC688.03c | 0.964889935 | 3.85623E-11 | 5.10701E-10 | SPAC688.03c |
| SPBPJ4664.02 | 0.962336518 | 7.72354E-30 | 3.43328E-28 | SPBPJ4664.02 |
| SPBC1683.12 | 0.949340697 | 1.36137E-11 | 1.91716E-10 | SPBC1683.12 |
| SPCC1906.04 | 0.937367596 | 0.000100012 | 0.000550926 | wtf20 |
| SPNCRNA.1558 | 0.934700189 | 0.008360892 | 0.028474938 | qcr10-antisense-1 |
| SPBC428.05c | 0.933048615 | 3.24357E-60 | 3.76844E-58 | arg12 |
| SPBC2D10.06 | 0.922017358 | 5.94575E-06 | 4.10288E-05 | rep1 |
| SPNCRNA.828 | 0.916610197 | 0.010692696 | 0.034860372 | SPNCRNA.828 |
| SPBC3H7.02 | 0.910074192 | 1.72196E-29 | 7.33556E-28 | SPBC3H7.02 |
| SPBC56F2.15 | 0.907679624 | 2.61548E-26 | 1.00529E-24 | tam13 |
| SPCC736.13 | 0.907381429 | 2.37147E-33 | 1.18852E-31 | SPCC736.13 |
| SPBC1773.05c | 0.902184732 | 7.33988E-21 | 2.1319E-19 | tms1 |
| SPAC14C4.03 | 0.900762139 | 0.008328058 | 0.028398027 | mek1 |
| SPBC106.02c | 0.895798805 | 3.04779E-05 | 0.000183513 | srx1 |
| SPBC337.08c | 0.895492874 | 1.20776E-54 | 1.18732E-52 | ubi4 |
| SPAC13G6.06c | 0.895028705 | 1.0835E-61 | 1.35094E-59 | gcv2 |
| SPBC1683.11c | 0.892498954 | 0.004213109 | 0.015674971 | SPBC1683.11c |
| SPNCRNA.1092 | 0.890214069 | 4.82862E-07 | 3.95575E-06 | SPNCRNA.1092 |
| SPAC22F3.12c | 0.888514109 | 2.80633E-11 | 3.77525E-10 | rgs1 |
| SPNCRNA.1188 | 0.883674591 | 0.00168335 | 0.006979146 | pbi2-antisense-1 |
| SPAC16.05c | 0.878013124 | 3.15773E-19 | 8.23587E-18 | sfp1 |
| SPAC11D3.16c | 0.871011944 | 3.03757E-15 | 5.92674E-14 | SPAC11D3.16c |
| SPAC15E1.10 | 0.868013165 | 0.000295833 | 0.001484099 | fub1 |
| SPCC285.04 | 0.865398766 | 2.26898E-10 | 2.78154E-09 | SPCC285.04 |
| SPAC977.16c | 0.864878541 | 3.49985E-11 | 4.65917E-10 | dak2 |
| SPBC1348.12 | 0.863998289 | 4.87915E-07 | 3.99075E-06 | SPBC1348.12 |
| SPCPB1C11.02 | 0.860294505 | 1.52469E-22 | 4.93306E-21 | SPCPB1C11.02 |
| SPNCRNA.1352 | 0.85450125 | 0.006763542 | 0.023763041 | SPNCRNA.1352 |
| SPAC3A11.10c | 0.848992243 | 1.04633E-30 | 4.73346E-29 | SPAC3A11.10c |
| SPAC26F1.04c | 0.842495035 | 2.49609E-26 | 9.66667E-25 | etr1 |
| SPCC70.09c | 0.836089158 | 0.004581093 | 0.016933152 | mug9 |
| SPAP8A3.04c | 0.835872822 | 9.23264E-38 | 5.89966E-36 | hsp9 |
| SPNCRNA.1565 | 0.835140375 | 0.001009936 | 0.004443025 | SPNCRNA.1565 |

| | | | | |
|---|---|---|---|---|
| SPCC191.11 | 0.832906909 | 1.65592E-14 | 3.02324E-13 | inv1 |
| SPAPB1A10.05 | 0.831175936 | 5.26191E-16 | 1.10798E-14 | SPAPB1A10.05 |
| SPBC8E4.04 | 0.829473903 | 2.20924E-34 | 1.20145E-32 | SPBC8E4.04 |
| SPAC17G6.13 | 0.824966033 | 3.14787E-33 | 1.56232E-31 | slt1 |
| SPCC576.02 | 0.823990959 | 1.63166E-13 | 2.75282E-12 | SPCC576.02 |
| SPAC1399.05c | 0.823408051 | 0.000129372 | 0.000699768 | toe1 |
| SPBPB21E7.09 | 0.814472381 | 4.14869E-18 | 1.00991E-16 | SPBPB21E7.09 |
| SPAC32A11.01 | 0.812896864 | 1.70081E-09 | 1.92784E-08 | mug8 |
| SPAC2E1P3.05c | 0.812858841 | 4.3625E-36 | 2.50574E-34 | SPAC2E1P3.05c |
| SPAC823.16c | 0.809507803 | 2.15611E-14 | 3.88101E-13 | atg1802 |
| SPAC1039.05c | 0.805180367 | 2.4712E-20 | 6.90314E-19 | klf1 |
| SPACUNK4.16c | 0.802876514 | 1.55237E-41 | 1.15011E-39 | SPACUNK4.16c |
| SPAPB24D3.04c | 0.801237866 | 0.000174718 | 0.000917942 | mag1 |
| SPBPB7E8.02 | 0.793685721 | 9.83375E-23 | 3.22245E-21 | SPBPB7E8.02 |
| SPAC1751.01c | 0.793152777 | 2.34036E-19 | 6.19892E-18 | gti1 |
| SPAC9E9.11 | 0.791920498 | 1.70311E-31 | 7.8435E-30 | plr1 |
| SPBC1683.08 | 0.791128722 | 1.06126E-14 | 1.98725E-13 | ght4 |
| SPCC24B10.14c | 0.782635646 | 0.001888181 | 0.007730955 | xlf1 |
| SPCC1620.02 | 0.780793327 | 0.000239672 | 0.001233838 | wtf23 |
| SPCC417.02 | 0.780408711 | 1.10095E-06 | 8.63195E-06 | dad5 |
| SPCC576.03c | 0.774935411 | 1.19273E-50 | 1.12912E-48 | tpx1 |
| SPAC589.07c | 0.773770723 | 0.000608329 | 0.002827072 | atg1801 |
| SPAC26F1.14c | 0.769364917 | 1.75464E-22 | 5.64133E-21 | aif1 |
| SPCC965.13 | 0.768868378 | 8.77253E-34 | 4.57604E-32 | SPCC965.13 |
| SPAC750.01 | 0.768535835 | 2.5401E-05 | 0.000154767 | SPAC750.01 |
| SPBC660.05 | 0.763671839 | 1.60926E-05 | 0.000103348 | SPBC660.05 |
| SPBC16A3.02c | 0.760681509 | 5.18411E-36 | 2.94458E-34 | SPBC16A3.02c |
| SPAC2F3.08 | 0.756390024 | 5.28877E-14 | 9.32283E-13 | sut1 |
| SPAC4H3.03c | 0.755565294 | 2.76623E-15 | 5.43884E-14 | SPAC4H3.03c |
| SPCC1235.14 | 0.752734689 | 9.73544E-06 | 6.49707E-05 | ght5 |
| SPBC1105.19 | 0.749583032 | 6.09624E-07 | 4.95855E-06 | tam12 |
| SPAC2E1P3.01 | 0.749081678 | 7.0326E-21 | 2.06613E-19 | SPAC2E1P3.01 |
| SPAC4D7.02c | 0.747974175 | 1.25776E-17 | 2.89626E-16 | pgc1 |

| | | | | |
|---|---|---|---|---|
| SPAC11H11.04 | 0.746652702 | 1.83942E-13 | 3.08299E-12 | mam2 |
| SPBC32F12.03c | 0.746152567 | 4.60682E-37 | 2.83736E-35 | gpx1 |
| SPAC29A4.17c | 0.742674027 | 3.85074E-07 | 3.22705E-06 | SPAC29A4.17c |
| SPBC660.06 | 0.74254899 | 1.92634E-33 | 9.74996E-32 | SPBC660.06 |
| SPNCRNA.904 | 0.741672521 | 0.001669089 | 0.006925637 | dmc1-antisense-1 |
| SPCC338.18 | 0.740811406 | 0.000342251 | 0.001696727 | SPCC338.18 |
| SPCC417.10 | 0.733830114 | 8.90987E-18 | 2.08932E-16 | dal51 |
| SPCC320.14 | 0.733117061 | 1.45787E-15 | 2.9226E-14 | sry1 |
| SPCC417.16 | 0.731482666 | 3.05648E-23 | 1.03475E-21 | SPCC417.16 |
| SPNCRNA.651 | 0.730325886 | 0.007490588 | 0.026048901 | SPNCRNA.651 |
| SPAC29B12.13 | 0.719979012 | 2.74702E-07 | 2.34046E-06 | SPAC29B12.13 |
| SPAC688.04c | 0.718068246 | 8.42859E-21 | 2.40709E-19 | gst3 |
| SPCC16A11.08 | 0.717416168 | 4.98033E-10 | 5.9208E-09 | atg20 |
| SPBC11C11.06c | 0.711515888 | 2.65601E-34 | 1.42921E-32 | SPBC11C11.06c |
| SPBC660.07 | 0.710554983 | 6.34664E-40 | 4.26895E-38 | ntp1 |
| SPAC3C7.13c | 0.709694602 | 1.8868E-16 | 4.06975E-15 | SPAC3C7.13c |
| SPBC1685.05 | 0.704214676 | 1.26429E-12 | 1.97044E-11 | SPBC1685.05 |
| SPCC191.01 | 0.703044648 | 7.81706E-24 | 2.75592E-22 | SPCC191.01 |
| SPBC8E4.01c | 0.70228162 | 7.80472E-25 | 2.89114E-23 | pho84 |
| SPCC757.12 | 0.696924541 | 9.63228E-30 | 4.20857E-28 | SPCC757.12 |
| SPBPB2B2.13 | 0.696105577 | 8.47912E-05 | 0.000473201 | gal1 |
| SPAC23D3.12 | 0.69481015 | 0.000298265 | 0.001493372 | SPAC23D3.12 |
| SPAC22A12.11 | 0.690583614 | 8.08888E-30 | 3.56469E-28 | dak1 |
| SPBC21C3.19 | 0.688733745 | 9.67799E-35 | 5.3776E-33 | SPBC21C3.19 |
| SPBC1347.11 | 0.688026253 | 6.78762E-10 | 7.995E-09 | sro1 |
| SPAC3A12.08 | 0.687018903 | 2.57365E-06 | 1.88489E-05 | SPAC3A12.08 |
| SPAC1B3.01c | 0.686802419 | 0.010823188 | 0.035195733 | SPAC1B3.01c |
| SPAC10F6.05c | 0.686557612 | 2.20071E-11 | 2.99202E-10 | ubc6 |
| SPAP11E10.01 | 0.686508846 | 4.17032E-15 | 7.98453E-14 | SPAP11E10.01 |
| SPAPB18E9.05c | 0.680947251 | 5.74566E-27 | 2.27688E-25 | SPAPB18E9.05c |
| SPBC19G7.06 | 0.68025031 | 7.54908E-10 | 8.85112E-09 | mbx1 |
| SPCC162.06c | 0.679115954 | 4.086E-12 | 6.07199E-11 | SPCC162.06c |
| SPAC22A12.17c | 0.67702964 | 5.33452E-05 | 0.000308835 | SPAC22A12.17c |

| | | | | |
|---|---|---|---|---|
| SPNCRNA.812 | 0.676907584 | 1.30168E-06 | 1.00668E-05 | SPAC6C3.02c-antisense-1 |
| SPCC965.09 | 0.676017024 | 4.40496E-07 | 3.65555E-06 | SPCC965.09 |
| SPNCRNA.738 | 0.675861197 | 0.009053376 | 0.030387958 | SPNCRNA.738 |
| SPAC458.04c | 0.675832572 | 0.009009693 | 0.030281097 | dli1 |
| SPBC1271.14 | 0.673973185 | 1.08272E-19 | 2.91309E-18 | SPBC1271.14 |
| SPBC365.12c | 0.671156267 | 3.23391E-19 | 8.39174E-18 | ish1 |
| SPAC977.14c | 0.66762079 | 2.70325E-40 | 1.9193E-38 | SPAC977.14c |
| SPCC553.05c | 0.665192256 | 0.005994718 | 0.021498499 | wtf6 |
| SPNCRNA.1560 | 0.657964585 | 0.000211754 | 0.001101209 | SPNCRNA.1560 |
| SPAC4F8.01 | 0.657756238 | 2.19501E-12 | 3.31979E-11 | did4 |
| SPAC19G12.04 | 0.656683477 | 1.96082E-05 | 0.000122539 | dal1 |
| SPCC4B3.01 | 0.653411142 | 9.85283E-17 | 2.18042E-15 | tum1 |
| SPCC417.09c | 0.652935073 | 9.11434E-13 | 1.44249E-11 | SPCC417.09c |
| SPAPB24D3.10c | 0.652278169 | 1.87942E-08 | 1.84407E-07 | agl1 |
| SPBC1683.07 | 0.649647499 | 5.68228E-06 | 3.94136E-05 | mal1 |
| SPAC977.18 | 0.648953309 | 0.011016101 | 0.035777831 | SPAC977.18 |
| SPAC3C7.05c | 0.648942914 | 2.85742E-14 | 5.0896E-13 | mug191 |
| SPAC1782.12c | 0.644335074 | 1.65737E-05 | 0.000105642 | SPAC1782.12c |
| SPBP19A11.01 | 0.643468402 | 4.20605E-31 | 1.91976E-29 | gcv3 |
| SPAC20G4.02c | 0.643375017 | 0.000228987 | 0.001184801 | fus1 |
| SPBC2A9.02 | 0.641945487 | 8.54799E-18 | 2.02302E-16 | SPBC2A9.02 |
| SPCC306.10 | 0.639865382 | 0.013162428 | 0.041611831 | wtf8 |
| SPAC31G5.12c | 0.638838787 | 3.35598E-08 | 3.23085E-07 | maf1 |
| SPBC21.07c | 0.637230124 | 3.01411E-06 | 2.19489E-05 | ppk24 |
| SPCC162.11c | 0.637097739 | 1.63034E-14 | 2.9872E-13 | SPCC162.11c |
| SPAC20G8.10c | 0.636858796 | 2.58888E-07 | 2.20941E-06 | atg6 |
| SPAC977.10 | 0.634643343 | 9.34949E-18 | 2.1824E-16 | sod2 |
| SPBC215.11c | 0.634578514 | 1.67956E-19 | 4.47183E-18 | SPBC215.11c |
| SPAPB18E9.04c | 0.63250102 | 5.20796E-09 | 5.53494E-08 | SPAPB18E9.04c |
| SPBC32H8.07 | 0.629891496 | 1.01618E-13 | 1.74907E-12 | git5 |
| SPBP8B7.24c | 0.626075705 | 1.63052E-13 | 2.75282E-12 | atg8 |
| SPACUNK4.10 | 0.626046622 | 1.16892E-22 | 3.80606E-21 | SPACUNK4.10 |
| SPAC1002.12c | 0.625938599 | 2.21042E-12 | 3.33323E-11 | SPAC1002.12c |

| | | | | |
|---|---|---|---|---|
| SPBC12D12.09 | 0.625247947 | 0.000318444 | 0.001584313 | rev7 |
| SPAC3H8.04 | 0.625042968 | 1.75753E-13 | 2.95542E-12 | SPAC3H8.04 |
| SPCPB1C11.03 | 0.624667244 | 2.56439E-07 | 2.19217E-06 | SPCPB1C11.03 |
| SPBC4F6.17c | 0.619951157 | 8.45072E-24 | 2.95891E-22 | SPBC4F6.17c |
| SPAC11D3.19 | 0.616942742 | 0.00151877 | 0.006369117 | SPAC11D3.19 |
| SPCC16A11.01 | 0.616154118 | 5.19103E-18 | 1.25766E-16 | sfk1 |
| SPCC306.11 | 0.612933513 | 1.54814E-24 | 5.61281E-23 | SPCC306.11 |
| SPNCRNA.1328 | 0.60852349 | 0.000388201 | 0.001888188 | SPBC660.08-antisense-1 |
| SPCC1739.06c | 0.607266791 | 6.21971E-18 | 1.49273E-16 | SPCC1739.06c |
| SPCPB1C11.01 | 0.606442936 | 0.016304998 | 0.049584266 | amt1 |
| SPBC17F3.01c | 0.603570401 | 1.11052E-06 | 8.69372E-06 | rga5 |
| SPAC11D3.08c | 0.603547239 | 1.55597E-13 | 2.64256E-12 | SPAC11D3.08c |
| SPBC36.02c | 0.603482848 | 3.73268E-09 | 4.02562E-08 | SPBC36.02c |
| SPCC830.07c | 0.599473747 | 9.59032E-23 | 3.18496E-21 | psi1 |
| SPCC306.08c | 0.598172816 | 1.28819E-25 | 4.87794E-24 | mdh1 |
| SPBC354.12 | 0.597022623 | 0.000278692 | 0.001411964 | gpd3 |
| SPAPB8E5.04c | 0.59679765 | 6.05351E-22 | 1.88692E-20 | npc2 |
| SPCC285.06c | 0.594093005 | 0.001965921 | 0.0080078 | wtf17 |
| SPAC26F1.07 | 0.5935564 | 3.80779E-32 | 1.78582E-30 | SPAC26F1.07 |
| SPBC31E1.04 | 0.593135914 | 2.68991E-14 | 4.80798E-13 | pep12 |
| SPNCRNA.1159 | 0.590119246 | 0.014833624 | 0.045985132 | SPNCRNA.1159 |
| SPBC23E6.01c | 0.587611169 | 1.92408E-11 | 2.62992E-10 | cxr1 |
| SPNCRNA.854 | 0.587312837 | 0.009527227 | 0.031658103 | SPNCRNA.854 |
| SPAC977.15 | 0.585117124 | 0.002735747 | 0.010684602 | SPAC977.15 |
| SPAC23G3.03 | 0.583668299 | 5.1721E-14 | 9.14871E-13 | sib2 |
| SPBC337.16 | 0.582522765 | 3.54848E-19 | 9.16154E-18 | cho1 |
| SPAC4H3.08 | 0.580634655 | 1.81899E-05 | 0.000114941 | SPAC4H3.08 |
| SPBC21B10.08c | 0.574962568 | 9.59475E-23 | 3.18496E-21 | SPBC21B10.08c |
| SPBC354.15 | 0.57446989 | 7.60514E-11 | 9.74373E-10 | fap1 |
| SPCC1450.13c | 0.573452177 | 1.46983E-10 | 1.82373E-09 | SPCC1450.13c |
| SPAC20G4.03c | 0.57266873 | 1.33709E-10 | 1.6712E-09 | hri1 |
| SPCC663.17 | 0.572435595 | 0.000564061 | 0.002638136 | wtf15 |
| SPCC162.04c | 0.570389467 | 0.001807183 | 0.007426301 | wtf13 |

| | | | | |
|---|---|---|---|---|
| SPAC821.10c | 0.569618723 | 3.77216E-20 | 1.04234E-18 | sod1 |
| SPBC1711.11 | 0.569211131 | 5.9883E-06 | 4.12563E-05 | SPBC1711.11 |
| SPBP4H10.12 | 0.568903936 | 3.96227E-11 | 5.20697E-10 | SPBP4H10.12 |
| SPAC6F12.03c | 0.568653861 | 1.87492E-08 | 1.84319E-07 | fsv1 |
| SPBP19A11.02c | 0.565925461 | 2.79831E-07 | 2.38019E-06 | SPBP19A11.02c |
| SPBC3D6.06c | 0.563737768 | 9.69739E-23 | 3.19826E-21 | prs5 |
| SPCC1442.16c | 0.562886494 | 1.38605E-16 | 3.02798E-15 | zta1 |
| SPAC4G9.19 | 0.562878523 | 5.84581E-06 | 4.04929E-05 | SPAC4G9.19 |
| SPAC17A2.13c | 0.559830502 | 1.29049E-29 | 5.59069E-28 | rad25 |
| SPBC19C2.05 | 0.559433853 | 1.39784E-06 | 1.07455E-05 | pat1 |
| SPAC10F6.06 | 0.557971586 | 1.57643E-29 | 6.77204E-28 | vip1 |
| SPBC13A2.04c | 0.557714322 | 0.005012871 | 0.018369747 | ptr2 |
| SPAC56E4.03 | 0.554456314 | 2.8953E-21 | 8.70633E-20 | SPAC56E4.03 |
| SPAC22H12.01c | 0.554027611 | 2.42663E-07 | 2.08837E-06 | mug35 |
| SPBC25B2.02c | 0.549597491 | 8.92554E-10 | 1.03463E-08 | mam1 |
| SPBC106.13 | 0.546537463 | 8.12658E-06 | 5.48786E-05 | gid9 |
| SPAC4H3.04c | 0.545479003 | 0.000329491 | 0.001636886 | SPAC4H3.04c |
| SPAC328.03 | 0.545285315 | 5.97311E-24 | 2.12045E-22 | tps1 |
| SPAC16E8.03 | 0.543891581 | 0.005216269 | 0.019019663 | gna1 |
| SPAC1687.02 | 0.543827893 | 0.000200506 | 0.001045902 | SPAC1687.02 |
| SPBC1652.01 | 0.54327143 | 1.54338E-08 | 1.53796E-07 | SPBC1652.01 |
| SPCC320.03 | 0.541010344 | 1.75916E-20 | 4.94111E-19 | SPCC320.03 |
| SPNCRNA.1473 | 0.538767369 | 0.008762333 | 0.029625029 | spo14-antisense-1 |
| SPAC7D4.07c | 0.536204436 | 9.38076E-20 | 2.53727E-18 | trx1 |
| SPAC630.07c | 0.533331471 | 7.08491E-06 | 4.82265E-05 | SPAC630.07c |
| SPBC428.11 | 0.53214524 | 1.11007E-15 | 2.25185E-14 | met17 |
| SPNCRNA.975 | 0.528624557 | 5.94437E-06 | 4.10288E-05 | SPNCRNA.975 |
| SPAC20H4.02 | 0.523430206 | 0.001604418 | 0.006684423 | dsc3 |
| SPBC8E4.02c | 0.518704705 | 1.95851E-05 | 0.000122539 | SPBC8E4.02c |
| SPBC16C6.14 | 0.516484336 | 0.001498777 | 0.006295602 | spo2 |
| SPCC830.02 | 0.516297097 | 0.002113598 | 0.008526159 | wtf24 |
| SPAC1F7.07c | 0.516254897 | 0.000653929 | 0.003017042 | fip1 |
| SPAC869.02c | 0.514867638 | 6.83151E-10 | 8.02821E-09 | SPAC869.02c |

| | | | | |
|---|---|---|---|---|
| SPAC26F1.10c | 0.514569761 | 1.28303E-11 | 1.81183E-10 | pyp1 |
| SPBC1773.06c | 0.514362233 | 4.37856E-07 | 3.64193E-06 | adh8 |
| SPAC1783.07c | 0.512856665 | 2.65291E-18 | 6.48885E-17 | pap1 |
| SPAC222.17 | 0.511311329 | 0.005475161 | 0.019892696 | SPAC222.17 |
| SPAC1635.01 | 0.511253734 | 4.21055E-23 | 1.41607E-21 | SPAC1635.01 |
| SPCC757.11c | 0.511148455 | 4.60208E-07 | 3.79449E-06 | SPCC757.11c |
| SPCC70.10 | 0.510643138 | 2.27024E-06 | 1.68195E-05 | SPCC70.10 |
| SPBC215.14c | 0.50578495 | 0.000201654 | 0.001050822 | vps20 |
| SPBP16F5.03c | 0.503886545 | 5.53337E-09 | 5.85643E-08 | tra1 |
| SPCC1739.15 | 0.502317089 | 0.013634522 | 0.042865731 | wtf21 |
| SPBP35G2.13c | 0.498279775 | 3.64033E-06 | 2.61486E-05 | swc2 |
| SPAC19A8.05c | 0.498246529 | 2.19427E-09 | 2.43322E-08 | sst4 |
| SPCC1235.01 | 0.497587626 | 1.96554E-25 | 7.38812E-24 | SPCC1235.01 |
| SPAC17G8.04c | 0.495132566 | 3.675E-15 | 7.08928E-14 | arc5 |
| SPCC1281.08 | 0.491324819 | 0.007384163 | 0.025722554 | wtf11 |
| SPCC417.11c | 0.490489876 | 7.47821E-05 | 0.000420557 | SPCC417.11c |
| SPBC947.15c | 0.489727137 | 1.20339E-15 | 2.43152E-14 | nde1 |
| SPNCRNA.622 | 0.489032111 | 0.001803422 | 0.007419667 | sty1-antisense-1 |
| SPCC364.02c | 0.487362546 | 3.68448E-05 | 0.000219203 | bis1 |
| SPNCRNA.1115 | 0.485415908 | 4.56192E-11 | 5.96433E-10 | SPNCRNA.1115 |
| SPCC1020.03 | 0.484463302 | 0.000594867 | 0.002767026 | mmt1 |
| SPAC1D4.11c | 0.481609375 | 3.19552E-09 | 3.49049E-08 | lkh1 |
| SPBPB2B2.10c | 0.481215378 | 7.44107E-05 | 0.000419391 | gal7 |
| SPBC16G5.09 | 0.480560484 | 2.42796E-10 | 2.96932E-09 | SPBC16G5.09 |
| SPBC1271.01c | 0.47704398 | 0.000688263 | 0.003158349 | pof13 |
| SPAC212.04c | 0.476085562 | 0.007081841 | 0.024762223 | SPAC212.04c |
| SPBPB7E8.01 | 0.475348103 | 1.27179E-18 | 3.17141E-17 | SPBPB7E8.01 |
| SPBC216.03 | 0.471806829 | 1.83241E-09 | 2.06783E-08 | SPBC216.03 |
| SPCC1183.11 | 0.471757245 | 1.21943E-10 | 1.53456E-09 | msy1 |
| SPAC3F10.19 | 0.469204025 | 0.013365211 | 0.042137073 | spd2 |
| SPAPB24D3.08c | 0.46782264 | 1.59591E-11 | 2.22297E-10 | SPAPB24D3.08c |
| SPAC23C4.16c | 0.467282781 | 1.47361E-09 | 1.67775E-08 | atg15 |
| SPBC32F12.16 | 0.465875154 | 0.011610887 | 0.037518871 | gem7 |

| | | | | |
|---|---|---|---|---|
| SPAC22F8.05 | 0.464728363 | 2.69753E-15 | 5.32423E-14 | SPAC22F8.05 |
| SPAC29E6.08 | 0.463879495 | 3.40224E-15 | 6.61303E-14 | tbp1 |
| SPAC25B8.10 | 0.463777443 | 5.59702E-09 | 5.91157E-08 | SPAC25B8.10 |
| SPCC162.08c | 0.462092167 | 0.00422654 | 0.015690684 | nup211 |
| SPCC24B10.03 | 0.460382385 | 0.011915648 | 0.038430784 | SPCC24B10.03 |
| SPCC330.21 | 0.458345301 | 0.001426307 | 0.006010949 | new25 |
| SPAC11G7.03 | 0.457482635 | 6.24472E-13 | 1.00387E-11 | idh1 |
| SPAC4G9.11c | 0.457228863 | 1.76592E-09 | 1.9972E-08 | cmb1 |
| SPBC26H8.03 | 0.456059423 | 6.4328E-18 | 1.52951E-16 | cho2 |
| SPAC2C4.17c | 0.45502322 | 1.99195E-08 | 1.94701E-07 | msy2 |
| SPBC1685.14c | 0.454029851 | 8.48848E-07 | 6.7696E-06 | SPBC1685.14c |
| SPBC16C6.04 | 0.450855103 | 0.000153281 | 0.000817928 | dbl6 |
| SPAPB1A10.10c | 0.450726217 | 1.57744E-06 | 1.19642E-05 | ypt71 |
| SPAC688.02c | 0.448036799 | 0.001006447 | 0.004431486 | mis14 |
| SPCC1682.11c | 0.446044539 | 7.82677E-07 | 6.26142E-06 | ctl1 |
| SPBC211.07c | 0.445650745 | 1.38419E-06 | 1.06566E-05 | ubc8 |
| SPBC23E6.09 | 0.445628937 | 9.81898E-12 | 1.39818E-10 | ssn6 |
| SPAC12B10.13 | 0.445405161 | 2.74916E-06 | 2.00767E-05 | gid8 |
| SPBC902.05c | 0.445250549 | 1.07508E-11 | 1.52661E-10 | idh2 |
| SPNCRNA.1133 | 0.444246668 | 0.004138846 | 0.015432372 | SPNCRNA.1133 |
| SPCC31H12.02c | 0.44367574 | 0.010826949 | 0.035195733 | mug73 |
| SPBC18H10.05 | 0.443247759 | 0.000241555 | 0.001241034 | SPBC18H10.05 |
| SPBC8D2.04 | 0.441292253 | 5.02918E-16 | 1.06677E-14 | hht2 |
| SPNCRNA.58 | 0.441008088 | 0.001801129 | 0.007419317 | prl58 |
| SPCC1020.05 | 0.440667775 | 1.53185E-08 | 1.52946E-07 | SPCC1020.05 |
| SPCC757.05c | 0.439777169 | 2.56835E-08 | 2.49135E-07 | SPCC757.05c |
| SPBC21B10.11 | 0.437013646 | 0.014819566 | 0.045985132 | dpm2 |
| SPCC550.07 | 0.436650275 | 2.80541E-05 | 0.000169719 | SPCC550.07 |
| SPBCPT2R1.04c | 0.436350249 | 0.011095995 | 0.036014429 | SPBCPT2R1.04c |
| SPCC1919.06c | 0.436251916 | 0.002984739 | 0.011489445 | wtf25 |
| SPAC18G6.01c | 0.43557381 | 1.25097E-05 | 8.17768E-05 | SPAC18G6.01c |
| SPBC660.09 | 0.43439007 | 4.58632E-06 | 3.22051E-05 | mug168 |
| SPAC4G9.20c | 0.433294029 | 3.55431E-08 | 3.40893E-07 | SPAC4G9.20c |

| | | | | |
|---|---|---|---|---|
| SPBC1271.03c | 0.432799371 | 2.55039E-06 | 1.87053E-05 | SPBC1271.03c |
| SPCC285.09c | 0.432447694 | 4.77462E-08 | 4.45399E-07 | cgs2 |
| SPAC23A1.14c | 0.431478569 | 1.93987E-10 | 2.38954E-09 | SPAC23A1.14c |
| SPAC1834.03c | 0.430743267 | 6.17412E-19 | 1.57025E-17 | hhf1 |
| SPAPB24D3.02c | 0.429833147 | 2.01487E-05 | 0.000125763 | SPAPB24D3.02c |
| SPBC2G5.06c | 0.429510209 | 1.67565E-18 | 4.15822E-17 | hmt2 |
| SPCP31B10.06 | 0.428106751 | 1.68134E-08 | 1.66247E-07 | mug190 |
| SPAC227.04 | 0.426302311 | 0.002910103 | 0.01124448 | atg10 |
| SPAC1687.12c | 0.42598996 | 3.56406E-08 | 3.41189E-07 | coq4 |
| SPCC18B5.11c | 0.42570103 | 6.44499E-05 | 0.000366891 | cds1 |
| SPBC800.14c | 0.423570605 | 0.000553102 | 0.002603552 | SPBC800.14c |
| SPCC576.06c | 0.423133775 | 0.000625538 | 0.00289389 | SPCC576.06c |
| SPBC428.10 | 0.422364993 | 5.48931E-09 | 5.82186E-08 | SPBC428.10 |
| SPCC757.07c | 0.419948662 | 2.76688E-13 | 4.54801E-12 | ctt1 |
| SPAC10F6.11c | 0.419735741 | 8.27008E-06 | 5.57739E-05 | atg17 |
| SPBC1685.13 | 0.417836715 | 2.04157E-12 | 3.13409E-11 | fhn1 |
| SPBC1539.07c | 0.417686893 | 1.11586E-10 | 1.40847E-09 | fmd1 |
| SPAC1486.01 | 0.417275278 | 1.71914E-10 | 2.1279E-09 | SPAC1486.01 |
| SPCC548.03c | 0.416903843 | 0.015497192 | 0.047752651 | wtf4 |
| SPCC1393.08 | 0.416885242 | 2.71679E-11 | 3.66444E-10 | SPCC1393.08 |
| SPBC359.03c | 0.416820712 | 2.49191E-07 | 2.13735E-06 | aat1 |
| SPBC2F12.17 | 0.415118103 | 1.96859E-06 | 1.47342E-05 | cox7 |
| SPAC25G10.03 | 0.414945186 | 1.53054E-14 | 2.8246E-13 | zip1 |
| SPBC17D1.17 | 0.414632167 | 3.64218E-09 | 3.95305E-08 | tam11 |
| SPCC576.04 | 0.414239537 | 1.38377E-07 | 1.21753E-06 | bxi1 |
| SPBC8D2.18c | 0.414005897 | 1.24058E-17 | 2.86962E-16 | SPBC8D2.18c |
| SPBC713.11c | 0.413233883 | 1.39142E-15 | 2.80037E-14 | pmp3 |
| SPBPB10D8.02c | 0.412366395 | 0.004927584 | 0.018083138 | SPBPB10D8.02c |
| SPBC12C2.12c | 0.41228865 | 2.12726E-09 | 2.37436E-08 | glo1 |
| SPBC14F5.05c | 0.411920815 | 1.87164E-18 | 4.62213E-17 | sam1 |
| SPAC227.17c | 0.409760034 | 1.17816E-05 | 7.74131E-05 | SPAC227.17c |
| SPAC57A10.14 | 0.409237136 | 0.000120577 | 0.00065434 | sgf11 |
| SPBC1778.06c | 0.408239944 | 1.55784E-16 | 3.38879E-15 | fim1 |

| | | | | |
|---|---|---|---|---|
| SPAC6C3.08 | 0.408233805 | 0.000438255 | 0.00210165 | nas6 |
| SPCC663.06c | 0.407428597 | 0.002330464 | 0.009285526 | osr1 |
| SPBC609.03 | 0.407132766 | 9.73195E-07 | 7.71313E-06 | iqw1 |
| SPBP8B7.13 | 0.406698401 | 3.74917E-06 | 2.67678E-05 | vac7 |
| SPAC1039.11c | 0.404350243 | 2.61382E-05 | 0.000158692 | gto1 |
| SPBC577.09 | 0.403940212 | 0.00036541 | 0.001790964 | ckn1 |
| SPAC6G9.16c | 0.402401157 | 0.004302894 | 0.015939415 | xrc4 |
| SPBC1271.05c | 0.401242285 | 6.85449E-06 | 4.68451E-05 | SPBC1271.05c |
| SPBP16F5.08c | 0.401017522 | 3.63913E-10 | 4.40722E-09 | SPBP16F5.08c |
| SPAC25B8.18 | 0.399957717 | 0.009332056 | 0.031200439 | SPAC25B8.18 |
| SPCC16C4.07 | 0.399244511 | 8.09811E-07 | 6.46836E-06 | scw1 |
| SPNCRNA.1215 | 0.399187765 | 0.00341917 | 0.013024441 | SPCC417.03-antisense-1 |
| SPAC23C4.13 | 0.398475253 | 3.24644E-05 | 0.000194558 | bet1 |
| SPCPJ732.02c | 0.397061053 | 5.62083E-08 | 5.20538E-07 | xks1 |
| SPAC22G7.08 | 0.396739924 | 0.000233705 | 0.001204333 | ppk8 |
| SPAC57A10.09c | 0.394870524 | 4.80051E-11 | 6.26026E-10 | nhp6 |
| SPBC23G7.16 | 0.394827581 | 1.71885E-07 | 1.49945E-06 | ctr6 |
| SPAC30D11.01c | 0.394792903 | 3.59216E-05 | 0.000214272 | gto2 |
| SPAC1006.01 | 0.393752516 | 7.59882E-10 | 8.88906E-09 | psp3 |
| SPCC61.03 | 0.393125714 | 1.0988E-06 | 8.6284E-06 | SPCC61.03 |
| SPAC19G12.10c | 0.392804022 | 6.58305E-16 | 1.37357E-14 | cpy1 |
| SPAC458.06 | 0.392303676 | 0.011706637 | 0.037804376 | atg1803 |
| SPBC15D4.15 | 0.392164985 | 3.24187E-10 | 3.93645E-09 | pho2 |
| SPAC1B3.07c | 0.390839089 | 0.000297952 | 0.001493265 | vps28 |
| SPAC4G8.13c | 0.390573465 | 2.29685E-08 | 2.23648E-07 | prz1 |
| SPBC1709.12 | 0.390491795 | 6.65024E-07 | 5.3659E-06 | rid1 |
| SPAC19G12.07c | 0.389768269 | 3.88798E-07 | 3.25292E-06 | rsd1 |
| SPBC4B4.10c | 0.389618049 | 9.37478E-05 | 0.000518657 | atg5 |
| SPAC4F10.02 | 0.389478947 | 1.65909E-07 | 1.44979E-06 | aap1 |
| SPNCRNA.833 | 0.388218816 | 0.010320078 | 0.033796438 | SPNCRNA.833 |
| SPCP1E11.03 | 0.387672427 | 0.006056474 | 0.021696353 | mug170 |
| SPAC222.18 | 0.387419798 | 0.00217664 | 0.008733897 | SPAC222.18 |
| SPBC21B10.07 | 0.387286468 | 3.44955E-09 | 3.75193E-08 | SPBC21B10.07 |

| | | | | |
|---|---|---|---|---|
| SPBC1289.05c | 0.385371563 | 2.44745E-06 | 1.80279E-05 | vma10 |
| SPAC27D7.09c | 0.384867956 | 1.28953E-05 | 8.41898E-05 | SPAC27D7.09c |
| SPAC13G7.07 | 0.383239514 | 0.002832589 | 0.011011556 | arb2 |
| SPBC6B1.09c | 0.381718939 | 0.00015682 | 0.000834614 | nbs1 |
| SPAC13G7.03 | 0.377989503 | 0.000507548 | 0.002409088 | upf3 |
| SPAC521.04c | 0.377973334 | 1.15512E-05 | 7.5997E-05 | sst1 |
| SPAC29B12.10c | 0.37789203 | 8.04351E-06 | 5.43894E-05 | pgt1 |
| SPAC3G9.11c | 0.37771113 | 0.008898977 | 0.030007631 | pdc201 |
| SPBC106.03 | 0.375990816 | 8.20867E-10 | 9.55871E-09 | SPBC106.03 |
| SPBC1198.13c | 0.375651494 | 3.91275E-06 | 2.78579E-05 | tfg2 |
| SPCC285.11 | 0.375039849 | 7.59824E-08 | 6.86258E-07 | dsc5 |
| SPAC17A2.09c | 0.375004492 | 4.68194E-07 | 3.84792E-06 | csx1 |
| SPCC1322.14c | 0.374456207 | 4.98831E-06 | 3.49318E-05 | vtc4 |
| SPBC215.10 | 0.373118306 | 5.32545E-06 | 3.71401E-05 | SPBC215.10 |
| SPBC36.04 | 0.372974175 | 8.99823E-11 | 1.14425E-09 | cys11 |
| SPCC63.08c | 0.372246567 | 1.64997E-05 | 0.000105431 | atg1 |
| SPCC777.09c | 0.370274065 | 1.18099E-13 | 2.01913E-12 | arg1 |
| SPBC4F6.16c | 0.367791226 | 5.30251E-08 | 4.92844E-07 | ero11 |
| SPAC17G8.11c | 0.366127238 | 3.53204E-05 | 0.000210932 | imt3 |
| SPAC23G3.02c | 0.366015748 | 7.14613E-06 | 4.85785E-05 | sib1 |
| SPAC1805.15c | 0.365912559 | 0.000853296 | 0.003836456 | pub2 |
| SPBC15C4.06c | 0.365796621 | 0.000560322 | 0.00262545 | SPBC15C4.06c |
| SPCP31B10.02 | 0.365056291 | 0.003817376 | 0.014348844 | SPCP31B10.02 |
| SPCC777.17c | 0.361581719 | 0.005613273 | 0.020293532 | SPCC777.17c |
| SPBC1709.11c | 0.360557198 | 4.29413E-05 | 0.000252317 | png2 |
| SPAC16E8.17c | 0.36032862 | 1.85627E-06 | 1.39139E-05 | SPAC16E8.17c |
| SPCC645.14c | 0.360296677 | 8.2884E-12 | 1.19018E-10 | sti1 |
| SPBC409.23 | 0.35962171 | 0.007934058 | 0.02725296 | tam7 |
| SPCC1682.08c | 0.359458843 | 1.66846E-05 | 0.000106216 | mcp2 |
| SPAC688.16 | 0.359412787 | 0.001501323 | 0.00630112 | SPAC688.16 |
| SPBP35G2.11c | 0.358814439 | 8.26666E-05 | 0.000462354 | nbr1 |
| SPAC3A12.18 | 0.35841227 | 1.23445E-11 | 1.74806E-10 | zwf1 |
| SPCC1020.10 | 0.3582779 | 1.20607E-05 | 7.91148E-05 | oca2 |

| | | | | |
|---|---|---|---|---|
| SPAPJ691.03 | 0.357841221 | 8.64701E-05 | 0.00048152 | SPAPJ691.03 |
| SPAC9.13c | 0.357278867 | 0.00035011 | 0.001727572 | cwf16 |
| SPAC11D3.05 | 0.355908185 | 0.007618932 | 0.026423326 | mfs2 |
| SPBC119.05c | 0.355722289 | 6.64347E-07 | 5.3659E-06 | lsb1 |
| SPBP23A10.04 | 0.355418693 | 0.002618866 | 0.010314054 | apc2 |
| SPAC17C9.11c | 0.355414832 | 3.93969E-05 | 0.00023256 | SPAC17C9.11c |
| SPAC2C4.09 | 0.354656568 | 0.001118419 | 0.004853446 | SPAC2C4.09 |
| SPAC6F12.04 | 0.353978467 | 2.31212E-08 | 2.24706E-07 | tvp15 |
| SPBC17D11.08 | 0.3533877 | 3.3824E-07 | 2.85799E-06 | SPBC17D11.08 |
| SPAC24C9.12c | 0.353182178 | 3.46442E-15 | 6.70838E-14 | shm1 |
| SPBC3E7.12c | 0.352807181 | 7.89797E-09 | 8.15645E-08 | cfh4 |
| SPBC887.07 | 0.352766153 | 0.00077257 | 0.00351681 | mrpl38 |
| SPAC1687.14c | 0.35230576 | 0.002698084 | 0.010585267 | SPAC1687.14c |
| SPCC330.06c | 0.34998062 | 5.38491E-08 | 4.99594E-07 | pmp20 |
| SPBC1711.12 | 0.349372777 | 4.45759E-07 | 3.69322E-06 | SPBC1711.12 |
| SPBC660.16 | 0.348883643 | 6.04978E-14 | 1.05192E-12 | SPBC660.16 |
| SPBC3E7.06c | 0.347116257 | 2.08635E-05 | 0.000129908 | fnx2 |
| SPCC126.04c | 0.34701603 | 0.000835326 | 0.003768922 | sgf73 |
| SPBC651.03c | 0.346898947 | 0.000101302 | 0.000556832 | gyp10 |
| SPBC12D12.01 | 0.344117764 | 1.64273E-05 | 0.000105102 | sad1 |
| SPBC18E5.01 | 0.343814227 | 9.52692E-08 | 8.49941E-07 | SPBC18E5.01 |
| SPCC645.03c | 0.343780828 | 1.58081E-07 | 1.38613E-06 | isa1 |
| SPBC83.13 | 0.343059258 | 4.35198E-09 | 4.66401E-08 | SPBC83.13 |
| SPCC1442.08c | 0.34264858 | 9.96176E-06 | 6.61357E-05 | cox12 |
| SPAC9E9.15 | 0.341184604 | 0.000787856 | 0.003573664 | SPAC9E9.15 |
| SPBC1703.13c | 0.340311775 | 7.30059E-08 | 6.61713E-07 | SPBC1703.13c |
| SPAC23D3.07 | 0.34023122 | 6.46501E-08 | 5.92278E-07 | pup1 |
| SPBC21B10.02 | 0.338694578 | 0.000548381 | 0.002586092 | SPBC21B10.02 |
| SPBC21B10.04c | 0.338387852 | 3.07979E-08 | 2.97616E-07 | nrf1 |
| SPAC2G11.04 | 0.33702673 | 0.008268707 | 0.028255102 | SPAC2G11.04 |
| SPBC21B10.15 | 0.335295644 | 0.0076279 | 0.026436492 | SPBC21B10.15 |
| SPBC17G9.06c | 0.334566363 | 3.7227E-08 | 3.55045E-07 | SPBC17G9.06c |
| SPBP8B7.26 | 0.334237599 | 2.43897E-05 | 0.000150217 | SPBP8B7.26 |

| | | | | |
|---|---|---|---|---|
| SPBC14C8.01c | 0.333918774 | 0.003740423 | 0.014069937 | cut2 |
| SPBC19C7.10 | 0.332224488 | 1.84656E-08 | 1.81881E-07 | bqt4 |
| SPBP35G2.05c | 0.332210776 | 3.27111E-06 | 2.36855E-05 | cki2 |
| SPCC4G3.13c | 0.332006417 | 6.7041E-05 | 0.00038037 | cue1 |
| SPBC16G5.03 | 0.331472638 | 0.001170606 | 0.005024466 | SPBC16G5.03 |
| SPCC70.02c | 0.331331517 | 6.16496E-05 | 0.000351733 | SPCC70.02c |
| SPAC1527.02 | 0.331005126 | 0.000257846 | 0.001316792 | sft2 |
| SPAC227.06 | 0.330682216 | 1.10573E-05 | 7.29354E-05 | yip5 |
| SPCC1739.09c | 0.328924149 | 3.4942E-07 | 2.94272E-06 | cox13 |
| SPBC776.05 | 0.327567631 | 0.002565144 | 0.010141543 | SPBC776.05 |
| SPBC1703.12 | 0.324053204 | 6.41623E-05 | 0.000365661 | ubp9 |
| SPBP8B7.07c | 0.323022744 | 0.001106364 | 0.004805209 | set6 |
| SPAC630.11 | 0.320429929 | 3.70538E-05 | 0.000219999 | vps55 |
| SPBC29A10.13 | 0.320423555 | 0.000610694 | 0.002835485 | atp7 |
| SPBC1711.08 | 0.320234217 | 4.51243E-08 | 4.24817E-07 | aha1 |
| SPBC409.13 | 0.319794239 | 1.53429E-05 | 9.89066E-05 | rib4 |
| SPBC31F10.15c | 0.319425309 | 2.32953E-05 | 0.000144346 | atp15 |
| SPBC1711.18 | 0.318578413 | 0.012267167 | 0.039354801 | tam9 |
| SPAC1639.02c | 0.318107782 | 0.000129895 | 0.000701186 | trk2 |
| SPAC12B10.14c | 0.318046319 | 5.06417E-05 | 0.000294852 | tea5 |
| SPBC839.16 | 0.317711601 | 5.80944E-09 | 6.11067E-08 | thf1 |
| SPAC17D4.01 | 0.31724991 | 1.92863E-10 | 2.38144E-09 | pex7 |
| SPBC947.01 | 0.317199367 | 9.46768E-05 | 0.000522665 | knk1 |
| SPAC139.04c | 0.316543143 | 0.001033828 | 0.004520897 | fap2 |
| SPAC23H4.06 | 0.315314314 | 6.2154E-06 | 4.27003E-05 | gln1 |
| SPAC2F3.17c | 0.315018644 | 0.003226171 | 0.012326 | lsm6 |
| SPAC29E6.05c | 0.31349012 | 0.002802275 | 0.01092695 | mxr1 |
| SPBC21C3.11 | 0.313309557 | 0.001364632 | 0.005765289 | ubx4 |
| SPBC1D7.05 | 0.312921972 | 0.000888228 | 0.003972546 | byr2 |
| SPAC25A8.02 | 0.312568896 | 0.00105915 | 0.004619776 | atg14 |
| SPAC17G8.10c | 0.310691967 | 0.000827501 | 0.003736911 | dma1 |
| SPAC1399.02 | 0.310359863 | 0.005525977 | 0.020048825 | SPAC1399.02 |
| SPCC70.12c | 0.309642526 | 0.007555273 | 0.026248471 | ecl1 |

| | | | | |
|---|---|---|---|---|
| SPAC5D6.08c | 0.308531688 | 0.009647109 | 0.031940428 | mes1 |
| SPCC61.05 | 0.307900294 | 0.000385577 | 0.001879001 | SPCC61.05 |
| SPBC577.10 | 0.30789745 | 1.32893E-06 | 1.02466E-05 | pre4 |
| SPCC132.03 | 0.307107981 | 0.000974529 | 0.00430851 | SPCC132.03 |
| SPBC839.17c | 0.306781306 | 5.85431E-08 | 5.40816E-07 | fkh1 |
| SPBC800.10c | 0.306439345 | 8.95426E-06 | 5.99923E-05 | SPBC800.10c |
| SPBC29A3.21 | 0.305988746 | 0.000975795 | 0.00430851 | SPBC29A3.21 |
| SPCC663.18 | 0.301819595 | 1.05037E-06 | 8.27348E-06 | SPCC663.18 |
| SPCC1235.16 | 0.301179124 | 0.01506991 | 0.046661041 | vma21 |
| SPBC36B7.02 | 0.300812765 | 8.67049E-05 | 0.000482302 | SPBC36B7.02 |
| SPCC16C4.10 | 0.300127418 | 3.34397E-07 | 2.83489E-06 | SPCC16C4.10 |
| SPAC1296.02 | 0.299851393 | 2.4178E-06 | 1.78352E-05 | cox4 |
| SPCC24B10.02c | 0.299764614 | 0.001025408 | 0.004495615 | SPCC24B10.02c |
| SPAC5D6.06c | 0.299386956 | 0.005977381 | 0.021458127 | alg14 |
| SPAC806.04c | 0.299319714 | 1.49869E-05 | 9.71017E-05 | SPAC806.04c |
| SPAC2F3.18c | 0.298589555 | 0.000110101 | 0.000600677 | SPAC2F3.18c |
| SPBC32H8.02c | 0.29830337 | 0.00232076 | 0.009254077 | nep2 |
| SPCC320.05 | 0.297891581 | 0.000665212 | 0.003060816 | SPCC320.05 |
| SPAC7D4.05 | 0.297193761 | 0.008920281 | 0.030059643 | SPAC7D4.05 |
| SPAC22F8.02c | 0.295381625 | 0.002728491 | 0.010671804 | pvg5 |
| SPCC594.04c | 0.294207428 | 0.001145938 | 0.004945232 | SPCC594.04c |
| SPCC1183.09c | 0.293952447 | 1.52E-05 | 9.83573E-05 | pmp31 |
| SPCC5E4.05c | 0.293507911 | 0.001915217 | 0.007832469 | mgl1 |
| SPAC56E4.02c | 0.292878714 | 0.014197606 | 0.044333514 | alg13 |
| SPAC1B3.14 | 0.292865392 | 1.32749E-06 | 1.02466E-05 | vma3 |
| SPAC1002.05c | 0.291319547 | 0.009680477 | 0.032009442 | jmj2 |
| SPAC1F3.03 | 0.290445618 | 0.000393508 | 0.001908551 | sro7 |
| SPAC13C5.05c | 0.290430283 | 2.49469E-05 | 0.000152912 | SPAC13C5.05c |
| SPAC22H10.07 | 0.290366628 | 0.003150174 | 0.01207173 | scd2 |
| SPAC23C4.05c | 0.290193531 | 8.61586E-06 | 5.79217E-05 | SPAC23C4.05c |
| SPAC12G12.11c | 0.289389507 | 0.000726764 | 0.003320124 | SPAC12G12.11c |
| SPAC1834.04 | 0.289274478 | 1.9195E-09 | 2.15659E-08 | hht1 |
| SPCC736.05 | 0.289221905 | 0.010260962 | 0.03364595 | wtf7 |

| | | | | |
|---|---|---|---|---|
| SPAC22F8.06 | 0.289000856 | 5.43341E-06 | 3.78414E-05 | pam1 |
| SPBP19A11.07c | 0.288978402 | 0.000844184 | 0.003804013 | SPBP19A11.07c |
| SPAC6C3.04 | 0.288347377 | 2.00492E-06 | 1.49623E-05 | cit1 |
| SPAC6G10.03c | 0.288056219 | 0.000281046 | 0.00141827 | cld1 |
| SPAC11E3.04c | 0.287965041 | 4.30694E-05 | 0.000252686 | ubc13 |
| SPAC4G9.13c | 0.286238287 | 0.004117011 | 0.015373381 | vps26 |
| SPAC922.04 | 0.285924975 | 1.06137E-08 | 1.08082E-07 | SPAC922.04 |
| SPCPB16A4.02c | 0.284383694 | 0.000156185 | 0.000832552 | SPCPB16A4.02c |
| SPCC126.07c | 0.284264688 | 0.01022086 | 0.033546959 | pbr1 |
| SPAC22F3.09c | 0.283896693 | 0.002860833 | 0.011079227 | res2 |
| SPAC1687.07 | 0.283545505 | 0.009520463 | 0.031658103 | SPAC1687.07 |
| SPAC4G9.10 | 0.282742657 | 3.1656E-05 | 0.000190159 | arg3 |
| SPAC22F3.13 | 0.281792873 | 0.000336588 | 0.001670524 | tsc1 |
| SPAC11E3.10 | 0.281443512 | 7.24485E-05 | 0.000408782 | SPAC11E3.10 |
| SPBC337.15c | 0.280919662 | 0.000143152 | 0.000767079 | coq7 |
| SPBC337.12 | 0.280254579 | 0.000398128 | 0.001925478 | red5 |
| SPCC1739.13 | 0.279354422 | 1.23106E-09 | 1.4142E-08 | ssa2 |
| SPCC569.01c | 0.278867964 | 0.001045636 | 0.004568626 | SPCC569.01c |
| SPBC26H8.05c | 0.277849068 | 0.001992023 | 0.008101209 | pph3 |
| SPBC19G7.18c | 0.276383268 | 0.016126682 | 0.049158974 | SPBC19G7.18c |
| SPBC24C6.03 | 0.276368155 | 0.000624393 | 0.002891209 | SPBC24C6.03 |
| SPCC622.11 | 0.27599766 | 0.0034598 | 0.013165337 | SPCC622.11 |
| SPAPB1A10.08 | 0.27580944 | 0.001800775 | 0.007419317 | SPAPB1A10.08 |
| SPCC1620.08 | 0.27514384 | 3.79235E-05 | 0.000224901 | SPCC1620.08 |
| SPCC18.16c | 0.274758599 | 0.012635116 | 0.040343981 | fmn1 |
| SPBC337.07c | 0.274529333 | 5.64007E-05 | 0.000325418 | ecm14 |
| SPBC887.13c | 0.27419104 | 0.004171225 | 0.015535686 | cem1 |
| SPAC23C11.08 | 0.274169186 | 0.001023001 | 0.00448891 | php3 |
| SPBC36.08c | 0.273859774 | 0.008782962 | 0.029635975 | cog2 |
| SPAC8E11.04c | 0.272741723 | 0.001848636 | 0.007584452 | SPAC8E11.04c |
| SPBC8D2.01 | 0.271874197 | 0.000344608 | 0.001702064 | gsk31 |
| SPBC16E9.02c | 0.271176405 | 3.44612E-06 | 2.4847E-05 | SPBC16E9.02c |
| SPBC14C8.02 | 0.27044723 | 0.001947725 | 0.007940007 | tim44 |

| | | | | |
|---|---|---|---|---|
| SPAC22E12.09c | 0.269653244 | 2.77301E-05 | 0.000168157 | krp1 |
| SPAC14C4.07 | 0.26939786 | 2.31455E-05 | 0.000143745 | SPAC14C4.07 |
| SPAPB15E9.01c | 0.26845885 | 9.41403E-08 | 8.41338E-07 | pfl2 |
| SPAC1039.03 | 0.268085449 | 0.001844255 | 0.007572556 | SPAC1039.03 |
| SPBC16C6.08c | 0.267816827 | 4.12842E-06 | 2.92711E-05 | qcr6 |
| SPBC1198.06c | 0.267356599 | 0.00042112 | 0.002025181 | SPBC1198.06c |
| SPAC1834.11c | 0.267294199 | 3.07649E-06 | 2.23713E-05 | sec18 |
| SPCC23B6.01c | 0.266043079 | 6.10118E-07 | 4.95855E-06 | SPCC23B6.01c |
| SPBC1703.10 | 0.264256362 | 1.22776E-05 | 8.03626E-05 | ypt1 |
| SPAC1B3.06c | 0.263251273 | 0.000966388 | 0.004277208 | SPAC1B3.06c |
