## Supplemental Table 5 for "Rapalink-1 reveals novel mTOR-dependent genes and an agmatinergic axis-based metabolic feedback regulating mTOR activity and lifespan"

**Supplementary Table 5**

**Rapalink-1 downregulated genes**

| gene_id | log2FoldChange | pvalue | padj | gene_name |
|---|---|---|---|---|
| SPNCRNA.189 | -1.854422844 | 0.005725908 | 0.020656912 | SPNCRNA.189 |
| SPNCRNA.1342 | -1.197127447 | 0.014114848 | 0.044185611 | SPNCRNA.1342 |
| SPAC869.11 | -1.176624212 | 6.0188E-40 | 4.15785E-38 | cat1 |
| SPBC1773.12 | -1.117788986 | 0.006425288 | 0.022746589 | SPBC1773.12 |
| SPAC17A2.01 | -0.981181565 | 9.23132E-41 | 6.64655E-39 | bsu1 |
| SPNCRNA.630 | -0.832823418 | 2.49001E-05 | 0.000152854 | SPNCRNA.630 |
| SPRRNA.46 | -0.826427656 | 0.002920607 | 0.011268032 | SPRRNA.46 |
| SPAC1B3.16c | -0.799537495 | 3.22376E-36 | 1.87271E-34 | vht1 |
| SPBC1685.07c | -0.782958507 | 3.15825E-07 | 2.68189E-06 | avt5 |
| SPNCRNA.779 | -0.774508344 | 0.000363639 | 0.001785706 | SPNCRNA.779 |
| SPCC31H12.06 | -0.726820569 | 2.16919E-09 | 2.41588E-08 | mug111 |
| SPAC977.12 | -0.712250674 | 1.13678E-14 | 2.12088E-13 | SPAC977.12 |
| SPAC8E11.10 | -0.71150469 | 1.09265E-28 | 4.54118E-27 | SPAC8E11.10 |
| SPAC186.05c | -0.689828979 | 9.3562E-07 | 7.43839E-06 | SPAC186.05c |
| SPAP8A3.13c | -0.678221527 | 0.009717129 | 0.032081049 | SPAP8A3.13c |
| SPAC23A1.03 | -0.66351469 | 8.93988E-28 | 3.65605E-26 | apt1 |
| SPCC126.03 | -0.653573119 | 3.87505E-15 | 7.4471E-14 | pus1 |
| SPNCRNA.1442 | -0.653379073 | 0.002050141 | 0.008317715 | SPNCRNA.1442 |
| SPBC530.07c | -0.651531731 | 1.73109E-26 | 6.75521E-25 | SPBC530.07c |
| SPCC1223.13 | -0.649553483 | 1.36568E-10 | 1.69863E-09 | cbf12 |
| SPBC1778.01c | -0.628610266 | 1.51567E-34 | 8.33131E-33 | zuo1 |
| SPBC1718.03 | -0.606577513 | 1.01613E-10 | 1.28895E-09 | ker1 |
| SPBPB8B6.05c | -0.601258848 | 1.72737E-08 | 1.7047E-07 | SPBPB8B6.05c |
| SPAC26H5.12 | -0.596298763 | 8.29828E-09 | 8.53538E-08 | rpo41 |
| SPAC4G8.07c | -0.596157969 | 3.98831E-08 | 3.79669E-07 | trm2 |
| SPAPB2B4.03 | -0.595602907 | 3.6593E-05 | 0.000218023 | cig2 |
| SPBC1198.02 | -0.587473279 | 4.22933E-20 | 1.16238E-18 | dea2 |
| SPBC36.01c | -0.58183789 | 7.10737E-09 | 7.38473E-08 | SPBC36.01c |
| SPCC74.04 | -0.573144633 | 1.45415E-06 | 1.11616E-05 | SPCC74.04 |
| SPBC23G7.07c | -0.572299343 | 1.2211E-08 | 1.22879E-07 | cms1 |
| SPAC1783.03 | -0.570545114 | 0.004004946 | 0.014998744 | fta2 |

| | | | | |
|---|---|---|---|---|
| SPNCRNA.1261 | -0.565929229 | 0.013551982 | 0.042647391 | rpl20-antisense-1 |
| SPAC57A10.10c | -0.561845772 | 1.60813E-14 | 2.95711E-13 | sla1 |
| SPBC11C11.09c | -0.552456765 | 8.03968E-29 | 3.3966E-27 | rpl502 |
| SPAC1093.01 | -0.547762077 | 2.21212E-13 | 3.67155E-12 | ppr5 |
| SPAC2F7.14c | -0.547250764 | 7.9976E-09 | 8.24269E-08 | rrp4 |
| SPAC57A10.12c | -0.545595306 | 4.74752E-26 | 1.81114E-24 | ura3 |
| SPAC19G12.05 | -0.540798921 | 2.09668E-15 | 4.17051E-14 | SPAC19G12.05 |
| SPAC57A7.12 | -0.537438252 | 8.93338E-24 | 3.10663E-22 | ssz1 |
| SPAP27G11.13c | -0.537220584 | 9.30683E-12 | 1.32895E-10 | nop10 |
| SPCP1E11.10 | -0.535145487 | 2.49075E-05 | 0.000152854 | SPCP1E11.10 |
| SPBC16H5.08c | -0.530539848 | 6.2847E-18 | 1.50128E-16 | SPBC16H5.08c |
| SPBC21C3.08c | -0.528693773 | 1.56346E-26 | 6.14801E-25 | car2 |
| SPAC1782.10c | -0.527937044 | 5.69459E-19 | 1.45554E-17 | nhp2 |
| SPBC8D2.16c | -0.52721537 | 1.52708E-05 | 9.86908E-05 | SPBC8D2.16c |
| SPAC6F12.16c | -0.527147157 | 4.15812E-12 | 6.16125E-11 | mtr4 |
| SPAC1783.05 | -0.522888662 | 1.45564E-05 | 9.47928E-05 | hrp1 |
| SPAC23G3.12c | -0.522034174 | 5.48105E-18 | 1.32166E-16 | SPAC23G3.12c |
| SPBC14F5.08 | -0.51472387 | 0.007755412 | 0.026787611 | med7 |
| SPBC16D10.06 | -0.513952136 | 3.89656E-11 | 5.14708E-10 | zrt1 |
| SPAC1006.07 | -0.509510481 | 2.26047E-27 | 9.09882E-26 | SPAC1006.07 |
| SPAC3A12.19 | -0.508436511 | 0.001603976 | 0.006684423 | SPAC3A12.19 |
| SPBC211.01 | -0.506694821 | 0.002114861 | 0.008526159 | rsm10 |
| SPBC21B10.10 | -0.505510549 | 2.18887E-27 | 8.88057E-26 | rps402 |
| SPBC25B2.09c | -0.504707523 | 3.0163E-22 | 9.63452E-21 | mrs1 |
| SPCC162.02c | -0.504693924 | 4.31598E-13 | 7.04897E-12 | SPCC162.02c |
| SPBC4C3.03 | -0.504182006 | 2.67826E-09 | 2.93804E-08 | thr1 |
| SPBC3F6.01c | -0.504166502 | 4.42275E-12 | 6.51559E-11 | SPBC3F6.01c |
| SPBC4C3.05c | -0.503556656 | 2.16697E-15 | 4.29362E-14 | nuc1 |
| SPBC2D10.10c | -0.497492976 | 7.30639E-22 | 2.26365E-20 | fib1 |
| SPBC8D2.10c | -0.495427221 | 9.26937E-16 | 1.8954E-14 | rmt3 |
| SPBC428.03c | -0.495378742 | 1.31385E-14 | 2.43348E-13 | pho4 |
| SPCC1223.08c | -0.493589226 | 9.30159E-13 | 1.46307E-11 | dfr1 |
| SPAC23G3.06 | -0.491132798 | 4.68982E-21 | 1.39386E-19 | nop58 |

| | | | | |
|---|---|---|---|---|
| SPBC1709.02c | -0.489628991 | 5.91804E-17 | 1.33273E-15 | vrs1 |
| SPBC1861.02 | -0.489336537 | 1.05189E-08 | 1.07545E-07 | abp2 |
| SPAC56F8.03 | -0.485859147 | 3.31402E-20 | 9.20721E-19 | SPAC56F8.03 |
| SPCC4G3.17 | -0.485530845 | 1.66132E-11 | 2.29532E-10 | hdd1 |
| SPAC1B3.17 | -0.48338678 | 1.39104E-05 | 9.07012E-05 | clr2 |
| SPAC3F10.06c | -0.481626377 | 0.005472975 | 0.019892696 | rit1 |
| SPAC3H1.09c | -0.48022028 | 4.34911E-12 | 6.42562E-11 | avt3 |
| SPBPB8B6.06c | -0.479819823 | 0.015869692 | 0.048607469 | SPBPB8B6.06c |
| SPAC17G8.01c | -0.479385722 | 3.82922E-07 | 3.21957E-06 | trl1 |
| SPBC725.15 | -0.47742319 | 1.7223E-11 | 2.37315E-10 | ura5 |
| SPAC4F8.12c | -0.47612605 | 1.54371E-06 | 1.17499E-05 | spp42 |
| SPBC646.10c | -0.473205201 | 3.4518E-21 | 1.03191E-19 | nop56 |
| SPBC23E6.10c | -0.47257767 | 2.51301E-09 | 2.76864E-08 | mri1 |
| SPBC19F5.02c | -0.471226295 | 0.004965539 | 0.018209351 | utp4 |
| SPAC664.14 | -0.470942439 | 0.000963799 | 0.004269445 | amt2 |
| SPAC29B12.07 | -0.469748845 | 8.82505E-05 | 0.000490366 | sec16 |
| SPAC5H10.06c | -0.468664516 | 5.91991E-14 | 1.03639E-12 | adh4 |
| SPAC16C9.07 | -0.466375304 | 8.85926E-09 | 9.07586E-08 | pom2 |
| SPBC20F10.01 | -0.466238028 | 1.1702E-08 | 1.17989E-07 | gar1 |
| SPAC3H1.07 | -0.464734587 | 1.17812E-16 | 2.59593E-15 | aru1 |
| SPAP8A3.07c | -0.464365094 | 1.50544E-07 | 1.3223E-06 | SPAP8A3.07c |
| SPAC664.09 | -0.463397264 | 5.38613E-11 | 6.9883E-10 | ggt1 |
| SPBC30B4.05 | -0.46238198 | 7.74551E-12 | 1.11535E-10 | kap109 |
| SPAC30C2.02 | -0.462238688 | 3.65388E-08 | 3.49134E-07 | mmd1 |
| SPBC216.06c | -0.461513179 | 0.008286881 | 0.028279397 | swi1 |
| SPAC5H10.12c | -0.460946352 | 0.000142683 | 0.000765365 | SPAC5H10.12c |
| SPCC622.16c | -0.460840046 | 0.001851391 | 0.007589664 | epe1 |
| SPAC20H4.01 | -0.460682607 | 0.000275219 | 0.001397141 | utp5 |
| SPBC15C4.05 | -0.459960443 | 2.28117E-09 | 2.5241E-08 | dhx29 |
| SPBC1703.03c | -0.459013074 | 6.71122E-09 | 6.98732E-08 | syo2 |
| SPCC1223.14 | -0.459010633 | 8.86532E-18 | 2.08846E-16 | SPCC1223.14 |
| SPBC409.08 | -0.456848571 | 4.95243E-11 | 6.44194E-10 | SPBC409.08 |
| SPAC3H5.07 | -0.454545007 | 3.05605E-22 | 9.64352E-21 | rpl702 |

| | | | | |
|---|---|---|---|---|
| SPBC2F12.07c | -0.453962154 | 7.26125E-21 | 2.12112E-19 | rpl802 |
| SPCC1223.11 | -0.453672475 | 1.87318E-07 | 1.623E-06 | ptc2 |
| SPAC6B12.02c | -0.451806132 | 0.000665163 | 0.003060816 | mus7 |
| SPAC24B11.09 | -0.45170916 | 7.74828E-06 | 5.25321E-05 | mpc2 |
| SPBC713.03 | -0.451240759 | 3.64682E-10 | 4.40722E-09 | SPBC713.03 |
| SPBP8B7.03c | -0.450708305 | 1.59377E-20 | 4.5013E-19 | rpl402 |
| SPAC27E2.03c | -0.450002255 | 7.89005E-19 | 1.98689E-17 | SPAC27E2.03c |
| SPCC645.08c | -0.44891474 | 7.06173E-16 | 1.46152E-14 | snd1 |
| SPBC1289.07c | -0.447317357 | 4.37606E-10 | 5.21455E-09 | rpc40 |
| SPAC9.10 | -0.447040835 | 1.94022E-05 | 0.000121549 | thi9 |
| SPCC825.01 | -0.445415774 | 1.75572E-12 | 2.71156E-11 | SPCC825.01 |
| SPCC553.03 | -0.442531706 | 1.35421E-08 | 1.3574E-07 | pex1 |
| SPCPB16A4.04c | -0.441072492 | 1.54459E-06 | 1.17499E-05 | trm8 |
| SPAC27D7.11c | -0.440179392 | 4.44801E-09 | 4.75696E-08 | SPAC27D7.11c |
| SPAC9E9.06c | -0.439909585 | 4.31806E-08 | 4.08021E-07 | SPAC9E9.06c |
| SPBC354.08c | -0.439783141 | 0.000206806 | 0.001076573 | rsn1 |
| SPCP31B10.07 | -0.439292016 | 1.10766E-17 | 2.5738E-16 | eft202 |
| SPBC30D10.18c | -0.438031243 | 0.000502195 | 0.002388112 | rpl102 |
| SPAC18G6.07c | -0.436536916 | 5.6855E-12 | 8.21025E-11 | mra1 |
| SPBC3B9.16c | -0.435562631 | 1.23563E-06 | 9.62884E-06 | nup120 |
| SPBC16D10.01c | -0.433038898 | 5.56894E-06 | 3.87325E-05 | SPBC16D10.01c |
| SPBC691.04 | -0.432039002 | 2.97717E-09 | 3.25895E-08 | mss116 |
| SPCP1E11.11 | -0.431554434 | 2.21624E-10 | 2.72341E-09 | puf6 |
| SPAC1002.11 | -0.429998023 | 3.13075E-05 | 0.000188287 | gaa1 |
| SPBC2F12.03c | -0.429286353 | 8.63015E-10 | 1.00267E-08 | ebs1 |
| SPBC1711.06 | -0.428783104 | 5.03659E-20 | 1.37685E-18 | rpl401 |
| SPAC1783.01 | -0.428745816 | 7.19084E-05 | 0.000406183 | SPAC1783.01 |
| SPBC3E7.16c | -0.428537331 | 4.0011E-13 | 6.55564E-12 | leu3 |
| SPAC4G9.08c | -0.427681752 | 1.4907E-06 | 1.14078E-05 | rpc2 |
| SPAC13D6.02c | -0.426980437 | 1.76584E-15 | 3.52615E-14 | byr3 |
| SPCC1393.11 | -0.426296178 | 0.000530782 | 0.002512369 | mrpl20 |
| SPCC18.14c | -0.42545969 | 2.98152E-19 | 7.81616E-18 | rpp0 |
| SPBC1709.05 | -0.425102953 | 6.63647E-19 | 1.67949E-17 | sks2 |

| | | | | |
|---|---|---|---|---|
| SPCC895.05 | -0.424854714 | 1.52905E-05 | 9.86933E-05 | for3 |
| SPCC16C4.08c | -0.424002219 | 6.90485E-05 | 0.000391326 | skb15 |
| SPBC16D10.02 | -0.423851049 | 7.01523E-06 | 4.78158E-05 | trm11 |
| SPAC1527.03 | -0.422660206 | 4.73633E-06 | 3.32128E-05 | SPAC1527.03 |
| SPCC584.01c | -0.421352848 | 2.87142E-05 | 0.000173302 | SPCC584.01c |
| SPBC19F8.05 | -0.421164938 | 0.00132884 | 0.005637369 | SPBC19F8.05 |
| SPAC12B10.10 | -0.420974288 | 0.015938558 | 0.048759969 | nod1 |
| SPCC1902.02 | -0.420453633 | 0.000109799 | 0.000599673 | mug72 |
| SPBC17G9.09 | -0.41955966 | 8.03235E-17 | 1.79307E-15 | tif213 |
| SPBC23G7.08c | -0.419513504 | 1.2946E-07 | 1.143E-06 | rga7 |
| SPBP8B7.09c | -0.418969434 | 1.72162E-06 | 1.29616E-05 | los1 |
| SPAC1250.05 | -0.418869921 | 1.06401E-08 | 1.08136E-07 | rpl3002 |
| SPAC3G9.09c | -0.417299551 | 6.86594E-14 | 1.18979E-12 | tif211 |
| SPCC1827.04 | -0.416342056 | 7.73701E-07 | 6.19931E-06 | vms1 |
| SPBC18H10.03 | -0.416027978 | 6.77388E-13 | 1.08552E-11 | tif35 |
| SPCC1827.06c | -0.415540818 | 1.05783E-15 | 2.15443E-14 | SPCC1827.06c |
| SPAC23H4.17c | -0.413814156 | 0.001022751 | 0.00448891 | srb10 |
| SPBC3H7.14 | -0.412418521 | 4.10442E-06 | 2.91414E-05 | mug176 |
| SPBC365.04c | -0.412335259 | 1.62812E-05 | 0.000104297 | SPBC365.04c |
| SPCC584.11c | -0.412052106 | 1.2505E-09 | 1.43331E-08 | SPCC584.11c |
| SPAC26F1.12c | -0.410651761 | 8.20321E-05 | 0.000459308 | hgh1 |
| SPCC191.03c | -0.410429943 | 0.001425817 | 0.006010949 | SPCC191.03c |
| SPBC1604.08c | -0.410296829 | 2.1847E-12 | 3.31979E-11 | imp1 |
| SPCC584.04 | -0.409397104 | 5.85312E-15 | 1.11646E-13 | sup35 |
| SPCC1827.05c | -0.408570865 | 2.58181E-06 | 1.88816E-05 | SPCC1827.05c |
| SPBC660.14 | -0.40694676 | 0.000168685 | 0.000891748 | mik1 |
| SPBC1105.02c | -0.405058267 | 4.61954E-17 | 1.05425E-15 | lys4 |
| SPBC1711.05 | -0.404517627 | 3.8427E-07 | 3.2256E-06 | SPBC1711.05 |
| SPAC4H3.06 | -0.404352683 | 0.001350791 | 0.005720997 | SPAC4H3.06 |
| SPCC4G3.18 | -0.402960696 | 0.00396917 | 0.0148975 | rix1 |
| SPCC794.06 | -0.40185326 | 1.2964E-06 | 1.00412E-05 | SPCC794.06 |
| SPAC328.10c | -0.40104225 | 5.80556E-17 | 1.31902E-15 | rps502 |
| SPBC16G5.14c | -0.400443849 | 1.6274E-17 | 3.7306E-16 | rps3 |

| | | | | |
|---|---|---|---|---|
| SPAC6B12.15 | -0.399517313 | 2.09896E-18 | 5.1586E-17 | cpc2 |
| SPBC1773.07c | -0.39494426 | 2.11403E-12 | 3.2356E-11 | sbp1 |
| SPCC1827.01c | -0.394866177 | 0.000474294 | 0.002259635 | utp25 |
| SPBC839.13c | -0.393536365 | 6.87563E-16 | 1.42879E-14 | rpl1601 |
| SPCC830.03 | -0.393526497 | 3.72986E-06 | 2.66672E-05 | grc3 |
| SPCC1682.02c | -0.393507016 | 6.24142E-09 | 6.52477E-08 | mcm3 |
| SPAC3A12.04c | -0.393446576 | 0.002943053 | 0.011346067 | rpp1 |
| SPCP31B10.08c | -0.393439861 | 5.26682E-16 | 1.10798E-14 | rpl35a |
| SPAC644.14c | -0.392469805 | 1.72569E-05 | 0.000109715 | rad51 |
| SPAC3C7.08c | -0.391974937 | 4.99265E-13 | 8.10235E-12 | elf1 |
| SPCC4G3.16 | -0.391587854 | 0.014830162 | 0.045985132 | rib2 |
| SPAC25G10.08 | -0.391069379 | 9.57641E-15 | 1.7998E-13 | SPAC25G10.08 |
| SPBC18H10.04c | -0.390906047 | 1.25769E-16 | 2.75936E-15 | sce3 |
| SPAPJ698.02c | -0.390715742 | 5.88879E-17 | 1.33201E-15 | rps002 |
| SPBP4H10.05c | -0.390334495 | 2.63689E-11 | 3.56608E-10 | spe2 |
| SPBC691.02c | -0.389908249 | 9.12186E-05 | 0.000505211 | drp1 |
| SPBC17D11.05 | -0.388968273 | 5.65443E-16 | 1.18465E-14 | tif32 |
| SPBC4F6.13c | -0.388062394 | 3.93338E-07 | 3.28553E-06 | erb1 |
| SPCC1223.05c | -0.384845494 | 1.62498E-11 | 2.25119E-10 | rpl3702 |
| SPBC211.02c | -0.383560775 | 0.007938118 | 0.02725296 | cwf3 |
| SPAC6G9.02c | -0.383124281 | 1.76591E-05 | 0.000111863 | nop9 |
| SPCC1682.14 | -0.382003574 | 3.78206E-16 | 8.0895E-15 | rpl1902 |
| SPAC1B1.03c | -0.381865357 | 6.13528E-11 | 7.92008E-10 | kap95 |
| SPAC3G9.14 | -0.381636848 | 2.54403E-07 | 2.1784E-06 | sak1 |
| SPAC9G1.12 | -0.380821157 | 4.56101E-07 | 3.7667E-06 | cpd1 |
| SPBC839.11c | -0.380656576 | 0.000102873 | 0.000564863 | hut1 |
| SPCC18.07 | -0.379378187 | 4.28378E-06 | 3.02468E-05 | rpc53 |
| SPAC110.04c | -0.378190229 | 1.65139E-16 | 3.57707E-15 | pss1 |
| SPBC1734.07c | -0.378167104 | 0.000581019 | 0.002712482 | trs8502 |
| SPBC460.03 | -0.377794637 | 2.1544E-05 | 0.000133981 | vba2 |
| SPAC4D7.08c | -0.377741031 | 2.12189E-12 | 3.23793E-11 | ade4 |
| SPBC685.06 | -0.37725126 | 8.53955E-15 | 1.61682E-13 | rps001 |
| SPCC895.03c | -0.377131493 | 0.000231741 | 0.001196625 | sua5 |

| | | | | |
|---|---|---|---|---|
| SPAC1F7.13c | -0.376628668 | 4.62166E-16 | 9.84413E-15 | rpl801 |
| SPBP8B7.21 | -0.376367111 | 1.00136E-09 | 1.15552E-08 | ubp3 |
| SPAC16C9.06c | -0.37628387 | 4.57059E-08 | 4.29501E-07 | upf1 |
| SPBC2D10.12 | -0.376258566 | 4.25003E-10 | 5.0762E-09 | rhp23 |
| SPAC637.07 | -0.376121574 | 1.89305E-14 | 3.44387E-13 | moe1 |
| SPCC1919.09 | -0.376006707 | 1.43562E-11 | 2.01618E-10 | tif6 |
| SPBC14C8.12 | -0.375844244 | 2.41541E-05 | 0.000149125 | rpb8 |
| SPAC630.10 | -0.375660291 | 0.00016645 | 0.000880842 | bmt2 |
| SPBC1347.02 | -0.375560037 | 3.85145E-09 | 4.13626E-08 | fkbp39 |
| SPBC29A3.16 | -0.375558741 | 0.000648709 | 0.002995664 | rrs1 |
| SPCC584.13 | -0.374513028 | 0.000725226 | 0.003316058 | SPCC584.13 |
| SPBC1289.03c | -0.374314415 | 9.07166E-16 | 1.86242E-14 | spi1 |
| SPAC2G11.02 | -0.373581412 | 6.03409E-05 | 0.000345423 | urb2 |
| SPBC19C7.06 | -0.373369884 | 7.22759E-13 | 1.15461E-11 | prs1 |
| SPCC1795.11 | -0.372505996 | 3.9938E-14 | 7.089E-13 | sum3 |
| SPBC336.04 | -0.371826181 | 0.002870181 | 0.011107013 | cdc6 |
| SPAC29A4.02c | -0.371015898 | 8.37648E-17 | 1.86176E-15 | SPAC29A4.02c |
| SPBC14F5.06 | -0.368884975 | 1.78509E-11 | 2.45306E-10 | rli1 |
| SPNCRNA.1599 | -0.368419971 | 0.011117563 | 0.036061536 | ptk1-antisense-1 |
| SPAC17A5.03 | -0.368405609 | 1.98567E-14 | 3.59956E-13 | rpl301 |
| SPBC17D1.05 | -0.36791158 | 1.05981E-08 | 1.08082E-07 | SPBC17D1.05 |
| SPAC24H6.10c | -0.367505848 | 3.4192E-12 | 5.11081E-11 | SPAC24H6.10c |
| SPCC970.05 | -0.367272495 | 8.35367E-14 | 1.4427E-12 | rpl3601 |
| SPAC10F6.03c | -0.366388987 | 1.54731E-08 | 1.53888E-07 | cts1 |
| SPAC890.04c | -0.366366322 | 7.18657E-07 | 5.78547E-06 | SPAC890.04c |
| SPAC167.01 | -0.366167863 | 0.010501734 | 0.034303426 | ire1 |
| SPBC13A2.01c | -0.365463717 | 0.001629278 | 0.006776947 | cbc2 |
| SPNCRNA.1096 | -0.364595837 | 0.000506423 | 0.002405978 | SPCC1884.01-antisense-1 |
| SPBC215.12 | -0.364552813 | 0.001280544 | 0.005450575 | cwf10 |
| SPCC1322.15 | -0.364059342 | 4.98192E-08 | 4.6389E-07 | rpl3402 |
| SPBC17G9.03c | -0.363696644 | 5.2784E-12 | 7.66568E-11 | krs1 |
| SPCC338.17c | -0.363640698 | 5.2331E-06 | 3.65459E-05 | rad21 |
| SPBC13E7.03c | -0.363356812 | 1.00324E-05 | 6.64909E-05 | SPBC13E7.03c |

| | | | | |
|---|---|---|---|---|
| SPBC3D6.15 | -0.362790912 | 2.58927E-10 | 3.15152E-09 | rps2501 |
| SPAC821.05 | -0.362474657 | 2.25996E-12 | 3.39792E-11 | tif38 |
| SPBC25H2.07 | -0.362362763 | 3.44906E-11 | 4.60355E-10 | tif11 |
| SPCC1259.03 | -0.362009698 | 0.000121705 | 0.000659761 | rpa12 |
| SPAC17G6.15c | -0.362007346 | 1.63447E-08 | 1.61926E-07 | SPAC17G6.15c |
| SPAC323.01c | -0.361708907 | 0.000447342 | 0.002142981 | pos5 |
| SPBC1711.04 | -0.36111322 | 3.45107E-07 | 2.9112E-06 | mtd1 |
| SPBC27B12.11c | -0.360860687 | 1.88734E-05 | 0.000118964 | pho7 |
| SPCC1235.11 | -0.359590461 | 3.77408E-09 | 4.06171E-08 | mpc1 |
| SPAC1D4.10 | -0.35868174 | 1.55836E-05 | 0.000100331 | trz1 |
| SPBC146.03c | -0.358650057 | 3.8697E-05 | 0.000228957 | cut3 |
| SPAC26A3.07c | -0.358616319 | 5.64029E-12 | 8.16803E-11 | rpl1101 |
| SPAC22A12.04c | -0.358485142 | 1.16416E-14 | 2.16406E-13 | rps2201 |
| SPCC132.01c | -0.358441936 | 6.65489E-07 | 5.3659E-06 | mtr1 |
| SPAC56F8.09 | -0.358276951 | 0.000559573 | 0.002624347 | rrp8 |
| SPCC1183.08c | -0.357876119 | 5.77612E-14 | 1.01469E-12 | rpl101 |
| SPCC132.02 | -0.355775843 | 0.000257343 | 0.001315537 | hst2 |
| SPAC13G7.08c | -0.355365633 | 9.86475E-06 | 6.55768E-05 | crb3 |
| SPAC12G12.09 | -0.355086118 | 0.001243602 | 0.005302163 | eti1 |
| SPBC14C8.03 | -0.354888638 | 1.10446E-08 | 1.11802E-07 | fma2 |
| SPBC12C2.09c | -0.354672598 | 0.000103286 | 0.000566523 | izh2 |
| SPCC1494.06c | -0.354657057 | 0.000116883 | 0.000636084 | dbp9 |
| SPAC664.04c | -0.354489861 | 4.36121E-13 | 7.10017E-12 | rps1602 |
| SPCC1494.07 | -0.354426662 | 7.30264E-07 | 5.86967E-06 | SPCC1494.07 |
| SPAC644.15 | -0.353345018 | 2.60977E-13 | 4.30359E-12 | rpp101 |
| SPAC1142.04 | -0.352682684 | 5.98664E-05 | 0.000343091 | SPAC1142.04 |
| SPCC5E4.07 | -0.352386099 | 9.02194E-15 | 1.70185E-13 | rpl2802 |
| SPBP23A10.07 | -0.3522546 | 6.82946E-08 | 6.2348E-07 | rpa2 |
| SPBC6B1.08c | -0.352206446 | 6.44268E-09 | 6.72143E-08 | ofd1 |
| SPBC1306.02 | -0.351688826 | 0.013087192 | 0.041450883 | rtt10 |
| SPBC23E6.05 | -0.350830892 | 4.38143E-07 | 3.64193E-06 | arx1 |
| SPBP35G2.08c | -0.350813802 | 0.000452335 | 0.002163083 | air1 |
| SPBC3B9.19 | -0.350761823 | 1.23631E-07 | 1.09722E-06 | mge1 |

| | | | | |
|---|---|---|---|---|
| SPCC962.04 | -0.350692982 | 9.22112E-13 | 1.45489E-11 | rps1201 |
| SPAC1071.07c | -0.350509625 | 7.15153E-15 | 1.35906E-13 | rps1502 |
| SPCC553.09c | -0.350158681 | 1.86769E-05 | 0.000117872 | spb70 |
| SPCC576.08c | -0.349645252 | 2.50173E-14 | 4.48732E-13 | rps2 |
| SPAC13G7.12c | -0.349292201 | 2.05956E-05 | 0.000128396 | eki1 |
| SPAC17G6.09 | -0.348952841 | 3.19132E-06 | 2.31404E-05 | sec62 |
| SPCC1223.07c | -0.348897321 | 3.68128E-10 | 4.42794E-09 | drs1 |
| SPAC1834.05 | -0.348410998 | 4.71745E-08 | 4.4087E-07 | alg9 |
| SPBPB10D8.07c | -0.348333523 | 0.014413787 | 0.044956242 | SPBPB10D8.07c |
| SPBC365.03c | -0.348091158 | 2.12314E-13 | 3.53534E-12 | rpl2101 |
| SPBC649.02 | -0.345338794 | 1.90046E-11 | 2.6046E-10 | rps1902 |
| SPCC4B3.07 | -0.345249372 | 4.18321E-10 | 5.01986E-09 | nro1 |
| SPAC343.14c | -0.345203025 | 1.63365E-06 | 1.2357E-05 | SPAC343.14c |
| SPAC26F1.13c | -0.34428955 | 4.64421E-08 | 4.35619E-07 | lrs1 |
| SPCC18.13 | -0.344061685 | 0.000244627 | 0.001255557 | SPCC18.13 |
| SPAC6F6.17 | -0.343508785 | 0.000374533 | 0.001832164 | rif1 |
| SPBC16H5.10c | -0.34207071 | 3.98567E-06 | 2.83376E-05 | prp43 |
| SPBC354.09c | -0.341879485 | 4.43076E-06 | 3.12414E-05 | SPBC354.09c |
| SPAP8A3.03 | -0.341769657 | 1.73268E-05 | 0.000109894 | zrt2 |
| SPCC330.07c | -0.341636331 | 0.000256837 | 0.001314264 | SPCC330.07c |
| SPCC1235.02 | -0.34103357 | 4.14584E-07 | 3.45734E-06 | bio2 |
| SPAC18G6.14c | -0.34082539 | 1.60553E-12 | 2.48711E-11 | rps7 |
| SPAC30C2.04 | -0.340116244 | 1.25432E-10 | 1.57159E-09 | asc1 |
| SPAC17G8.06c | -0.339531137 | 2.69447E-12 | 4.03933E-11 | SPAC17G8.06c |
| SPAC23H3.09c | -0.339352444 | 4.92173E-12 | 7.18847E-11 | gly1 |
| SPAC8F11.09c | -0.33899557 | 0.000129495 | 0.000699768 | nnt1 |
| SPCC297.05 | -0.338957882 | 1.04482E-06 | 8.25522E-06 | SPCC297.05 |
| SPAC13G6.07c | -0.338860077 | 1.15112E-13 | 1.97468E-12 | rps601 |
| SPAPB2B4.04c | -0.338415945 | 4.97943E-05 | 0.00029058 | pmc1 |
| SPAC3A12.10 | -0.338059127 | 6.36248E-10 | 7.52893E-09 | rpl2001 |
| SPBC365.01 | -0.337201542 | 1.78812E-07 | 1.55457E-06 | crs103 |
| SPBC2F12.04 | -0.33707877 | 5.63005E-13 | 9.10785E-12 | rpl1701 |
| SPAC1952.06c | -0.336224412 | 5.58623E-05 | 0.000322676 | SPAC1952.06c |

| | | | | |
|---|---|---|---|---|
| SPAC4A8.16c | -0.336157358 | 1.9779E-11 | 2.69627E-10 | tif33 |
| SPAC56F8.02 | -0.33615536 | 0.005017195 | 0.018372422 | SPAC56F8.02 |
| SPBP19A11.06 | -0.335908786 | 0.005640064 | 0.020361588 | lid2 |
| SPAC31G5.02 | -0.335472042 | 0.008408534 | 0.028599085 | SPAC31G5.02 |
| SPAC18G6.11c | -0.334909738 | 2.39655E-05 | 0.00014814 | rrn3 |
| SPBC16D10.11c | -0.334882261 | 3.38706E-11 | 4.53263E-10 | rps1801 |
| SPAC19A8.01c | -0.334829337 | 0.005545238 | 0.020090188 | sec73 |
| SPBP8B7.30c | -0.334318313 | 0.000169133 | 0.000893192 | thi5 |
| SPCC63.06 | -0.334079564 | 7.41596E-07 | 5.9514E-06 | SPCC63.06 |
| SPCC576.11 | -0.333831623 | 4.93575E-12 | 7.18847E-11 | rpl15 |
| SPAC139.01c | -0.333585741 | 3.1027E-08 | 2.99264E-07 | ath2 |
| SPBC27B12.13 | -0.333223699 | 4.63636E-07 | 3.8166E-06 | tom40 |
| SPAC144.12 | -0.33210817 | 5.90224E-08 | 5.43644E-07 | rki1 |
| SPBC839.08c | -0.332045146 | 6.24164E-07 | 5.06464E-06 | its8 |
| SPBC32C12.03c | -0.331428466 | 0.000749461 | 0.0034177 | ppk25 |
| SPAC5D6.01 | -0.33119705 | 1.44909E-09 | 1.65352E-08 | rps2202 |
| SPAC19B12.03 | -0.329087635 | 2.18145E-07 | 1.88053E-06 | bgs3 |
| SPBC1289.13c | -0.328840915 | 9.91455E-05 | 0.000546744 | gmh6 |
| SPCP25A2.02c | -0.328700084 | 1.08149E-05 | 7.14289E-05 | rhp26 |
| SPAC22F8.10c | -0.328612865 | 0.000214805 | 0.001115937 | sap145 |
| SPBC1604.02c | -0.328550148 | 5.41734E-05 | 0.000313274 | ppr1 |
| SPBC18H10.13 | -0.328064187 | 7.82094E-13 | 1.24163E-11 | rps1402 |
| SPCC162.03 | -0.327237083 | 0.000175678 | 0.000922037 | SPCC162.03 |
| SPAC31G5.11 | -0.327231118 | 3.03905E-08 | 2.94235E-07 | pac2 |
| SPCC1672.01 | -0.327154872 | 0.000157307 | 0.000835918 | SPCC1672.01 |
| SPCC1682.13 | -0.326932248 | 0.001632701 | 0.006785665 | laf2 |
| SPAPB1A10.11c | -0.326563394 | 0.000620336 | 0.002875028 | mse1 |
| SPBC16A3.08c | -0.326304515 | 1.00881E-12 | 1.58191E-11 | oga1 |
| SPAC27E2.05 | -0.325626005 | 0.000156898 | 0.000834614 | cdc1 |
| SPBC19F8.08 | -0.325423694 | 4.49042E-12 | 6.59627E-11 | rps401 |
| SPAC15A10.04c | -0.325401077 | 1.652E-05 | 0.000105431 | zpr1 |
| SPAC890.07c | -0.324819268 | 1.76355E-07 | 1.53582E-06 | rmt1 |
| SPBC106.18 | -0.324631879 | 2.18867E-12 | 3.31979E-11 | rpl2501 |

| | | | | |
|---|---|---|---|---|
| SPAC4F8.03 | -0.324454268 | 1.60878E-05 | 0.000103348 | sdo1 |
| SPAC17D4.04 | -0.32413331 | 1.05853E-05 | 7.00027E-05 | trm401 |
| SPAC1783.08c | -0.323944035 | 1.25182E-12 | 1.95698E-11 | rpl1502 |
| SPCC553.11c | -0.323899226 | 2.48842E-05 | 0.000152854 | toa2 |
| SPAC56F8.06c | -0.323148256 | 0.003011402 | 0.011583362 | alg10 |
| SPAC1834.08 | -0.322582944 | 0.000173256 | 0.000913078 | mak1 |
| SPBC30B4.07c | -0.322281819 | 0.002858355 | 0.011078023 | tfb4 |
| SPCC757.08 | -0.3221634 | 0.002839204 | 0.011028883 | rrp45 |
| SPAC694.02 | -0.321572148 | 4.51953E-07 | 3.73848E-06 | SPAC694.02 |
| SPBC660.15 | -0.321111732 | 3.64198E-06 | 2.61486E-05 | msi2 |
| SPCC1450.04 | -0.321034424 | 1.98782E-12 | 3.06077E-11 | tef5 |
| SPBC8D2.06 | -0.32059534 | 5.86096E-08 | 5.40816E-07 | irs1 |
| SPAC11G7.04 | -0.320594321 | 3.20354E-11 | 4.2983E-10 | ubi1 |
| SPBC3H7.07c | -0.320351066 | 8.97092E-05 | 0.00049739 | ser2 |
| SPAC3H1.02c | -0.319623943 | 6.45286E-05 | 0.00036693 | SPAC3H1.02c |
| SPBC30B4.01c | -0.318988012 | 0.000876868 | 0.003925175 | wsc1 |
| SPAC1A6.05c | -0.318649226 | 3.37175E-05 | 0.000201595 | ptl3 |
| SPBC18H10.12c | -0.318570477 | 8.49327E-12 | 1.21618E-10 | rpl701 |
| SPAC12G12.06c | -0.318473838 | 2.45185E-07 | 2.10653E-06 | rcl1 |
| SPBC16H5.11c | -0.318031473 | 4.17783E-06 | 2.95804E-05 | skb1 |
| SPAC23C4.18c | -0.317968722 | 0.00635259 | 0.022520417 | rad4 |
| SPAC3F10.03 | -0.317863422 | 2.44696E-10 | 2.98541E-09 | grs1 |
| SPBC713.10 | -0.317634569 | 0.000511867 | 0.002425084 | tim16 |
| SPAC13G6.02c | -0.317492357 | 7.44372E-13 | 1.18543E-11 | rps101 |
| SPAC4D7.05 | -0.317360563 | 5.12981E-09 | 5.46324E-08 | sum1 |
| SPNCRNA.1616 | -0.317336864 | 0.002825608 | 0.010992776 | SPNCRNA.1616 |
| SPBC4C3.07 | -0.317029651 | 5.41961E-10 | 6.42809E-09 | eif6 |
| SPCC338.07c | -0.3164893 | 7.29204E-08 | 6.61713E-07 | naa15 |
| SPBC20F10.02c | -0.316452015 | 0.007785438 | 0.026868856 | SPBC20F10.02c |
| SPAC521.05 | -0.316155668 | 1.60448E-11 | 2.22883E-10 | rps802 |
| SPBC839.05c | -0.315860034 | 3.95456E-11 | 5.20697E-10 | rps1701 |
| SPAC1565.05 | -0.315421076 | 0.000411092 | 0.001982549 | utp8 |
| SPAC23H4.10c | -0.314746243 | 4.24034E-06 | 2.99815E-05 | thi4 |

| | | | | |
|---|---|---|---|---|
| SPAC24C9.09 | -0.313843383 | 0.001125188 | 0.00487868 | SPAC24C9.09 |
| SPBC146.01 | -0.313738567 | 6.59288E-05 | 0.000374476 | med15 |
| SPAC959.08 | -0.313539515 | 6.60583E-11 | 8.50605E-10 | rpl2102 |
| SPBC337.05c | -0.313344985 | 2.2958E-09 | 2.5348E-08 | cct8 |
| SPBC3B8.06 | -0.313279555 | 3.83777E-05 | 0.000227331 | SPBC3B8.06 |
| SPCC1322.11 | -0.313180505 | 1.34492E-10 | 1.67689E-09 | rpl2302 |
| SPBC776.01 | -0.311756691 | 1.85389E-09 | 2.08746E-08 | rpl29 |
| SPBP35G2.07 | -0.310893523 | 1.6341E-08 | 1.61926E-07 | ilv1 |
| SPBPB2B2.09c | -0.310607043 | 0.000281567 | 0.001419496 | pan5 |
| SPBC16G5.10 | -0.310376531 | 0.009486573 | 0.031593068 | rrp42 |
| SPCC417.08 | -0.310339995 | 4.61554E-12 | 6.76065E-11 | tef3 |
| SPAPB1E7.12 | -0.31009994 | 5.98216E-11 | 7.74197E-10 | rps602 |
| SPAC6F12.10c | -0.3098635 | 4.47729E-08 | 4.22286E-07 | ade3 |
| SPCC364.03 | -0.309741086 | 1.56675E-11 | 2.18831E-10 | rpl1702 |
| SPCC16C4.11 | -0.309229385 | 0.005764573 | 0.020781731 | pef1 |
| SPAC23C11.13c | -0.308708149 | 1.48617E-06 | 1.13903E-05 | hpt1 |
| SPBC17G9.10 | -0.308106811 | 1.11331E-09 | 1.28181E-08 | rpl1102 |
| SPAC8C9.06c | -0.308088894 | 0.000402238 | 0.00194168 | ppr4 |
| SPAC1952.01 | -0.307287173 | 0.009720959 | 0.032081049 | gab1 |
| SPAC3G6.04 | -0.307057166 | 0.004839295 | 0.017784669 | rnp24 |
| SPAC15F9.02 | -0.306721245 | 0.000586395 | 0.002735083 | seh1 |
| SPBC16H5.02 | -0.306637206 | 3.53886E-08 | 3.40049E-07 | pfk1 |
| SPAC1B2.03c | -0.306149717 | 6.11015E-09 | 6.40064E-08 | SPAC1B2.03c |
| SPBC25H2.13c | -0.306074141 | 0.007167748 | 0.025045472 | cdc20 |
| SPAC1071.03c | -0.305612909 | 0.000342864 | 0.001696727 | sil1 |
| SPBC106.19 | -0.305439816 | 0.000355137 | 0.001747314 | SPBC106.19 |
| SPAC959.07 | -0.305209826 | 8.80711E-11 | 1.12274E-09 | rps403 |
| SPCC895.07 | -0.304667212 | 8.00071E-05 | 0.000448953 | alp14 |
| SPBC4F6.14 | -0.304280041 | 0.000116166 | 0.000633093 | nop4 |
| SPCC306.03c | -0.304215382 | 0.000813546 | 0.003680397 | cnd2 |
| SPAC31G5.17c | -0.303339221 | 4.46748E-11 | 5.85584E-10 | rps1001 |
| SPBC646.09c | -0.303243539 | 3.6876E-09 | 3.99386E-08 | int6 |
| SPBC405.01 | -0.302801372 | 3.7277E-09 | 4.02562E-08 | ade1 |

| | | | | |
|---|---|---|---|---|
| SPAC23H4.15 | -0.30243044 | 0.000554903 | 0.002609628 | tsr1 |
| SPAPB8E5.06c | -0.302336709 | 1.22177E-10 | 1.53456E-09 | rpl302 |
| SPBC947.12 | -0.301259085 | 0.016055401 | 0.049029398 | kms2 |
| SPCC16A11.13 | -0.300935537 | 0.000952443 | 0.004226467 | luc7 |
| SPAC6G9.10c | -0.299682414 | 0.000904708 | 0.004039184 | sen1 |
| SPBC776.11 | -0.299665957 | 1.52017E-09 | 1.72691E-08 | rpl2801 |
| SPBC428.04 | -0.299621116 | 0.00294948 | 0.011362276 | apq12 |
| SPBC2G5.07c | -0.299364576 | 0.000312722 | 0.001558122 | rpc25 |
| SPBC685.07c | -0.299280972 | 2.56046E-09 | 2.81486E-08 | rpl2701 |
| SPBC16A3.03c | -0.298605166 | 7.74598E-05 | 0.000435137 | ppr7 |
| SPBC25D12.05 | -0.298544935 | 0.000354022 | 0.001743791 | trm1 |
| SPAC688.10 | -0.29820069 | 0.01221922 | 0.039236589 | rev3 |
| SPCC63.05 | -0.297211619 | 0.012498748 | 0.039958473 | tap42 |
| SPBC18H10.16 | -0.29705423 | 0.000364298 | 0.001787229 | can1 |
| SPAC2C4.12c | -0.296850656 | 0.004820134 | 0.017726996 | tpt1 |
| SPAP7G5.02c | -0.296240393 | 4.67576E-08 | 4.37775E-07 | gua2 |
| SPAC21E11.06 | -0.296180946 | 3.15824E-06 | 2.29332E-05 | tif224 |
| SPAC6G9.03c | -0.296065967 | 0.007990561 | 0.027414594 | mug183 |
| SPCC757.09c | -0.295315745 | 5.58852E-07 | 4.55638E-06 | rnc1 |
| SPAC23A1.11 | -0.295277843 | 1.02828E-10 | 1.30113E-09 | rpl1602 |
| SPAC27F1.06c | -0.295193485 | 0.000844592 | 0.003804013 | SPAC27F1.06c |
| SPCC1322.01 | -0.295107924 | 0.005907656 | 0.021252595 | rpm1 |
| SPBC18E5.06 | -0.295096877 | 1.39072E-09 | 1.59046E-08 | rps21 |
| SPBP8B7.11 | -0.294985955 | 1.18227E-07 | 1.05109E-06 | nxt3 |
| SPAC20G4.07c | -0.294845272 | 5.94726E-06 | 4.10288E-05 | erg4 |
| SPCC622.14 | -0.294691747 | 1.91752E-05 | 0.000120275 | SPCC622.14 |
| SPAC22G7.06c | -0.294006129 | 8.67396E-08 | 7.77917E-07 | ura1 |
| SPAC5D6.12 | -0.293851654 | 8.08591E-05 | 0.000453237 | mtf2 |
| SPBC1A4.03c | -0.293693435 | 5.02321E-05 | 0.000292801 | top2 |
| SPAC22H12.04c | -0.293556523 | 2.46054E-11 | 3.33641E-10 | rps102 |
| SPBC839.15c | -0.293288136 | 1.47581E-11 | 2.06695E-10 | tef103 |
| SPAC3H5.10 | -0.293054246 | 2.01521E-09 | 2.25916E-08 | rpl3202 |
| SPCC330.05c | -0.293009351 | 5.18756E-06 | 3.62775E-05 | ura4 |

| | | | | |
|---|---|---|---|---|
| SPAC13D6.03c | -0.292225431 | 0.00974271 | 0.032132086 | trm9 |
| SPAC3H1.05 | -0.292168363 | 2.32768E-06 | 1.71952E-05 | SPAC3H1.05 |
| SPBC25H2.05 | -0.292116472 | 7.56658E-09 | 7.83003E-08 | egd2 |
| SPCC18.17c | -0.291677781 | 0.000594752 | 0.002767026 | SPCC18.17c |
| SPCC18B5.04 | -0.290349233 | 0.00037109 | 0.001817061 | rsm18 |
| SPAC3H5.09c | -0.289097583 | 0.002142696 | 0.008625844 | SPAC3H5.09c |
| SPAC6G10.02c | -0.288564538 | 0.000132587 | 0.000714208 | tea3 |
| SPBC28F2.02 | -0.288532208 | 7.4752E-06 | 5.07479E-05 | mep33 |
| SPBC119.11c | -0.288272941 | 0.005572979 | 0.020162115 | pac1 |
| SPAC8F11.04 | -0.288156088 | 0.002097366 | 0.00847568 | SPAC8F11.04 |
| SPCC613.01 | -0.288035194 | 0.000188443 | 0.000984989 | SPCC613.01 |
| SPAC637.12c | -0.28791195 | 0.002675791 | 0.010513944 | mst1 |
| SPCC1795.12c | -0.287243408 | 0.015660532 | 0.048110961 | SPCC1795.12c |
| SPBC13E7.05 | -0.287030215 | 0.003545705 | 0.013446325 | gpi14 |
| SPAC1751.03 | -0.287001815 | 8.4778E-09 | 8.70251E-08 | tif313 |
| SPBC14F5.03c | -0.286855607 | 6.22295E-06 | 4.27003E-05 | kap123 |
| SPAC23D3.03c | -0.286743553 | 2.52421E-05 | 0.000154237 | SPAC23D3.03c |
| SPAC18G6.06 | -0.286656944 | 0.004088514 | 0.015289308 | utp11 |
| SPCC31H12.05c | -0.28629869 | 9.75785E-06 | 6.50354E-05 | sds21 |
| SPAC4F8.14c | -0.286195594 | 1.55302E-06 | 1.17965E-05 | hcs1 |
| SPBC1604.06c | -0.285540544 | 0.005786788 | 0.020847118 | noc4 |
| SPBC119.13c | -0.285005021 | 0.00066289 | 0.00305563 | prp31 |
| SPAC1A6.10 | -0.285003103 | 6.15783E-05 | 0.000351719 | tcd1 |
| SPBC17G9.07 | -0.284632493 | 4.20963E-10 | 5.03973E-09 | rps2402 |
| SPBP8B7.29 | -0.283996357 | 0.015748359 | 0.048293707 | abz1 |
| SPAC13G6.09 | -0.283550393 | 0.002490016 | 0.009882735 | SPAC13G6.09 |
| SPAC4D7.04c | -0.283021316 | 0.000299173 | 0.001496448 | rer2 |
| SPBC13G1.10c | -0.282829268 | 0.001146342 | 0.004945232 | mug81 |
| SPAC1A6.09c | -0.282784617 | 1.49156E-05 | 9.67624E-05 | lag1 |
| SPAC22H10.09 | -0.282753828 | 0.001171688 | 0.005024892 | SPAC22H10.09 |
| SPAC2C4.16c | -0.282322194 | 6.66384E-10 | 7.86733E-09 | rps801 |
| SPAP8A3.14c | -0.281819694 | 0.004736679 | 0.017445174 | sls1 |
| SPAC23C4.15 | -0.281724659 | 7.17021E-05 | 0.000405466 | rpb5 |

| | | | | |
|---|---|---|---|---|
| SPBP4H10.13 | -0.281596675 | 1.39832E-08 | 1.39886E-07 | rps2302 |
| SPAC6G10.07 | -0.280899297 | 6.38455E-06 | 4.37505E-05 | cbc1 |
| SPBC646.02 | -0.280807142 | 0.001182092 | 0.005060884 | cwf11 |
| SPBC651.10 | -0.280324457 | 0.01450449 | 0.045184005 | nse5 |
| SPAC589.10c | -0.279888128 | 5.87423E-09 | 6.16613E-08 | ubi5 |
| SPAC19A8.08 | -0.279616076 | 0.000378535 | 0.001848205 | upf2 |
| SPCC330.14c | -0.279232858 | 7.36509E-09 | 7.63699E-08 | rpl2402 |
| SPBC19G7.08c | -0.278713688 | 0.008036341 | 0.027516258 | art1 |
| SPAC2F3.12c | -0.27782002 | 0.010079803 | 0.033136947 | plp1 |
| SPAC3H1.10 | -0.277799762 | 0.002290033 | 0.009152968 | pcs2 |
| SPBC405.07 | -0.277641155 | 7.80646E-08 | 7.03821E-07 | rpl3602 |
| SPAP27G11.04c | -0.277549939 | 0.000930399 | 0.004139426 | tad3 |
| SPAC1B3.15c | -0.276657598 | 0.009170696 | 0.030721232 | SPAC1B3.15c |
| SPBC36.03c | -0.276475475 | 5.98147E-05 | 0.000343091 | mfs3 |
| SPBC800.04c | -0.276335715 | 9.96748E-10 | 1.1528E-08 | rpl4301 |
| SPAC1639.01c | -0.276319661 | 2.30491E-06 | 1.70516E-05 | SPAC1639.01c |
| SPAC328.02 | -0.275657666 | 0.000896847 | 0.004007591 | dbl4 |
| SPBC16H5.14c | -0.27561289 | 0.008957541 | 0.030145073 | SPBC16H5.14c |
| SPCC550.11 | -0.275430179 | 1.26057E-06 | 9.79335E-06 | SPCC550.11 |
| SPAC31A2.05c | -0.275259031 | 0.010546753 | 0.034428481 | mis4 |
| SPAC25G10.05c | -0.274478767 | 4.08783E-08 | 3.877E-07 | his1 |
| SPAC31G5.13 | -0.274467581 | 1.48652E-05 | 9.66807E-05 | rpn11 |
| SPBC11G11.03 | -0.274390767 | 1.90088E-05 | 0.000119377 | mrt4 |
| SPBP22H7.08 | -0.274138636 | 1.29654E-08 | 1.30215E-07 | rps1002 |
| SPCC1494.03 | -0.273848952 | 0.016093713 | 0.049087745 | arz1 |
| SPBC18H10.02 | -0.273218725 | 1.4908E-05 | 9.67624E-05 | lcf1 |
| SPBC12C2.06 | -0.273057363 | 6.56E-06 | 4.48926E-05 | dbp5 |
| SPCC1919.12c | -0.27288333 | 0.010281416 | 0.033691409 | SPCC1919.12c |
| SPAC3H5.12c | -0.272158494 | 5.07579E-09 | 5.417E-08 | rpl501 |
| SPBC17A3.01c | -0.271267341 | 2.78715E-05 | 0.000168814 | tim50 |
| SPBC25D12.06 | -0.271237093 | 0.001183051 | 0.005060884 | SPBC25D12.06 |
| SPAC607.03c | -0.271235775 | 7.09726E-08 | 6.45573E-07 | snu13 |
| SPAC29E6.06c | -0.271037292 | 2.45542E-06 | 1.80431E-05 | SPAC29E6.06c |

| | | | | |
|---|---|---|---|---|
| SPBP8B7.31 | -0.270707614 | 0.013851525 | 0.043494469 | SPBP8B7.31 |
| SPBC1E8.02 | -0.270546919 | 0.000628261 | 0.002903861 | SPBC1E8.02 |
| SPCC18B5.06 | -0.270054789 | 0.001288368 | 0.005474762 | dom34 |
| SPAC1D4.14 | -0.269862167 | 0.007199331 | 0.025136325 | tho2 |
| SPAC1B3.05 | -0.269728179 | 1.89109E-05 | 0.000118964 | not3 |
| SPAC1B9.03c | -0.269076409 | 0.013556731 | 0.042647391 | SPAC1B9.03c |
| SPBC29A10.01 | -0.269066759 | 1.71674E-06 | 1.29439E-05 | ccr1 |
| SPAC25H1.08c | -0.268830063 | 0.000435402 | 0.002089929 | SPAC25H1.08c |
| SPAC3F10.18c | -0.268673554 | 0.00217016 | 0.008714736 | rpl4102 |
| SPAC9.03c | -0.268576807 | 0.001796056 | 0.007410361 | brr2 |
| SPAC1071.05 | -0.268527364 | 0.000453516 | 0.002166705 | SPAC1071.05 |
| SPAC1805.14 | -0.268252191 | 0.003461309 | 0.013165337 | SPAC1805.14 |
| SPCC1259.11c | -0.267811402 | 0.000148111 | 0.000792821 | gyp2 |
| SPCC613.05c | -0.267778373 | 1.0692E-08 | 1.08448E-07 | rpl35 |
| SPAC22A12.03c | -0.267374699 | 0.007558245 | 0.026248471 | csn4 |
| SPAC139.06 | -0.266035173 | 0.003957727 | 0.014865468 | hat1 |
| SPBP4H10.06c | -0.265992192 | 0.00080202 | 0.003634686 | cut14 |
| SPBC543.02c | -0.265149696 | 7.88161E-06 | 5.33653E-05 | SPBC543.02c |
| SPAC4D7.09 | -0.265023405 | 3.68769E-05 | 0.000219203 | tif223 |
| SPAC23H4.14 | -0.265013829 | 0.004857326 | 0.017838111 | vam6 |
| SPAC17H9.04c | -0.264755733 | 8.87071E-05 | 0.000492368 | nrp1 |
| SPAC15E1.08 | -0.264114085 | 0.000920656 | 0.004099644 | naa10 |
| SPBP8B7.06 | -0.263121052 | 2.12157E-08 | 2.06975E-07 | rpp201 |
