## Supplemental Table 6 for "Rapalink-1 reveals novel mTOR-dependent genes and an agmatinergic axis-based metabolic feedback regulating mTOR activity and lifespan"

Supplementary Table 6

| Gene name | Agmatine fitness ratio | Putrescine fitness ratio |
|---|---|---|
| SPBC800.04c | 0.931046459 | 0.069510511 |
| SPCC285.17 | 0.459010401 | 0.091671921 |
| SPBC651.03c | 0.997306482 | 0.097250859 |
| SPAC3F10.17 | 0.206422018 | 0.121698113 |
| SPAC29B12.05c | 2.898686756 | 0.127053159 |
| SPAC57A7.12 | 0.307852784 | 0.158125117 |
| SPAC2C4.16c | 0.223124043 | 0.200277566 |
| SPCC1393.03 | 0.335946218 | 0.210514275 |
| SPCP1E11.09c | 1.002891759 | 0.225926166 |
| SPAC4G8.13c | 4.491630659 | 0.239689536 |
| SPBC1539.03c | 0.317875481 | 0.265977444 |
| SPBC21H7.06c | 0.912374582 | 0.275397157 |
| SPBC8E4.01c | 0.57467891 | 0.276812664 |
| SPAC13D6.01 | 1.025352378 | 0.281091492 |
| SPBC21C3.08c | 0.590783503 | 0.282489399 |
| SPCC18.06c | 0.527524457 | 0.286707134 |
| SPCC622.11 | 0.817699011 | 0.288267672 |
| SPAC1071.07c | 0.324615271 | 0.293452744 |
| SPBC713.05 | 0.266304049 | 0.297184775 |
| SPAC17A5.16 | 0.275323834 | 0.298410954 |
| SPAC926.03 | 1.478482697 | 0.30171362 |
| SPAC140.02 | 0.713927561 | 0.306704477 |
| SPBC839.05c | 0.205118652 | 0.317336358 |
| SPAC144.11 | 0.343606517 | 0.341586699 |
| SPAC12G12.13c | 0.544554375 | 0.343575339 |
| SPCP20C8.01c | 0.644721517 | 0.349010066 |
| SPAC30C2.08 | 0.391887487 | 0.351690452 |
| SPBC21B10.06c | 0.723441653 | 0.358332792 |
| SPBC25D12.02c | 0.348690559 | 0.363512229 |
| SPAC23A1.09 | 0.860829242 | 0.364668576 |
| SPAC644.15 | 0.17812787 | 0.365021741 |
| SPAPJ696.02 | 0.659797386 | 0.376521567 |
| SPBC1734.06 | 0.963808688 | 0.388630688 |
| SPCC364.05 | 0.778115012 | 0.397839704 |
| SPAC24C9.02c | 0.475258533 | 0.413307539 |
| SPBC30B4.04c | 2.383146973 | 0.420239527 |
| SPAC1805.04 | 0.374258511 | 0.427563744 |
| SPBC8D2.04 | 0.519327895 | 0.43272039 |
| SPCC1494.09c | 0.578028631 | 0.438481589 |
| SPAC20H4.06c | 0.938872953 | 0.447574953 |
| SPCC338.12 | 0.826767034 | 0.449425368 |
| SPCC550.03c | 0.553211142 | 0.458455948 |
| SPAC3A11.10c | 1.017537093 | 0.463060733 |
| SPBC11C11.02 | 0.302761279 | 0.465060829 |
| SPCC330.14c | 0.41448468 | 0.474593054 |
| SPCC1235.05c | 0.44318959 | 0.477070187 |

| | | |
|---|---|---|
| SPBC1778.05c | 0.4628 | 0.478354978 |
| SPAPB1E7.12 | 0.418403548 | 0.480912062 |
| SPCC18B5.07c | 1.348618559 | 0.487268445 |
| SPBC16C6.10 | 0.547019574 | 0.488474388 |
| SPCC736.13 | 0.876232546 | 0.489317504 |
| SPBC530.04 | 0.724673043 | 0.490164593 |
| SPAC23C11.04c | 1.208370989 | 0.494989301 |
| SPBC16D10.11c | 0.748851328 | 0.495108316 |
| SPBC646.12c | 0.924308281 | 0.502269421 |
| SPBC409.06 | 0.733351686 | 0.502543265 |
| SPCC1739.05 | 0.279932812 | 0.510568119 |
| SPBC56F2.12 | 0.838720012 | 0.511165846 |
| SPCC4B3.02c | 0.788671809 | 0.516783668 |
| SPCC777.10c | 0.645833333 | 0.517780172 |
| SPBC12D12.06 | 0.97949402 | 0.519657352 |
| SPBC20F10.10 | 1.117133155 | 0.523451327 |
| SPAC4G9.02 | 0.622683493 | 0.525829804 |
| SPAC26H5.07c | 0.737773168 | 0.5274799 |
| SPAC27F1.08 | 0.718905841 | 0.530672328 |
| SPBC1815.01 | 0.612858908 | 0.530916481 |
| SPBC16A3.17c | 0.572091222 | 0.534027556 |
| SPAC3A12.10 | 0.711544039 | 0.538087643 |
| SPAC30D11.09 | 1.517787982 | 0.541774929 |
| SPAC1296.06 | 0.534078484 | 0.542443195 |
| SPAC3H8.03 | 0.506216196 | 0.545544965 |
| SPCC970.02 | 0.84718186 | 0.547388362 |
| SPCC622.08c | 0.397967618 | 0.54793519 |
| SPCC16C4.01 | 0.678364987 | 0.549571635 |
| SPAC13G7.13c | 0.830973813 | 0.550600572 |
| SPBC19C7.12c | 0.827981995 | 0.551032258 |
| SPAC13G7.05 | 0.687335809 | 0.557838847 |
| SPBC18H10.02 | 0.372891388 | 0.560559951 |
| SPAC17G6.15c | 0.709136866 | 0.561705814 |
| SPBC32H8.08c | 0.558708073 | 0.563716284 |
| SPAC4G8.05 | 0.57017442 | 0.564509604 |
| SPAC13A11.04c | 1.202790071 | 0.565598804 |
| SPAC1F7.07c | 1.016638306 | 0.5666268 |
| SPAC644.13c | 0.615321568 | 0.56718858 |
| SPAC694.02 | 0.670323747 | 0.567387171 |
| SPBC1709.18 | 0.656900233 | 0.570331985 |
| SPAC13C5.01c | 0.523513277 | 0.573298754 |
| SPAC2E1P3.05c | 0.809233114 | 0.57480396 |
| SPAC11D3.03c | 0.617748451 | 0.57509481 |
| SPAC2H10.01 | 0.641058496 | 0.577166498 |
| SPBC12D12.09 | 0.595112936 | 0.578669458 |
| SPAC1002.18 | 0.636723964 | 0.580026087 |
| SPBC577.14c | 0.560169423 | 0.581499313 |
| SPAC1F5.10 | 4.725368847 | 0.581653035 |

| | | |
|---|---|---|
| SPAC25B8.15c | 0.493773913 | 0.581893565 |
| SPBC887.06c | 0.576640125 | 0.582947668 |
| SPBC6B1.03c | 0.610181071 | 0.583364951 |
| SPAC4G8.10 | 0.252006168 | 0.584173207 |
| SPBP22H7.04 | 0.770439035 | 0.586614244 |
| SPCC1450.16c | 0.964690877 | 0.588706762 |
| SPAC24H6.07 | 0.554319649 | 0.588724619 |
| SPCC364.02c | 0.626610945 | 0.591947914 |
| SPBC13G1.10c | 0.707537029 | 0.592126575 |
| SPBC16E9.14c | 0.621823607 | 0.592275495 |
| SPCC663.04 | 0.87387234 | 0.592898913 |
| SPBC8D2.10c | 0.633811413 | 0.595085063 |
| SPBC2F12.12c | 0.575667023 | 0.597068436 |
| SPBC1711.11 | 0.692829896 | 0.599852723 |
| SPAC26F1.04c | 0.669001796 | 0.601087375 |
| SPBPB8B6.04c | 0.98398876 | 0.601111366 |
| SPBC16E9.15 | 0.645822103 | 0.601333653 |
| SPCC191.09c | 0.399970757 | 0.603500786 |
| SPAC3A11.08 | 0.84653746 | 0.603875422 |
| SPBC1861.07 | 0.597844367 | 0.603970588 |
| SPCC1235.13 | 0.860286225 | 0.604153846 |
| SPCC777.04 | 0.715637491 | 0.606029868 |
| SPBPB2B2.12c | 0.598487614 | 0.607487428 |
| SPBC1683.08 | 0.817274128 | 0.609674116 |
| SPAC15E1.05c | 0.487279317 | 0.610267232 |
| SPBC691.01 | 0.633989008 | 0.61033772 |
| SPAC22A12.04c | 0.434431028 | 0.610386937 |
| SPAC57A7.07c | 1.005346141 | 0.61046973 |
| SPAC15A10.03c | 0.318065365 | 0.611774292 |
| SPBC31F10.08 | 0.608688749 | 0.613024414 |
| SPBC12D12.05c | 0.928740309 | 0.614302706 |
| SPBC1347.03 | 0.716359072 | 0.61441283 |
| SPAC630.14c | 0.663900345 | 0.614873857 |
| SPAC167.06c | 0.641389985 | 0.616597471 |
| SPBC25B2.06c | 0.372611238 | 0.617757552 |
| SPAC23D3.04c | 0.776644788 | 0.618826824 |
| SPAC17C9.16c | 0.585680149 | 0.623361611 |
| SPBC27B12.07 | 0.736395904 | 0.626503722 |
| SPCC1223.09 | 0.741405197 | 0.627252333 |
| SPAC3H5.04 | 1.03431383 | 0.62730246 |
| SPAC19B12.04 | 0.77088362 | 0.627645021 |
| SPAC4A8.07c | 0.745956541 | 0.628443938 |
| SPCC1672.03c | 1.226318412 | 0.628589702 |
| SPBC713.09 | 0.728430588 | 0.628918861 |
| SPBPB2B2.10c | 0.622803523 | 0.629751649 |
| SPBC106.05c | 0.796858474 | 0.629843936 |
| SPAC23D3.03c | 0.678542966 | 0.631308489 |
| SPAC6G9.10c | 1.198979779 | 0.631331667 |

| | | |
|---|---|---|
| SPCC132.04c | 0.621644076 | 0.633032832 |
| SPBC29A3.10c | 0.592665261 | 0.633277631 |
| SPBC337.16 | 0.670033932 | 0.63332834 |
| SPCC162.03 | 0.577686989 | 0.634540541 |
| SPAC1B9.02c | 0.819145076 | 0.635066195 |
| SPBC1347.12 | 0.718809504 | 0.635590679 |
| SPBPB2B2.11 | 0.599361499 | 0.635847344 |
| SPCC1442.16c | 0.771883779 | 0.636973541 |
| SPAC11D3.05 | 0.629738796 | 0.637147488 |
| SPBC1271.11 | 0.665589223 | 0.637835596 |
| SPAC222.14c | 0.649116251 | 0.640700817 |
| SPAC664.12c | 0.959049226 | 0.642700315 |
| SPBPB2B2.05 | 0.62695724 | 0.643450252 |
| SPAC17G6.17 | 0.939454362 | 0.646742597 |
| SPAC29A4.20 | 0.819245242 | 0.647030772 |
| SPBC19G7.06 | 0.976244035 | 0.64723235 |
| SPBC16E9.07 | 0.920718816 | 0.647636364 |
| SPBC119.08 | 0.585855164 | 0.647719256 |
| SPBC530.01 | 0.457552675 | 0.647774346 |
| SPCC1753.05 | 0.707910995 | 0.648090383 |
| SPBC428.05c | 0.884665794 | 0.648163746 |
| SPBC660.10 | 0.700168977 | 0.648626263 |
| SPAC13C5.06c | 0.657389 | 0.649043241 |
| SPAC16C9.07 | 0.675997694 | 0.649979801 |
| SPAC24C9.05c | 0.615310962 | 0.650890832 |
| SPBC30B4.02c | 0.655221272 | 0.65098535 |
| SPAC16E8.18 | 0.705813632 | 0.651385522 |
| SPAC1687.09 | 0.832920006 | 0.651656974 |
| SPCC16A11.01 | 0.845959255 | 0.652706036 |
| SPAC17G8.08c | 0.672756867 | 0.653763695 |
| SPBC16A3.13 | 1.169427989 | 0.655025557 |
| SPAC19D5.02c | 0.802802446 | 0.655524517 |
| SPCC736.04c | 0.52578362 | 0.655652174 |
| SPAC13C5.05c | 0.623635031 | 0.656009527 |
| SPBC1105.09 | 0.87625042 | 0.656312488 |
| SPBP8B7.30c | 0.603205048 | 0.657133976 |
| SPBC20F10.03 | 0.728301753 | 0.658896094 |
| SPCC1450.05c | 0.676519219 | 0.659548053 |
| SPBC1289.11 | 0.759821068 | 0.659604874 |
| SPAC1F12.06c | 0.703925207 | 0.66002314 |
| SPBC83.18c | 0.815302419 | 0.660225527 |
| SPAC30.01c | 0.60677042 | 0.661731199 |
| SPAC823.15 | 0.795235478 | 0.661874092 |
| SPBC1734.05c | 0.695958894 | 0.661981655 |
| SPAC2G11.06 | 0.889565217 | 0.662119565 |
| SPAC25B8.01 | 0.687278068 | 0.662569997 |
| SPAC6G10.06 | 0.793069877 | 0.662934978 |
| SPAC6F6.02c | 0.899245139 | 0.663255738 |

| | | |
|---|---|---|
| SPCC4G3.13c | 0.668016727 | 0.66360562 |
| SPAC14C4.10c | 0.781940937 | 0.665708272 |
| SPCC584.16c | 0.790077519 | 0.667112727 |
| SPCC14G10.04 | 0.593636054 | 0.668186359 |
| SPBC1778.10c | 0.646787009 | 0.668465537 |
| SPBC8D2.18c | 0.814871274 | 0.668704906 |
| SPAC30.02c | 0.662768137 | 0.669058416 |
| SPBC725.15 | 0.66870396 | 0.669339409 |
| SPBC651.04 | 0.811890937 | 0.669956311 |
| SPBC29A3.05 | 0.761132559 | 0.670323413 |
| SPAC4A8.05c | 0.63066884 | 0.671299237 |
| SPAC13D6.03c | 0.68187444 | 0.673466748 |
| SPAC926.06c | 1.222901821 | 0.673683962 |
| SPAC31G5.11 | 0.661976013 | 0.674056236 |
| SPAC11E3.06 | 0.91255107 | 0.674250457 |
| SPBC29A10.03c | 0.973030573 | 0.674312684 |
| SPAC11E3.13c | 0.851803763 | 0.676155477 |
| SPAC23C11.02c | 0.60076189 | 0.677525749 |
| SPAC6F6.11c | 0.895210272 | 0.677572058 |
| SPAC17G8.06c | 0.794518518 | 0.678125 |
| SPAC20G4.03c | 1.003111041 | 0.679076445 |
| SPCC162.10 | 0.780724612 | 0.679619429 |
| SPAC56F8.05c | 0.753193111 | 0.679797952 |
| SPBC23E6.03c | 0.726897912 | 0.680102781 |
| SPAC20H4.09 | 0.753272314 | 0.680540492 |
| SPBC1683.04 | 0.707149203 | 0.680605964 |
| SPAC4G9.06c | 0.633046314 | 0.681757773 |
| SPAC4F8.08 | 0.794795658 | 0.68202227 |
| SPBC530.14c | 1.188278717 | 0.682325279 |
| SPCC24B10.09 | 0.656594724 | 0.682343537 |
| SPAC12G12.15 | 0.698497854 | 0.683945816 |
| SPAC664.04c | 0.594614533 | 0.685021671 |
| SPBC56F2.06 | 0.896440886 | 0.685356938 |
| SPAC1786.04 | 0.81086422 | 0.685668498 |
| SPBC354.08c | 0.721923879 | 0.685780759 |
| SPAC637.13c | 0.809302444 | 0.685890649 |
| SPBC3D6.10 | 0.648188643 | 0.686719787 |
| SPAC22E12.11c | 0.6660879 | 0.687533912 |
| SPAC6G9.09c | 0.774912172 | 0.687782086 |
| SPBC15C4.06c | 0.803278344 | 0.688119519 |
| SPCC31H12.04c | 1.053045646 | 0.688175055 |
| SPCPJ732.02c | 0.609810283 | 0.688310394 |
| SPAC589.03c | 0.51215451 | 0.688427147 |
| SPBC543.03c | 0.490635061 | 0.688865076 |
| SPCP31B10.06 | 0.315081637 | 0.689438289 |
| SPAC4C5.02c | 0.696026612 | 0.68961162 |
| SPAC1039.02 | 0.848280722 | 0.689964472 |
| SPBC20F10.07 | 1.043628463 | 0.69047768 |

| | | |
|---|---|---|
| SPAC14C4.03 | 0.668438812 | 0.690544662 |
| SPCC962.01 | 0.682012714 | 0.691027825 |
| SPBC1683.11c | 0.852898329 | 0.691121362 |
| SPAC17A5.04c | 0.984179333 | 0.691254206 |
| SPAC6G10.03c | 0.71664699 | 0.691656617 |
| SPAC26H5.09c | 0.772901921 | 0.691925831 |
| SPCC1223.02 | 0.740640825 | 0.69195432 |
| SPBC13E7.11 | 0.8019488 | 0.692093824 |
| SPCC285.13c | 1.510160307 | 0.693101942 |
| SPAC3H1.10 | 0.715896684 | 0.693194906 |
| SPAPB2B4.06 | 1.029745275 | 0.694197636 |
| SPCC320.04c | 0.689569278 | 0.694796858 |
| SPAC144.01 | 0.65767444 | 0.69483199 |
| SPAC23G3.08c | 0.748407653 | 0.696016242 |
| SPAC1142.03c | 0.586750408 | 0.696034892 |
| SPAC22F3.04 | 0.712793282 | 0.697672265 |
| SPAC22E12.01 | 0.627964427 | 0.697844614 |
| SPBC211.06 | 0.733847715 | 0.698014852 |
| SPBC83.10 | 0.807738333 | 0.69873058 |
| SPBC1198.14c | 0.341586758 | 0.698856685 |
| SPAC694.03 | 0.682505821 | 0.698938995 |
| SPAC8E11.07c | 0.683393102 | 0.698973922 |
| SPBPJ4664.03 | 0.715680273 | 0.699174279 |
| SPAC6C3.06c | 1.159148221 | 0.699325301 |
| SPAPB2B4.07 | 0.833550463 | 0.69978546 |
| SPBP23A10.05 | 0.687204564 | 0.699915953 |
| SPAC1F8.01 | 0.802087362 | 0.700156969 |
| SPBC12C2.05c | 0.769708458 | 0.70017258 |
| SPAPB8E5.02c | 0.963099928 | 0.700300568 |
| SPBC1348.01 | 0.874227335 | 0.700418344 |
| SPBC16E9.08 | 0.6597528 | 0.700463499 |
| SPBC1105.02c | 0.787853161 | 0.70185124 |
| SPAC12B10.11 | 0.651478634 | 0.702105093 |
| SPCC61.05 | 0.76256536 | 0.703118386 |
| SPAC22E12.14c | 0.650865811 | 0.703299284 |
| SPAC343.09 | 0.608800306 | 0.703464045 |
| SPBC1289.16c | 0.475273464 | 0.703896484 |
| SPAC1556.02c | 1.157189274 | 0.703939414 |
| SPAC222.15 | 0.690294394 | 0.703950278 |
| SPAP27G11.06c | 0.540943497 | 0.704085187 |
| SPAC32A11.02c | 0.647264292 | 0.704293408 |
| SPBC713.03 | 0.624872299 | 0.704580059 |
| SPAC18B11.03c | 0.778047138 | 0.704614689 |
| SPAC18B11.09c | 0.732617459 | 0.705040761 |
| SPCC4E9.02 | 0.481933336 | 0.705143165 |
| SPAC1071.09c | 0.616917218 | 0.706241018 |
| SPAC1039.10 | 0.739698668 | 0.706705501 |
| SPBC29A10.09c | 0.672489836 | 0.706807532 |

| | | |
|---|---|---|
| SPAC1071.02 | 0.691601233 | 0.706940092 |
| SPBC577.04 | 0.666067158 | 0.708128697 |
| SPAC3G6.09c | 0.776333397 | 0.708807827 |
| SPCC364.06 | 1.434487018 | 0.709409602 |
| SPAC3G9.11c | 0.794638287 | 0.709805126 |
| SPCC70.02c | 0.813452734 | 0.710090551 |
| SPBC106.19 | 1.028312212 | 0.711620605 |
| SPBC9B6.11c | 0.319026641 | 0.71218265 |
| SPAPB2B4.03 | 0.738295619 | 0.712979165 |
| SPCC777.15 | 0.723279661 | 0.713390564 |
| SPBC3E7.05c | 0.740611072 | 0.71348751 |
| SPCC162.04c | 0.761188713 | 0.713537148 |
| SPAPJ695.01c | 0.721063215 | 0.713805709 |
| SPAC3H5.05c | 0.745322189 | 0.714818406 |
| SPCPB16A4.03c | 0.794415898 | 0.714925366 |
| SPAC16E8.13 | 0.633174309 | 0.71525838 |
| SPBP8B7.23 | 0.657820918 | 0.715270866 |
| SPBC1685.07c | 0.623733184 | 0.715613611 |
| SPBC1D7.05 | 0.645751086 | 0.715728934 |
| SPCC613.01 | 0.895347693 | 0.715729198 |
| SPBC337.10c | 0.753237035 | 0.715938973 |
| SPAC14C4.15c | 0.660595529 | 0.71622343 |
| SPCC1672.12c | 0.805130098 | 0.716260643 |
| SPBC800.08 | 0.980992979 | 0.717088766 |
| SPBC18A7.02c | 0.353204322 | 0.717422242 |
| SPBC16D10.07c | 0.640688372 | 0.7176 |
| SPBP8B7.21 | 0.671752868 | 0.717750886 |
| SPBC115.02c | 0.67068289 | 0.717756128 |
| SPBC16C6.06 | 0.623126801 | 0.71818714 |
| SPCC4G3.19 | 0.744876571 | 0.719173891 |
| SPAC1039.03 | 0.677318077 | 0.719448204 |
| SPBP8B7.07c | 0.84964787 | 0.719889843 |
| SPBC15D4.15 | 0.672077357 | 0.719901257 |
| SPAC1142.08 | 0.833432886 | 0.719920072 |
| SPBP22H7.05c | 0.692013017 | 0.720135324 |
| SPAC20G8.07c | 0.774726955 | 0.720413942 |
| SPAC17G6.06 | 0.723593053 | 0.720528328 |
| SPBC800.12c | 0.753387842 | 0.720558323 |
| SPBC13E7.07 | 0.740996202 | 0.721301534 |
| SPAC458.02c | 0.956836887 | 0.721439168 |
| SPBC1773.03c | 0.69137817 | 0.7214622 |
| SPAC24H6.09 | 0.792750472 | 0.721845594 |
| SPBC1921.01c | 0.754890439 | 0.721873538 |
| SPBC3H7.07c | 0.778367441 | 0.722010503 |
| SPCC757.05c | 0.553089001 | 0.722713638 |
| SPBC29A10.06c | 0.552188086 | 0.722869225 |
| SPAC16E8.08 | 0.812262284 | 0.723103163 |
| SPAC23H3.03c | 0.690670417 | 0.723244198 |

| | | |
|---|---|---|
| SPAC22G7.08 | 0.784855273 | 0.723332774 |
| SPAC10F6.07c | 0.872820783 | 0.723355109 |
| SPCP1E11.11 | 0.800064935 | 0.723370257 |
| SPAC3H1.13 | 0.69518023 | 0.723393198 |
| SPAC23H4.02 | 0.823191093 | 0.723525059 |
| SPCC4B3.01 | 0.703162486 | 0.723657306 |
| SPAC4G8.06c | 0.708109824 | 0.723657389 |
| SPAC806.07 | 0.530329823 | 0.72377113 |
| SPAC22A12.14c | 0.596044795 | 0.723855875 |
| SPBC25H2.03 | 1.072057728 | 0.72493086 |
| SPAC16E8.17c | 0.717427993 | 0.725134851 |
| SPAC521.05 | 0.761342058 | 0.725403472 |
| SPAC2E1P5.03 | 0.665718671 | 0.726186799 |
| SPBC4F6.12 | 0.789827692 | 0.726308131 |
| SPAC4G9.20c | 0.890768873 | 0.72663909 |
| SPBC17A3.03c | 1.20478221 | 0.727530101 |
| SPAC12B10.07 | 0.81237619 | 0.728262078 |
| SPBC13E7.09 | 0.868431564 | 0.728617521 |
| SPAC2H10.02c | 0.935218124 | 0.728857184 |
| SPAC1782.06c | 0.796052263 | 0.72934107 |
| SPBC29A3.14c | 0.75513629 | 0.729651864 |
| SPCC5E4.10c | 0.718940809 | 0.730096702 |
| SPAC6B12.03c | 0.709694627 | 0.731026351 |
| SPAC2E1P3.01 | 0.712999196 | 0.731141511 |
| SPBC36B7.04 | 0.701474359 | 0.731219952 |
| SPCC4B3.08 | 0.601602374 | 0.731305638 |
| SPBC23E6.02 | 0.772582323 | 0.732354853 |
| SPAC13D6.02c | 0.888917286 | 0.73256932 |
| SPBC21C3.14c | 1.092056907 | 0.733772521 |
| SPAC1071.06 | 0.70018978 | 0.734005764 |
| SPAC869.07c | 0.727494638 | 0.734294883 |
| SPAC17G8.10c | 0.865159573 | 0.734550419 |
| SPAC3C7.13c | 0.724346522 | 0.735381972 |
| SPBC11C11.08 | 0.762763026 | 0.735403358 |
| SPBC31F10.15c | 0.928844086 | 0.735697994 |
| SPBC1198.12 | 0.757665511 | 0.736052687 |
| SPAC4F8.15 | 0.571801759 | 0.736104648 |
| SPBC354.14c | 0.442680934 | 0.736566293 |
| SPBC12C2.09c | 0.783853912 | 0.736778808 |
| SPAC3C7.08c | 1.305418917 | 0.736953025 |
| SPAC1F7.06 | 0.899312578 | 0.737402046 |
| SPCC576.02 | 0.784665797 | 0.737928396 |
| SPBC32H8.11 | 0.653389504 | 0.738218125 |
| SPCC1235.08c | 0.767344912 | 0.738273903 |
| SPAC20G4.01 | 0.755202506 | 0.738512992 |
| SPCC126.13c | 0.946139286 | 0.738570883 |
| SPAC11E3.10 | 0.990947704 | 0.73896567 |
| SPBC216.04c | 0.828111512 | 0.739430119 |

| | | |
|---|---|---|
| SPBC25B2.10 | 0.819323872 | 0.739783807 |
| SPAPJ696.01c | 0.835114635 | 0.740251767 |
| SPAC458.05 | 0.980166976 | 0.740284006 |
| SPCC18B5.05c | 1.145454545 | 0.740727273 |
| SPAC4A8.02c | 0.879167214 | 0.740831393 |
| SPAP7G5.05 | 0.735314902 | 0.740928547 |
| SPAC3C7.06c | 0.717212876 | 0.741320872 |
| SPCC1020.12c | 0.727725538 | 0.74132817 |
| SPAC12B10.09 | 0.921377349 | 0.741733441 |
| SPAC22G7.05 | 0.828200371 | 0.741929499 |
| SPBC887.05c | 0.756196245 | 0.742385843 |
| SPBC215.13 | 1.011126639 | 0.742602576 |
| SPCC1322.03 | 0.961730232 | 0.742782633 |
| SPBC25B2.03 | 0.894301125 | 0.742852071 |
| SPBC947.05c | 1.0656574 | 0.742870337 |
| SPAPB1E7.04c | 0.879964275 | 0.742885627 |
| SPAC29E6.09 | 1.020404731 | 0.742954807 |
| SPCC576.13 | 0.140388123 | 0.74328845 |
| SPAC24C9.07c | 0.769890527 | 0.744327216 |
| SPBC17D11.03c | 0.721071309 | 0.744441643 |
| SPAC589.07c | 0.613128824 | 0.744935555 |
| SPBC691.05c | 0.889938611 | 0.745234142 |
| SPAC144.13c | 0.753044611 | 0.745726148 |
| SPAC1556.04c | 0.776378036 | 0.746106742 |
| SPAC57A10.08c | 0.683512939 | 0.746590753 |
| SPCC962.05 | 0.755311837 | 0.746727306 |
| SPAC806.03c | 0.630199765 | 0.746768508 |
| SPAC8F11.10c | 1.310593752 | 0.747439782 |
| SPAC3A11.06 | 0.731282609 | 0.747520214 |
| SPAC22F3.03c | 0.572807618 | 0.747945125 |
| SPBC36B7.05c | 0.760023691 | 0.748042371 |
| SPAC11G7.04 | 0.82693725 | 0.748421532 |
| SPAC6G9.01c | 0.820875378 | 0.748705876 |
| SPBC2G2.02 | 0.387917637 | 0.749001656 |
| SPAC2F7.03c | 0.702062807 | 0.749793593 |
| SPAC17G8.14c | 0.9420216 | 0.749844502 |
| SPBC2A9.05c | 0.812777166 | 0.750110458 |
| SPCC285.15c | 0.596301218 | 0.750641201 |
| SPBC23E6.08 | 0.942810645 | 0.750923601 |
| SPAC25B8.09 | 0.759811978 | 0.751047172 |
| SPBC2G5.04c | 0.891238766 | 0.752149739 |
| SPAC13F5.01c | 0.886339184 | 0.753765258 |
| SPAC22H10.13 | 0.68227886 | 0.753794978 |
| SPAC5H10.12c | 0.686388344 | 0.754313253 |
| SPCC126.07c | 0.696532572 | 0.754934596 |
| SPAC630.04c | 0.698445152 | 0.75564557 |
| SPAC29A4.11 | 0.708485535 | 0.755809796 |
| SPAC1F8.08 | 0.795155815 | 0.755847692 |

| | | |
|---|---|---|
| SPAC4H3.14c | 0.71753668 | 0.756120172 |
| SPCC663.03 | 0.750918076 | 0.756541387 |
| SPAC29A4.02c | 0.802680067 | 0.756790412 |
| SPAC30.03c | 0.793121958 | 0.756841849 |
| SPBP35G2.06c | 0.816110806 | 0.757179367 |
| SPAC25B8.10 | 0.814926985 | 0.757309733 |
| SPCP31B10.05 | 0.818792104 | 0.75734005 |
| SPAC1002.20 | 1.02360371 | 0.757468887 |
| SPBC1683.03c | 0.694574307 | 0.758130918 |
| SPBP8B7.09c | 0.741829673 | 0.758181932 |
| SPAP27G11.02 | 0.564341085 | 0.758742138 |
| SPBC336.14c | 0.96131326 | 0.75886845 |
| SPAPJ691.03 | 1.036929313 | 0.758918861 |
| SPAC30D11.01c | 0.784469063 | 0.759247221 |
| SPBP26C9.02c | 0.40966371 | 0.759256963 |
| SPBC428.11 | 0.805877923 | 0.75992211 |
| SPAC3C7.02c | 0.706995705 | 0.760285886 |
| SPBC27.06c | 0.784371391 | 0.760765338 |
| SPBC1703.06 | 0.823527447 | 0.760977517 |
| SPCC4B3.05c | 0.786565458 | 0.76130896 |
| SPCC11E10.08 | 0.932184148 | 0.761335826 |
| SPBC3F6.05 | 0.759633501 | 0.761511716 |
| SPCC830.04c | 0.873160411 | 0.761517323 |
| SPBC146.06c | 0.964529191 | 0.761821825 |
| SPBC3E7.07c | 0.78952212 | 0.762081222 |
| SPAC23H3.12c | 0.738225932 | 0.762169199 |
| SPAC13G6.01c | 0.883005486 | 0.7622476 |
| SPBC1347.06c | 0.713674276 | 0.762309155 |
| SPCC126.06 | 0.807417504 | 0.762514457 |
| SPAC16.05c | 0.71125 | 0.762890625 |
| SPAC18G6.02c | 0.832237833 | 0.76321909 |
| SPBC16H5.11c | 0.739093328 | 0.763239335 |
| SPAC926.05c | 0.692859218 | 0.763805198 |
| SPAC10F6.16 | 0.672236068 | 0.763992695 |
| SPAC16C9.01c | 0.831807952 | 0.764129032 |
| SPCC126.12 | 0.830374567 | 0.764754486 |
| SPAC5D6.02c | 0.678697097 | 0.764879814 |
| SPAC2E1P3.04 | 0.695358098 | 0.764886411 |
| SPAC1002.01 | 0.758703445 | 0.764994377 |
| SPBC3B8.03 | 0.887588762 | 0.765531001 |
| SPBC19C7.08c | 0.73591535 | 0.765602735 |
| SPAC1002.17c | 0.9470776 | 0.765785467 |
| SPBC1271.10c | 0.768755595 | 0.765806849 |
| SPCC576.04 | 0.73613592 | 0.765909712 |
| SPAC14C4.09 | 0.72512964 | 0.765937801 |
| SPAC1751.01c | 0.696401129 | 0.766278718 |
| SPBC4C3.09 | 0.785396704 | 0.766385589 |
| SPBC25D12.05 | 0.799114826 | 0.766587721 |

| | | |
|---|---|---|
| SPCC663.10 | 1.36144542 | 0.76671336 |
| SPBC21D10.08c | 0.808138805 | 0.76690872 |
| SPAC1751.04 | 0.827610969 | 0.767151095 |
| SPBC15D4.12c | 0.637811251 | 0.767387642 |
| SPCC162.02c | 0.470638986 | 0.767442126 |
| SPBC8E4.03 | 0.490959279 | 0.768021758 |
| SPCC736.11 | 0.447201385 | 0.768068379 |
| SPAC4G8.04 | 0.772942509 | 0.768220586 |
| SPAC20H4.10 | 0.731146511 | 0.768434685 |
| SPAC11E3.09 | 0.901953021 | 0.768445729 |
| SPCC622.15c | 0.74528872 | 0.768834269 |
| SPBC776.14 | 0.755696604 | 0.769557382 |
| SPAC222.12c | 1.060273272 | 0.769704858 |
| SPCC417.12 | 1.081326787 | 0.769857145 |
| SPBC16C6.05 | 0.84740638 | 0.769994732 |
| SPAC22H12.05c | 0.986936911 | 0.770176323 |
| SPAC8C9.11 | 0.802676072 | 0.770653908 |
| SPCC63.06 | 0.789763578 | 0.77114315 |
| SPBC713.02c | 0.800027231 | 0.771160943 |
| SPAC806.08c | 0.780019275 | 0.771397134 |
| SPAC18B11.08c | 0.790840272 | 0.772213285 |
| SPBP4H10.19c | 0.764987895 | 0.772220033 |
| SPAC11E3.12 | 1.02613423 | 0.772352068 |
| SPBC651.12c | 0.794846328 | 0.772940554 |
| SPAC30C2.02 | 1.111238674 | 0.772944221 |
| SPBC119.12 | 0.702362772 | 0.773092512 |
| SPBC2D10.06 | 0.869599645 | 0.77330836 |
| SPBPJ4664.05 | 0.773711543 | 0.773598467 |
| SPBC32F12.03c | 0.564530588 | 0.773713902 |
| SPCC297.06c | 0.770108695 | 0.773771764 |
| SPACUNK4.15 | 0.669412534 | 0.77411552 |
| SPACUNK4.17 | 1.061430424 | 0.774206613 |
| SPAC1805.09c | 0.596571848 | 0.775416294 |
| SPAC25G10.02 | 0.656182461 | 0.775546039 |
| SPCC1393.09c | 0.501973109 | 0.775813072 |
| SPAC5H10.06c | 0.87769033 | 0.775849688 |
| SPCC1442.02 | 0.927108927 | 0.775901876 |
| SPCC24B10.22 | 0.777386411 | 0.776013148 |
| SPCC1322.12c | 0.592764858 | 0.776501938 |
| SPAC1296.01c | 1.107973964 | 0.776880195 |
| SPAC8E11.10 | 0.684750282 | 0.77705665 |
| SPAC25H1.06 | 1.014584871 | 0.777439657 |
| SPAP8A3.04c | 0.735159336 | 0.777807098 |
| SPBC365.12c | 0.748799872 | 0.777977933 |
| SPCC18B5.01c | 0.970049965 | 0.778042622 |
| SPAC5D6.05 | 0.968044212 | 0.778159195 |
| SPCC613.03 | 0.720843922 | 0.778284536 |
| SPAPB2C8.01 | 0.732178038 | 0.778294432 |

| | | |
|---|---|---|
| SPAC17A5.18c | 0.752003568 | 0.778295967 |
| SPAC328.03 | 0.713878469 | 0.778322581 |
| SPCC4B3.15 | 0.765624226 | 0.778517723 |
| SPCC1827.07c | 0.791958533 | 0.778728195 |
| SPBC947.11c | 0.699648212 | 0.778750965 |
| SPAC922.04 | 0.794888109 | 0.779264151 |
| SPAC7D4.14c | 0.751124522 | 0.779330014 |
| SPCC417.05c | 0.999726402 | 0.779945701 |
| SPCC1259.09c | 0.857909916 | 0.780067446 |
| SPAC6G9.03c | 0.782656461 | 0.780219825 |
| SPAC3A11.14c | 0.782417988 | 0.780790746 |
| SPAC6G9.16c | 0.770137194 | 0.781547516 |
| SPCC330.12c | 0.847355393 | 0.781645994 |
| SPBC336.03 | 0.800160578 | 0.781883955 |
| SPBC1105.05 | 1.042921164 | 0.782060936 |
| SPAC20G8.04c | 0.860848322 | 0.782309536 |
| SPAC22F8.07c | 0.763409771 | 0.782340479 |
| SPAC3H8.07c | 0.587723445 | 0.782496276 |
| SPCC1840.05c | 0.935935766 | 0.782755031 |
| SPAC1039.04 | 0.7681749 | 0.782864149 |
| SPBC9B6.07 | 1.093967143 | 0.783239745 |
| SPCC1919.05 | 0.688059586 | 0.783268033 |
| SPCC1020.13c | 0.754412214 | 0.783578244 |
| SPAC23H3.13c | 0.548564057 | 0.78358832 |
| SPACUNK4.12c | 0.877805045 | 0.783666764 |
| SPBC29B5.04c | 0.911574571 | 0.7837111 |
| SPAC23C4.17 | 0.841385736 | 0.784349081 |
| SPAC29B12.13 | 0.796419598 | 0.785892352 |
| SPCC737.09c | 0.931674282 | 0.786002464 |
| SPAC13F5.03c | 0.217632826 | 0.786214961 |
| SPAC5H10.10 | 0.994426137 | 0.786671478 |
| SPAC29B12.11c | 0.883924392 | 0.78721808 |
| SPBC1685.11 | 0.782409178 | 0.787225143 |
| SPAC23D3.11 | 1.201018754 | 0.787422477 |
| SPAC6C3.03c | 0.770845813 | 0.788156114 |
| SPCC965.11c | 0.753497627 | 0.788202521 |
| SPAC23G3.02c | 0.846526046 | 0.788203896 |
| SPBC21D10.09c | 0.878175517 | 0.788228533 |
| SPAC9G1.07 | 0.760571361 | 0.78871066 |
| SPAC1B3.15c | 0.866416017 | 0.788762858 |
| SPAC869.05c | 0.739560475 | 0.789136519 |
| SPAC1834.09 | 0.909802217 | 0.789233645 |
| SPBC29A10.14 | 0.809664095 | 0.789330469 |
| SPAC19B12.09 | 0.772381288 | 0.789342732 |
| SPAC750.05c | 0.921110177 | 0.789937151 |
| SPAC26F1.09 | 0.854355872 | 0.789957419 |
| SPCC1919.12c | 0.710384737 | 0.790097947 |
| SPCC895.08c | 0.710717621 | 0.790293256 |

| | | |
|---|---|---|
| SPAC31A2.09c | 0.897323948 | 0.790293742 |
| SPAPB1A10.05 | 0.611053633 | 0.790416436 |
| SPCC569.08c | 0.655058316 | 0.790482028 |
| SPAC1952.12c | 0.68372093 | 0.790758621 |
| SPCC1281.08 | 0.826666667 | 0.790979381 |
| SPBP8B7.18c | 0.794203422 | 0.791034513 |
| SPCC417.03 | 0.824672645 | 0.791179942 |
| SPBC725.09c | 0.639373966 | 0.791229087 |
| SPAC1687.13c | 0.592668527 | 0.791673525 |
| SPCPB16A4.02c | 0.873964338 | 0.79182171 |
| SPAC1399.02 | 0.836481989 | 0.792020547 |
| SPBC3E7.06c | 0.787431783 | 0.792417336 |
| SPCC550.15c | 0.767564833 | 0.792556601 |
| SPAC8C9.16c | 0.839317132 | 0.792715754 |
| SPBC19G7.01c | 0.881001183 | 0.792717777 |
| SPAC3H8.08c | 1.43728665 | 0.792971175 |
| SPCC4G3.05c | 0.764159684 | 0.793026311 |
| SPAC227.03c | 0.705372331 | 0.793034333 |
| SPBC4F6.11c | 0.626870221 | 0.793420093 |
| SPAC16.01 | 0.835161826 | 0.793426378 |
| SPBP4G3.02 | 0.660809669 | 0.794059656 |
| SPAC11D3.06 | 0.915849826 | 0.794215427 |
| SPBP4H10.16c | 0.893836614 | 0.794252852 |
| SPAC17C9.11c | 0.87563856 | 0.794851494 |
| SPBC1773.06c | 0.677101794 | 0.794858952 |
| SPCC663.06c | 1.240713441 | 0.794938983 |
| SPBC887.01 | 0.901284339 | 0.795096195 |
| SPAPB8E5.05 | 0.912597901 | 0.795244445 |
| SPAC4D7.02c | 0.802486533 | 0.795359663 |
| SPBP8B7.06 | 0.78348875 | 0.795598665 |
| SPAC144.17c | 0.89755712 | 0.795625137 |
| SPAC20H4.07 | 0.884629878 | 0.79581721 |
| SPAC23G3.10c | 0.820384726 | 0.795969136 |
| SPCC737.04 | 0.739932387 | 0.796126129 |
| SPBC18H10.10c | 0.82424505 | 0.796193235 |
| SPAC4C5.01 | 0.717459187 | 0.796222292 |
| SPBC1703.03c | 0.952266428 | 0.79641969 |
| SPBC685.06 | 0.871637373 | 0.796769417 |
| SPAC56F8.09 | 0.786513196 | 0.797563125 |
| SPAC3H1.12c | 0.835743077 | 0.797733023 |
| SPAC19A8.04 | 0.795297667 | 0.797733469 |
| SPAC139.03 | 0.775221141 | 0.798114902 |
| SPAC23H3.08c | 0.794870368 | 0.798535885 |
| SPCC31H12.03c | 0.753665208 | 0.798626024 |
| SPAC32A11.03c | 0.846418275 | 0.798746112 |
| SPCC1442.17c | 0.462140089 | 0.799383949 |
| SPAC1782.11 | 0.780040382 | 0.799591496 |
| SPAC694.06c | 0.843475397 | 0.799657813 |

| | | |
|---|---|---|
| SPBC26H8.08c | 1.149804512 | 0.799686028 |
| SPCC1223.03c | 1.107840736 | 0.800562252 |
| SPAC29E6.10c | 0.687572842 | 0.800564543 |
| SPAP27G11.12 | 0.840159075 | 0.800595043 |
| SPAC2F7.06c | 0.805083448 | 0.800627404 |
| SPAPB17E12.02 | 0.84981412 | 0.801442798 |
| SPAC821.03c | 0.909190093 | 0.801535396 |
| SPAC11H11.05c | 0.77388837 | 0.801749153 |
| SPAC1039.09 | 0.765350422 | 0.801803137 |
| SPCC962.04 | 0.856305693 | 0.801931931 |
| SPAC2F7.11 | 0.895222069 | 0.802192807 |
| SPAC6G10.08 | 1.11179168 | 0.802338028 |
| SPAC4G9.14 | 0.922734145 | 0.802933998 |
| SPAC22F3.07c | 0.901924255 | 0.802974478 |
| SPBC530.15c | 0.727357862 | 0.803029314 |
| SPAC2C4.10c | 1.308474633 | 0.803149708 |
| SPBC1685.15c | 0.800999473 | 0.803154178 |
| SPBC1306.02 | 0.547646215 | 0.80343646 |
| SPBC1348.14c | 0.960371904 | 0.803642575 |
| SPCC613.11c | 0.827675975 | 0.80377064 |
| SPBC115.03 | 0.908744344 | 0.803872598 |
| SPBC660.17c | 1.348841407 | 0.804216606 |
| SPCC1919.13c | 0.928878968 | 0.804224215 |
| SPBC1685.01 | 0.899616506 | 0.804430113 |
| SPBC119.03 | 0.748660934 | 0.804662696 |
| SPAC3A11.11c | 0.760277275 | 0.804710972 |
| SPAC21E11.03c | 0.69187468 | 0.804792176 |
| SPBC14F5.09c | 0.90213712 | 0.804867784 |
| SPAC1A6.06c | 0.809491682 | 0.805093381 |
| SPAC25H1.07 | 1.20914344 | 0.805094905 |
| SPBC16G5.07c | 1.164004689 | 0.805153271 |
| SPAC4H3.05 | 0.670208024 | 0.8053724 |
| SPCPB1C11.03 | 0.67841902 | 0.806137356 |
| SPAC56F8.14c | 0.848723958 | 0.806241246 |
| SPAC17A5.09c | 0.940275006 | 0.806365055 |
| SPCC1020.03 | 0.876837829 | 0.807048695 |
| SPBC1289.14 | 0.714763647 | 0.807117091 |
| SPCC1183.04c | 0.888865248 | 0.807351712 |
| SPAC343.12 | 0.834226094 | 0.807424525 |
| SPBC887.08 | 0.878430273 | 0.808091223 |
| SPAPB1A11.01 | 0.678693841 | 0.808212223 |
| SPAC343.18 | 0.875428788 | 0.808277603 |
| SPAC12B10.06c | 0.774445342 | 0.808774383 |
| SPBP18G5.03 | 0.715047687 | 0.808945602 |
| SPAC19G12.05 | 0.855486238 | 0.809483945 |
| SPAC17D4.01 | 0.457242601 | 0.809851081 |
| SPCC63.04 | 0.856737589 | 0.809949298 |
| SPBC543.10 | 0.838284779 | 0.810456912 |

| | | |
|---|---|---|
| SPBC8D2.03c | 0.821321131 | 0.811405192 |
| SPAC513.05 | 0.899650638 | 0.811688697 |
| SPAC3F10.06c | 0.748828105 | 0.812045928 |
| SPBCPT2R1.03 | 0.919535902 | 0.812115767 |
| SPCC1322.07c | 0.744134615 | 0.812146187 |
| SPAC11E3.01c | 0.969676235 | 0.812235263 |
| SPCC663.15c | 0.899166261 | 0.812532322 |
| SPBC3E7.15c | 0.932722583 | 0.813210874 |
| SPBC23G7.10c | 0.77422035 | 0.813234359 |
| SPBC16G5.06 | 0.905098761 | 0.813287035 |
| SPBC16E9.18 | 0.675426775 | 0.813384434 |
| SPCC569.07 | 0.740512404 | 0.814213671 |
| SPAC1F12.05 | 0.770441887 | 0.814235425 |
| SPCC550.09 | 0.858606272 | 0.814274191 |
| SPAC821.10c | 0.678196982 | 0.814495029 |
| SPCC594.01 | 0.748312308 | 0.814503069 |
| SPAC1006.01 | 0.901 | 0.8146875 |
| SPAC12G12.01c | 0.581256048 | 0.815599105 |
| SPAC1952.05 | 0.735474549 | 0.81570176 |
| SPBPB10D8.02c | 0.793408133 | 0.815732563 |
| SPAC926.07c | 0.741875294 | 0.815867745 |
| SPBC36.07 | 0.766362888 | 0.816130074 |
| SPAC14C4.07 | 0.85368871 | 0.816363453 |
| SPAC57A10.09c | 0.940711723 | 0.816914877 |
| SPCC576.01c | 0.995921753 | 0.816969974 |
| SPBC31F10.13c | 0.616110305 | 0.817568595 |
| SPCC737.03c | 0.686327151 | 0.817598905 |
| SPBC776.16 | 1.232552687 | 0.81761313 |
| SPBC1709.16c | 1.169330643 | 0.8177794 |
| SPBC23G7.07c | 0.872025694 | 0.81782988 |
| SPAPB24D3.10c | 0.740365554 | 0.818259811 |
| SPBC25H2.05 | 0.733207983 | 0.818499738 |
| SPBC1604.07 | 0.82377261 | 0.818666667 |
| SPAC9G1.08c | 0.507329372 | 0.818793025 |
| SPBC15D4.13c | 0.786075486 | 0.818946958 |
| SPCC1840.08c | 0.870435772 | 0.819109895 |
| SPAC4F10.19c | 0.953823371 | 0.819214532 |
| SPCC622.18 | 0.872937185 | 0.819284209 |
| SPBC28E12.06c | 0.890787111 | 0.819343676 |
| SPAC1527.02 | 0.546812941 | 0.819364202 |
| SPBC543.05c | 0.783968258 | 0.81946656 |
| SPBPB21E7.01c | 0.7873479 | 0.820165455 |
| SPAC13G6.04 | 0.914949354 | 0.820206078 |
| SPAC1399.01c | 0.917577108 | 0.820257817 |
| SPCC1223.05c | 0.941204674 | 0.820321932 |
| SPBC21C3.06 | 0.882053931 | 0.820366685 |
| SPAC9.11 | 0.804793297 | 0.820723236 |
| SPAC821.07c | 0.880825628 | 0.820902156 |

| | | |
|---|---|---|
| SPCC24B10.03 | 0.765719658 | 0.821073151 |
| SPAC1687.14c | 0.75758427 | 0.821464624 |
| SPBC30D10.09c | 0.895431805 | 0.821873845 |
| SPAC18G6.01c | 0.901540037 | 0.821891903 |
| SPAC637.03 | 0.688839844 | 0.822093484 |
| SPBC19G7.17 | 0.891388343 | 0.822110933 |
| SPBP35G2.02 | 0.838084217 | 0.822231128 |
| SPCC645.07 | 0.811955416 | 0.822298388 |
| SPBC11B10.05c | 1.006980483 | 0.822304535 |
| SPAC2F7.07c | 0.940912755 | 0.822400039 |
| SPAC1F7.11c | 0.732559899 | 0.822709175 |
| SPAC1B3.05 | 0.721115837 | 0.822818042 |
| SPAPJ691.02 | 0.737848917 | 0.822931785 |
| SPBC1773.08c | 0.860645622 | 0.822942238 |
| SPBC1685.02c | 0.976785766 | 0.823046052 |
| SPAC1D4.02c | 0.914327553 | 0.823084788 |
| SPAC23C4.08 | 0.795598743 | 0.823119105 |
| SPAC23C4.07 | 1.308894932 | 0.823263824 |
| SPCC777.08c | 0.877573143 | 0.823403226 |
| SPBC12C2.07c | 0.739336493 | 0.823620853 |
| SPBC2F12.03c | 0.850533783 | 0.824445716 |
| SPAC1D4.05c | 0.81129825 | 0.824474358 |
| SPAC212.02 | 0.778581978 | 0.824542756 |
| SPBC651.06 | 0.852100596 | 0.824597261 |
| SPBC1198.07c | 0.86164726 | 0.824665532 |
| SPBC11C11.11c | 0.365447543 | 0.824858075 |
| SPBC23G7.11 | 0.90848908 | 0.825620169 |
| SPAC8C9.04 | 1.025697339 | 0.825639144 |
| SPBPB21E7.09 | 0.802645294 | 0.825666731 |
| SPCC1795.10c | 0.808477532 | 0.825711077 |
| SPAC22G7.11c | 0.827857287 | 0.82613822 |
| SPBC1A4.02c | 0.771392767 | 0.826194951 |
| SPAC31A2.12 | 0.646124167 | 0.826212721 |
| SPBC8D2.16c | 0.77624925 | 0.826535525 |
| SPAC29A4.13 | 0.900380361 | 0.826712294 |
| SPAC589.09 | 1.036838153 | 0.826723155 |
| SPBC2F12.09c | 0.821116866 | 0.827094185 |
| SPAC15E1.07c | 0.885415965 | 0.827140872 |
| SPBC16C6.08c | 0.855842897 | 0.827264082 |
| SPAC19D5.03 | 0.824511787 | 0.82730569 |
| SPBC18H10.20c | 0.782997225 | 0.827394834 |
| SPCC1322.16 | 0.78102401 | 0.827443361 |
| SPBC839.11c | 0.706406639 | 0.827504919 |
| SPCC1183.11 | 0.788528357 | 0.827634921 |
| SPBC1198.09 | 0.775881684 | 0.827680603 |
| SPAC22E12.04 | 0.97689152 | 0.827746068 |
| SPAC4H3.06 | 0.917493436 | 0.827847593 |
| SPCC645.13 | 0.929447274 | 0.827909744 |

| | | |
|---|---|---|
| SPAC1565.03 | 0.837703289 | 0.827979345 |
| SPBC543.02c | 0.839667253 | 0.828028719 |
| SPBC1778.09 | 0.929810434 | 0.828051471 |
| SPAC4G9.11c | 0.834137855 | 0.828154772 |
| SPBC1773.13 | 1.195528046 | 0.828539502 |
| SPCC1739.09c | 1.1036699 | 0.828763111 |
| SPCC70.03c | 0.924784224 | 0.828887184 |
| SPBC11B10.02c | 0.997908172 | 0.828911432 |
| SPCC74.09 | 0.885402744 | 0.829042825 |
| SPBC1773.16c | 0.825835755 | 0.829043533 |
| SPCC338.02 | 0.94992163 | 0.829055643 |
| SPBC16E9.06c | 0.908970135 | 0.829131796 |
| SPAC6F6.09 | 0.677381445 | 0.82931779 |
| SPBC530.09c | 0.739801165 | 0.829327295 |
| SPCPB16A4.05c | 0.821380184 | 0.829389219 |
| SPAC22H10.02 | 0.224662444 | 0.829664279 |
| SPAC11E3.03 | 0.604463875 | 0.829864964 |
| SPCC1795.02c | 0.769703901 | 0.830363405 |
| SPAC5H10.05c | 0.812349094 | 0.830365335 |
| SPBP4H10.08 | 0.873426564 | 0.830629135 |
| SPCC965.14c | 0.751088697 | 0.830651925 |
| SPCP1E11.05c | 0.744792706 | 0.831514966 |
| SPAC2C4.06c | 1.217267044 | 0.83164092 |
| SPAC1002.02 | 0.725860895 | 0.832040062 |
| SPAC11D3.01c | 0.854775177 | 0.832671405 |
| SPAC19A8.10 | 0.778615848 | 0.832698051 |
| SPAC1327.01c | 0.564800902 | 0.833008546 |
| SPAC17G8.07 | 0.85397849 | 0.833141078 |
| SPBC32F12.12c | 0.875489782 | 0.833214787 |
| SPAPJ760.02c | 0.838274781 | 0.833377161 |
| SPAC17H9.03c | 0.724235808 | 0.833378821 |
| SPAC328.09 | 0.771925595 | 0.833492248 |
| SPAC24C9.12c | 0.641000134 | 0.833903661 |
| SPAC1420.03 | 1.016313085 | 0.834094172 |
| SPBC9B6.03 | 0.730461073 | 0.834126132 |
| SPBC17G9.05 | 0.801733035 | 0.834201262 |
| SPCC4B3.14 | 0.626976295 | 0.834611301 |
| SPAC1F3.09 | 0.89491497 | 0.834819374 |
| SPAC5H10.04 | 0.836260162 | 0.835522876 |
| SPBC25B2.07c | 1.254354765 | 0.836078431 |
| SPAC1002.12c | 0.921966704 | 0.836490251 |
| SPBP8B7.25 | 0.735231615 | 0.836565097 |
| SPBC577.05c | 0.742508203 | 0.837016952 |
| SPCC825.01 | 0.619392584 | 0.83705945 |
| SPAC22E12.06c | 0.801631913 | 0.837206546 |
| SPAC7D4.12c | 0.828415685 | 0.837317615 |
| SPAC15A10.16 | 0.947042596 | 0.837665147 |
| SPBC651.05c | 0.621758618 | 0.837708043 |

| | | |
|---|---|---|
| SPBC1105.10 | 0.141658591 | 0.837827241 |
| SPBC216.02 | 0.807562192 | 0.837856849 |
| SPAC1952.16 | 0.821198146 | 0.837986511 |
| SPAC1A6.03c | 0.527853211 | 0.838279816 |
| SPBC582.05c | 0.892989262 | 0.83829667 |
| SPBC776.03 | 0.749548382 | 0.838461937 |
| SPAC57A10.12c | 0.685048 | 0.838525669 |
| SPAC13G7.12c | 0.768279688 | 0.838660038 |
| SPAC26F1.10c | 0.903506118 | 0.838842904 |
| SPAC5D6.10c | 0.947759496 | 0.838920942 |
| SPAC3H1.06c | 0.885489305 | 0.839244442 |
| SPAC2G11.07c | 1.131210322 | 0.839324661 |
| SPAC186.08c | 0.945547463 | 0.839402135 |
| SPBC32F12.08c | 1.045002791 | 0.839481312 |
| SPAC16A10.04 | 0.841613025 | 0.839632242 |
| SPAP11E10.01 | 0.770590039 | 0.839765792 |
| SPBC1711.03 | 0.818625423 | 0.839813374 |
| SPCC24B10.15 | 0.834756097 | 0.839957627 |
| SPBC21C3.15c | 0.736272745 | 0.840364887 |
| SPAC343.07 | 0.742965081 | 0.840376938 |
| SPCC126.01c | 1.206426755 | 0.840521065 |
| SPBC1703.08c | 0.903579583 | 0.841043975 |
| SPAC27D7.04 | 0.865794979 | 0.841069561 |
| SPAC23A1.02c | 0.887270901 | 0.841302096 |
| SPBC106.04 | 0.989084802 | 0.841860831 |
| SPCC320.06 | 0.869774501 | 0.841879878 |
| SPBC2D10.04 | 0.71660799 | 0.841959921 |
| SPAC212.04c | 0.851986846 | 0.842015803 |
| SPAPYUG7.03c | 1.040618198 | 0.84205752 |
| SPCC1183.02 | 1.005119469 | 0.842528427 |
| SPBC839.07 | 0.959472717 | 0.842792206 |
| SPCC74.02c | 0.679111187 | 0.842842717 |
| SPAC11D3.14c | 0.790335793 | 0.843113552 |
| SPAC139.02c | 0.92018346 | 0.843139776 |
| SPAP27G11.16 | 0.738503904 | 0.843848901 |
| SPBC17G9.10 | 0.962581717 | 0.843972432 |
| SPAC1687.10 | 1.096616973 | 0.844304421 |
| SPAC57A10.04 | 1.054588608 | 0.845203718 |
| SPAP8A3.13c | 0.462837645 | 0.845423612 |
| SPAC1786.01c | 0.868776078 | 0.845803633 |
| SPCC1682.12c | 0.876949942 | 0.845823632 |
| SPAC3H1.03 | 0.923935791 | 0.845915827 |
| SPCC338.18 | 0.768823856 | 0.84617258 |
| SPAPB17E12.04c | 0.829524438 | 0.846272717 |
| SPAC821.06 | 0.863517527 | 0.847011414 |
| SPCC132.02 | 0.93792662 | 0.847063507 |
| SPAC1783.01 | 0.958524637 | 0.847066668 |
| SPCC737.05 | 0.770041631 | 0.847197609 |

| | | |
|---|---|---|
| SPAC1783.02c | 0.715929878 | 0.847287062 |
| SPCC1795.09 | 0.701609113 | 0.847374129 |
| SPAC644.08 | 0.888130969 | 0.847689291 |
| SPAC5D6.07c | 0.604603887 | 0.847815481 |
| SPAC343.06c | 0.858246374 | 0.848564254 |
| SPAC3F10.02c | 0.718682167 | 0.848866657 |
| SPCC1223.04c | 0.760196966 | 0.849178273 |
| SPCC736.09c | 0.878749103 | 0.849652003 |
| SPBC36.06c | 1.336597307 | 0.84967852 |
| SPAC19D5.01 | 0.898825253 | 0.849734091 |
| SPCC23B6.02c | 0.90505683 | 0.849837315 |
| SPACUNK4.14 | 0.877371274 | 0.850183824 |
| SPBC13E7.08c | 0.687048329 | 0.850450562 |
| SPBC582.06c | 0.918007532 | 0.850682557 |
| SPAC1142.06 | 0.821364212 | 0.850754812 |
| SPAC4F8.11 | 0.966317447 | 0.85109677 |
| SPAC1F5.08c | 1.048441811 | 0.851152034 |
| SPAC10F6.17c | 0.930964164 | 0.851390551 |
| SPAC12B10.14c | 0.877928335 | 0.852003147 |
| SPBC337.07c | 0.753885561 | 0.852278713 |
| SPAC1952.10c | 0.787952289 | 0.852859599 |
| SPBC28E12.02 | 0.865948897 | 0.852986122 |
| SPBC800.11 | 1.100488535 | 0.853283764 |
| SPBC2G2.15c | 0.909120507 | 0.853496945 |
| SPCC1442.01 | 0.806904534 | 0.853720119 |
| SPBC1773.04 | 0.788399549 | 0.853736773 |
| SPBP35G2.13c | 1.450100446 | 0.853774524 |
| SPAC1610.03c | 0.997492233 | 0.854058981 |
| SPAC186.07c | 0.814540817 | 0.855776239 |
| SPBC31F10.05 | 0.889568302 | 0.855982304 |
| SPCC16C4.06c | 0.887096141 | 0.856407935 |
| SPCC825.04c | 0.83055389 | 0.856484465 |
| SPBC19C2.02 | 0.861306153 | 0.856844564 |
| SPCC16C4.04 | 0.81325427 | 0.856953284 |
| SPAC3H8.10 | 0.916653709 | 0.857115446 |
| SPBC409.11 | 0.853360001 | 0.857185449 |
| SPAC20G8.08c | 0.940144372 | 0.857538256 |
| SPBC1773.14 | 0.936099851 | 0.857562359 |
| SPBC29A10.02 | 0.866036357 | 0.857575758 |
| SPAC27F1.06c | 0.931069472 | 0.85761123 |
| SPBC18E5.10 | 0.88942206 | 0.857860186 |
| SPCC70.04c | 1.341278244 | 0.857889257 |
| SPAC8C9.03 | 1.080256589 | 0.858112445 |
| SPBP35G2.10 | 0.899625934 | 0.858437644 |
| SPBC660.06 | 0.784984854 | 0.858796243 |
| SPCC1322.06 | 0.826882845 | 0.859309623 |
| SPBC4.02c | 0.82963194 | 0.859365253 |
| SPAC25A8.03c | 0.89425093 | 0.859982668 |

| | | |
|---|---|---|
| SPCP31B10.02 | 0.781399573 | 0.86003811 |
| SPBC1198.03c | 0.884129854 | 0.860410317 |
| SPBC8D2.11 | 0.889659125 | 0.860613422 |
| SPBC18H10.18c | 0.861321924 | 0.860670312 |
| SPBC17D11.02c | 1.245593804 | 0.860978836 |
| SPAC1296.04 | 1.105787719 | 0.860999005 |
| SPBC530.07c | 0.858626944 | 0.861001865 |
| SPAC13A11.03 | 1.078156289 | 0.861055398 |
| SPAC4G9.13c | 0.827146983 | 0.861228758 |
| SPAC19G12.03 | 0.958437146 | 0.861233013 |
| SPAC4D7.07c | 0.801265958 | 0.861398176 |
| SPAC26A3.14c | 0.808387343 | 0.862013915 |
| SPCC794.09c | 0.935700535 | 0.862537448 |
| SPCC18.02 | 0.970514288 | 0.862875688 |
| SPAC17A5.10 | 0.806218235 | 0.863294702 |
| SPCC757.12 | 0.843668084 | 0.863418732 |
| SPCC1902.02 | 0.548817782 | 0.863541351 |
| SPAC1F3.06c | 0.773369084 | 0.863545192 |
| SPAC31A2.15c | 0.878556015 | 0.863586185 |
| SPAC821.04c | 0.922619474 | 0.863626998 |
| SPCC736.02 | 0.895696551 | 0.863857081 |
| SPAC2C4.07c | 0.944640355 | 0.863860963 |
| SPAC1F7.10 | 0.783220016 | 0.8638921 |
| SPBC428.04 | 0.868664086 | 0.864100796 |
| SPAC664.14 | 0.879039143 | 0.864217266 |
| SPCC1259.08 | 0.722940036 | 0.86421842 |
| SPAPB24D3.07c | 1.095890432 | 0.864531704 |
| SPCC13B11.04c | 0.774648832 | 0.86458434 |
| SPBC2D10.05 | 0.895948896 | 0.86496161 |
| SPBC83.01 | 0.906676978 | 0.865898836 |
| SPBP4H10.12 | 0.913475 | 0.866460465 |
| SPAC17C9.10 | 0.926534724 | 0.866682802 |
| SPBC18A7.01 | 0.892121027 | 0.867386756 |
| SPAC22F3.02 | 0.923288275 | 0.867396627 |
| SPAC57A7.05 | 0.846886795 | 0.867541719 |
| SPAC3F10.11c | 0.981720742 | 0.867555723 |
| SPAC57A10.03 | 0.883320759 | 0.867607849 |
| SPBC1861.06c | 1.021295483 | 0.867769384 |
| SPCC4B3.06c | 0.860555699 | 0.86878127 |
| SPCC364.01 | 0.971484097 | 0.868935933 |
| SPAC17C9.08 | 0.879275473 | 0.869122934 |
| SPAC6G9.15c | 0.912818703 | 0.869378124 |
| SPBC17A3.10 | 1.104454381 | 0.870052423 |
| SPCC1840.04 | 0.760822519 | 0.87051549 |
| SPAC12B10.13 | 0.954156158 | 0.870676949 |
| SPCC63.14 | 1.148987247 | 0.870709678 |
| SPBC17F3.01c | 0.847055173 | 0.870853766 |
| SPAC26F1.01 | 0.897363416 | 0.870918379 |

| | | |
|---|---|---|
| SPCC1450.06c | 0.873259392 | 0.871197773 |
| SPCC1840.02c | 0.887164724 | 0.871747905 |
| SPAC1556.03 | 0.346779191 | 0.872141399 |
| SPCC4B3.03c | 0.986470227 | 0.872507559 |
| SPAC589.10c | 0.830328882 | 0.872510461 |
| SPAC12B10.03 | 0.852464895 | 0.872598326 |
| SPCC1795.06 | 0.870044843 | 0.872837924 |
| SPCC1442.11c | 0.566581423 | 0.873946804 |
| SPCC1020.01c | 0.877678269 | 0.874105787 |
| SPBC106.11c | 0.698769007 | 0.874144328 |
| SPAC14C4.05c | 1.010218254 | 0.874670274 |
| SPBC359.03c | 0.859384808 | 0.875182323 |
| SPCC1450.08c | 0.912584249 | 0.875270589 |
| SPBC216.03 | 0.550897171 | 0.875485249 |
| SPCC126.10 | 0.960463365 | 0.876202302 |
| SPBC2G2.09c | 0.925632815 | 0.876526633 |
| SPAC1399.04c | 0.93948654 | 0.876605393 |
| SPCC1442.05c | 0.821877455 | 0.876669285 |
| SPAC14C4.08 | 0.917955509 | 0.876742608 |
| SPCC1223.12c | 1.812737702 | 0.876820375 |
| SPBC660.07 | 0.649594009 | 0.87688308 |
| SPBC1773.05c | 0.829839973 | 0.877209273 |
| SPCC18.17c | 1.828024975 | 0.877228532 |
| SPBC1773.02c | 0.831633303 | 0.877244432 |
| SPAC3F10.16c | 0.994167299 | 0.877245298 |
| SPAC3H1.05 | 0.940432213 | 0.877393665 |
| SPAC1782.05 | 0.964413849 | 0.877882792 |
| SPCC965.13 | 0.770280927 | 0.878043708 |
| SPBC23E6.05 | 0.89200574 | 0.87810216 |
| SPAC2G11.13 | 0.843220038 | 0.878185653 |
| SPAPB24D3.04c | 0.943695444 | 0.878473367 |
| SPAC17G6.03 | 0.953337725 | 0.879157468 |
| SPAC13G7.09c | 0.867516826 | 0.879184378 |
| SPAC688.14 | 0.785047489 | 0.879227978 |
| SPCC1529.01 | 1.026662315 | 0.879454942 |
| SPAC343.20 | 0.891045854 | 0.879688967 |
| SPBP23A10.12 | 0.919491526 | 0.879756356 |
| SPAC4F10.06 | 0.184214966 | 0.879823834 |
| SPAC17H9.11 | 0.875753358 | 0.879874062 |
| SPBP35G2.04c | 0.941306254 | 0.879938889 |
| SPAC21E11.04 | 0.766842105 | 0.87997291 |
| SPBC28E12.04 | 0.731591131 | 0.880117048 |
| SPCC1620.04c | 0.990844668 | 0.880246586 |
| SPBC1685.04 | 0.898944892 | 0.88086187 |
| SPAC23H4.16c | 0.855904877 | 0.8808948 |
| SPCC188.12 | 0.847069019 | 0.880903531 |
| SPBC428.03c | 0.851828517 | 0.880912799 |
| SPAC20G4.05c | 1.044962447 | 0.88129567 |

| | | |
|---|---|---|
| SPAC227.10 | 0.684690393 | 0.881305009 |
| SPBC660.05 | 0.826666667 | 0.881685127 |
| SPBPB2B2.07c | 0.880218254 | 0.88193156 |
| SPAC1D4.13 | 0.75118431 | 0.882468876 |
| SPAC1A6.08c | 0.886544422 | 0.882613929 |
| SPCC18.01c | 0.865054248 | 0.883009716 |
| SPBC1604.04 | 0.776031792 | 0.883024866 |
| SPAC30D11.10 | 3.206452236 | 0.883890252 |
| SPAC25H1.02 | 0.908543322 | 0.883949176 |
| SPBC16E9.16c | 0.773872006 | 0.884116077 |
| SPAC27D7.08c | 0.777070752 | 0.884858607 |
| SPCC622.16c | 0.906047816 | 0.884982283 |
| SPAC11D3.13 | 0.883922126 | 0.885599726 |
| SPAC3A11.03 | 1.054175859 | 0.885705563 |
| SPCC16C4.11 | 0.967588723 | 0.886051395 |
| SPAC3C7.14c | 0.91426327 | 0.886071778 |
| SPAC3A12.08 | 0.958074148 | 0.886399456 |
| SPAC3G6.05 | 1.098479376 | 0.886941019 |
| SPBC16A3.14 | 0.858340611 | 0.886951965 |
| SPAC140.04 | 0.847448206 | 0.88700101 |
| SPAC222.08c | 1.253979925 | 0.887047957 |
| SPAC25H1.04 | 1.039526343 | 0.887422261 |
| SPBP35G2.11c | 0.82153723 | 0.887697658 |
| SPAC19D5.07 | 0.811533736 | 0.887705346 |
| SPAC19B12.12c | 0.842260573 | 0.887885412 |
| SPAC1687.06c | 0.96512141 | 0.888350036 |
| SPAC222.04c | 1.472236034 | 0.888686922 |
| SPCC1902.01 | 0.984419258 | 0.888741012 |
| SPBC16C6.03c | 1.10071336 | 0.888971261 |
| SPBP4H10.13 | 0.991062651 | 0.889400787 |
| SPCC1183.06 | 0.880893231 | 0.889918908 |
| SPAC1782.08c | 0.99335449 | 0.889938847 |
| SPAC17A5.05c | 0.964725312 | 0.890048105 |
| SPCP1E11.10 | 1.153585015 | 0.89035165 |
| SPBC11C11.01 | 0.83720079 | 0.890688695 |
| SPAC9E9.08 | 1.056414496 | 0.890979838 |
| SPACUNK4.19 | 0.773007473 | 0.891098427 |
| SPAC11E3.08c | 0.742171848 | 0.891181872 |
| SPAC20G4.08 | 0.951633672 | 0.891261172 |
| SPAC11H11.03c | 0.716577909 | 0.891806334 |
| SPAC19B12.07c | 0.999116955 | 0.891900574 |
| SPAC6C3.05 | 0.896439929 | 0.8922184 |
| SPAC22G7.06c | 1.013970319 | 0.892232188 |
| SPAC19A8.03 | 0.847600503 | 0.892358842 |
| SPBC14C8.03 | 0.85668435 | 0.892378754 |
| SPAC19A8.11c | 0.810538908 | 0.892403247 |
| SPAC23H3.14 | 0.886523684 | 0.892445566 |
| SPBC409.03 | 0.861047159 | 0.892487467 |

| | | |
|---|---|---|
| SPAC521.03 | 1.12691275 | 0.89261232 |
| SPAC1834.07 | 0.999594769 | 0.892626251 |
| SPBC1347.13c | 0.989897959 | 0.892684866 |
| SPBC887.11 | 0.884049981 | 0.892964771 |
| SPBC18H10.15 | 1.112190274 | 0.893055893 |
| SPCC4B3.11c | 0.747319395 | 0.893068185 |
| SPCC24B10.16c | 0.794810296 | 0.893207832 |
| SPAC24B11.13 | 1.030685104 | 0.893275676 |
| SPBC32F12.09 | 0.713531484 | 0.893684211 |
| SPBC1271.03c | 0.920537943 | 0.89373105 |
| SPAC3G9.04 | 0.866813426 | 0.894002273 |
| SPAPB1E7.02c | 0.931734193 | 0.8941699 |
| SPBP8B7.26 | 0.882657686 | 0.895159438 |
| SPBC1734.13 | 0.781777071 | 0.895309999 |
| SPAC1B3.16c | 0.943098421 | 0.895392625 |
| SPBC17G9.08c | 0.947100692 | 0.895866101 |
| SPBC4B4.07c | 0.962835123 | 0.896319747 |
| SPAC19A8.02 | 0.109352396 | 0.896410444 |
| SPAC29A4.18 | 1.478766431 | 0.896434783 |
| SPBC1347.09 | 0.726907379 | 0.896473441 |
| SPAC2E12.03c | 0.784111494 | 0.897056442 |
| SPAC3H8.05c | 0.932415585 | 0.897263641 |
| SPBC23G7.06c | 0.789797798 | 0.897317463 |
| SPCC4G3.10c | 0.806199631 | 0.897502848 |
| SPCC70.09c | 0.846675766 | 0.897710368 |
| SPAC20H4.05c | 0.882257415 | 0.897747921 |
| SPAC328.07c | 1.238232071 | 0.897989318 |
| SPBC1539.07c | 0.863490909 | 0.898295455 |
| SPAC227.07c | 0.832174145 | 0.898727171 |
| SPAC1527.03 | 0.866010243 | 0.89900111 |
| SPAPB1A10.09 | 1.024690083 | 0.899006595 |
| SPBC146.02 | 0.965142703 | 0.899026408 |
| SPAC7D4.02c | 0.855473031 | 0.899053908 |
| SPBC365.01 | 0.852259379 | 0.899072248 |
| SPAC5D6.09c | 1.15706619 | 0.899076923 |
| SPBC13G1.04c | 0.908147471 | 0.899240241 |
| SPAC5H10.02c | 0.8393 | 0.899367965 |
| SPAC1039.06 | 1.047412594 | 0.899449228 |
| SPBC902.05c | 0.907444313 | 0.89953072 |
| SPAPYUG7.02c | 1.001609729 | 0.899563614 |
| SPAPB24D3.03 | 0.862087736 | 0.899603971 |
| SPAC26H5.10c | 0.816973775 | 0.899622127 |
| SPCC285.11 | 0.702571226 | 0.899759139 |
| SPBC649.02 | 1.339679277 | 0.899838096 |
| SPAC4F10.15c | 1.083236388 | 0.899890351 |
| SPCC191.03c | 0.796249441 | 0.899922001 |
| SPAPB24D3.01 | 0.921087036 | 0.899924602 |
| SPAC10F6.06 | 0.860529685 | 0.900035868 |

| | | |
|---|---|---|
| SPBPB2B2.01 | 0.994831016 | 0.900095836 |
| SPAC227.04 | 0.816552256 | 0.900406627 |
| SPBC8D2.02c | 0.790645102 | 0.900474534 |
| SPAPB8E5.08 | 0.906326449 | 0.900586072 |
| SPAC10F6.04 | 0.686831993 | 0.900746494 |
| SPBC8D2.19 | 0.949209829 | 0.90092663 |
| SPAC26A3.04 | 0.728590786 | 0.901220238 |
| SPAC105.01c | 0.918477948 | 0.901947228 |
| SPBC651.02 | 0.842430903 | 0.902097963 |
| SPCC11E10.09c | 1.21151406 | 0.902117597 |
| SPBC18E5.11c | 0.848072492 | 0.902336785 |
| SPAC1142.07c | 0.854308446 | 0.902336842 |
| SPAC11H11.02c | 0.970797684 | 0.902668415 |
| SPBC543.08 | 0.978559482 | 0.902672142 |
| SPBC19G7.18c | 0.662258393 | 0.902721262 |
| SPAC1805.10 | 0.884239733 | 0.903163152 |
| SPAC23C11.01 | 0.869107143 | 0.903754783 |
| SPAC12G12.09 | 0.846375394 | 0.903909704 |
| SPCC1223.13 | 1.030781562 | 0.904155955 |
| SPBC19F8.02 | 0.831574231 | 0.904237092 |
| SPBC23G7.08c | 0.792778043 | 0.904389 |
| SPAC3H1.09c | 1.174812428 | 0.904477201 |
| SPAC26H5.02c | 0.39206033 | 0.904632632 |
| SPBC725.01 | 0.828926124 | 0.904632999 |
| SPAC29A4.05 | 0.862171624 | 0.904915591 |
| SPAC25A8.02 | 0.573483413 | 0.90599056 |
| SPCC965.08c | 0.919434622 | 0.906026201 |
| SPCC330.03c | 0.934058303 | 0.90611831 |
| SPCC970.05 | 2.56388962 | 0.906350479 |
| SPAC12B10.10 | 0.974797416 | 0.906452569 |
| SPCC23B6.03c | 0.946039034 | 0.906484036 |
| SPAC17G6.04c | 0.925694444 | 0.906521267 |
| SPCC1682.13 | 0.93213638 | 0.906556039 |
| SPCC1919.11 | 0.932440322 | 0.906892894 |
| SPAC57A7.08 | 1.279540169 | 0.907106384 |
| SPAC1250.02 | 0.834575361 | 0.90728852 |
| SPAC343.11c | 0.835598384 | 0.907477094 |
| SPBC2A9.13 | 0.881045141 | 0.907841495 |
| SPBCPT2R1.01c | 0.989194435 | 0.907939178 |
| SPAC186.06 | 0.757337662 | 0.907975862 |
| SPBC4F6.16c | 0.77015625 | 0.908203125 |
| SPCC794.06 | 0.767983969 | 0.908218021 |
| SPAC19B12.08 | 0.96189241 | 0.908406144 |
| SPBC336.05c | 0.970415473 | 0.908723585 |
| SPBPB10D8.05c | 1.179069767 | 0.90883146 |
| SPCC191.06 | 1.020375598 | 0.908878947 |
| SPBP8B7.02 | 0.868284383 | 0.909493785 |
| SPBC354.09c | 1.016355052 | 0.909942472 |

| | | |
|---|---|---|
| SPCC1223.15c | 0.752443185 | 0.91025237 |
| SPBC725.06c | 0.974137391 | 0.910309537 |
| SPAC6B12.06c | 0.916129299 | 0.910362756 |
| SPBC713.06 | 0.840377761 | 0.910391534 |
| SPCC594.02c | 1.01116279 | 0.9104 |
| SPBC2D10.15c | 0.973698352 | 0.910595602 |
| SPCC417.11c | 0.7629054 | 0.91069384 |
| SPCC330.11 | 0.924802674 | 0.910857994 |
| SPAC977.16c | 0.975067541 | 0.911405254 |
| SPAC26H5.05 | 0.989546969 | 0.911462569 |
| SPAC31G5.10 | 0.552560694 | 0.911759486 |
| SPAC1486.10 | 0.802416813 | 0.912084063 |
| SPAC869.03c | 0.988400245 | 0.912307726 |
| SPACUNK4.07c | 0.935306864 | 0.912310229 |
| SPBC725.07 | 1.166471239 | 0.912337148 |
| SPAC30C2.07 | 1.105978315 | 0.912438862 |
| SPAC30D11.02c | 1.110167789 | 0.912549234 |
| SPCC1906.02c | 0.986818906 | 0.912967015 |
| SPAC3F10.15c | 0.891698713 | 0.913509372 |
| SPAC607.06c | 1.027296567 | 0.913745186 |
| SPAC3A11.05c | 1.038907584 | 0.914291143 |
| SPAC26A3.01 | 0.814074528 | 0.914449784 |
| SPBC13G1.14c | 0.913265306 | 0.914540816 |
| SPBC1683.10c | 0.950363669 | 0.914748993 |
| SPBC685.03 | 1.256340016 | 0.914769655 |
| SPBC577.08c | 0.7956815 | 0.914865742 |
| SPBC1711.08 | 0.790824813 | 0.915490422 |
| SPCC31H12.06 | 0.787778726 | 0.916028648 |
| SPBC839.02 | 0.951774634 | 0.916092703 |
| SPBC337.09 | 1.003278154 | 0.91625455 |
| SPBC3F6.01c | 1.10653807 | 0.916989839 |
| SPAC3F10.09 | 0.864552196 | 0.917019421 |
| SPAC1782.01 | 0.928226527 | 0.917061012 |
| SPAC1687.05 | 1.026699189 | 0.917111142 |
| SPCC970.06 | 0.97452639 | 0.917118495 |
| SPCC830.08c | 0.943427353 | 0.917413842 |
| SPCC594.04c | 1.405575415 | 0.91776505 |
| SPAC1952.07 | 0.975831504 | 0.917766377 |
| SPBC3E7.10 | 0.707122454 | 0.917837838 |
| SPAC139.01c | 0.993791019 | 0.917881145 |
| SPAC19G12.16c | 0.904310665 | 0.917990524 |
| SPBC36.11 | 0.908177439 | 0.918080147 |
| SPBPB21E7.05 | 0.996282542 | 0.918130947 |
| SPBC1105.11c | 0.863632861 | 0.918321962 |
| SPACUNK12.02c | 0.841420468 | 0.918374602 |
| SPBC23E6.01c | 0.409812238 | 0.918577869 |
| SPCC417.07c | 0.802831788 | 0.919436396 |
| SPAC30D11.06c | 0.948893224 | 0.919634623 |

| | | |
|---|---|---|
| SPAC2F3.05c | 0.960568205 | 0.919681611 |
| SPBC947.10 | 1.557821662 | 0.919733945 |
| SPAC4C5.04 | 1.007305917 | 0.919888699 |
| SPAC26A3.06 | 0.951420094 | 0.920291967 |
| SPAC22F8.02c | 0.393254039 | 0.920519084 |
| SPAC922.07c | 0.895703236 | 0.920566157 |
| SPAPB24D3.08c | 1.00577897 | 0.920869331 |
| SPCP25A2.02c | 0.752376758 | 0.921067754 |
| SPAC4F10.05c | 0.903472859 | 0.921261445 |
| SPBC215.04 | 1.093408604 | 0.921324372 |
| SPAC19D5.06c | 1.080355798 | 0.921515608 |
| SPCC74.05 | 1.113487796 | 0.921521066 |
| SPAC4H3.01 | 1.104030885 | 0.92205242 |
| SPBC428.08c | 0.732361618 | 0.92209036 |
| SPAC806.04c | 1.216929988 | 0.922197109 |
| SPBC428.12c | 1.091266581 | 0.922287444 |
| SPAC27D7.03c | 1.027487923 | 0.922388668 |
| SPAC186.03 | 0.96866221 | 0.92286347 |
| SPBC14C8.05c | 0.785676325 | 0.923258447 |
| SPBC1921.05 | 0.737658959 | 0.923661012 |
| SPCC790.02 | 0.849397159 | 0.923693526 |
| SPCPJ732.01 | 1.110979015 | 0.923729736 |
| SPBC1734.15 | 0.879084305 | 0.92405214 |
| SPBC25H2.16c | 0.851737349 | 0.924768657 |
| SPCC1739.15 | 0.85060241 | 0.925075301 |
| SPBC1711.14 | 1.069345817 | 0.92520058 |
| SPCC1259.13 | 0.962973531 | 0.92522795 |
| SPAC1F3.05 | 0.821165757 | 0.925357533 |
| SPAC22A12.10 | 0.975938164 | 0.925625423 |
| SPAC7D4.03c | 0.848198326 | 0.92619761 |
| SPAPB17E12.03 | 0.946432719 | 0.926309782 |
| SPAC26F1.14c | 1.273116531 | 0.926363316 |
| SPCC794.15 | 0.961410777 | 0.926439961 |
| SPAC2G11.04 | 0.8921432 | 0.926509437 |
| SPBC660.12c | 0.894782911 | 0.926535022 |
| SPBC1A4.04 | 0.968696726 | 0.927090704 |
| SPAC3A12.03c | 0.999169997 | 0.927215389 |
| SPBP35G2.12 | 0.931098743 | 0.927288271 |
| SPAC17A2.11 | 0.915938713 | 0.927302411 |
| SPBC1539.06 | 1.131064926 | 0.927381495 |
| SPAC13G6.10c | 0.329163671 | 0.92784726 |
| SPAC6G10.02c | 1.003968043 | 0.927969754 |
| SPAC607.09c | 0.7134417 | 0.92803696 |
| SPBPB21E7.04c | 1.174166308 | 0.928101139 |
| SPAC8E11.04c | 0.902536541 | 0.928153308 |
| SPAC22G7.03 | 0.907459114 | 0.928356576 |
| SPCC645.14c | 0.769897186 | 0.92862393 |
| SPAC12G12.16c | 0.892146315 | 0.92862426 |

| | | |
|---|---|---|
| SPCC364.07 | 0.927835096 | 0.928820059 |
| SPBC1703.11 | 0.951094594 | 0.928863059 |
| SPAC9.07c | 0.808651649 | 0.928899363 |
| SPAC17C9.07 | 0.873101534 | 0.929919137 |
| SPAPB1A10.07c | 0.987620818 | 0.930216078 |
| SPAC26F1.07 | 0.945723043 | 0.930228707 |
| SPAC5D6.04 | 0.802543023 | 0.930366 |
| SPAC1952.15c | 0.881015396 | 0.930397872 |
| SPAC17H9.06c | 0.894707425 | 0.930410232 |
| SPAC10F6.12c | 0.917968234 | 0.930414333 |
| SPCC663.14c | 1.069470631 | 0.930465059 |
| SPBC146.11c | 0.962176311 | 0.930667825 |
| SPCC584.13 | 0.789628834 | 0.931237823 |
| SPCC306.07c | 0.878427454 | 0.931622972 |
| SPCC645.12c | 0.897701704 | 0.931747671 |
| SPBC31F10.14c | 0.900277542 | 0.93239258 |
| SPBC1105.01 | 0.920601442 | 0.932644969 |
| SPAC18G6.04c | 1.228807202 | 0.932739664 |
| SPBC18H10.05 | 0.893626182 | 0.932875785 |
| SPBC25B2.08 | 0.925475767 | 0.933217761 |
| SPBC3H7.14 | 0.972749793 | 0.933551817 |
| SPAC31G5.04 | 0.939497357 | 0.933664254 |
| SPCC24B10.18 | 0.864560956 | 0.934432525 |
| SPAC13G7.11 | 1.056121353 | 0.934608959 |
| SPAC869.09 | 0.922125814 | 0.935242012 |
| SPBC530.03c | 0.98915581 | 0.935249375 |
| SPBP8B7.31 | 0.918515599 | 0.935305386 |
| SPAC9E9.14 | 0.864645728 | 0.93543942 |
| SPAC27E2.09 | 0.860937283 | 0.935726571 |
| SPCC553.03 | 0.97460528 | 0.936450909 |
| SPAC22H12.01c | 0.817648686 | 0.937002257 |
| SPAC688.10 | 0.727257167 | 0.937006874 |
| SPAC513.07 | 1.050298028 | 0.937143748 |
| SPAC227.13c | 1.039536878 | 0.937178387 |
| SPBC3B8.07c | 1.695254273 | 0.937409639 |
| SPBC21B10.10 | 1.120520334 | 0.937589331 |
| SPBC1683.13c | 0.946382979 | 0.937593985 |
| SPBC646.06c | 0.892385496 | 0.937633855 |
| SPAC11D3.02c | 0.943429603 | 0.937973826 |
| SPAC4H3.02c | 0.831288433 | 0.938134117 |
| SPAC750.08c | 1.00500804 | 0.938585179 |
| SPCC1739.13 | 0.958498058 | 0.938781405 |
| SPAC922.05c | 0.823074009 | 0.93904886 |
| SPCC338.08 | 2.544712325 | 0.939498471 |
| SPBC530.06c | 1.113053135 | 0.939599047 |
| SPAC1783.08c | 0.841995731 | 0.939801228 |
| SPCC2H8.05c | 0.976015273 | 0.940220149 |
| SPACUNK4.13c | 0.848097637 | 0.94069583 |

| | | |
|---|---|---|
| SPAC1093.01 | 0.793383796 | 0.940772762 |
| SPBC947.06c | 0.882651939 | 0.940839126 |
| SPCC613.12c | 1.086390575 | 0.941134499 |
| SPCC126.11c | 0.829494025 | 0.941240379 |
| SPBC3B8.05 | 1.276530886 | 0.941803576 |
| SPBC6B1.10 | 1.00630734 | 0.94183658 |
| SPAC328.05 | 1.08571237 | 0.942063591 |
| SPAC11D3.16c | 0.877510189 | 0.942189237 |
| SPAC23H4.10c | 0.956345238 | 0.942599464 |
| SPAC1F5.09c | 0.818688382 | 0.942646834 |
| SPAC26A3.17c | 0.95556701 | 0.942783505 |
| SPAC1687.16c | 1.003518489 | 0.9428126 |
| SPBC1D7.01 | 0.978233603 | 0.942926474 |
| SPBC428.07 | 0.871945047 | 0.94353308 |
| SPBC4B4.10c | 0.879235311 | 0.943563664 |
| SPCC663.13c | 1.144896626 | 0.943566853 |
| SPBC1347.11 | 0.938265023 | 0.943568841 |
| SPAPYUK71.03c | 0.82791767 | 0.943607745 |
| SPBC83.04 | 0.983168126 | 0.943666058 |
| SPAC1486.04c | 1.003640722 | 0.943746672 |
| SPBC28F2.05c | 1.218878645 | 0.943833897 |
| SPAC1002.05c | 1.001604739 | 0.94470151 |
| SPBP8B7.05c | 0.980390711 | 0.944825839 |
| SPBC776.04 | 0.826160656 | 0.944894282 |
| SPAC1F8.05 | 0.880313049 | 0.945027098 |
| SPBC1604.16c | 0.930490111 | 0.94503276 |
| SPAC1687.07 | 0.928745319 | 0.945237748 |
| SPCC191.05c | 0.904877717 | 0.945503822 |
| SPBC31E1.02c | 0.960866297 | 0.946135101 |
| SPAC1805.02c | 0.897172232 | 0.946215883 |
| SPBPB2B2.02 | 0.878489087 | 0.946245433 |
| SPBC651.11c | 1.846175784 | 0.946396163 |
| SPAC23G3.12c | 0.939802254 | 0.946557954 |
| SPBC106.12c | 0.855324413 | 0.946661499 |
| SPAC11E3.05 | 0.816085694 | 0.947644336 |
| SPCC663.08c | 0.510326629 | 0.947755738 |
| SPAC1071.03c | 0.600015339 | 0.94777471 |
| SPAP27G11.08c | 0.920203572 | 0.947964335 |
| SPBC428.14 | 0.713251973 | 0.948264371 |
| SPCC24B10.04 | 0.76540072 | 0.948464548 |
| SPBPJ4664.06 | 0.718906469 | 0.948846548 |
| SPBC17D1.05 | 0.851273387 | 0.948928079 |
| SPAC3G9.07c | 1.177230965 | 0.948992877 |
| SPCC1235.01 | 0.895454546 | 0.94921875 |
| SPBC106.10 | 0.910831941 | 0.94934435 |
| SPBC3E7.16c | 1.026954714 | 0.949740825 |
| SPBC691.03c | 0.759060308 | 0.949875452 |
| SPAC9E9.15 | 0.863075944 | 0.950065078 |

| | | |
|---|---|---|
| SPBC3H7.15 | 0.664108038 | 0.950276374 |
| SPBC1685.05 | 1.033916924 | 0.950291291 |
| SPAC144.04c | 1.063722045 | 0.950589928 |
| SPAC1F7.12 | 0.882114853 | 0.950904224 |
| SPAPB24D3.02c | 1.204774606 | 0.951230332 |
| SPAC1039.07c | 0.829898939 | 0.951260504 |
| SPAC31A2.16 | 0.968863278 | 0.951281133 |
| SPBC21B10.08c | 1.036489164 | 0.951406857 |
| SPAPB24D3.09c | 0.936989926 | 0.951455445 |
| SPBC13A2.02 | 0.811372065 | 0.951505453 |
| SPBC29A3.02c | 1.522269175 | 0.951542671 |
| SPCC550.08 | 1.091243342 | 0.951687151 |
| SPAC30D11.07 | 1.191030094 | 0.951752684 |
| SPBC1198.01 | 0.843623742 | 0.951832836 |
| SPBC31F10.02 | 0.936278143 | 0.952330898 |
| SPBC1D7.03 | 0.933561214 | 0.953395116 |
| SPBC18H10.13 | 1.001126155 | 0.953464764 |
| SPBC215.11c | 1.107716702 | 0.953590909 |
| SPBC23G7.14 | 0.997607296 | 0.953848726 |
| SPBC31F10.07 | 1.160492345 | 0.954352257 |
| SPAC31G5.09c | 0.928635891 | 0.954368159 |
| SPCC970.01 | 0.960930085 | 0.954369338 |
| SPCC24B10.13 | 0.913351531 | 0.954562152 |
| SPAC1952.06c | 1.121033792 | 0.954575024 |
| SPBPB8B6.05c | 1.001885949 | 0.954641726 |
| SPAC589.11 | 1.005417577 | 0.955042145 |
| SPCC1494.03 | 1.051089291 | 0.955076327 |
| SPBC215.06c | 0.956358238 | 0.955230685 |
| SPBC887.17 | 1.476621787 | 0.955705263 |
| SPAC688.12c | 0.960338863 | 0.955965026 |
| SPAC1F8.02c | 0.904334633 | 0.956152471 |
| SPAC11H11.04 | 1.058127111 | 0.956382009 |
| SPAC17A2.14 | 1.086566369 | 0.956732981 |
| SPBC56F2.14 | 0.998426427 | 0.956768946 |
| SPACUNK4.08 | 1.053741451 | 0.956952998 |
| SPAC683.02c | 1.091832325 | 0.957027874 |
| SPAC2E1P5.01c | 0.974197898 | 0.957138212 |
| SPAC2G11.09 | 0.941443805 | 0.957221661 |
| SPBC4.05 | 1.033498191 | 0.957332879 |
| SPAC23C11.13c | 0.8502 | 0.957416179 |
| SPCC16A11.03c | 1.078480506 | 0.95763382 |
| SPAC1783.06c | 1.181120083 | 0.958561715 |
| SPAC14C4.16 | 0.824639591 | 0.95890088 |
| SPCC1235.06 | 1.331754137 | 0.958920728 |
| SPBC1683.02 | 1.008467759 | 0.959044979 |
| SPBC557.02c | 0.843513627 | 0.959134908 |
| SPAC11D3.04c | 1.002621232 | 0.959231806 |
| SPAC144.14 | 0.970901109 | 0.959312421 |

| | | |
|---|---|---|
| SPBC776.05 | 1.143968295 | 0.959349122 |
| SPCC1753.02c | 0.30570167 | 0.9597458 |
| SPAC1F8.04c | 1.09947705 | 0.959827629 |
| SPCC4G3.02 | 0.896624539 | 0.959918228 |
| SPBC646.02 | 1.00635407 | 0.960589405 |
| SPBC36.10 | 0.916158752 | 0.960854447 |
| SPBC1604.20c | 1.717499261 | 0.961083215 |
| SPCC1442.03 | 0.203412073 | 0.961122047 |
| SPAC1687.15 | 0.15330446 | 0.961200393 |
| SPAC22A12.02c | 0.962102009 | 0.961263524 |
| SPAC1296.03c | 0.891646627 | 0.961379602 |
| SPAC6B12.15 | 0.820418766 | 0.961431667 |
| SPAC12B10.16c | 0.923326713 | 0.961807642 |
| SPBPB7E8.02 | 1.117210357 | 0.961829983 |
| SPAC1B3.08 | 0.855788661 | 0.961987866 |
| SPBC947.15c | 1.06977907 | 0.962325 |
| SPAC3A12.06c | 0.974588941 | 0.962488588 |
| SPCP1E11.04c | 1.283632055 | 0.962828025 |
| SPAC13G6.12c | 1.056483367 | 0.962830957 |
| SPCC569.01c | 0.872870264 | 0.963393551 |
| SPBC9B6.09c | 0.861148078 | 0.963405941 |
| SPAC688.04c | 1.057345412 | 0.963789878 |
| SPCC4G3.15c | 1.217804241 | 0.96404617 |
| SPAC1805.12c | 0.998348616 | 0.964308855 |
| SPBP35G2.03c | 0.913217569 | 0.964568773 |
| SPBC1683.01 | 0.88105306 | 0.96464956 |
| SPAC27E2.02 | 1.0815079 | 0.964712929 |
| SPAC683.03 | 1.00892506 | 0.965449608 |
| SPCC306.02c | 1.004396866 | 0.965733974 |
| SPAC869.06c | 1.184515141 | 0.9658102 |
| SPAC821.05 | 1.033542758 | 0.965828639 |
| SPBC25B2.01 | 1.185034124 | 0.966019005 |
| SPAC24C9.14 | 0.99970467 | 0.967079607 |
| SPAC19G12.11 | 1.180048176 | 0.967344463 |
| SPAC29B12.12 | 1.117812242 | 0.967802807 |
| SPAC1805.16c | 0.966682586 | 0.967901584 |
| SPAC32A11.01 | 0.935510242 | 0.96814445 |
| SPAC11E3.14 | 0.960691849 | 0.968151031 |
| SPBC530.02 | 0.853678382 | 0.968224263 |
| SPAC15F9.01c | 0.964433921 | 0.968320715 |
| SPAC3G6.11 | 0.860014357 | 0.968349499 |
| SPAC6F6.13c | 0.709193508 | 0.968649444 |
| SPAC13G6.15c | 1.199785497 | 0.968819254 |
| SPBC16A3.12c | 0.957711593 | 0.96895942 |
| SPBC776.06c | 0.912407045 | 0.969129159 |
| SPBC1348.07 | 0.846117309 | 0.969595457 |
| SPBC16G5.09 | 0.922552886 | 0.970139387 |
| SPAC607.07c | 0.896759259 | 0.970258137 |

| | | |
|---|---|---|
| SPAC2C4.17c | 0.999958581 | 0.970264954 |
| SPCC11E10.03 | 1.01613325 | 0.970323118 |
| SPAC2C4.05 | 0.909149015 | 0.970464635 |
| SPCC777.12c | 0.150086015 | 0.971046575 |
| SPAC1A6.07 | 1.132347448 | 0.971641751 |
| SPCC18B5.09c | 0.947905239 | 0.971679574 |
| SPBC428.17c | 0.61956052 | 0.971753953 |
| SPAC6C3.02c | 1.054618329 | 0.972107722 |
| SPBC4B4.11 | 0.883682251 | 0.972325226 |
| SPBC15D4.07c | 1.020691583 | 0.972367158 |
| SPCC63.03 | 0.912042662 | 0.97249696 |
| SPAC23H4.09 | 1.021631817 | 0.97250537 |
| SPCC18B5.10c | 0.824112288 | 0.973799617 |
| SPCC1906.04 | 0.904979408 | 0.973891199 |
| SPCC1259.03 | 1.095353096 | 0.974055151 |
| SPAC25B8.05 | 0.9812363 | 0.974695857 |
| SPAC1071.05 | 0.915190351 | 0.974926488 |
| SPCC645.11c | 1.397572553 | 0.974963653 |
| SPBC685.02 | 0.82793401 | 0.975029394 |
| SPAC4H3.07c | 1.199629124 | 0.975193144 |
| SPBC4C3.06 | 0.844575079 | 0.975515961 |
| SPCC569.06 | 0.956463415 | 0.975838415 |
| SPCC11E10.07c | 0.965911493 | 0.975949593 |
| SPCC16C4.03 | 0.925191509 | 0.976041858 |
| SPCC576.17c | 0.510825982 | 0.976524138 |
| SPCC11E10.05c | 0.987994581 | 0.976525796 |
| SPBC2G2.05 | 1.038376374 | 0.976545021 |
| SPBC11B10.06 | 0.891821681 | 0.976828095 |
| SPAC1834.10c | 1.413776761 | 0.976953368 |
| SPBC15D4.03 | 0.929779409 | 0.977051455 |
| SPAC12G12.12 | 1.128403498 | 0.977105077 |
| SPCC4G3.08 | 0.985566232 | 0.977324456 |
| SPAC750.06c | 0.994051747 | 0.977623306 |
| SPAC3A12.12 | 1.161070118 | 0.977624256 |
| SPAC3H1.07 | 0.881879724 | 0.977659375 |
| SPAC17A2.13c | 0.785961081 | 0.977712251 |
| SPBC11C11.10 | 1.037876226 | 0.977923148 |
| SPAC1F12.10c | 0.793971501 | 0.977952594 |
| SPAC19E9.01c | 0.646405898 | 0.978198427 |
| SPAC328.06 | 0.953636933 | 0.978300755 |
| SPCC1393.05 | 0.91005352 | 0.978592904 |
| SPAC607.10 | 0.832000263 | 0.97859712 |
| SPAC23E2.03c | 0.394293709 | 0.97863985 |
| SPBC25H2.10c | 0.989025362 | 0.978804351 |
| SPAC926.02 | 0.859790622 | 0.978935412 |
| SPAC3F10.13 | 0.567232508 | 0.979905246 |
| SPAC3F10.05c | 0.725736472 | 0.980134384 |
| SPAC25G10.09c | 0.975544975 | 0.980327058 |

| | | |
|---|---|---|
| SPAC4F8.10c | 1.167458973 | 0.980428975 |
| SPBC18E5.08 | 1.284989953 | 0.980497726 |
| SPAC17A5.11 | 0.987743404 | 0.980644614 |
| SPBC609.05 | 0.982595572 | 0.980887435 |
| SPCC737.07c | 0.883018868 | 0.981132075 |
| SPBC31E1.01c | 0.667803238 | 0.98135912 |
| SPAC4H3.04c | 0.82661971 | 0.981754569 |
| SPCC569.05c | 1.073562829 | 0.981976294 |
| SPAC27D7.06 | 0.962091773 | 0.982065997 |
| SPAC824.05 | 1.036640814 | 0.982279868 |
| SPAC824.04 | 1.027877664 | 0.982310498 |
| SPCC1020.05 | 0.832107634 | 0.982418848 |
| SPAC3A11.02 | 0.815789474 | 0.982828457 |
| SPAC1093.06c | 0.987692592 | 0.982976168 |
| SPBC83.11 | 0.937689925 | 0.982977204 |
| SPAC26A3.16 | 1.007613069 | 0.983052853 |
| SPAC17A5.02c | 1.397738783 | 0.983085118 |
| SPBC29A10.12 | 0.531573525 | 0.98319995 |
| SPBC1347.08c | 0.97608235 | 0.983707993 |
| SPBC577.03c | 1.020758614 | 0.984018212 |
| SPBC56F2.10c | 0.822331618 | 0.984156203 |
| SPAC7D4.08 | 0.810522115 | 0.984287025 |
| SPAPB17E12.12c | 0.923105098 | 0.984748122 |
| SPBC725.04 | 0.868088272 | 0.984905661 |
| SPAC29E6.01 | 0.963715111 | 0.986465351 |
| SPBC4F6.05c | 0.942655475 | 0.986860931 |
| SPAC4D7.10c | 0.563051172 | 0.986880045 |
| SPAC607.08c | 0.966457898 | 0.986951983 |
| SPBC21C3.11 | 0.995689594 | 0.987306396 |
| SPCC4G3.11 | 2.181963087 | 0.987442324 |
| SPAPB8E5.03 | 1.026640003 | 0.987458014 |
| SPBPJ4664.02 | 0.949757322 | 0.987539526 |
| SPAC23H3.15c | 0.902588066 | 0.987553918 |
| SPAC821.11 | 0.80276066 | 0.988063223 |
| SPBC1289.09 | 1.057689335 | 0.988214838 |
| SPBC15C4.05 | 0.949973844 | 0.988444509 |
| SPBC56F2.01 | 0.875481213 | 0.98850827 |
| SPBC8D2.01 | 0.954855581 | 0.988550786 |
| SPCC1259.11c | 0.899847798 | 0.989324252 |
| SPBC1706.01 | 1.176996643 | 0.989425676 |
| SPBC18H10.08c | 1.000818182 | 0.989836679 |
| SPAC3C7.09 | 0.80772573 | 0.989989324 |
| SPAC15A10.09c | 1.169382127 | 0.990055509 |
| SPCC188.09c | 0.892671201 | 0.990206976 |
| SPAC1D4.03c | 1.061106281 | 0.990794393 |
| SPBC106.02c | 0.910100937 | 0.991967975 |
| SPBC646.13 | 1.009046493 | 0.992369813 |
| SPBC21H7.07c | 1.042836539 | 0.992497086 |

| | | |
|---|---|---|
| SPAC24C9.16c | 1.279776786 | 0.99259431 |
| SPAC6F6.04c | 0.986706005 | 0.992624089 |
| SPAC16E8.12c | 1.56756374 | 0.993105967 |
| SPAC17A2.01 | 0.995661759 | 0.99312564 |
| SPBP26C9.03c | 0.935505097 | 0.993526586 |
| SPAC13G6.13 | 0.939205458 | 0.993629615 |
| SPAC20G8.10c | 1.177845217 | 0.994041582 |
| SPAC20G4.02c | 0.990701971 | 0.99410214 |
| SPBC337.03 | 0.963277449 | 0.994153004 |
| SPBC19G7.08c | 1.109145596 | 0.994200309 |
| SPBC2G5.01 | 0.971675468 | 0.994910089 |
| SPAC1039.05c | 0.867775886 | 0.995016612 |
| SPAC6G9.05 | 0.980131878 | 0.995304421 |
| SPBC2G2.10c | 1.327949283 | 0.995877423 |
| SPBC19C7.11 | 0.974306085 | 0.995906408 |
| SPBC336.10c | 0.879937556 | 0.99605923 |
| SPAPB2B4.04c | 1.056275714 | 0.996214697 |
| SPBC582.09 | 1.097161617 | 0.99624857 |
| SPCC1393.12 | 1.014765013 | 0.996750024 |
| SPAC1805.08 | 1.122389007 | 0.997552332 |
| SPBC3E7.09 | 0.865822111 | 0.997663636 |
| SPAC12G12.11c | 0.813350842 | 0.997778043 |
| SPBC14F5.03c | 0.929728314 | 0.997801684 |
| SPCC1322.09 | 0.981565129 | 0.99792923 |
| SPBC1271.05c | 0.916753268 | 0.99816379 |
| SPCC548.07c | 0.967956212 | 0.998298493 |
| SPCC18B5.03 | 1.09253486 | 0.998993917 |
| SPAC13F5.04c | 0.938995582 | 0.999403962 |
| SPBC725.12 | 0.919026549 | 0.999534896 |
| SPAC869.04 | 0.978408725 | 0.999860997 |
| SPAC4G9.09c | 3.189915808 | 0.999863989 |
| SPAC6F12.06 | 0.884683426 | 0.999869064 |
| SPAC17H9.08 | 1.002292022 | 0.999959566 |
| SPCC162.06c | 1.492843955 | 1.000168035 |
| SPAPB17E12.08 | 1.015133731 | 1.000242367 |
| SPAC31G5.14 | 1.060601918 | 1.000599007 |
| SPAC3C7.05c | 0.702594131 | 1.000943873 |
| SPBC1D7.04 | 1.125582155 | 1.001229456 |
| SPBC21H7.04 | 1.091555761 | 1.001316045 |
| SPAC1851.03 | 0.936006527 | 1.002037034 |
| SPAC1610.01 | 0.982613108 | 1.002481169 |
| SPBC17A3.06 | 1.130573341 | 1.00305905 |
| SPAC4D7.01c | 0.976727533 | 1.003088077 |
| SPAC23D3.10c | 0.92258723 | 1.003220281 |
| SPBC3B9.04 | 1.188114177 | 1.00345946 |
| SPAC13A11.01c | 0.962719361 | 1.003553362 |
| SPBC354.01 | 0.881908704 | 1.003681546 |
| SPAC9.12c | 1.037952301 | 1.003798352 |

| | | |
|---|---|---|
| SPBC215.08c | 3.608250322 | 1.003895466 |
| SPBC839.13c | 1.434559665 | 1.003983607 |
| SPCC1322.10 | 0.923337725 | 1.003987219 |
| SPCC16A11.16c | 0.977605681 | 1.004168456 |
| SPAC6C3.07 | 0.963173765 | 1.004253044 |
| SPCC1753.03c | 0.999161266 | 1.004376186 |
| SPBC418.01c | 0.461760283 | 1.004405814 |
| SPBC2G2.01c | 0.850533783 | 1.004562952 |
| SPBC146.10 | 0.900613659 | 1.004576681 |
| SPBC1289.15 | 1.072795275 | 1.004601378 |
| SPAC144.02 | 1.082670107 | 1.004951493 |
| SPAC23G3.07c | 0.961527067 | 1.00524357 |
| SPCC550.01c | 1.034773601 | 1.005721242 |
| SPCC1442.15c | 1.052432324 | 1.005777952 |
| SPAC22A12.16 | 0.919471004 | 1.005966974 |
| SPCC825.02 | 1.008775782 | 1.006041309 |
| SPCC553.07c | 0.908562563 | 1.006111356 |
| SPCC1739.06c | 0.502243695 | 1.006126761 |
| SPAC23D3.13c | 1.046546418 | 1.006345662 |
| SPBC83.17 | 0.847592087 | 1.006784976 |
| SPCC757.13 | 0.913109143 | 1.006785762 |
| SPAC922.03 | 0.971411535 | 1.00723601 |
| SPCC285.10c | 1.02176142 | 1.007366813 |
| SPCC594.07c | 0.874070067 | 1.007375447 |
| SPCP31B10.07 | 1.043468828 | 1.007652441 |
| SPAC26H5.04 | 0.995402581 | 1.007872777 |
| SPAC977.15 | 0.969814726 | 1.007989653 |
| SPAC8C9.10c | 0.890648858 | 1.008261552 |
| SPAC23D3.01 | 1.103034023 | 1.008976859 |
| SPBC660.14 | 1.028030605 | 1.00910984 |
| SPCC4B3.07 | 1.060339384 | 1.009110899 |
| SPBC8E4.02c | 1.267580073 | 1.009229974 |
| SPBC1347.01c | 1.081155128 | 1.00927897 |
| SPCC4B3.13 | 0.676086535 | 1.009727122 |
| SPAC17D4.04 | 0.986589669 | 1.009775024 |
| SPBP8B7.24c | 1.037612146 | 1.009866718 |
| SPBC30B4.01c | 1.06691978 | 1.010070917 |
| SPAC630.13c | 0.719069736 | 1.010118767 |
| SPAPYUG7.06 | 0.907362876 | 1.010255079 |
| SPBC36B7.03 | 0.900587581 | 1.010647849 |
| SPBC56F2.03 | 1.113691246 | 1.010782679 |
| SPAC1399.05c | 0.952082522 | 1.011361425 |
| SPAC22A12.11 | 1.146137583 | 1.011524475 |
| SPAC22F8.03c | 0.912015104 | 1.01165036 |
| SPBC2D10.09 | 0.917918675 | 1.01184695 |
| SPAC1420.01c | 0.996491918 | 1.012570706 |
| SPBC25H2.15 | 0.873903925 | 1.012767192 |
| SPAC56E4.03 | 0.996741048 | 1.013497158 |

| | | |
|---|---|---|
| SPAC644.14c | 1.108215269 | 1.013790358 |
| SPBC32H8.13c | 1.009040306 | 1.013945328 |
| SPAC139.05 | 0.802727432 | 1.013989611 |
| SPCC4G3.12c | 0.804843784 | 1.014070425 |
| SPBC31F10.03 | 1.504148873 | 1.014074215 |
| SPAC3A12.09c | 0.964210495 | 1.01415692 |
| SPBC1921.03c | 1.281375015 | 1.014368136 |
| SPAC8F11.03 | 1.059881553 | 1.014391649 |
| SPAC1006.06 | 0.998008778 | 1.014425258 |
| SPBC16G5.16 | 1.1274392 | 1.014944884 |
| SPAC23H3.04 | 0.903393214 | 1.015157186 |
| SPAC25B8.11 | 1.089342699 | 1.015216925 |
| SPBC27B12.05 | 0.931951862 | 1.015419207 |
| SPAC1B3.01c | 1.097829492 | 1.015852671 |
| SPBC16D10.01c | 1.234127581 | 1.016033278 |
| SPBC2D10.20 | 1.072138434 | 1.01668364 |
| SPAC3C7.10 | 1.1051865 | 1.017008676 |
| SPAPB1E7.07 | 0.995920605 | 1.017128091 |
| SPCC191.11 | 0.794037696 | 1.017305265 |
| SPCC1919.01 | 0.900743816 | 1.017309017 |
| SPCC965.07c | 1.360217118 | 1.017513065 |
| SPBC1604.08c | 1.088589874 | 1.017587384 |
| SPAC1002.03c | 1.048605767 | 1.017952066 |
| SPBC31F10.09c | 1.173033152 | 1.018174068 |
| SPBC16G5.03 | 0.866176673 | 1.018175051 |
| SPAC227.05 | 0.985684147 | 1.018676537 |
| SPAC688.06c | 0.635884111 | 1.018680533 |
| SPBC902.02c | 0.95415196 | 1.019251565 |
| SPAC23C11.08 | 1.165496799 | 1.019469837 |
| SPBC106.07c | 1.199879032 | 1.019618225 |
| SPAC890.06 | 0.983183119 | 1.020172603 |
| SPBC336.01 | 0.836596119 | 1.020670355 |
| SPBC56F2.04 | 0.801124126 | 1.021198286 |
| SPAC27D7.12c | 1.013262208 | 1.021472773 |
| SPAC1B1.04c | 0.824814894 | 1.021590709 |
| SPBC1734.12c | 0.926155659 | 1.021619603 |
| SPCC1235.03 | 1.077220121 | 1.021979848 |
| SPBP8B7.04 | 0.990562337 | 1.022366523 |
| SPBC1703.14c | 1.137956797 | 1.022671182 |
| SPBC27B12.14 | 1.023279279 | 1.023209459 |
| SPCC1827.02c | 1.361702246 | 1.02345187 |
| SPAC6B12.04c | 1.043678262 | 1.023800452 |
| SPAP8A3.14c | 1.142011494 | 1.023868535 |
| SPBC1703.12 | 1.092936871 | 1.024085323 |
| SPBC17D1.07c | 1.096247844 | 1.024105873 |
| SPAC637.06 | 0.834007816 | 1.024121528 |
| SPAC1002.07c | 0.823415818 | 1.024208778 |
| SPBC16G5.02c | 1.054312829 | 1.024660505 |

| | | |
|---|---|---|
| SPAC3H5.10 | 1.095305686 | 1.02475523 |
| SPAC31G5.19 | 0.902791806 | 1.024915476 |
| SPBC4C3.08 | 1.022252011 | 1.025146017 |
| SPAC12G12.10 | 1.931343986 | 1.025166578 |
| SPAC4D7.03 | 1.217011682 | 1.025260445 |
| SPCC1494.07 | 0.923916598 | 1.025385834 |
| SPBC12C2.12c | 1.067358626 | 1.02561292 |
| SPCC306.11 | 0.923244554 | 1.025943129 |
| SPAC630.06c | 1.013774176 | 1.026028293 |
| SPAC24B11.08c | 0.540427523 | 1.02602831 |
| SPAC13D6.04c | 0.964057906 | 1.026032232 |
| SPBC577.13 | 1.126579059 | 1.026087899 |
| SPBC1347.02 | 0.891302285 | 1.026196394 |
| SPCC285.04 | 0.945554511 | 1.026210005 |
| SPAC2E1P5.02c | 0.903903994 | 1.026421311 |
| SPBC23G7.04c | 1.120202181 | 1.026501286 |
| SPAC3G9.05 | 0.677159555 | 1.02659876 |
| SPBC530.05 | 0.878179163 | 1.026803109 |
| SPAC2F7.04 | 0.788744884 | 1.026901432 |
| SPAC3G6.13c | 1.05936624 | 1.027017691 |
| SPBC17D11.08 | 0.968475392 | 1.027138893 |
| SPAC9G1.05 | 0.989595014 | 1.02756482 |
| SPAC6F12.12 | 1.023553493 | 1.027853267 |
| SPBC1604.03c | 1.114453945 | 1.028252141 |
| SPAC821.09 | 1.10682237 | 1.028497482 |
| SPBC660.11 | 0.901530612 | 1.028821625 |
| SPBC56F2.05c | 1.073991789 | 1.028974956 |
| SPAC227.18 | 0.942795076 | 1.029077661 |
| SPCC1223.06 | 0.96977653 | 1.029186635 |
| SPAC1834.05 | 0.85185223 | 1.029275257 |
| SPCC70.06 | 1.01126488 | 1.029325108 |
| SPAC212.03 | 1.091493951 | 1.029617877 |
| SPAC2F3.02 | 1.162312538 | 1.029672174 |
| SPAC6G9.13c | 0.886826204 | 1.030094248 |
| SPAC3C7.07c | 1.010742932 | 1.030167754 |
| SPBC2D10.03c | 1.026736671 | 1.030292955 |
| SPBC30B4.03c | 1.147788457 | 1.0307567 |
| SPBP8B7.27 | 1.086984399 | 1.030835703 |
| SPAC17A2.07c | 0.845725838 | 1.031027309 |
| SPAC15A10.05c | 1.194355892 | 1.03152809 |
| SPAC343.04c | 1.070343191 | 1.031643878 |
| SPAC18G6.12c | 0.989956846 | 1.031838466 |
| SPAC19G12.09 | 1.232457919 | 1.032105779 |
| SPAC664.10 | 1.070943598 | 1.032391914 |
| SPCC777.03c | 1.318256267 | 1.032546025 |
| SPBC19F8.04c | 1.061742655 | 1.032617258 |
| SPBC18E5.01 | 1.093532159 | 1.033124136 |
| SPAC1B3.10c | 1.076323752 | 1.033235628 |

| | | |
|---|---|---|
| SPAC26A3.09c | 1.13676482 | 1.033254461 |
| SPAC18B11.02c | 0.606295075 | 1.033549703 |
| SPBC17D1.06 | 1.113339923 | 1.033555772 |
| SPAC4G9.12 | 1.0385 | 1.033794643 |
| SPBC215.10 | 0.836672869 | 1.033909701 |
| SPAC513.01c | 1.187973704 | 1.033932714 |
| SPAC22G7.02 | 1.237228305 | 1.034130669 |
| SPACUNK4.11c | 0.982929638 | 1.034598475 |
| SPAPB21F2.02 | 0.974280822 | 1.035294234 |
| SPBC23E6.09 | 1.078156468 | 1.035534469 |
| SPAC11D3.07c | 1.159677867 | 1.036092724 |
| SPAC16E8.06c | 1.140339818 | 1.036100363 |
| SPBC16C6.04 | 1.426900945 | 1.036233392 |
| SPAC823.05c | 1.158555857 | 1.036296439 |
| SPAC959.07 | 1.116119052 | 1.036638876 |
| SPAC227.11c | 1.256850282 | 1.036711913 |
| SPAC22A12.01c | 0.97911414 | 1.036744037 |
| SPAC29B12.14c | 0.943542256 | 1.037161164 |
| SPBC8E4.04 | 1.116572037 | 1.037504436 |
| SPAC4G9.19 | 0.86922736 | 1.037544453 |
| SPAC105.03c | 0.999310447 | 1.037559143 |
| SPAC25B8.18 | 0.958200995 | 1.037699778 |
| SPBP23A10.16 | 1.329420945 | 1.038255033 |
| SPAC3G9.03 | 0.849107675 | 1.038327489 |
| SPAC1B3.03c | 1.003604651 | 1.03845 |
| SPAC890.05 | 1.57981584 | 1.038671675 |
| SPCC16C4.09 | 0.896363867 | 1.03875767 |
| SPCC777.07 | 0.997917024 | 1.038960633 |
| SPBC1105.08 | 0.897477558 | 1.039535903 |
| SPAC16A10.02 | 1.034248573 | 1.03962916 |
| SPBC2F12.13 | 0.843117862 | 1.039792375 |
| SPAC2G11.05c | 0.995538838 | 1.039812087 |
| SPBC839.17c | 0.925200791 | 1.040291192 |
| SPAC824.02 | 1.099665346 | 1.040317243 |
| SPAC22F8.04 | 1.137586372 | 1.040437922 |
| SPAPB1E7.08c | 1.060768133 | 1.040457621 |
| SPCC132.01c | 1.14245074 | 1.040603694 |
| SPBC3B9.13c | 2.594327848 | 1.040780487 |
| SPAP11E10.02c | 1.001183632 | 1.040824738 |
| SPAC18G6.13 | 1.048172706 | 1.040900665 |
| SPAP27G11.10c | 0.864868017 | 1.041166223 |
| SPBC19C2.06c | 1.152166432 | 1.041810544 |
| SPCP1E11.02 | 1.174139138 | 1.041971705 |
| SPAC23G3.05c | 1.049795968 | 1.042046693 |
| SPBC16G5.05c | 1.127173209 | 1.042184465 |
| SPAC20G4.04c | 1.316237665 | 1.04231454 |
| SPAC14C4.04 | 0.883286496 | 1.042420486 |
| SPAC12B10.02c | 0.995764618 | 1.042459239 |

| | | |
|---|---|---|
| SPAC9E9.12c | 1.181654001 | 1.042524462 |
| SPBC2G5.03 | 1.097847103 | 1.043045174 |
| SPAC607.02c | 0.956362033 | 1.043187034 |
| SPAC23C11.14 | 1.206646449 | 1.043225513 |
| SPBC21.07c | 1.003299293 | 1.043327769 |
| SPCC320.03 | 1.26778689 | 1.043872034 |
| SPAC31G5.18c | 1.109634459 | 1.043941939 |
| SPBC428.10 | 1.089753984 | 1.044582678 |
| SPAC13G7.04c | 0.918233724 | 1.044727475 |
| SPAC17A5.07c | 1.66696591 | 1.044758736 |
| SPBC31F10.12 | 0.909376127 | 1.045136677 |
| SPBC215.05 | 0.879760052 | 1.045311517 |
| SPBC2F12.15c | 1.003217429 | 1.045805335 |
| SPBPB10D8.06c | 1.065789604 | 1.045836517 |
| SPCC576.14 | 0.906618005 | 1.046532847 |
| SPAC8C9.14 | 0.986159527 | 1.046663252 |
| SPBC530.13 | 1.13750496 | 1.046753917 |
| SPBC27.02c | 0.980305233 | 1.046774066 |
| SPCC31H12.08c | 1.415059504 | 1.046822214 |
| SPAPB21F2.03 | 0.68512962 | 1.047358468 |
| SPBC2G2.17c | 1.209549796 | 1.047717002 |
| SPAC15E1.02c | 0.816089563 | 1.047735896 |
| SPAC13G6.02c | 0.937444347 | 1.048560049 |
| SPAC631.01c | 1.122863993 | 1.048689841 |
| SPBC25H2.08c | 0.990380273 | 1.049285806 |
| SPCC1183.09c | 1.071077758 | 1.049358478 |
| SPAP27G11.15 | 1.042373406 | 1.050048344 |
| SPAC3A12.13c | 0.652453042 | 1.050228846 |
| SPCC757.07c | 0.699777629 | 1.050336317 |
| SPAC23H4.14 | 0.957110321 | 1.050405964 |
| SPBPB2B2.13 | 1.267292551 | 1.050806961 |
| SPAC1006.03c | 1.096480398 | 1.050934618 |
| SPBC19F5.01c | 1.104170828 | 1.051087406 |
| SPAC15A10.15 | 0.89447898 | 1.051195109 |
| SPAC589.12 | 1.074978659 | 1.051649527 |
| SPBC1711.01c | 0.97653399 | 1.051757407 |
| SPAC17C9.15c | 1.139039133 | 1.051926472 |
| SPAC9G1.02 | 1.176084171 | 1.051997026 |
| SPCC1020.07 | 1.341600825 | 1.05228169 |
| SPBC428.02c | 1.085343714 | 1.052351028 |
| SPAC6F6.06c | 1.103775805 | 1.052490136 |
| SPCC191.10 | 1.154460145 | 1.052513859 |
| SPCC417.06c | 1.103963745 | 1.052655131 |
| SPCC970.10c | 1.007245544 | 1.052973829 |
| SPAC3H8.02 | 1.048273026 | 1.053161578 |
| SPBC21C3.17c | 1.07946249 | 1.053203564 |
| SPAC17G6.13 | 1.081903858 | 1.053276113 |
| SPAC9E9.13 | 0.833474025 | 1.053468697 |

| | | |
|---|---|---|
| SPCC550.11 | 0.978251344 | 1.053641893 |
| SPAC3F10.10c | 1.108025794 | 1.053930209 |
| SPBC4F6.15c | 0.97527281 | 1.053985188 |
| SPAC31G5.15 | 0.91165936 | 1.054266045 |
| SPCC330.02 | 1.110720789 | 1.054281093 |
| SPAC1687.08 | 0.973853643 | 1.054331813 |
| SPAC23E2.01 | 1.098630933 | 1.054372958 |
| SPAC22H10.03c | 0.965304886 | 1.055061897 |
| SPBC106.20 | 1.107677602 | 1.055210523 |
| SPCC1235.12c | 0.973368189 | 1.055936874 |
| SPAC1F7.08 | 1.06416516 | 1.056340416 |
| SPAC144.05 | 1.154660682 | 1.056569087 |
| SPAC1B1.02c | 1.080214726 | 1.056839781 |
| SPCC663.12 | 1.064172691 | 1.056855996 |
| SPBC16A3.08c | 0.980789079 | 1.056988192 |
| SPAC9G1.04 | 1.133212209 | 1.057106195 |
| SPAC167.01 | 1.041151877 | 1.057402833 |
| SPBC13E7.03c | 1.045819334 | 1.057534639 |
| SPBP19A11.02c | 1.030468946 | 1.057798375 |
| SPAC17H9.14c | 1.069422584 | 1.058537029 |
| SPAC27F1.05c | 1.157533092 | 1.058567789 |
| SPAC644.11c | 0.914044095 | 1.058571428 |
| SPCC23B6.01c | 1.057036485 | 1.058662695 |
| SPBC1105.18c | 1.126026277 | 1.058701213 |
| SPAP7G5.04c | 1.014004598 | 1.058770746 |
| SPBC1683.12 | 1.353893103 | 1.059504012 |
| SPAC732.02c | 0.542837382 | 1.059524163 |
| SPAC6G9.08 | 1.205419062 | 1.059741276 |
| SPCC4G3.03 | 0.854205494 | 1.059962822 |
| SPCC1682.07 | 0.927684047 | 1.06012087 |
| SPBP23A10.14c | 0.521878528 | 1.060194175 |
| SPBC1703.07 | 0.858230634 | 1.060423086 |
| SPCC63.13 | 1.250294117 | 1.060731189 |
| SPBC1773.09c | 1.231286282 | 1.061096471 |
| SPAC323.04 | 1.032639356 | 1.061786434 |
| SPBC19F8.03c | 1.075421262 | 1.061861543 |
| SPAC1805.06c | 1.073703259 | 1.061876227 |
| SPBC30D10.05c | 1.244823292 | 1.062283896 |
| SPAC977.14c | 1.186789618 | 1.062332932 |
| SPBC342.03 | 1.218585526 | 1.062358655 |
| SPBC839.14c | 0.936118918 | 1.06267114 |
| SPBC30D10.16 | 1.005694212 | 1.062707784 |
| SPAC4A8.06c | 1.186294786 | 1.063362081 |
| SPAC19A8.14 | 1.008513462 | 1.063381572 |
| SPAC13G7.02c | 1.114748977 | 1.063559647 |
| SPBC1683.06c | 1.075175376 | 1.063714918 |
| SPBC1861.03 | 1.090985578 | 1.064306116 |
| SPAC1687.21 | 1.01965001 | 1.064310391 |

| | | |
|---|---|---|
| SPCC1827.08c | 1.292337662 | 1.064366883 |
| SPBC21.05c | 0.98136606 | 1.064658212 |
| SPBC839.06 | 0.59950862 | 1.06500868 |
| SPBC947.02 | 0.829464861 | 1.065250242 |
| SPBC16E9.19 | 0.895481614 | 1.065281455 |
| SPAPB18E9.04c | 0.949571141 | 1.065326372 |
| SPAC4H3.13 | 1.151875733 | 1.065932818 |
| SPAC25H1.09 | 1.069343486 | 1.065959816 |
| SPBC1709.13c | 0.740728106 | 1.06618636 |
| SPAC890.07c | 0.925652473 | 1.066198569 |
| SPAC8E11.05c | 0.854583325 | 1.066307362 |
| SPBP4H10.04 | 1.137884332 | 1.066360174 |
| SPAC30D11.12 | 1.302943235 | 1.067843936 |
| SPAC630.10 | 1.160246322 | 1.067845536 |
| SPCC1322.05c | 1.239012739 | 1.067861764 |
| SPAC4G9.10 | 0.938768594 | 1.068372936 |
| SPAC2F7.10 | 1.103127329 | 1.068388271 |
| SPAPB8E5.04c | 1.167120936 | 1.068498522 |
| SPAC2G11.10c | 1.37038177 | 1.068873217 |
| SPBC20F10.02c | 1.261788174 | 1.069583578 |
| SPBPB7E8.01 | 1.878530512 | 1.069944436 |
| SPAP7G5.03 | 0.925518018 | 1.070093707 |
| SPAC6F12.09 | 1.240269069 | 1.070131044 |
| SPAC1687.19c | 1.124095167 | 1.070200485 |
| SPAC23C4.16c | 1.118333528 | 1.07032557 |
| SPBC1539.08 | 1.457311865 | 1.07068018 |
| SPAC823.02 | 1.122939391 | 1.071644894 |
| SPAC18B11.07c | 1.025468816 | 1.071819762 |
| SPCC1620.12c | 0.941975445 | 1.072310988 |
| SPAC15A10.08 | 1.468722371 | 1.072744232 |
| SPBC18H10.09 | 1.071614923 | 1.072878619 |
| SPAC13C5.03 | 1.172973536 | 1.073603911 |
| SPBC3D6.13c | 1.155714534 | 1.073778582 |
| SPAC24H6.13 | 0.968385078 | 1.074272194 |
| SPCC126.15c | 1.335240696 | 1.074374925 |
| SPAC3A12.17c | 0.887045622 | 1.074379619 |
| SPAC17A2.06c | 1.095776879 | 1.074648568 |
| SPAC824.03c | 0.883110513 | 1.075154349 |
| SPAC1F12.04c | 1.077834822 | 1.075380721 |
| SPAC227.06 | 1.315383752 | 1.076190885 |
| SPBC30B4.08 | 1.047489216 | 1.076315132 |
| SPBC29B5.02c | 1.225754282 | 1.076335134 |
| SPCC24B10.20 | 0.703268803 | 1.07675266 |
| SPBC609.02 | 1.172491579 | 1.076790425 |
| SPCC4B3.10c | 0.505697246 | 1.077069457 |
| SPBC32F12.11 | 1.517008503 | 1.077568282 |
| SPBC1A4.05 | 1.089305109 | 1.077895537 |
| SPAC22H10.04 | 1.139202219 | 1.078083208 |

| | | |
|---|---|---|
| SPAC26H5.08c | 1.105018122 | 1.078318178 |
| SPCC622.19 | 1.068927336 | 1.079240796 |
| SPAC513.06c | 1.042591185 | 1.07938646 |
| SPAC23C4.05c | 1.11236815 | 1.079482133 |
| SPAC27E2.01 | 0.855823656 | 1.080139027 |
| SPBC3H7.13 | 1.037738488 | 1.080259334 |
| SPCC622.01c | 1.120055147 | 1.080357901 |
| SPAC1565.02c | 1.082894737 | 1.080592105 |
| SPBC3B9.06c | 0.987322057 | 1.080848994 |
| SPAC31G5.07 | 1.095022422 | 1.081308269 |
| SPAC1952.09c | 0.956039375 | 1.082480693 |
| SPBC409.08 | 1.082765802 | 1.082590577 |
| SPAC4F10.17 | 1.308157445 | 1.082656018 |
| SPBC3E7.08c | 0.90234411 | 1.082816828 |
| SPBC1711.12 | 1.032693981 | 1.08315989 |
| SPAC343.16 | 0.704751934 | 1.083831094 |
| SPAC2F3.01 | 1.149359728 | 1.083981108 |
| SPBC29A3.01 | 0.902998299 | 1.084145375 |
| SPAC3F10.18c | 0.986645341 | 1.084178828 |
| SPBC16H5.13 | 1.031209985 | 1.084342061 |
| SPAC589.05c | 1.360701639 | 1.084371111 |
| SPAC22E12.18 | 1.161031323 | 1.084399158 |
| SPAPB1A10.15 | 1.042451876 | 1.084433017 |
| SPBC21B10.03c | 0.947783029 | 1.084664371 |
| SPBC19G7.02 | 1.040225086 | 1.084770426 |
| SPAC4D7.06c | 1.022529476 | 1.084869279 |
| SPAC22F3.06c | 1.074813783 | 1.084959028 |
| SPAC823.03 | 1.282407714 | 1.085270926 |
| SPAC227.01c | 0.899406911 | 1.085343249 |
| SPBC6B1.09c | 1.056778232 | 1.085343578 |
| SPBC530.08 | 0.952281744 | 1.08540643 |
| SPCC24B10.19c | 1.100626669 | 1.08552861 |
| SPBC14C8.17c | 2.647520972 | 1.085595308 |
| SPBC4C3.12 | 1.064861097 | 1.085804464 |
| SPAC19G12.04 | 1.203962414 | 1.086230604 |
| SPCC965.12 | 0.987956037 | 1.086712374 |
| SPCC24B10.17 | 1.019210134 | 1.0875357 |
| SPAC11D3.15 | 1.083450574 | 1.087826269 |
| SPBC16G5.11c | 1.234427199 | 1.087960488 |
| SPAPB2B4.02 | 0.989208828 | 1.088343417 |
| SPAP27G11.05c | 0.980175776 | 1.088439436 |
| SPAC1A6.10 | 1.124184702 | 1.088561232 |
| SPBC365.13c | 0.962734257 | 1.088754372 |
| SPCC338.05c | 1.025286763 | 1.089087901 |
| SPAC140.01 | 1.331280035 | 1.089367628 |
| SPAC13G6.03 | 0.988078646 | 1.090115885 |
| SPBC405.05 | 1.360195741 | 1.090246793 |
| SPBC4F6.09 | 0.957225896 | 1.090292822 |

| | | |
|---|---|---|
| SPBC21C3.18 | 1.193220086 | 1.090340399 |
| SPBC21D10.10 | 1.206849286 | 1.090357249 |
| SPBC2D10.19c | 1.030481072 | 1.090989748 |
| SPBC3H7.05c | 1.114568444 | 1.091241754 |
| SPAC343.19 | 1.380231523 | 1.091245548 |
| SPBC1E8.02 | 1.010076557 | 1.091591068 |
| SPBC15C4.01c | 0.993877163 | 1.092291079 |
| SPCC757.09c | 1.673071803 | 1.093225589 |
| SPCC13B11.03c | 1.061608486 | 1.093745495 |
| SPAC1F7.09c | 0.798288038 | 1.093771111 |
| SPBC1271.09 | 0.68544596 | 1.094064968 |
| SPBC6B1.02 | 1.103087046 | 1.094365716 |
| SPAC4F8.01 | 1.319346214 | 1.094368763 |
| SPBC1709.14 | 1.052308046 | 1.094555625 |
| SPBC609.04 | 0.846170567 | 1.094639807 |
| SPCC297.04c | 1.354070067 | 1.094810088 |
| SPAC17A2.12 | 1.186816782 | 1.094932362 |
| SPBC2G2.14 | 0.631683014 | 1.095244813 |
| SPBC1706.03 | 1.161133802 | 1.095512793 |
| SPAC12B10.01c | 0.675260616 | 1.095556359 |
| SPBC29A3.12 | 1.207610448 | 1.095584086 |
| SPAC22F3.12c | 1.338691448 | 1.096861241 |
| SPAC1002.06c | 1.220462301 | 1.096956127 |
| SPBP23A10.10 | 1.135780675 | 1.097367703 |
| SPBP4G3.03 | 1.263349072 | 1.097519798 |
| SPAC11D3.10 | 1.172252836 | 1.097602327 |
| SPBC26H8.11c | 1.085584171 | 1.098058653 |
| SPCC1450.07c | 1.112370486 | 1.098600182 |
| SPCPB1C11.01 | 1.063274426 | 1.098798069 |
| SPCC31H12.02c | 1.482621317 | 1.098915367 |
| SPBC1604.01 | 1.13533762 | 1.098954984 |
| SPAC1952.08c | 1.241711852 | 1.099611993 |
| SPAC6B12.09 | 1.187360231 | 1.099985375 |
| SPBP4H10.18c | 1.352427649 | 1.100713986 |
| SPBC342.06c | 1.128823113 | 1.101040751 |
| SPAC1486.01 | 1.089512384 | 1.10119279 |
| SPCC338.04 | 0.882614543 | 1.10167349 |
| SPBC359.05 | 1.857738573 | 1.101682759 |
| SPBC557.04 | 1.13809224 | 1.101748454 |
| SPBC1271.01c | 1.198682229 | 1.101880177 |
| SPBC15D4.02 | 1.124154402 | 1.101946801 |
| SPAC9E9.03 | 1.060963935 | 1.101991827 |
| SPCC576.12c | 1.001003062 | 1.102597403 |
| SPAC3C7.01c | 0.675165551 | 1.102835543 |
| SPBC3B9.15c | 1.131018019 | 1.102847682 |
| SPAC3G6.04 | 1.021401595 | 1.102970388 |
| SPAC19E9.03 | 1.013087201 | 1.103123523 |
| SPAC24H6.04 | 0.901669962 | 1.103207342 |

| | | |
|---|---|---|
| SPAC10F6.15 | 0.887532015 | 1.10322467 |
| SPAC6B12.07c | 1.19199798 | 1.103271892 |
| SPAC328.04 | 1.041426342 | 1.103347726 |
| SPBC1271.14 | 1.639462734 | 1.103394933 |
| SPBP4H10.07 | 1.078043143 | 1.103993019 |
| SPAC23A1.04c | 0.886733929 | 1.104203101 |
| SPBC83.02c | 1.027377615 | 1.104233682 |
| SPBC428.15 | 1.140712407 | 1.104358006 |
| SPCC285.05 | 1.149050426 | 1.104831778 |
| SPAP8A3.03 | 1.068646845 | 1.105363115 |
| SPAP14E8.04 | 1.075259197 | 1.105499639 |
| SPAC5H10.08c | 1.068191111 | 1.105775291 |
| SPBP35G2.14 | 1.066230418 | 1.1061601 |
| SPBC16H5.07c | 1.034711031 | 1.107010503 |
| SPAC18B11.04 | 1.183261887 | 1.10712852 |
| SPAC19G12.12 | 1.154990326 | 1.107465135 |
| SPBC577.11 | 1.04752411 | 1.107836428 |
| SPAC9E9.11 | 0.98033903 | 1.108033398 |
| SPBP16F5.04 | 1.332517186 | 1.10827358 |
| SPBC365.08c | 0.911997207 | 1.108509919 |
| SPAC11D3.17 | 1.363688105 | 1.108950932 |
| SPAC8E11.03c | 1.210543897 | 1.10918294 |
| SPBC1347.07 | 0.967226883 | 1.109546913 |
| SPAC17H9.09c | 1.22616975 | 1.11000408 |
| SPAC11D3.18c | 1.340810848 | 1.11005667 |
| SPBC1778.01c | 1.02866743 | 1.110457286 |
| SPBC18E5.07 | 1.415165822 | 1.11072296 |
| SPBC609.03 | 1.126025354 | 1.110785913 |
| SPBC1685.06 | 1.207444457 | 1.110934303 |
| SPAC4H3.03c | 1.254021978 | 1.110996369 |
| SPAC13G7.07 | 1.357530286 | 1.111468735 |
| SPAC167.04 | 1.506503744 | 1.111566101 |
| SPAC6B12.16 | 1.156809079 | 1.111669727 |
| SPAC1687.23c | 1.306679526 | 1.11172977 |
| SPCC1259.10 | 1.189649247 | 1.112014529 |
| SPBC2F12.04 | 1.126195652 | 1.112083569 |
| SPBC902.04 | 1.119098869 | 1.112090085 |
| SPCC663.09c | 1.108025725 | 1.112121035 |
| SPAC9.10 | 1.042701455 | 1.112149617 |
| SPBC1652.01 | 1.121088423 | 1.112336757 |
| SPAC1B3.04c | 1.146595581 | 1.112414255 |
| SPBC18H10.06c | 1.342551657 | 1.112418754 |
| SPAC9G1.10c | 0.991991252 | 1.112933781 |
| SPBC800.03 | 0.941586841 | 1.112951899 |
| SPBC3H7.06c | 1.089853191 | 1.113619999 |
| SPAC1071.12c | 1.09460177 | 1.11363385 |
| SPBC1711.15c | 0.757188547 | 1.113824565 |
| SPBC29A10.05 | 1.251279273 | 1.114259756 |

| | | |
|---|---|---|
| SPBPB2B2.14c | 1.171241283 | 1.114305194 |
| SPBC1778.04 | 0.835366746 | 1.114373109 |
| SPAC869.08 | 1.148012593 | 1.115149055 |
| SPAC15E1.06 | 1.080842199 | 1.115445989 |
| SPAC22G7.07c | 1.298904618 | 1.11573913 |
| SPCC1840.12 | 1.029806969 | 1.115841816 |
| SPBC24C6.09c | 0.981282626 | 1.11786372 |
| SPAPB1A11.02 | 1.105524115 | 1.117942701 |
| SPBC16E9.02c | 1.350570825 | 1.118945455 |
| SPBC887.10 | 1.311253003 | 1.119168726 |
| SPAC5H10.09c | 1.230534804 | 1.119313389 |
| SPCC1393.07c | 1.054426643 | 1.12011266 |
| SPCC777.17c | 1.114718871 | 1.120183179 |
| SPCC16C4.12 | 1.139253661 | 1.120217289 |
| SPCC757.02c | 1.116466726 | 1.120345558 |
| SPBC354.15 | 0.971794309 | 1.120468555 |
| SPAC24B11.10c | 1.008362426 | 1.120754717 |
| SPBC947.04 | 1.112565235 | 1.121198816 |
| SPAC17H9.04c | 1.149452244 | 1.121284457 |
| SPAC6F6.17 | 1.201113732 | 1.121341039 |
| SPAC24H6.02c | 0.841010401 | 1.121347201 |
| SPCC18B5.11c | 1.272839207 | 1.121362085 |
| SPAP8A3.05 | 1.059299218 | 1.121449089 |
| SPBC17A3.08 | 1.161826087 | 1.122065217 |
| SPBPB10D8.01 | 1.148977285 | 1.122200991 |
| SPAC1093.03 | 0.911231864 | 1.122451912 |
| SPAC23C11.06c | 1.120875558 | 1.122681901 |
| SPAC1002.14 | 1.064084696 | 1.122933439 |
| SPAC3A11.04 | 1.03146778 | 1.123627168 |
| SPAC1805.07c | 1.224831482 | 1.123715057 |
| SPBC30D10.13c | 0.630197519 | 1.123773633 |
| SPCC548.06c | 0.95931911 | 1.123773814 |
| SPCC16A11.07 | 1.323981015 | 1.123854873 |
| SPAC29E6.05c | 1.130938967 | 1.124363182 |
| SPAC23C11.07 | 1.122135417 | 1.124441964 |
| SPBC16G5.17 | 0.934483596 | 1.124650248 |
| SPAC30.04c | 1.078077695 | 1.124808494 |
| SPBP4H10.10 | 1.124944978 | 1.125495816 |
| SPCC1322.08 | 1.068979671 | 1.12597859 |
| SPBC337.04 | 1.145987966 | 1.126120444 |
| SPAC16C9.05 | 1.148800058 | 1.126391437 |
| SPAC589.02c | 2.346538515 | 1.126755825 |
| SPAC22A12.17c | 1.053328242 | 1.126781397 |
| SPCP20C8.02c | 1.096238921 | 1.126943895 |
| SPCC553.12c | 1.11566068 | 1.127044629 |
| SPAC458.04c | 1.178152147 | 1.127374946 |
| SPAP32A8.02 | 1.190798687 | 1.127573544 |
| SPCC24B10.12 | 1.408909462 | 1.12777201 |

| | | |
|---|---|---|
| SPAC6G10.11c | 1.235339885 | 1.12817943 |
| SPAC926.09c | 1.170205378 | 1.128439589 |
| SPAC1805.11c | 1.095901357 | 1.12890265 |
| SPAC8E11.01c | 1.017449687 | 1.128907104 |
| SPBC19G7.04 | 1.233991627 | 1.129343951 |
| SPAC1F5.03c | 1.213200935 | 1.129453855 |
| SPBC31A8.01c | 0.392616343 | 1.129737359 |
| SPAC2E1P3.02c | 1.164149318 | 1.130026608 |
| SPBC16E9.13 | 1.304831665 | 1.130141402 |
| SPAC22H10.07 | 1.099150246 | 1.130430367 |
| SPBC30D10.14 | 1.075155436 | 1.130678168 |
| SPAC1610.04 | 1.152160102 | 1.130926382 |
| SPAC4A8.09c | 1.375041528 | 1.131 |
| SPAC56F8.12 | 1.073536024 | 1.131296175 |
| SPBC3D6.05 | 1.201731641 | 1.13136357 |
| SPAC17H9.01 | 1.235608705 | 1.131559284 |
| SPAC869.02c | 0.932538871 | 1.131565666 |
| SPAC22H10.08 | 1.140466518 | 1.131660488 |
| SPAC26A3.02 | 1.138863594 | 1.131812931 |
| SPAC1142.01 | 1.013958736 | 1.131855402 |
| SPAC1565.04c | 1.401881973 | 1.131976879 |
| SPAC12B10.15c | 1.393353888 | 1.132033659 |
| SPBC21B10.04c | 1.021699485 | 1.132987749 |
| SPBC2A9.04c | 1.03751708 | 1.133023691 |
| SPAC6G9.04 | 1.047419006 | 1.133137225 |
| SPAC22H10.09 | 1.427684012 | 1.133444919 |
| SPAC144.03 | 1.174297057 | 1.133635057 |
| SPAC1805.03c | 1.240502092 | 1.134244417 |
| SPBC83.19c | 1.127252969 | 1.1342683 |
| SPBC19C7.02 | 1.232320086 | 1.134490446 |
| SPBC1198.08 | 1.126973152 | 1.134676748 |
| SPBC19F8.08 | 1.330677532 | 1.134718958 |
| SPBC2A9.03 | 1.240890736 | 1.134918317 |
| SPAC23D3.09 | 1.384805485 | 1.134940822 |
| SPAC57A10.10c | 1.141369383 | 1.135392517 |
| SPBC16A3.06 | 0.849591684 | 1.135600425 |
| SPAC8C9.19 | 1.308310389 | 1.135897392 |
| SPBC21B10.12 | 1.355439331 | 1.136095338 |
| SPCC1450.03 | 0.863072137 | 1.136713376 |
| SPCC1450.09c | 1.130112973 | 1.136851953 |
| SPAC26H5.03 | 1.149118296 | 1.137088997 |
| SPBC4F6.06 | 1.75690332 | 1.137351716 |
| SPAC29A4.09 | 1.897327748 | 1.137540373 |
| SPAC26F1.02 | 1.234278233 | 1.137593369 |
| SPCPB16A4.04c | 1.020440033 | 1.137728936 |
| SPBC11C11.06c | 1.141184809 | 1.138319496 |
| SPBC6B1.08c | 1.0304013 | 1.138635512 |
| SPBPB10D8.07c | 1.066981478 | 1.138646862 |

| | | |
|---|---|---|
| SPAC1F8.03c | 1.07656202 | 1.139124177 |
| SPAC824.07 | 0.99517696 | 1.139401632 |
| SPAC5H10.07 | 1.296257089 | 1.139425804 |
| SPAC2F7.02c | 1.079075319 | 1.139632924 |
| SPAC630.15 | 1.114605721 | 1.140306648 |
| SPCC1919.04 | 1.091651516 | 1.140338072 |
| SPAC3F10.12c | 1.265510286 | 1.140694332 |
| SPBC26H8.05c | 1.393816954 | 1.140780479 |
| SPBC16C6.09 | 1.413472333 | 1.141241379 |
| SPBC27.03 | 1.081475264 | 1.14138423 |
| SPBC2D10.11c | 1.171047989 | 1.142144412 |
| SPBC30D10.03c | 1.078371151 | 1.143106274 |
| SPBC354.10 | 0.736549866 | 1.143536575 |
| SPBC11G11.02c | 1.240535675 | 1.143672808 |
| SPBC947.03c | 1.211070835 | 1.144424913 |
| SPBC18H10.16 | 1.183189033 | 1.144886364 |
| SPBC1718.02 | 1.229088925 | 1.146012467 |
| SPBC685.04c | 0.683630575 | 1.146549491 |
| SPAC13G6.08 | 1.222739019 | 1.146654655 |
| SPBC16H5.05c | 1.18154316 | 1.146821376 |
| SPAC23H4.01c | 1.136383141 | 1.14690153 |
| SPBP16F5.08c | 1.091549842 | 1.147380543 |
| SPAC29E6.07 | 1.268490153 | 1.147419115 |
| SPAC30C2.05 | 1.16898142 | 1.147434517 |
| SPAPB17E12.14c | 1.021907496 | 1.147775516 |
| SPBC725.03 | 1.02519685 | 1.14800689 |
| SPAC3H1.04c | 1.296372481 | 1.148120222 |
| SPBC16G5.13 | 0.961062513 | 1.148353211 |
| SPAC14C4.11 | 1.326838499 | 1.148840859 |
| SPAC2G11.15c | 0.938208776 | 1.148935028 |
| SPBP16F5.03c | 1.241260349 | 1.149172688 |
| SPBC17A3.02 | 1.05030596 | 1.14926173 |
| SPBC31F10.17c | 1.416355556 | 1.149583333 |
| SPAC1834.03c | 1.064988381 | 1.14978781 |
| SPBP22H7.06 | 1.99625213 | 1.149874894 |
| SPCC417.09c | 1.131894186 | 1.149939499 |
| SPAC823.14 | 1.275078859 | 1.150724861 |
| SPAC513.04 | 1.051434117 | 1.15118289 |
| SPAC1A6.05c | 0.670356408 | 1.151345099 |
| SPAC2F3.15 | 0.882111505 | 1.151666935 |
| SPAC19B12.10 | 1.080578835 | 1.151810043 |
| SPAC22F8.09 | 1.117776989 | 1.151946324 |
| SPAC17C9.14 | 1.286811995 | 1.151956154 |
| SPBC106.03 | 1.132518189 | 1.152311954 |
| SPCC1620.08 | 1.093021452 | 1.152356902 |
| SPCC330.07c | 1.25224727 | 1.152438013 |
| SPBC3B8.06 | 1.071806634 | 1.152518673 |
| SPAC5D6.08c | 1.16133983 | 1.152770188 |

| | | |
|---|---|---|
| SPBC36B7.02 | 1.191413287 | 1.15287641 |
| SPBC3B9.05 | 1.039055929 | 1.153042098 |
| SPAC17G8.09 | 1.0005468 | 1.153061756 |
| SPBC342.05 | 1.211904121 | 1.15330256 |
| SPBC16H5.04 | 1.214536957 | 1.153385632 |
| SPAC26F1.08c | 1.330456268 | 1.154170173 |
| SPCC548.04 | 1.04038424 | 1.154172449 |
| SPBC16E9.17c | 0.863730588 | 1.154462984 |
| SPBC16H5.12c | 1.155458557 | 1.154575854 |
| SPAC17H9.12c | 1.149544073 | 1.154675966 |
| SPBC1711.09c | 0.91089993 | 1.154860404 |
| SPBC1105.12 | 1.110244912 | 1.155251395 |
| SPBP8B7.11 | 1.06161478 | 1.155341259 |
| SPBC839.15c | 1.748110133 | 1.15549597 |
| SPAC4F10.08 | 1.239679962 | 1.155551087 |
| SPAC23A1.14c | 1.175802673 | 1.155804756 |
| SPBC216.06c | 1.173005696 | 1.156002495 |
| SPAC31A2.02 | 1.459777361 | 1.156049817 |
| SPCC757.03c | 1.181802486 | 1.156312628 |
| SPAC2F3.12c | 1.147807607 | 1.156322958 |
| SPAC1A6.04c | 0.93098818 | 1.156783626 |
| SPBC1703.04 | 1.261621651 | 1.157051148 |
| SPBP4H10.09 | 1.534352665 | 1.157457659 |
| SPAC5H10.13c | 1.022784755 | 1.157633696 |
| SPBC1778.07 | 0.972633331 | 1.157905686 |
| SPCC31H12.05c | 0.613388531 | 1.157939267 |
| SPAC4F10.18 | 1.11709637 | 1.158171972 |
| SPBC577.06c | 0.921690383 | 1.158230272 |
| SPCC1223.01 | 1.048315451 | 1.158356429 |
| SPAC2F7.09c | 1.090819028 | 1.158714863 |
| SPAC4G9.05 | 1.224545838 | 1.159062104 |
| SPBC16D10.08c | 1.177123235 | 1.159896414 |
| SPAC5H10.11 | 1.142788817 | 1.160484683 |
| SPAC823.13c | 1.282391752 | 1.160759948 |
| SPAC11G7.01 | 0.666659488 | 1.160805476 |
| SPAC1F8.06 | 0.870504476 | 1.160857051 |
| SPBC1683.07 | 0.849780012 | 1.160864865 |
| SPBC14F5.13c | 1.272582857 | 1.160870692 |
| SPCC320.07c | 0.979127828 | 1.1609266 |
| SPBC2G5.06c | 1.177707229 | 1.161186153 |
| SPAC3G6.06c | 1.307917692 | 1.161512827 |
| SPBC30D10.18c | 1.031495414 | 1.162437508 |
| SPBC36.04 | 1.214106303 | 1.16306965 |
| SPAC25G10.01 | 1.10082581 | 1.163354521 |
| SPBC3D6.04c | 0.810627836 | 1.163387129 |
| SPCC1259.14c | 0.994195212 | 1.163728849 |
| SPAC18G6.05c | 1.042152244 | 1.163895971 |
| SPAC1565.01 | 1.107142857 | 1.163907385 |

| | | |
|---|---|---|
| SPAC22G7.04 | 1.233564014 | 1.164008434 |
| SPCC320.14 | 1.267787187 | 1.164278223 |
| SPAC139.04c | 1.344879639 | 1.164511412 |
| SPAC688.13 | 1.50257176 | 1.164531446 |
| SPAC16C9.02c | 1.050540629 | 1.164846909 |
| SPCC338.16 | 2.210484264 | 1.165615308 |
| SPCC737.06c | 0.885568745 | 1.165719281 |
| SPCC191.01 | 1.134918608 | 1.166189408 |
| SPAC17G8.11c | 1.198928586 | 1.167332933 |
| SPCC1450.12 | 0.92430056 | 1.167378598 |
| SPBC17D1.02 | 1.024501424 | 1.167467949 |
| SPBC30D10.10c | 0.84323829 | 1.167489632 |
| SPCC1494.01 | 1.202327201 | 1.167638052 |
| SPCC70.10 | 1.258618264 | 1.167640975 |
| SPBC14C8.09c | 1.113381755 | 1.167864514 |
| SPAC1B3.06c | 1.394882608 | 1.168265628 |
| SPCC285.14 | 1.059416678 | 1.168435839 |
| SPBC4B4.04 | 1.164806677 | 1.168515643 |
| SPCC1739.01 | 1.312374133 | 1.168660837 |
| SPBC2D10.07c | 1.341509434 | 1.169221698 |
| SPBC14C8.04 | 1.175685037 | 1.169748137 |
| SPBC3B8.10c | 1.303649387 | 1.170353992 |
| SPAC823.16c | 1.393575231 | 1.170465007 |
| SPBC20F10.06 | 1.25265057 | 1.1705981 |
| SPAC23A1.03 | 1.095548648 | 1.170837413 |
| SPAC323.01c | 0.901726857 | 1.171718359 |
| SPAC8C9.12c | 1.259695852 | 1.171772218 |
| SPBC2D10.17 | 0.705609257 | 1.171777798 |
| SPBC2G5.02c | 1.356342494 | 1.172095041 |
| SPBCPT2R1.02 | 1.10841456 | 1.172238476 |
| SPAC29B12.02c | 1.223030754 | 1.173621729 |
| SPBC359.01 | 1.075987989 | 1.174024902 |
| SPBC19C7.05 | 1.101980506 | 1.174173746 |
| SPAC27F1.10 | 1.28302546 | 1.174184497 |
| SPAC1039.01 | 1.369968944 | 1.174233307 |
| SPBC1289.10c | 1.671538722 | 1.174326313 |
| SPBC19F8.01c | 1.293811218 | 1.174368463 |
| SPCC74.04 | 1.192824808 | 1.174592065 |
| SPAC24B11.06c | 1.196257897 | 1.174669966 |
| SPBC29A3.21 | 1.1047434 | 1.174887202 |
| SPAC343.10 | 1.191922518 | 1.175280488 |
| SPAC977.10 | 1.211503899 | 1.175725742 |
| SPCC622.14 | 1.711885756 | 1.175762243 |
| SPAC664.07c | 0.997398067 | 1.175814443 |
| SPBC14F5.07 | 1.232642666 | 1.176287926 |
| SPBC36B7.08c | 1.268432158 | 1.176374671 |
| SPAC13F5.07c | 1.398924642 | 1.176447074 |
| SPAC959.05c | 1.356755984 | 1.176872074 |

| | | |
|---|---|---|
| SPAC2F7.08c | 1.249819946 | 1.176918291 |
| SPBC27B12.03c | 1.0126226 | 1.177492876 |
| SPCC2H8.02 | 0.974781342 | 1.177540562 |
| SPBC582.10c | 0.772733135 | 1.17844152 |
| SPAC24B11.07c | 1.096273453 | 1.178698448 |
| SPAC19B12.11c | 1.266138023 | 1.178810655 |
| SPBC21B10.05c | 1.094814224 | 1.179048312 |
| SPCC1020.10 | 0.998838074 | 1.179108272 |
| SPCC18.03 | 1.279354454 | 1.179630055 |
| SPCC1682.11c | 1.348049864 | 1.180080189 |
| SPBC2G2.06c | 0.494996914 | 1.180390508 |
| SPAC2F3.18c | 1.177854916 | 1.180464867 |
| SPBC13E7.06 | 1.626778634 | 1.180507546 |
| SPAC3H1.08c | 1.197184043 | 1.180816293 |
| SPBC23G7.13c | 1.208743371 | 1.181658544 |
| SPBC21B10.07 | 1.322856587 | 1.181701918 |
| SPAC14C4.12c | 1.263855436 | 1.181823073 |
| SPAC6F6.12 | 1.682800512 | 1.181825448 |
| SPCC4G3.09c | 1.133121427 | 1.182155367 |
| SPCC1442.04c | 1.668215891 | 1.182398511 |
| SPBC359.04c | 1.181907817 | 1.182959894 |
| SPAC630.05 | 1.15791041 | 1.183308245 |
| SPBC31F10.16 | 1.258495034 | 1.183337983 |
| SPCC569.02c | 1.256380491 | 1.183477117 |
| SPCC794.12c | 1.337866028 | 1.18373422 |
| SPBC16E9.09c | 1.190357636 | 1.184301477 |
| SPAC15A10.06 | 1.287332905 | 1.184313763 |
| SPBC29A10.11c | 1.04686709 | 1.184559518 |
| SPAC1782.04 | 1.255141712 | 1.184923174 |
| SPCC364.03 | 1.363342175 | 1.185626658 |
| SPBPB2B2.08 | 1.191844461 | 1.186341757 |
| SPAC2C4.09 | 1.177662979 | 1.18641439 |
| SPBC1A4.03c | 1.158340342 | 1.186524357 |
| SPAC12B10.05 | 1.193835645 | 1.186901116 |
| SPBC1198.11c | 1.121936315 | 1.187041159 |
| SPAC1783.04c | 1.183681723 | 1.187049112 |
| SPBC32C12.03c | 1.273806487 | 1.18706839 |
| SPAC8F11.05c | 1.258519737 | 1.187155863 |
| SPBC17G9.09 | 1.092315196 | 1.187672894 |
| SPAC23C4.02 | 1.274574433 | 1.187689938 |
| SPBC336.06c | 1.051965679 | 1.18821109 |
| SPBC14C8.15 | 1.268229618 | 1.188478996 |
| SPAC1A6.01c | 1.180079323 | 1.188508461 |
| SPAC16A10.03c | 1.357731541 | 1.188615624 |
| SPAC25H1.03 | 1.349809693 | 1.188752855 |
| SPBC27.08c | 1.068470492 | 1.188757864 |
| SPBC1348.02 | 1.034676676 | 1.188955484 |
| SPBC1539.10 | 1.064080949 | 1.190715919 |

| | | |
|---|---|---|
| SPAC27E2.11c | 1.103026486 | 1.190779613 |
| SPBC21.03c | 1.12174869 | 1.190956335 |
| SPBC36.01c | 1.034033336 | 1.191524177 |
| SPBC4C3.04c | 1.135756182 | 1.19159385 |
| SPAC23H3.11c | 1.151275101 | 1.191839426 |
| SPCC825.05c | 1.096460412 | 1.192352365 |
| SPBC28E12.03 | 1.219727806 | 1.192749317 |
| SPBC405.03c | 0.963377335 | 1.192946614 |
| SPAC1952.02 | 1.333685678 | 1.193017398 |
| SPAC15E1.09 | 1.450768961 | 1.193069754 |
| SPCC1672.09 | 1.031369297 | 1.194080083 |
| SPAC19A8.08 | 1.262546644 | 1.194104092 |
| SPBC1861.02 | 0.768177716 | 1.194137314 |
| SPBC32F12.05c | 1.155763328 | 1.194261627 |
| SPAC18B11.11 | 1.184253316 | 1.194831441 |
| SPBC56F2.09c | 1.349916174 | 1.195299725 |
| SPBC3H7.11 | 1.137933663 | 1.196094717 |
| SPAC16C9.06c | 1.150432172 | 1.196257263 |
| SPBC15D4.06 | 1.39161344 | 1.196526273 |
| SPBPB10D8.04c | 1.160802009 | 1.198072376 |
| SPAC4A8.04 | 1.279087322 | 1.198386066 |
| SPBC27.04 | 1.114698458 | 1.198727127 |
| SPBC19F8.06c | 1.412146548 | 1.198787252 |
| SPAC1250.05 | 1.206452193 | 1.198988478 |
| SPCC1322.14c | 1.089692634 | 1.199543448 |
| SPBP4H10.17c | 1.564633599 | 1.199726219 |
| SPAC12B10.04 | 1.231711999 | 1.200648325 |
| SPAC13F5.05 | 1.067617824 | 1.201137266 |
| SPCC5E4.05c | 1.128684301 | 1.20116738 |
| SPBC2G2.13c | 1.03166896 | 1.201236599 |
| SPAC6F12.02 | 1.432325317 | 1.201833643 |
| SPAC57A7.13 | 1.213479628 | 1.202204041 |
| SPBP35G2.05c | 1.22186059 | 1.202914073 |
| SPBC16H5.08c | 1.192076796 | 1.203191059 |
| SPBC16D10.05 | 0.819802616 | 1.203321177 |
| SPAC1399.03 | 1.245904325 | 1.203840971 |
| SPBC146.04 | 1.080973961 | 1.203991476 |
| SPBC577.12 | 1.189817233 | 1.204614067 |
| SPBC146.09c | 1.211030382 | 1.204747677 |
| SPBC2F12.05c | 1.091501102 | 1.205213565 |
| SPBC23E6.10c | 1.269191061 | 1.205512483 |
| SPBC2D10.12 | 0.938756486 | 1.205756373 |
| SPAC105.02c | 1.203853755 | 1.206151186 |
| SPCC1795.03 | 1.044408161 | 1.206217876 |
| SPAC1805.01c | 0.307501826 | 1.207213224 |
| SPAC23A1.06c | 1.234334374 | 1.207809055 |
| SPBC1711.05 | 0.807685525 | 1.208145355 |
| SPCC777.09c | 1.123030303 | 1.208251345 |

| | | |
|---|---|---|
| SPAC323.07c | 1.171095781 | 1.208675934 |
| SPBC29A10.13 | 2.001153152 | 1.209370885 |
| SPCC1739.08c | 1.267503622 | 1.209391403 |
| SPBC13G1.02 | 1.169092351 | 1.209905661 |
| SPAC227.15 | 1.140212367 | 1.209985553 |
| SPCC1739.04c | 1.182404566 | 1.210179754 |
| SPBC776.11 | 1.086522921 | 1.210315756 |
| SPBC365.06 | 0.422643227 | 1.210579046 |
| SPBC1604.09c | 1.411689736 | 1.210841586 |
| SPAC1952.17c | 1.176243116 | 1.211141682 |
| SPBC902.06 | 1.393350456 | 1.211497731 |
| SPAC1006.04c | 1.373901393 | 1.211716238 |
| SPAC1705.02 | 1.27553565 | 1.21259453 |
| SPBC14C8.11c | 1.150183212 | 1.21301868 |
| SPBC13A2.04c | 1.228661829 | 1.213543113 |
| SPBC18E5.14c | 1.165756371 | 1.214149406 |
| SPAC2C4.08 | 1.129807397 | 1.214185148 |
| SPAC824.09c | 1.30935533 | 1.214300186 |
| SPAC3G6.01 | 0.949308485 | 1.21448909 |
| SPCC1442.07c | 1.300088039 | 1.214746651 |
| SPAC22G7.01c | 1.062684269 | 1.214897869 |
| SPAC14C4.06c | 1.759935021 | 1.215117647 |
| SPBC4B4.12c | 0.82478801 | 1.215771569 |
| SPBC27B12.08 | 1.261204188 | 1.215828549 |
| SPBC1709.12 | 0.991025141 | 1.217165694 |
| SPAC17G6.02c | 1.169020708 | 1.217186088 |
| SPAC1527.01 | 1.40388611 | 1.217739996 |
| SPCC1450.11c | 1.251247898 | 1.217932759 |
| SPCP1E11.03 | 1.688389225 | 1.218252424 |
| SPBC17D11.04c | 1.254037877 | 1.218679666 |
| SPBPB2B2.09c | 1.440233604 | 1.218836817 |
| SPAC13G7.06 | 1.176949013 | 1.219345035 |
| SPBC12D12.07c | 1.3482446 | 1.219432476 |
| SPAC15A10.07 | 1.213434057 | 1.219520653 |
| SPAC4G8.07c | 1.276216968 | 1.220705842 |
| SPCC1827.04 | 1.12759672 | 1.220989327 |
| SPAC3H1.14 | 1.174831639 | 1.221123928 |
| SPCC16C4.20c | 1.0472251 | 1.221137384 |
| SPBC16G5.15c | 1.240887915 | 1.221437489 |
| SPBC713.11c | 1.328179529 | 1.221560477 |
| SPAC630.07c | 1.084343132 | 1.221983864 |
| SPAC17C9.09c | 1.138458395 | 1.222002986 |
| SPBC11B10.08 | 1.332177745 | 1.222109082 |
| SPCC4F11.04c | 1.424379585 | 1.222872483 |
| SPAC458.06 | 1.631128547 | 1.222896945 |
| SPAC27D7.02c | 1.082028195 | 1.222927785 |
| SPAC6B12.12 | 1.39232411 | 1.223316456 |
| SPBC29A10.08 | 1.161410392 | 1.223403727 |

| | | |
|---|---|---|
| SPAC8E11.06 | 1.010385369 | 1.223434438 |
| SPBP8B7.08c | 1.098180636 | 1.223577478 |
| SPBC16A3.02c | 1.054801345 | 1.223591769 |
| SPBC776.15c | 0.768684517 | 1.223624854 |
| SPAPB8E5.06c | 1.231918605 | 1.2243 |
| SPCC1620.07c | 1.158985586 | 1.225084804 |
| SPCC1739.03 | 1.05152 | 1.225275 |
| SPAC18G6.15 | 1.219656841 | 1.225313923 |
| SPBC16C6.01c | 1.140351928 | 1.226053328 |
| SPCC1739.14 | 1.459698558 | 1.226800154 |
| SPBC215.07c | 1.197103803 | 1.227328646 |
| SPAC823.11 | 1.087584185 | 1.22734915 |
| SPCC550.14 | 1.378812549 | 1.227385362 |
| SPAC1071.08 | 1.147631866 | 1.227720721 |
| SPAC10F6.08c | 0.978863421 | 1.228585252 |
| SPAC10F6.11c | 1.213944141 | 1.228849229 |
| SPCC23B6.05c | 1.335687104 | 1.228858351 |
| SPCC4G3.17 | 1.272138229 | 1.229245411 |
| SPCC338.11c | 1.133884573 | 1.229497974 |
| SPAPB1A11.04c | 1.186569522 | 1.229684005 |
| SPAP8A3.02c | 1.076470626 | 1.230188363 |
| SPCC757.04 | 1.243838833 | 1.230334997 |
| SPBC21B10.02 | 1.177448071 | 1.230341246 |
| SPCC794.10 | 1.285399568 | 1.23081447 |
| SPAC25B8.08 | 0.964525483 | 1.231220082 |
| SPCC16A11.15c | 1.120101744 | 1.231777912 |
| SPCC24B10.11c | 1.150658448 | 1.232024097 |
| SPBC216.05 | 1.196984615 | 1.232409452 |
| SPAC1002.19 | 1.065128205 | 1.232760989 |
| SPBC4.06 | 0.903458793 | 1.232884804 |
| SPAC57A10.06 | 1.190764994 | 1.232945345 |
| SPAC328.01c | 1.259825464 | 1.233733973 |
| SPBC4B4.02c | 1.208357156 | 1.234029213 |
| SPAC9E9.10c | 1.292465835 | 1.234113266 |
| SPAC19G12.02c | 1.384905564 | 1.234358297 |
| SPAC1786.02 | 1.177906554 | 1.234609136 |
| SPCC550.10 | 1.204531703 | 1.234644996 |
| SPAPB1A10.12c | 1.329405288 | 1.234831694 |
| SPCC965.05c | 0.791453961 | 1.23503523 |
| SPBC1604.18c | 1.502046471 | 1.235130309 |
| SPBC27B12.11c | 1.187016748 | 1.235454917 |
| SPAC4F8.03 | 3.637816286 | 1.236163886 |
| SPAP27G11.07c | 1.034310435 | 1.23634775 |
| SPAC688.03c | 1.211580217 | 1.236354041 |
| SPBC25D12.06 | 1.088234812 | 1.237597458 |
| SPBC11B10.07c | 0.837143571 | 1.237924207 |
| SPAC1952.11c | 0.607251788 | 1.237959652 |
| SPBC3B9.08c | 1.262923657 | 1.238713198 |

| | | |
|---|---|---|
| SPAC56F8.06c | 1.288779615 | 1.238719502 |
| SPAC630.09c | 1.222861625 | 1.238993326 |
| SPAC630.11 | 1.170538597 | 1.239028219 |
| SPBC29A3.17 | 1.168733417 | 1.239048919 |
| SPAC12G12.07c | 1.15079145 | 1.239135594 |
| SPCC320.05 | 1.037585287 | 1.239289542 |
| SPAC3C7.12 | 1.240002892 | 1.239542788 |
| SPAC977.06 | 1.31650641 | 1.240046575 |
| SPCC1827.03c | 0.969262528 | 1.240964943 |
| SPBC25H2.09 | 1.611987565 | 1.241425105 |
| SPCC338.06c | 1.17312297 | 1.241446858 |
| SPBC4B4.06 | 1.213843857 | 1.241789787 |
| SPAC521.04c | 1.25965337 | 1.242030487 |
| SPBC405.04c | 0.315890449 | 1.24229064 |
| SPCC18.13 | 1.09738292 | 1.24229511 |
| SPBC1734.07c | 1.234455994 | 1.242524276 |
| SPAC27F1.03c | 1.165882825 | 1.242688253 |
| SPAC22F3.11c | 0.962038833 | 1.242970341 |
| SPCC16C4.17 | 1.20556298 | 1.243778539 |
| SPCC584.01c | 1.542381557 | 1.244508506 |
| SPAC29A4.14c | 1.349439733 | 1.245185748 |
| SPAC13A11.05 | 1.24967344 | 1.245385069 |
| SPAC9G1.06c | 0.965783628 | 1.245611181 |
| SPBC3E7.01 | 1.441562017 | 1.245853764 |
| SPBC29A10.01 | 1.408966546 | 1.246748663 |
| SPBC19C2.10 | 1.170900023 | 1.247050462 |
| SPBC25B2.02c | 1.219726899 | 1.247264509 |
| SPBC32H8.01c | 1.174360821 | 1.247479295 |
| SPAC15A10.10 | 1.285369148 | 1.24767279 |
| SPBC3E7.02c | 1.325111607 | 1.247698103 |
| SPAC10F6.14c | 1.460978409 | 1.248162997 |
| SPBP35G2.07 | 1.444400718 | 1.248545703 |
| SPBC28F2.02 | 1.010053971 | 1.249010936 |
| SPAC24B11.12c | 1.201701067 | 1.249766445 |
| SPCC18.15 | 1.09202864 | 1.249895585 |
| SPBC56F2.11 | 2.28664705 | 1.24990072 |
| SPBC1921.07c | 1.157775 | 1.250512791 |
| SPCC4F11.02 | 0.928502287 | 1.251279489 |
| SPAC17G6.08 | 1.27498144 | 1.251391982 |
| SPCC1281.07c | 1.328176744 | 1.251432 |
| SPAC6F6.01 | 0.69652513 | 1.252082226 |
| SPCC1442.14c | 1.285106383 | 1.25265341 |
| SPAC27D7.11c | 1.228164898 | 1.253614209 |
| SPAC2F3.16 | 1.179315099 | 1.254308704 |
| SPAC3A11.09 | 1.208486313 | 1.254615232 |
| SPAC56E4.06c | 1.335050727 | 1.254981625 |
| SPBCPT2R1.08c | 1.134435492 | 1.256406229 |
| SPAC186.02c | 1.148403221 | 1.257411136 |

| | | |
|---|---|---|
| SPAC1250.03 | 1.249671593 | 1.257466133 |
| SPAC22F8.05 | 1.037256618 | 1.257521872 |
| SPAC29B12.10c | 1.144236049 | 1.258465057 |
| SPBC2D10.13 | 1.325380303 | 1.258832065 |
| SPAC7D4.04 | 1.186982639 | 1.258865939 |
| SPBC1685.14c | 1.137554022 | 1.259583987 |
| SPAC31F12.01 | 1.124601872 | 1.259746131 |
| SPAC6B12.02c | 1.855161301 | 1.259927253 |
| SPAC22E12.03c | 1.484055393 | 1.260143623 |
| SPCC1020.08 | 1.314810831 | 1.260188595 |
| SPAPB8E5.10 | 1.331158911 | 1.260230551 |
| SPBC776.01 | 1.383676384 | 1.260519467 |
| SPAC4C5.03 | 1.65876253 | 1.260747926 |
| SPBC16D10.02 | 1.20181197 | 1.261246697 |
| SPAC9.05 | 1.346712634 | 1.261607999 |
| SPAC12B10.12c | 0.948282567 | 1.261949125 |
| SPCC1682.08c | 1.272767843 | 1.261952888 |
| SPCC1450.02 | 1.045410386 | 1.262659629 |
| SPCC1682.15 | 0.948721026 | 1.262734017 |
| SPBC16H5.02 | 1.023006114 | 1.262975206 |
| SPBC29B5.03c | 1.306256597 | 1.263266055 |
| SPBC19C2.04c | 1.489250043 | 1.263384942 |
| SPBC582.08 | 1.518026264 | 1.263443585 |
| SPBC17D11.01 | 1.193486136 | 1.264511748 |
| SPCC1223.10c | 1.513246517 | 1.264523001 |
| SPAC22A12.06c | 0.566087875 | 1.265026251 |
| SPBC21C3.13 | 1.41443556 | 1.265504526 |
| SPBP8B7.10c | 1.240775168 | 1.265584144 |
| SPBC16E9.03c | 1.223448187 | 1.265621063 |
| SPBC21C3.12c | 1.24693234 | 1.265651878 |
| SPAC3G9.01 | 1.09345666 | 1.265745953 |
| SPAC17D4.03c | 1.166534071 | 1.265760856 |
| SPAC8C9.06c | 1.046599737 | 1.267106167 |
| SPBC646.08c | 0.897314233 | 1.268002881 |
| SPAC29B12.04 | 1.001628978 | 1.268251077 |
| SPAC17H9.10c | 1.844938164 | 1.26968918 |
| SPBC1778.03c | 1.459954257 | 1.269993707 |
| SPBC660.09 | 1.59975923 | 1.270026585 |
| SPBC21C3.03 | 1.161640584 | 1.270218418 |
| SPBC1773.01 | 1.410977243 | 1.270268574 |
| SPAC20H4.08 | 1.525376973 | 1.27058398 |
| SPBC418.02 | 1.171609536 | 1.270733321 |
| SPCC1884.02 | 0.923103182 | 1.271132736 |
| SPCC16A11.04 | 0.972082756 | 1.272581602 |
| SPBC23G7.16 | 1.43122865 | 1.272900237 |
| SPCC1682.01 | 1.525204549 | 1.273702432 |
| SPBC8D2.17 | 1.236269875 | 1.273904168 |
| SPAC14C4.01c | 0.948316157 | 1.274899506 |

| | | |
|---|---|---|
| SPAC343.15 | 1.271960825 | 1.274920918 |
| SPAC824.08 | 1.363813362 | 1.27499139 |
| SPBC409.10 | 1.223417278 | 1.275270393 |
| SPCC965.10 | 1.296654472 | 1.276167484 |
| SPAC6G9.14 | 1.245530931 | 1.276250425 |
| SPBC725.05c | 1.309063173 | 1.27632167 |
| SPBC365.14c | 1.064173154 | 1.276472716 |
| SPBC3H7.12 | 1.153669108 | 1.276711133 |
| SPAC27D7.13c | 1.296432774 | 1.277583731 |
| SPCC830.07c | 1.017584994 | 1.278795428 |
| SPAC25B8.07c | 0.819616837 | 1.279047586 |
| SPBC1271.08c | 1.210171078 | 1.280275862 |
| SPAC6F12.03c | 1.449926621 | 1.28035465 |
| SPAC16E8.01 | 1.415379336 | 1.280394215 |
| SPBC17G9.12c | 1.219507856 | 1.28049828 |
| SPBC365.07c | 0.511711096 | 1.280997095 |
| SPAC922.06 | 1.222789937 | 1.281336765 |
| SPAC644.06c | 1.061598952 | 1.281382365 |
| SPAC23D3.12 | 0.524766493 | 1.282407733 |
| SPAC19B12.06c | 1.150579305 | 1.283022873 |
| SPAC8F11.08c | 1.072893487 | 1.283273834 |
| SPBC1711.06 | 1.412848156 | 1.283508747 |
| SPCC1235.15 | 1.364680073 | 1.283638025 |
| SPBC25H2.14 | 0.94043285 | 1.284042279 |
| SPCC569.04 | 1.326007135 | 1.284972193 |
| SPBC1289.01c | 1.092079791 | 1.285269171 |
| SPAC8F11.09c | 1.242688922 | 1.285310939 |
| SPAC521.02 | 1.21597482 | 1.285844293 |
| SPBC32F12.02 | 1.327050018 | 1.286648171 |
| SPBC1709.06 | 1.092938423 | 1.288867264 |
| SPCC364.04c | 1.366108583 | 1.289349882 |
| SPAC11E3.11c | 1.37904422 | 1.289416281 |
| SPCC63.08c | 2.258765522 | 1.289661705 |
| SPCC18B5.06 | 1.231554464 | 1.290087695 |
| SPCC1322.15 | 1.212485776 | 1.290805428 |
| SPAC212.01c | 1.1413592 | 1.291701911 |
| SPAC19E9.02 | 1.236440443 | 1.292354357 |
| SPCPB16A4.06c | 1.342608774 | 1.292571267 |
| SPBC21H7.03c | 1.212890769 | 1.29293338 |
| SPBC24C6.06 | 1.022324787 | 1.292945424 |
| SPAC4F10.07c | 1.307963872 | 1.292972029 |
| SPCC1393.08 | 1.329772613 | 1.293113837 |
| SPBC83.16c | 1.264226912 | 1.293231702 |
| SPAC664.01c | 1.292167407 | 1.293370789 |
| SPAC589.08c | 1.337460164 | 1.293614968 |
| SPBC1709.01 | 1.345271694 | 1.294011842 |
| SPBC800.05c | 0.468649937 | 1.294173756 |
| SPBC32H8.05 | 1.363632856 | 1.294295592 |

| | | |
|---|---|---|
| SPCC965.06 | 1.604129986 | 1.294572335 |
| SPAC13C5.04 | 1.359891687 | 1.296371051 |
| SPAC17H9.19c | 2.423423786 | 1.296723467 |
| SPAC15F9.02 | 1.295545377 | 1.297416959 |
| SPAC8F11.02c | 1.034901366 | 1.298036798 |
| SPBC24C6.08c | 1.300983271 | 1.298229868 |
| SPAC24H6.08 | 2.39402985 | 1.298507462 |
| SPAC323.03c | 1.21944584 | 1.298727632 |
| SPCC126.02c | 0.990931239 | 1.298996256 |
| SPAC110.02 | 0.918137758 | 1.299299562 |
| SPBC16A3.16 | 1.427629368 | 1.299536628 |
| SPAC1851.02 | 1.347987421 | 1.300534757 |
| SPCC1020.11c | 1.79902037 | 1.301215222 |
| SPBC2A9.11c | 1.149983405 | 1.301525421 |
| SPAC24C9.08 | 1.32654398 | 1.301777191 |
| SPCC1620.02 | 1.28520534 | 1.302193687 |
| SPAC823.12 | 0.2507 | 1.302337662 |
| SPBC8D2.12c | 0.82707951 | 1.30246198 |
| SPCC1739.10 | 1.41592636 | 1.303957884 |
| SPCC970.07c | 0.914225404 | 1.304141406 |
| SPBC18H10.07 | 1.414716024 | 1.304820318 |
| SPBC354.13 | 1.091591529 | 1.305242126 |
| SPCPB1C11.02 | 0.782044956 | 1.305288805 |
| SPAC30D11.13 | 1.148083334 | 1.306346255 |
| SPAC30D11.14c | 1.585092619 | 1.307427962 |
| SPBC19C7.09c | 1.050976798 | 1.307668755 |
| SPBC26H8.09c | 1.265626129 | 1.307959839 |
| SPCC895.05 | 1.368607748 | 1.30806811 |
| SPBC3B9.11c | 1.263998602 | 1.308967405 |
| SPCC1682.14 | 1.911238144 | 1.309005586 |
| SPBC19C7.04c | 1.066671374 | 1.309720746 |
| SPBC1215.01 | 1.533323886 | 1.310338167 |
| SPBC365.11 | 1.219815452 | 1.310654194 |
| SPAC3H5.09c | 1.438332371 | 1.311155849 |
| SPCC24B10.02c | 1.310624316 | 1.311473165 |
| SPCC1795.01c | 1.376588532 | 1.311946533 |
| SPCC1620.13 | 1.259969339 | 1.312552027 |
| SPAP14E8.05c | 1.420067008 | 1.313228013 |
| SPAC24H6.10c | 1.19008763 | 1.31400598 |
| SPBC119.14 | 1.330630985 | 1.314255956 |
| SPBC16A3.18 | 0.983570261 | 1.314566598 |
| SPBPB2B2.19c | 1.527883638 | 1.314864418 |
| SPAC1565.07c | 1.309880518 | 1.315108806 |
| SPAC6C3.08 | 1.259908219 | 1.316019186 |
| SPBC21D10.12 | 1.184326402 | 1.316209483 |
| SPBC2A9.02 | 1.401382913 | 1.316475876 |
| SPCC126.03 | 1.369800132 | 1.316584466 |
| SPAC23G3.03 | 1.397253283 | 1.316919725 |

| | | |
|---|---|---|
| SPCC550.07 | 1.277584925 | 1.31718511 |
| SPAC140.03 | 1.575735201 | 1.317200827 |
| SPAC890.03 | 1.452603395 | 1.317757878 |
| SPCC126.09 | 0.967460096 | 1.319850639 |
| SPCC1840.09 | 1.500713477 | 1.320593 |
| SPAC1B3.17 | 0.620727455 | 1.322117982 |
| SPCC61.03 | 1.286189661 | 1.323268191 |
| SPAC5D6.01 | 1.181979778 | 1.324114191 |
| SPAC4F10.16c | 0.795169126 | 1.324266617 |
| SPBC29A10.10c | 1.602292723 | 1.324787125 |
| SPAC20H4.04 | 1.215867855 | 1.325486124 |
| SPCP31B10.04 | 1.173422929 | 1.32604721 |
| SPBC2A9.07c | 1.109890376 | 1.326081166 |
| SPBC2D10.14c | 1.458823529 | 1.326807598 |
| SPBC16A3.19 | 0.918163924 | 1.327151996 |
| SPAC9E9.09c | 1.285957555 | 1.327373406 |
| SPBC18E5.05c | 1.273003048 | 1.329574142 |
| SPAP32A8.03c | 1.266187887 | 1.329933667 |
| SPBP8B7.22 | 1.28344146 | 1.330114348 |
| SPAC23A1.07 | 1.512012372 | 1.330343605 |
| SPCC16A11.08 | 1.213949201 | 1.330678592 |
| SPAP14E8.02 | 1.251231462 | 1.3307463 |
| SPBC17G9.02c | 1.176867114 | 1.331382673 |
| SPBC646.09c | 1.169285376 | 1.331400851 |
| SPBC947.09 | 0.882596627 | 1.331867603 |
| SPAC24C9.15c | 1.116247479 | 1.332020035 |
| SPAPB1E7.06c | 1.739036253 | 1.332597285 |
| SPAC1851.04c | 0.534905641 | 1.332736274 |
| SPCC1235.11 | 1.645889847 | 1.332821413 |
| SPAC25G10.05c | 1.290678295 | 1.33409585 |
| SPAC9.08c | 1.584241658 | 1.334725222 |
| SPCC1919.10c | 1.653006191 | 1.334794279 |
| SPAC6G10.12c | 1.794627786 | 1.334854665 |
| SPAC23A1.16c | 1.282668041 | 1.336142931 |
| SPAPB1E7.05 | 0.986847218 | 1.336822322 |
| SPCC548.05c | 1.448727954 | 1.336993289 |
| SPAC637.10c | 1.097020943 | 1.337809264 |
| SPBC29A10.07 | 1.30754909 | 1.338436264 |
| SPCC24B10.10c | 1.367048012 | 1.340782258 |
| SPBC1685.13 | 1.239110292 | 1.341033679 |
| SPAC6B12.14c | 1.183414954 | 1.341101691 |
| SPBC106.01 | 1.276840069 | 1.341108811 |
| SPBC1711.13 | 1.307469643 | 1.341461238 |
| SPBC947.01 | 1.452024935 | 1.342163498 |
| SPCC1281.04 | 1.305462306 | 1.342451119 |
| SPBC21B10.09 | 1.243450581 | 1.342588533 |
| SPBC1289.08 | 1.420833334 | 1.344597583 |
| SPBC21D10.11c | 1.263858822 | 1.346290034 |

| | | |
|---|---|---|
| SPBC947.08c | 1.259242355 | 1.346345847 |
| SPBC691.04 | 1.516735835 | 1.346495653 |
| SPCP1E11.06 | 1.258845042 | 1.346702495 |
| SPAC10F6.05c | 1.010920326 | 1.346726167 |
| SPAC19G12.15c | 2.201672892 | 1.347012638 |
| SPAC22E12.19 | 1.396517954 | 1.347184439 |
| SPAC3F10.04 | 1.321501737 | 1.348268914 |
| SPAC644.09 | 1.5099631 | 1.348862238 |
| SPAC513.02 | 1.366969549 | 1.34973884 |
| SPBC83.13 | 1.081990259 | 1.350587371 |
| SPCC613.02 | 1.490622414 | 1.351699485 |
| SPAC186.01 | 0.99965419 | 1.352400385 |
| SPAC2G11.03c | 1.368302304 | 1.352406121 |
| SPBC409.20c | 1.353862141 | 1.353779594 |
| SPBC1271.12 | 1.075874933 | 1.354192288 |
| SPAC664.02c | 1.308717261 | 1.354856959 |
| SPBC1709.10c | 1.157496837 | 1.354886639 |
| SPBC12C2.03c | 1.331199505 | 1.355219216 |
| SPAC20H4.03c | 1.676021505 | 1.357658769 |
| SPBP4H10.05c | 1.132704581 | 1.357663979 |
| SPAC26F1.05 | 1.493511111 | 1.357972222 |
| SPBC28F2.11 | 1.627758331 | 1.35816566 |
| SPCC285.16c | 1.47866498 | 1.360207003 |
| SPCC736.08 | 1.495779561 | 1.361342467 |
| SPCC1840.10 | 1.417226406 | 1.361919199 |
| SPBC1861.05 | 0.982185616 | 1.363000346 |
| SPBC11G11.03 | 1.509092792 | 1.363069309 |
| SPBC20F10.05 | 1.28398528 | 1.363319837 |
| SPAC1F3.03 | 0.952405255 | 1.364714017 |
| SPBC23G7.15c | 1.465478512 | 1.365176375 |
| SPBP4H10.14c | 1.018378857 | 1.365350877 |
| SPAC25B8.13c | 1.57718698 | 1.365790148 |
| SPCC74.03c | 1.585750897 | 1.365982189 |
| SPAC9.06c | 1.089823381 | 1.366353027 |
| SPCC1494.08c | 1.984317094 | 1.366891454 |
| SPAC186.05c | 1.037377417 | 1.368289156 |
| SPAC652.01 | 1.084476775 | 1.370210712 |
| SPBC1709.04c | 3.16075746 | 1.370255946 |
| SPCC4B3.17 | 1.929532669 | 1.370683432 |
| SPBC32H8.02c | 1.445655054 | 1.370916949 |
| SPBC21C3.01c | 1.523043478 | 1.370991848 |
| SPBC16H5.14c | 1.499306244 | 1.371035679 |
| SPAC694.05c | 1.434825426 | 1.371167189 |
| SPBC354.04 | 1.466949907 | 1.37137476 |
| SPAC1782.09c | 1.294483411 | 1.372743183 |
| SPAC22F3.10c | 1.178508858 | 1.374157208 |
| SPCC777.06c | 1.069350094 | 1.375329729 |
| SPBC21C3.07c | 1.818942505 | 1.375545431 |

| | | |
|---|---|---|
| SPAC16.03c | 1.678112806 | 1.377350125 |
| SPAC31G5.17c | 1.555668361 | 1.379339775 |
| SPBC106.08c | 1.074245394 | 1.379642787 |
| SPBC3B8.04c | 1.332291571 | 1.379703058 |
| SPCC24B10.08c | 1.580514857 | 1.380033793 |
| SPACUNK4.16c | 1.513089775 | 1.380215244 |
| SPAC15E1.04 | 1.305558912 | 1.380249244 |
| SPBC32H8.06 | 1.535309473 | 1.381122156 |
| SPAC823.10c | 1.086816195 | 1.381478822 |
| SPCC306.05c | 1.3525919 | 1.381736145 |
| SPAC56F8.16 | 1.561124665 | 1.382655603 |
| SPBC1652.02 | 1.340729302 | 1.384493429 |
| SPCC1919.07 | 1.950901287 | 1.384670353 |
| SPBC29A3.13 | 1.415829255 | 1.384674519 |
| SPAC4G8.03c | 1.450068587 | 1.384953704 |
| SPBP4H10.11c | 0.990360168 | 1.38501657 |
| SPAC27E2.03c | 1.388478709 | 1.385369282 |
| SPAC1071.04c | 0.947560975 | 1.38645362 |
| SPBC1709.19c | 1.192719162 | 1.386688921 |
| SPBC2D10.16 | 1.520395365 | 1.386811334 |
| SPCC330.19c | 1.370558118 | 1.388177713 |
| SPAC3C7.03c | 0.435067604 | 1.38827907 |
| SPBC651.09c | 1.081163515 | 1.388518876 |
| SPAC30C2.06c | 1.589141414 | 1.388541667 |
| SPAC23H3.09c | 1.490325842 | 1.388572335 |
| SPCC965.09 | 1.360023853 | 1.389322248 |
| SPCC16C4.10 | 1.770381011 | 1.389914725 |
| SPAC17A2.05 | 1.471000274 | 1.390299233 |
| SPBC14F5.10c | 1.292961556 | 1.390720076 |
| SPBC56F2.08c | 1.614210537 | 1.390820563 |
| SPBC1604.19c | 1.550576423 | 1.390979454 |
| SPBC337.11 | 1.111576638 | 1.391651106 |
| SPAC977.11 | 1.463940625 | 1.393615215 |
| SPBC1921.06c | 1.331578014 | 1.395505204 |
| SPAC7D4.13c | 1.387385433 | 1.395913636 |
| SPCC1840.07c | 1.186602605 | 1.396165288 |
| SPAC8C9.05 | 0.941698135 | 1.396552109 |
| SPAC4F10.02 | 1.552206926 | 1.396911397 |
| SPAC1093.02 | 1.165811015 | 1.397496656 |
| SPBC83.12 | 1.496213288 | 1.397524377 |
| SPCC794.11c | 1.610869118 | 1.398746597 |
| SPBC83.09c | 1.064847512 | 1.399478331 |
| SPAC31G5.21 | 1.38514768 | 1.400159132 |
| SPAC29A4.16 | 1.566696589 | 1.40094816 |
| SPCC613.06 | 1.668040414 | 1.403036376 |
| SPCC162.11c | 1.53470852 | 1.40316704 |
| SPCC18.10 | 1.449744681 | 1.404893617 |
| SPBP16F5.05c | 1.595380355 | 1.405564424 |

| | | |
|---|---|---|
| SPBC215.01 | 1.429194068 | 1.405984658 |
| SPCC663.11 | 1.674112039 | 1.407396588 |
| SPBC32F12.01c | 1.362294298 | 1.407397713 |
| SPBC13G1.03c | 1.481545741 | 1.408049093 |
| SPCC794.02 | 1.597836089 | 1.408212422 |
| SPAC144.06 | 1.642529511 | 1.409506745 |
| SPAC6G9.12 | 1.327436643 | 1.409770222 |
| SPCC777.13 | 1.189373602 | 1.409780481 |
| SPAC23H3.05c | 1.80610687 | 1.410152672 |
| SPAC222.07c | 1.003521863 | 1.41121448 |
| SPAC57A7.09 | 1.666196552 | 1.411923421 |
| SPBC1711.04 | 1.523895222 | 1.413342985 |
| SPBC1778.02 | 1.475831453 | 1.413351739 |
| SPBC359.06 | 1.440731001 | 1.413621445 |
| SPCC1442.13c | 1.350107514 | 1.414534798 |
| SPBC839.04 | 1.633882353 | 1.415514706 |
| SPBC1539.04 | 0.972146139 | 1.415524915 |
| SPAC16.04 | 1.531224729 | 1.416040604 |
| SPAC6F12.04 | 1.179444481 | 1.416465657 |
| SPBC19C7.10 | 1.488396026 | 1.417028921 |
| SPCC1620.03 | 1.395597622 | 1.417614783 |
| SPAC3H5.12c | 1.475275077 | 1.418687963 |
| SPBC21C3.19 | 1.245872461 | 1.419312441 |
| SPAC57A10.07 | 1.207652589 | 1.419798557 |
| SPAC27D7.05c | 1.268987818 | 1.420683014 |
| SPAC222.13c | 1.345850561 | 1.421657219 |
| SPCC895.07 | 1.347527978 | 1.421743596 |
| SPAC1783.05 | 1.384420936 | 1.421815713 |
| SPAPJ760.03c | 1.103077657 | 1.42228143 |
| SPBC902.03 | 1.29617772 | 1.422829593 |
| SPAC7D4.05 | 1.301526243 | 1.423405324 |
| SPAC17G6.05c | 1.506448172 | 1.424682875 |
| SPAC19A8.05c | 1.301254316 | 1.425414321 |
| SPAC1834.04 | 1.619 | 1.4259375 |
| SPBC24C6.10c | 1.561507204 | 1.426525823 |
| SPAC30D11.05 | 1.211652105 | 1.427191567 |
| SPBC19C2.09 | 1.37912117 | 1.428144201 |
| SPAC25G10.03 | 1.595732507 | 1.429033382 |
| SPAC22F3.08c | 1.319447134 | 1.429710158 |
| SPCC1235.02 | 1.265178571 | 1.430454 |
| SPBC19C2.13c | 1.4608555 | 1.431187096 |
| SPBC32H8.07 | 1.384369214 | 1.432239925 |
| SPAC29A4.19c | 1.28105166 | 1.433105102 |
| SPBC336.13c | 1.467721252 | 1.433272571 |
| SPBC14C8.16c | 1.687542662 | 1.433287116 |
| SPAC20G8.02 | 1.415096851 | 1.433417458 |
| SPBC6B1.04 | 1.468607921 | 1.433817383 |
| SPAC2G11.12 | 1.596429737 | 1.434574352 |

| | | |
|---|---|---|
| SPAC1296.05c | 1.382129964 | 1.434764215 |
| SPBC17A3.05c | 2.202219357 | 1.434783228 |
| SPAC23C4.06c | 1.593946726 | 1.434992769 |
| SPCC1259.07 | 1.753404706 | 1.436170213 |
| SPAC17C9.05c | 1.651882971 | 1.436635937 |
| SPAC6B12.05c | 1.691921799 | 1.436965123 |
| SPAP8A3.12c | 1.531957238 | 1.437022823 |
| SPAC23C4.12 | 1.260571832 | 1.437224232 |
| SPBC3D6.02 | 1.157349708 | 1.437279807 |
| SPBC11B10.10c | 1.78782669 | 1.437337475 |
| SPCC4E9.01c | 1.140353468 | 1.438006631 |
| SPBC725.11c | 1.773610773 | 1.439591899 |
| SPBC19G7.07c | 1.277721867 | 1.43983017 |
| SPCP1E11.07c | 1.51955478 | 1.441073767 |
| SPAC1782.02c | 1.45766655 | 1.443393109 |
| SPAC4F10.13c | 1.373923754 | 1.443735507 |
| SPAC24H6.11c | 1.175990854 | 1.445312501 |
| SPAC17A2.02c | 1.894482011 | 1.445945791 |
| SPAPJ698.02c | 1.505482929 | 1.446745079 |
| SPBC1105.14 | 1.346156965 | 1.447588137 |
| SPBC28F2.03 | 1.401599458 | 1.448203148 |
| SPCC1393.13 | 1.494239184 | 1.448612091 |
| SPBC16E9.12c | 1.134043032 | 1.449335557 |
| SPAC29B12.06c | 1.411479221 | 1.449901613 |
| SPAC17A2.10c | 1.421879764 | 1.4510061 |
| SPBC725.02 | 1.419437584 | 1.451171713 |
| SPBC21C3.09c | 1.488519199 | 1.452079113 |
| SPBC26H8.01 | 1.41148777 | 1.45385753 |
| SPAC823.09c | 1.172221969 | 1.453866705 |
| SPBC16A3.10 | 1.206322983 | 1.454811413 |
| SPBC83.03c | 1.815628726 | 1.455066741 |
| SPBC337.13c | 1.317909658 | 1.456771776 |
| SPAC1D4.09c | 1.716518143 | 1.458867647 |
| SPBC800.02 | 1.204934956 | 1.459547068 |
| SPAPB1A10.08 | 1.410897436 | 1.459700226 |
| SPAC8C9.17c | 1.454748603 | 1.459890014 |
| SPCC63.02c | 0.539337341 | 1.460752524 |
| SPAC959.04c | 1.543741589 | 1.461599933 |
| SPBC25B2.04c | 1.628665719 | 1.461766016 |
| SPCC1223.11 | 1.326677669 | 1.462539146 |
| SPAC186.09 | 1.110681245 | 1.462902123 |
| SPBC2F12.11c | 1.443310768 | 1.464328435 |
| SPCC569.03 | 1.525139474 | 1.464674565 |
| SPAC11G7.03 | 1.598150568 | 1.466166535 |
| SPCC4F11.03c | 0.438628342 | 1.467078239 |
| SPBC3E7.11c | 1.275360367 | 1.467657398 |
| SPAC1071.11 | 1.166813325 | 1.469772973 |
| SPBC19G7.03c | 1.446634562 | 1.47024971 |

| | | |
|---|---|---|
| SPBC1539.02 | 1.358727209 | 1.470761147 |
| SPAC3H8.04 | 1.273671264 | 1.470797631 |
| SPAPB18E9.01 | 1.518202765 | 1.471661666 |
| SPCC645.08c | 1.827377332 | 1.472650908 |
| SPBC2A9.06c | 1.693385754 | 1.477321346 |
| SPAC23H4.08 | 1.184821041 | 1.480144815 |
| SPAC4A8.14 | 1.367769743 | 1.480349366 |
| SPAC890.02c | 0.932169287 | 1.480408658 |
| SPAC1635.01 | 1.603615812 | 1.480685746 |
| SPCC18.09c | 1.418941856 | 1.480710167 |
| SPCC70.08c | 1.815330882 | 1.481506239 |
| SPBC428.06c | 1.959259836 | 1.48225136 |
| SPBC106.17c | 1.212274296 | 1.482772702 |
| SPBC1709.11c | 1.121725732 | 1.482921417 |
| SPBP8B7.13 | 1.623659454 | 1.483516483 |
| SPCC126.08c | 1.340858223 | 1.485564716 |
| SPAC56F8.02 | 1.261111789 | 1.486122883 |
| SPBC1773.15 | 1.351458856 | 1.486159324 |
| SPAC23G3.04 | 1.788668562 | 1.48642164 |
| SPAC16E8.14c | 1.709311054 | 1.48885811 |
| SPAC2C4.14c | 1.427648323 | 1.489037709 |
| SPBC36B7.06c | 1.716789898 | 1.490888809 |
| SPCC1235.09 | 1.245798114 | 1.491836253 |
| SPAC4F10.20 | 1.810753564 | 1.493966395 |
| SPAC7D4.06c | 1.421767142 | 1.495467998 |
| SPBC25B2.11 | 1.59459513 | 1.496989734 |
| SPBC21C3.20c | 1.686833615 | 1.497976085 |
| SPAPB17E12.05 | 1.52464019 | 1.500445445 |
| SPCC338.14 | 1.521131124 | 1.502613621 |
| SPBC119.06 | 1.617160164 | 1.502805064 |
| SPBC1921.04c | 1.37682732 | 1.50351118 |
| SPAC4A8.10 | 1.11213278 | 1.504186811 |
| SPAC1952.03 | 1.810853965 | 1.504723222 |
| SPBC1861.09 | 1.707910601 | 1.507588423 |
| SPBC19C2.14 | 1.512195122 | 1.508173326 |
| SPBC32F12.07c | 1.455001753 | 1.508933751 |
| SPBC3B9.09 | 0.798355398 | 1.509035138 |
| SPBC1778.06c | 1.545749128 | 1.510574229 |
| SPBC215.03c | 1.703460591 | 1.510886486 |
| SPAC5D6.06c | 1.780506431 | 1.512698453 |
| SPAC13C5.07 | 1.743055884 | 1.514893507 |
| SPAC25H1.05 | 1.985852671 | 1.515922282 |
| SPBC409.17c | 1.87843874 | 1.516814617 |
| SPCP31B10.03c | 1.995603446 | 1.51936445 |
| SPBC405.02c | 1.613566503 | 1.519595018 |
| SPAC1805.05 | 1.74904199 | 1.520064717 |
| SPBC1685.08 | 1.881035714 | 1.520591518 |
| SPAC11D3.09 | 1.488322015 | 1.52194992 |

| | | |
|---|---|---|
| SPBC365.16 | 2.019248093 | 1.522181818 |
| SPAPB1A10.14 | 1.464616665 | 1.523493606 |
| SPAC644.07 | 1.321845619 | 1.523898662 |
| SPBC3B8.08 | 1.667691724 | 1.524843394 |
| SPAC11G7.06c | 2.319890109 | 1.52657967 |
| SPBC23G7.12c | 1.642386579 | 1.528827382 |
| SPBC317.01 | 1.882048861 | 1.530455275 |
| SPCC1259.12c | 1.351271132 | 1.530629041 |
| SPAC1556.01c | 1.902785618 | 1.531322993 |
| SPAC17A5.08 | 1.183910139 | 1.531648452 |
| SPAC19D5.11c | 1.740241358 | 1.531661446 |
| SPCC777.02 | 1.585409683 | 1.532773512 |
| SPCC553.01c | 1.603670332 | 1.533554996 |
| SPAC12G12.03 | 1.396761503 | 1.53861548 |
| SPAPB1A11.03 | 1.955401564 | 1.539878731 |
| SPBC1734.09 | 1.559963178 | 1.542105348 |
| SPAC1834.08 | 1.700889465 | 1.542555084 |
| SPAC24H6.03 | 1.497058324 | 1.542556575 |
| SPBC713.07c | 0.575208093 | 1.547877573 |
| SPCC1739.07 | 1.371939239 | 1.550095749 |
| SPAC1F3.10c | 1.678225461 | 1.552159237 |
| SPAC2F3.08 | 1.682281138 | 1.552606332 |
| SPAC57A7.04c | 1.835316781 | 1.553898223 |
| SPAC1687.17c | 1.800446761 | 1.55483293 |
| SPAC694.04c | 1.355034247 | 1.556173279 |
| SPAC23C4.09c | 1.574520363 | 1.558282144 |
| SPBC1289.06c | 1.866953536 | 1.561252652 |
| SPBC15D4.05 | 1.374796931 | 1.562153901 |
| SPAC23C4.03 | 1.339124467 | 1.562917843 |
| SPBC21C3.02c | 0.951138493 | 1.563681872 |
| SPBC1718.03 | 1.855291015 | 1.564051638 |
| SPAC1F7.01c | 0.912781955 | 1.564088271 |
| SPAC4F10.04 | 2.000297214 | 1.564221957 |
| SPBC83.05 | 1.643756312 | 1.565902236 |
| SPBC24C6.05 | 0.904436237 | 1.570493932 |
| SPAC227.17c | 2.048723186 | 1.573352911 |
| SPAC664.13 | 1.378065 | 1.574750769 |
| SPAC18G6.10 | 1.211217413 | 1.575374503 |
| SPBC1861.01c | 2.148045959 | 1.576911665 |
| SPAC1805.14 | 1.450395055 | 1.577695273 |
| SPAC1142.02c | 1.630118212 | 1.578478188 |
| SPAC20G8.09c | 1.761095958 | 1.579551249 |
| SPAC8E11.02c | 1.599099629 | 1.581466259 |
| SPBC29A3.11c | 1.313800784 | 1.581846528 |
| SPCC4B3.12 | 1.423342437 | 1.581907444 |
| SPAC4A8.03c | 1.146240255 | 1.582556658 |
| SPBC26H8.03 | 1.193832391 | 1.584602951 |
| SPAC14C4.13 | 1.759214182 | 1.585863729 |

| | | |
|---|---|---|
| SPBC215.02 | 1.516350607 | 1.586166789 |
| SPBC29A3.03c | 1.609609573 | 1.591428356 |
| SPBC409.16c | 1.57260514 | 1.593745196 |
| SPAC6F6.03c | 1.537211923 | 1.596371601 |
| SPCC895.09c | 1.432484234 | 1.596392643 |
| SPAC1F5.05c | 1.532259331 | 1.600704316 |
| SPBC887.04c | 1.602675292 | 1.601172058 |
| SPAC959.08 | 1.814939356 | 1.604573073 |
| SPBC11G11.05 | 1.664724733 | 1.604755273 |
| SPAC513.03 | 1.109763314 | 1.605029585 |
| SPAC22A12.03c | 1.433746985 | 1.608266874 |
| SPAC11G7.02 | 1.082794205 | 1.613775576 |
| SPAC11E3.04c | 1.168360902 | 1.61588784 |
| SPBC3D6.09 | 1.741708451 | 1.616377197 |
| SPAC29B12.08 | 1.847864623 | 1.618562516 |
| SPBC530.11c | 1.779686688 | 1.620683517 |
| SPBC1773.12 | 2.053367653 | 1.622064028 |
| SPBP4H10.20 | 1.693237533 | 1.624750489 |
| SPBC15D4.01c | 1.918839005 | 1.62524641 |
| SPCC23B6.04c | 0.719663594 | 1.626925914 |
| SPCC895.06 | 0.906294118 | 1.6275 |
| SPAC15A10.11 | 1.752106857 | 1.62758731 |
| SPBC146.13c | 2.638954051 | 1.630245111 |
| SPBC30B4.06c | 1.942742982 | 1.631049454 |
| SPBC4F6.08c | 1.799070681 | 1.63646769 |
| SPBC6B1.05c | 1.405125281 | 1.636469417 |
| SPCC11E10.06c | 1.913085081 | 1.637345812 |
| SPBC2G2.08 | 2.128775039 | 1.638306909 |
| SPAC1B3.02c | 1.799808748 | 1.641398053 |
| SPBC1198.06c | 1.550486674 | 1.642727765 |
| SPBC29A10.16c | 1.883265811 | 1.645686823 |
| SPBC649.04 | 2.265850975 | 1.647202657 |
| SPAC2F7.17 | 1.923195592 | 1.64927686 |
| SPBC18E5.04 | 1.889110889 | 1.650552572 |
| SPACUNK4.10 | 1.995978054 | 1.654529673 |
| SPBC106.16 | 1.607712766 | 1.656665558 |
| SPAC1805.15c | 1.013561083 | 1.657723062 |
| SPAC22F3.13 | 1.68734197 | 1.659986577 |
| SPBC1734.08 | 1.940079894 | 1.661451398 |
| SPAC13C5.02 | 1.808228956 | 1.662695966 |
| SPBC11G11.01 | 1.511560153 | 1.663370523 |
| SPBC216.01c | 1.663931935 | 1.6680102 |
| SPCC162.12 | 2.459571245 | 1.669420195 |
| SPCC1020.09 | 1.952531203 | 1.672128172 |
| SPBC3H7.10 | 1.969570654 | 1.672276971 |
| SPBC3D6.08c | 1.962852228 | 1.672843666 |
| SPBC365.20c | 1.67806595 | 1.67548166 |
| SPAPB1A10.03 | 1.732906339 | 1.677647727 |

| | | |
|---|---|---|
| SPAC6B12.08 | 1.587974676 | 1.684989791 |
| SPAC19G12.08 | 1.789312356 | 1.686210545 |
| SPBC119.04 | 1.792815543 | 1.687578294 |
| SPAC767.01c | 1.533044083 | 1.687679237 |
| SPAC11H11.01 | 2.09427975 | 1.687930584 |
| SPAC26F1.12c | 1.776308446 | 1.702322141 |
| SPAC167.07c | 1.750661147 | 1.704726268 |
| SPAP8A3.07c | 1.571534193 | 1.706463896 |
| SPAC16C9.04c | 1.87034291 | 1.707657554 |
| SPBC1271.07c | 1.979181909 | 1.712557424 |
| SPBC30D10.04 | 1.880876403 | 1.713406773 |
| SPAPB1A10.13 | 1.809226186 | 1.714820182 |
| SPAC227.14 | 1.19741351 | 1.715516019 |
| SPCC188.08c | 2.676559147 | 1.718984216 |
| SPBC29A3.08 | 1.929120299 | 1.71921913 |
| SPBC29A3.09c | 1.8084229 | 1.719393846 |
| SPBC29A3.18 | 1.781372249 | 1.721002796 |
| SPAC31A2.14 | 1.443706208 | 1.721928182 |
| SPBC24C6.04 | 1.63274219 | 1.723041601 |
| SPAC1B3.07c | 1.951384997 | 1.724049021 |
| SPAC8C9.09c | 1.80149905 | 1.724637264 |
| SPAC869.11 | 1.118108273 | 1.729144636 |
| SPCC594.05c | 1.356714199 | 1.730715819 |
| SPCC338.10c | 2.342875432 | 1.730860832 |
| SPAC22F3.09c | 1.782297298 | 1.732284233 |
| SPAC29B12.03 | 1.225421612 | 1.732916423 |
| SPAC22E12.05c | 1.510668489 | 1.736295431 |
| SPCC1840.06 | 1.69740634 | 1.736491354 |
| SPAC869.01 | 1.476944122 | 1.737465851 |
| SPAC5H10.01 | 1.578629793 | 1.738767567 |
| SPAC5D6.13 | 2.101879373 | 1.739306411 |
| SPAC328.10c | 2.031923792 | 1.739984438 |
| SPAC19G12.13c | 2.021818182 | 1.742045455 |
| SPCC191.07 | 2.012993189 | 1.744623444 |
| SPAC1639.02c | 1.566909848 | 1.747426253 |
| SPBP22H7.08 | 1.943105191 | 1.749027929 |
| SPAC664.15 | 1.784541114 | 1.750154607 |
| SPCC1259.02c | 1.904381779 | 1.75635846 |
| SPAC9.13c | 1.578333563 | 1.759843156 |
| SPAC664.03 | 1.580284746 | 1.761087842 |
| SPAC22F8.12c | 1.880249999 | 1.761616161 |
| SPCC1322.01 | 2.053884216 | 1.763019545 |
| SPBC13E7.04 | 1.451522722 | 1.773451993 |
| SPAC3H8.09c | 1.741500336 | 1.774663834 |
| SPCC1742.01 | 1.752321853 | 1.777205627 |
| SPCC306.04c | 1.659783976 | 1.779215115 |
| SPBC1683.09c | 1.938137072 | 1.780833007 |
| SPAP27G11.14c | 2.592998776 | 1.785515789 |

| | | |
|---|---|---|
| SPCC594.06c | 1.489897904 | 1.78790969 |
| SPBC405.06 | 1.482284744 | 1.788369234 |
| SPAC3A11.07 | 2.139245868 | 1.794862291 |
| SPAC637.11 | 0.92916727 | 1.803302316 |
| SPAPB1A10.10c | 1.6092407 | 1.806702853 |
| SPAC1D4.06c | 2.269075414 | 1.807743149 |
| SPAC25A8.01c | 1.748453569 | 1.810784122 |
| SPAC1F5.07c | 2.121639382 | 1.811478205 |
| SPCC1919.03c | 2.056170213 | 1.814834077 |
| SPBPJ4664.01 | 1.950192678 | 1.817106214 |
| SPBC2G2.03c | 2.292462985 | 1.822095547 |
| SPBC36.03c | 1.653858007 | 1.822211283 |
| SPAC637.09 | 1.674680963 | 1.824862402 |
| SPAC6C3.04 | 1.913414038 | 1.831025587 |
| SPBC18H10.04c | 1.812765849 | 1.831036382 |
| SPCC550.12 | 1.587034302 | 1.846134091 |
| SPCC794.03 | 1.646861192 | 1.850419777 |
| SPAC3A11.13 | 1.787676688 | 1.850521113 |
| SPAPB1E7.11c | 2.276581367 | 1.869001017 |
| SPBC11C11.07 | 1.278028173 | 1.880536831 |
| SPAC144.08 | 2.187753303 | 1.883301211 |
| SPAC23H4.12 | 2.072556257 | 1.887554408 |
| SPBC19C7.01 | 2.272119144 | 1.889672399 |
| SPAC1610.02c | 2.286335957 | 1.90836104 |
| SPBC354.03 | 1.56276946 | 1.916766017 |
| SPBC1604.11 | 2.471142857 | 1.945803571 |
| SPAC22H10.11c | 2.198735804 | 1.947612302 |
| SPAC167.05 | 1.924154964 | 1.950533522 |
| SPAC26A3.11 | 1.885907145 | 1.952044269 |
| SPAC2C4.15c | 1.322002876 | 1.955634068 |
| SPAC16E8.05c | 2.444109937 | 1.966923636 |
| SPBC13G1.08c | 1.546851863 | 1.984686958 |
| SPCC1672.06c | 2.243125905 | 1.989372287 |
| SPBC12C2.08 | 1.244838521 | 1.996087592 |
| SPBC19G7.16 | 2.121497172 | 2.014408611 |
| SPCC584.11c | 1.94129325 | 2.01546479 |
| SPBC13G1.12 | 2.308283271 | 2.026806988 |
| SPAC23C11.10 | 2.274560716 | 2.029324165 |
| SPCC188.02 | 2.00253825 | 2.029968221 |
| SPBC1604.02c | 1.544759476 | 2.042805013 |
| SPBC29A3.07c | 1.910187524 | 2.068278181 |
| SPAC1D4.01 | 1.263829302 | 2.072698494 |
| SPAC4G8.11c | 3.456717174 | 2.076980037 |
| SPCC297.05 | 1.922468589 | 2.080223516 |
| SPAC5D6.12 | 2.617591339 | 2.092185385 |
| SPCC320.12 | 1.694603131 | 2.110211947 |
| SPAC3H5.07 | 2.253524935 | 2.114454357 |
| SPBC4B4.03 | 1.974323249 | 2.136713473 |

| | | |
|---|---|---|
| SPAC17G8.13c | 1.935466796 | 2.159798876 |
| SPAC30D11.11 | 2.182098899 | 2.179056779 |
| SPBC15D4.10c | 4.325390307 | 2.182473736 |
| SPAC4D7.11 | 2.431443334 | 2.20489006 |
| SPBC725.10 | 1.995784872 | 2.211370898 |
| SPAC1039.08 | 2.691932057 | 2.221337575 |
| SPBC12C2.01c | 4.313404756 | 2.224898876 |
| SPAC25G10.06 | 2.529225176 | 2.228262174 |
| SPBC15C4.04c | 1.797536291 | 2.241281799 |
| SPBC3B8.02 | 1.901659186 | 2.275576113 |
| SPBC543.07 | 2.404097561 | 2.281666666 |
| SPBC577.02 | 2.754907503 | 2.30500251 |
| SPCC1183.10 | 2.59703388 | 2.310067805 |
| SPAC22H12.02 | 1.881765169 | 2.313331722 |
| SPBC713.08 | 2.448492646 | 2.315359476 |
| SPAC9G1.12 | 3.024118458 | 2.330765181 |
| SPCC1393.02c | 2.170137257 | 2.378331364 |
| SPCC794.08 | 2.148729509 | 2.391457758 |
| SPCC16A11.10c | 2.428381375 | 2.394854363 |
| SPBC16A3.07c | 2.861920688 | 2.426531592 |
| SPAC31G5.03 | 2.422781883 | 2.454813515 |
| SPAC10F6.13c | 4.072070949 | 2.468569521 |
| SPAC328.02 | 2.968244668 | 2.469288719 |
| SPAC13G6.09 | 2.964713004 | 2.504571557 |
| SPCC11E10.04 | 1.585992106 | 2.589870741 |
| SPAC18G6.09c | 2.075486473 | 2.688439209 |
| SPBC21B10.13c | 3.392907797 | 2.711779446 |
| SPBC409.19c | 3.258490718 | 2.717158064 |
| SPAC222.05c | 3.186584294 | 2.727913201 |
| SPBC26H8.14c | 2.575381197 | 2.785898992 |
| SPBC3H7.09 | 2.545939468 | 2.790223539 |
| SPBC32H8.03 | 2.298578904 | 2.902379765 |
| SPBC776.09 | 2.371413827 | 2.941755957 |
| SPCC736.07c | 1.962632659 | 2.951976692 |
| SPAC1486.08 | 2.599652064 | 3.000563605 |
| SPAC16A10.05c | 3.723976674 | 3.081982206 |
| SPAC26F1.03 | 2.407731873 | 3.20131803 |
| SPCC4G3.04c | 4.103009538 | 3.269707983 |
| SPCC794.07 | 2.821100919 | 3.339622644 |
| SPBC16A3.03c | 4.26357107 | 3.538338349 |
| SPBC839.03c | 4.319926471 | 3.979596383 |
| SPAC3G6.02 | 0.948876728 | 5.617029284 |
