## Supplemental Table 7 for "Rapalink-1 reveals novel mTOR-dependent genes and an agmatinergic axis-based metabolic feedback regulating mTOR activity and lifespan"

| Systematic gene name | *agm1* interaction value |
|---|---|
| SPAC17A2.09c | -2 |
| SPAC824.07 | -1.925437511 |
| SPAC683.03 | -1.802254236 |
| SPAC2F3.15 | -1.736206329 |
| SPBC30D10.13c | -1.734351688 |
| SPBPJ4664.02 | -1.56315012 |
| SPCC970.10c | -1.556528371 |
| SPAC23A1.11 | -1.522140221 |
| SPAC5D6.01 | -1.47704468 |
| SPAC6G10.12c | -1.41067768 |
| SPBP4H10.09 | -1.293537046 |
| SPBC21C3.08c | -1.267179492 |
| SPBP8B7.22 | -1.185660438 |
| SPAC212.03 | -1.179457746 |
| SPBC651.02 | -1.065402297 |
| SPAC15A10.03c | -0.991025825 |
| SPAC144.02 | -0.969322733 |
| SPAC10F6.16 | -0.968303647 |
| SPAC5D6.12 | -0.957580109 |
| SPBPJ4664.01 | -0.83703687 |
| SPBC1604.08c | -0.836524642 |
| SPAC227.13c | -0.817893336 |
| SPAC227.17c | -0.814607767 |
| SPAC2F7.10 | -0.770654166 |
| SPCC1840.06 | -0.767059783 |
| SPBC21B10.13c | -0.675890116 |
| SPBC26H8.14c | -0.674407152 |
| SPBC577.02 | -0.649232079 |
| SPCC1442.14c | -0.600583695 |
| SPAC343.04c | -0.594504157 |
| SPBC1105.10 | -0.582917626 |
| SPAC26H5.02c | -0.572903498 |
| SPAC16C9.06c | -0.571230025 |
| SPAC17D4.03c | -0.549099384 |
| SPBC16E9.07 | -0.544004861 |
| SPAC16C9.05 | -0.537875936 |
| SPBC3D6.04c | -0.535863676 |
| SPBC29A10.12 | -0.530520955 |
| SPAC15A10.11 | -0.529295257 |
| SPBPB8B6.05c | -0.523575803 |
| SPAC1851.02 | -0.503019869 |
| SPAC1486.08 | -0.493490853 |
| SPCC191.07 | -0.487528689 |
| SPAC824.04 | -0.473994275 |
| SPAC8C9.14 | -0.473500823 |
| SPAPB1E7.05 | -0.470614337 |

| | |
|---|---|
| SPAC23C11.14 | -0.450276509 |
| SPBC4C3.12 | -0.441805866 |
| SPAC8E11.07c | -0.43340922 |
| SPBC26H8.12 | -0.432125384 |
| SPAC31A2.16 | -0.430260516 |
| SPAC23C11.10 | -0.422411078 |
| SPCC1682.08c | -0.419832093 |
| SPAC1142.07c | -0.417292175 |
| SPBC36.07 | -0.406340922 |
| SPAC8C9.19 | -0.396550277 |
| SPBC23G7.12c | -0.392319118 |
| SPBC29A3.18 | -0.387507679 |
| SPAC24B11.08c | -0.384950084 |
| SPAC9G1.02 | -0.382651166 |
| SPBC2D10.17 | -0.374103928 |
| SPBC4B4.10c | -0.366964304 |
| SPCC320.12 | -0.362349841 |
| SPAC1805.12c | -0.360988407 |
| SPCC162.12 | -0.357943725 |
| SPBC119.06 | -0.357882278 |
| SPAC2G11.05c | -0.356364076 |
| SPAC222.07c | -0.349108387 |
| SPBP16F5.05c | -0.338868283 |
| SPBC354.12 | -0.337438849 |
| SPAC8E11.05c | -0.3374016 |
| SPBC1539.06 | -0.332879435 |
| SPBC713.07c | -0.328372622 |
| SPAC630.13c | -0.31899287 |
| SPAC5D6.08c | -0.317737175 |
| SPAC630.15 | -0.31057621 |
| SPAC3F10.02c | -0.30808781 |
| SPAC3C7.08c | -0.307082176 |
| SPCC576.12c | -0.306843001 |
| SPBC23G7.13c | -0.306032948 |
| SPAC22H10.07 | -0.304213061 |
| SPAC4H3.03c | -0.301756626 |
| SPBC16G5.05c | -0.295171362 |
| SPAC11G7.02 | -0.295141612 |
| SPBC3H7.03c | -0.292007147 |
| SPAC27D7.02c | -0.287454759 |
| SPBC19C7.02 | -0.286199642 |
| SPAC19A8.11c | -0.282781839 |
| SPBC1271.15c | -0.278747114 |
| SPAC1F8.02c | -0.274989889 |
| SPBC365.10 | -0.272798527 |
| SPBC1306.02 | -0.264252873 |
| SPAC27F1.05c | -0.263617306 |
| SPAC167.05 | -0.258114157 |

| | |
|---|---|
| SPAC1486.04c | -0.255636919 |
| SPAC13G6.15c | -0.251822823 |
| SPBC23E6.09 | -0.250228195 |
| SPAC57A7.04c | -0.246608572 |
| SPCC1739.14 | -0.245639393 |
| SPAC1B3.17 | -0.242789328 |
| SPCC594.02c | -0.242292281 |
| SPBP23A10.14c | -0.235169184 |
| SPCC736.08 | -0.232521714 |
| SPAC12B10.07 | -0.230708646 |
| SPAC2F7.03c | -0.230183702 |
| SPAC959.04c | -0.228935062 |
| SPAC1751.04 | -0.2260781 |
| SPBC1198.01 | -0.225197945 |
| SPBC106.16 | -0.223168345 |
| SPAC4F10.02 | -0.222378359 |
| SPBC19F8.08 | -0.220357326 |
| SPCC1672.06c | -0.216673295 |
| SPBC16D10.08c | -0.214222882 |
| SPAC630.09c | -0.212798325 |
| SPCC1235.05c | -0.209672012 |
| SPAC13G6.03 | -0.209016621 |
| SPAC29E6.10c | -0.20634714 |
| SPCC777.09c | -0.205240679 |
| SPCC74.02c | -0.204519315 |
| SPBC409.19c | -0.203701648 |
| SPAC1782.11 | -0.201615502 |
| SPAC26F1.09 | -0.201612382 |
| SPAC2F7.17 | -0.201511633 |
| SPBC3D6.09 | -0.199822723 |
| SPCC1450.11c | -0.199022601 |
| SPBC1683.07 | -0.198781078 |
| SPAC4G8.13c | -0.19612139 |
| SPBC15D4.06 | -0.195306544 |
| SPAC14C4.06c | -0.19317666 |
| SPBC428.08c | -0.18951501 |
| SPBC106.17c | -0.189347154 |
| SPCC126.09 | -0.189188275 |
| SPAC5H10.07 | -0.18855875 |
| SPAC821.11 | -0.187397125 |
| SPAC30C2.06c | -0.18675464 |
| SPBC21C3.02c | -0.18637856 |
| SPAC1F3.09 | -0.185705586 |
| SPCC1223.13 | -0.185351377 |
| SPAC24B11.10c | -0.185051671 |
| SPAC11G7.03 | -0.181111684 |
| SPCC1682.01 | -0.180734458 |
| SPAC19E9.01c | -0.180119448 |

| | |
|---|---|
| SPAC3G9.07c | -0.179520706 |
| SPAC29A4.17c | -0.179501472 |
| SPAC19B12.10 | -0.179024357 |
| SPCC794.01c | -0.178387611 |
| SPAC13F5.07c | -0.177078446 |
| SPBC21H7.04 | -0.177039917 |
| SPBP4H10.08 | -0.175282712 |
| SPAC1039.08 | -0.175072433 |
| SPBC106.12c | -0.170360837 |
| SPAC4G8.11c | -0.169715676 |
| SPAC12G12.15 | -0.169615448 |
| SPBC14C8.16c | -0.169200514 |
| SPAC4G9.13c | -0.168305852 |
| SPAC30.02c | -0.168108207 |
| SPAC30D11.07 | -0.167718575 |
| SPAC977.11 | -0.16655261 |
| SPAC343.20 | -0.165345556 |
| SPAPB21F2.03 | -0.164013089 |
| SPAC5H10.09c | -0.163039609 |
| SPAC1705.02 | -0.162646087 |
| SPBC2G2.13c | -0.161819767 |
| SPBC800.08 | -0.160988905 |
| SPCC1223.05c | -0.159188918 |
| SPBC18E5.04 | -0.159073743 |
| SPBC4F6.16c | -0.158927383 |
| SPAC110.02 | -0.158302884 |
| SPAC11E3.04c | -0.156420152 |
| SPAC6B12.16 | -0.156209538 |
| SPBC16C6.09 | -0.155984549 |
| SPAPB1A11.03 | -0.15560037 |
| SPCC4F11.04c | -0.155346686 |
| SPCC737.06c | -0.153451822 |
| SPAC167.04 | -0.152007229 |
| SPBC577.12 | -0.151968865 |
| SPCC23B6.01c | -0.151276944 |
| SPAC750.08c | -0.151177772 |
| SPBC887.10 | -0.150642112 |
| SPBC18E5.08 | -0.149432177 |
| SPAC977.10 | -0.148068791 |
| SPBC4F6.06 | -0.147543973 |
| SPBC660.17c | -0.146996789 |
| SPAC1F3.10c | -0.146970268 |
| SPBC2G2.09c | -0.146517931 |
| SPBC21B10.07 | -0.146352101 |
| SPAC19D5.11c | -0.145658115 |
| SPAC212.08c | -0.145564463 |
| SPBC23G7.08c | -0.145493706 |
| SPAC1B3.16c | -0.144036113 |

| | |
|---|---|
| SPBP22H7.04 | -0.143976402 |
| SPAC1296.05c | -0.143512885 |
| SPBP4H10.13 | -0.142219486 |
| SPAC15A10.06 | -0.142203211 |
| SPCC4G3.04c | -0.142195154 |
| SPAC1F5.03c | -0.142157 |
| SPCC777.10c | -0.142060995 |
| SPAC13G6.13 | -0.141532086 |
| SPCP1E11.05c | -0.141295119 |
| SPAC144.08 | -0.140943194 |
| SPAC5H10.10 | -0.140578104 |
| SPAC26H5.05 | -0.140382932 |
| SPAC3C7.13c | -0.140083873 |
| SPBC83.03c | -0.139282192 |
| SPBC18H10.02 | -0.138885637 |
| SPAC2F3.12c | -0.138746649 |
| SPAC24B11.07c | -0.137623344 |
| SPAC1071.08 | -0.137609207 |
| SPAC23D3.09 | -0.136898935 |
| SPAC22H10.04 | -0.136889528 |
| SPAC1006.03c | -0.136602361 |
| SPAC16E8.08 | -0.136520823 |
| SPAC31A2.15c | -0.136171381 |
| SPAC26F1.05 | -0.135843084 |
| SPAC328.04 | -0.135698218 |
| SPBPB21E7.04c | -0.13556065 |
| SPAC3A11.08 | -0.135325571 |
| SPAP11E10.02c | -0.135099243 |
| SPAC750.05c | -0.134948449 |
| SPAC8E11.02c | -0.134899445 |
| SPAC869.11 | -0.133899007 |
| SPAC17A5.01 | -0.133591415 |
| SPAC19E9.03 | -0.133161777 |
| SPBC21B10.05c | -0.133025856 |
| SPAC15A10.05c | -0.133008492 |
| SPAC22G7.05 | -0.132900807 |
| SPAC5H10.13c | -0.132642965 |
| SPBC887.05c | -0.132571923 |
| SPCC1739.03 | -0.132508872 |
| SPAC13G7.02c | -0.132333244 |
| SPCC1739.09c | -0.131662501 |
| SPBC146.09c | -0.131587707 |
| SPBC1683.13c | -0.131570178 |
| SPAC22G7.06c | -0.131549629 |
| SPBP4H10.18c | -0.130210987 |
| SPAC17A5.08 | -0.130060633 |
| SPBC1652.02 | -0.129697887 |
| SPBC21H7.06c | -0.129129197 |

| | |
|---|---|
| SPAC22G7.04 | -0.129111798 |
| SPAP27G11.05c | -0.128922271 |
| SPBCPT2R1.03 | -0.128326353 |
| SPAC17H9.10c | -0.127411007 |
| SPCC548.04 | -0.12736564 |
| SPAC823.02 | -0.127066218 |
| SPAC1071.12c | -0.126982048 |
| SPAC57A10.03 | -0.126127954 |
| SPBPB10D8.06c | -0.125931589 |
| SPCC70.06 | -0.125442616 |
| SPBC215.03c | -0.124851172 |
| SPAC9G1.10c | -0.124737863 |
| SPAC31G5.21 | -0.124532746 |
| SPAC32A11.03c | -0.124391331 |
| SPBC26H8.08c | -0.124053633 |
| SPAC806.04c | -0.123513308 |
| SPAC1783.05 | -0.123355145 |
| SPAC26A3.16 | -0.122903389 |
| SPAC6G9.09c | -0.122485695 |
| SPBC1711.01c | -0.122403494 |
| SPAPB17E12.12c | -0.122054333 |
| SPBP35G2.07 | -0.122032358 |
| SPCC970.07c | -0.121432215 |
| SPAC6G9.04 | -0.120913191 |
| SPBC115.02c | -0.120619746 |
| SPAC22F3.13 | -0.120532043 |
| SPAC607.09c | -0.120292155 |
| SPBC13E7.04 | -0.119880418 |
| SPBC1709.10c | -0.119878726 |
| SPAP14E8.02 | -0.119617303 |
| SPBC31F10.05 | -0.119386031 |
| SPBC31F10.07 | -0.119297881 |
| SPCC126.06 | -0.118763391 |
| SPAC24H6.11c | -0.118344743 |
| SPAC5H10.12c | -0.118333359 |
| SPAC227.06 | -0.118134374 |
| SPAC1F8.03c | -0.117879329 |
| SPBC29A10.06c | -0.117486866 |
| SPBC1711.05 | -0.117397302 |
| SPBC28F2.02 | -0.117386995 |
| SPBC23G7.15c | -0.117308575 |
| SPAC977.14c | -0.117180096 |
| SPBC1709.16c | -0.116665955 |
| SPBC83.09c | -0.116654437 |
| SPCC297.04c | -0.116421366 |
| SPBC1289.06c | -0.115886438 |
| SPBC2A9.06c | -0.115534972 |
| SPCC24B10.22 | -0.115532716 |

| | |
|---|---|
| SPBC1289.08 | -0.115528936 |
| SPCC757.07c | -0.115255427 |
| SPAC17A2.13c | -0.115116999 |
| SPCC1223.06 | -0.115066363 |
| SPAC227.01c | -0.114705631 |
| SPAC23D3.01 | -0.11467594 |
| SPAC1093.06c | -0.114670945 |
| SPCC1827.02c | -0.113795905 |
| SPBC428.05c | -0.113713168 |
| SPAC8C9.10c | -0.113473962 |
| SPBC21C3.11 | -0.112986587 |
| SPAC31A2.12 | -0.111859477 |
| SPACUNK12.02c | -0.111838493 |
| SPAC328.03 | -0.111434662 |
| SPBC1D7.04 | -0.111355201 |
| SPAC17C9.09c | -0.111332422 |
| SPAC458.05 | -0.111264459 |
| SPAC328.10c | -0.111240004 |
| SPAC513.03 | -0.110940613 |
| SPAC823.11 | -0.109943877 |
| SPAC19G12.05 | -0.109877953 |
| SPAC23C4.16c | -0.108884645 |
| SPAC5D6.06c | -0.108782693 |
| SPBC4B4.07c | -0.108415673 |
| SPBC16E9.08 | -0.107979968 |
| SPAC343.07 | -0.107954337 |
| SPBC1604.09c | -0.107688661 |
| SPAC12G12.09 | -0.107541104 |
| SPAC589.11 | -0.106991145 |
| SPAPB1A10.10c | -0.10672786 |
| SPAC11E3.03 | -0.106066769 |
| SPAC1071.11 | -0.105841216 |
| SPAC1782.02c | -0.105770839 |
| SPBC1604.16c | -0.105545174 |
| SPAC630.06c | -0.105245956 |
| SPBC336.13c | -0.10499418 |
| SPBP35G2.06c | -0.104814831 |
| SPBP23A10.05 | -0.104750756 |
| SPBP8B7.23 | -0.104722413 |
| SPAC323.04 | -0.104495878 |
| SPCC794.11c | -0.104467299 |
| SPBC3B9.13c | -0.104115433 |
| SPBC21C3.07c | -0.104024267 |
| SPAC22F3.12c | -0.103968854 |
| SPAC25H1.02 | -0.103736836 |
| SPBC12C2.01c | -0.10345716 |
| SPAC24B11.13 | -0.103280481 |
| SPBP8B7.06 | -0.103127528 |

| | |
|---|---|
| SPAC56E4.06c | -0.103034864 |
| SPBC36B7.03 | -0.10288417 |
| SPAC31A2.09c | -0.102842104 |
| SPBC23G7.06c | -0.102761691 |
| SPCC790.02 | -0.102530802 |
| SPCC1620.07c | -0.102406065 |
| SPBC24C6.04 | -0.102351516 |
| SPAC13G6.10c | -0.101997395 |
| SPCC1259.02c | -0.10199474 |
| SPAC17H9.14c | -0.101882468 |
| SPAC18B11.04 | -0.101520268 |
| SPAC458.02c | -0.101504098 |
| SPAC7D4.13c | -0.101116631 |
| SPAPB1A10.14 | -0.100983173 |
| SPAC19A8.05c | -0.100865221 |
| SPAC30C2.05 | -0.100396967 |
| SPAC4G9.12 | -0.100111372 |
| SPAPB17E12.04c | -0.099954346 |
| SPBC19C2.02 | -0.099797285 |
| SPCC1682.14 | -0.099680732 |
| SPBC17D11.02c | -0.09911269 |
| SPBC32H8.13c | -0.098813611 |
| SPBC3E7.10 | -0.098409667 |
| SPBC409.03 | -0.098356599 |
| SPAC1F5.09c | -0.098162451 |
| SPAC23H3.03c | -0.09770479 |
| SPAC521.04c | -0.097482932 |
| SPBC83.02c | -0.09715649 |
| SPBC3D6.02 | -0.097075917 |
| SPAC824.05 | -0.097028963 |
| SPAC13G7.07 | -0.096916704 |
| SPBC12D12.07c | -0.096666371 |
| SPAC1A6.01c | -0.096565492 |
| SPBP8B7.10c | -0.096247617 |
| SPBC2D10.13 | -0.096059582 |
| SPBC646.08c | -0.095877278 |
| SPAC17A5.07c | -0.095771396 |
| SPCC1620.04c | -0.095625517 |
| SPAC13D6.03c | -0.095398859 |
| SPAC16C9.04c | -0.095100859 |
| SPAC1687.13c | -0.095093821 |
| SPBC1709.11c | -0.094509101 |
| SPCC31H12.02c | -0.09426451 |
| SPAC1D4.02c | -0.094244789 |
| SPCC126.15c | -0.093511275 |
| SPBC32F12.07c | -0.093408367 |
| SPAC4C5.04 | -0.092912315 |
| SPBC17A3.06 | -0.09275332 |

| | |
|---|---|
| SPAC24H6.08 | -0.092640269 |
| SPBP26C9.02c | -0.092529899 |
| SPCC1494.08c | -0.092445796 |
| SPBC685.03 | -0.092231126 |
| SPBC1685.11 | -0.092218801 |
| SPBC4F6.08c | -0.092029206 |
| SPAC22G7.07c | -0.091511125 |
| SPBC354.13 | -0.09135363 |
| SPAC17G8.06c | -0.091256782 |
| SPBC1861.09 | -0.091248476 |
| SPCC1753.03c | -0.091237419 |
| SPCP20C8.01c | -0.091142874 |
| SPAC1071.05 | -0.090874649 |
| SPAC24B11.06c | -0.090534535 |
| SPBPJ4664.03 | -0.090350255 |
| SPCC622.12c | -0.090268533 |
| SPAC22H12.02 | -0.090134584 |
| SPBC25D12.06 | -0.090069286 |
| SPCC1919.07 | -0.089802288 |
| SPAC24H6.07 | -0.089655176 |
| SPBC13G1.08c | -0.089530252 |
| SPBC216.05 | -0.089328134 |
| SPAC26F1.02 | -0.089185672 |
| SPBC1604.02c | -0.089083892 |
| SPBC16A3.01 | -0.088951056 |
| SPAC630.05 | -0.08871061 |
| SPAC26A3.14c | -0.088640526 |
| SPBC16E9.15 | -0.088583878 |
| SPAC2G11.03c | -0.088244174 |
| SPAC589.09 | -0.088031157 |
| SPAC4F10.20 | -0.088000637 |
| SPAC1039.09 | -0.087613235 |
| SPBC1A4.04 | -0.087566568 |
| SPBC2G5.02c | -0.08715902 |
| SPAC806.08c | -0.086836175 |
| SPCC736.11 | -0.086693628 |
| SPCC757.04 | -0.086642787 |
| SPBC19G7.07c | -0.086423345 |
| SPBC947.09 | -0.086106988 |
| SPBC36.03c | -0.086048243 |
| SPCC965.04c | -0.085763691 |
| SPAC22H10.09 | -0.085667732 |
| SPBC16A3.17c | -0.085404423 |
| SPAC18B11.11 | -0.084664959 |
| SPAC18B11.08c | -0.084473458 |
| SPCC1450.03 | -0.08440272 |
| SPBC16H5.02 | -0.084401878 |
| SPAC5D6.05 | -0.084318581 |

| | |
|---|---|
| SPAC16C9.07 | -0.084266151 |
| SPBC27.06c | -0.084247724 |
| SPAPB1A10.07c | -0.083833006 |
| SPAC17G6.08 | -0.083594504 |
| SPBC29A3.07c | -0.083254208 |
| SPAC8F11.08c | -0.08323072 |
| SPAC2C4.09 | -0.082947702 |
| SPAC806.07 | -0.082845995 |
| SPBC15C4.05 | -0.082585892 |
| SPBC4F6.09 | -0.082517882 |
| SPCC126.13c | -0.082488677 |
| SPBC1289.15 | -0.082480025 |
| SPAC17A2.01 | -0.082080743 |
| SPBC1289.09 | -0.081793359 |
| SPBC337.11 | -0.08140646 |
| SPBC19C7.12c | -0.080874366 |
| SPAC4A8.04 | -0.08066737 |
| SPBC16G5.13 | -0.080513871 |
| SPBC2F12.15c | -0.080396383 |
| SPBC405.06 | -0.080334099 |
| SPBC887.17 | -0.080321951 |
| SPAC212.01c | -0.080138504 |
| SPAC15E1.05c | -0.079962258 |
| SPBC4F6.11c | -0.079828381 |
| SPCC1183.11 | -0.079728186 |
| SPAC23A1.09 | -0.079485407 |
| SPBC12D12.05c | -0.07936497 |
| SPBC3E7.12c | -0.0792829 |
| SPAC22F3.04 | -0.079180792 |
| SPBC29A3.11c | -0.079145071 |
| SPBC216.06c | -0.078731741 |
| SPBC21C3.09c | -0.078688059 |
| SPAC3F10.12c | -0.078404329 |
| SPAC1783.08c | -0.077835338 |
| SPAC20G8.09c | -0.077773182 |
| SPAC6F6.03c | -0.077522312 |
| SPBC21B10.04c | -0.077368555 |
| SPAC14C4.10c | -0.077352234 |
| SPCC1494.07 | -0.077028392 |
| SPAC11H11.03c | -0.076917579 |
| SPBC14C8.04 | -0.076570061 |
| SPAC664.01c | -0.076082281 |
| SPCC830.07c | -0.075940346 |
| SPBC660.06 | -0.075924337 |
| SPAC18G6.02c | -0.075914964 |
| SPBC25B2.03 | -0.075545575 |
| SPAPB1E7.08c | -0.074881579 |
| SPBC119.03 | -0.074448763 |

| | |
|---|---|
| SPBC16A3.16 | -0.074410788 |
| SPCC1682.12c | -0.074215498 |
| SPAC5H10.11 | -0.074050394 |
| SPBC660.05 | -0.07400802 |
| SPBPB2B2.12c | -0.073890528 |
| SPAC3H8.09c | -0.073805024 |
| SPAC13G6.01c | -0.073747646 |
| SPCC320.14 | -0.073694965 |
| SPAC6G9.13c | -0.073525338 |
| SPBC30B4.04c | -0.073507481 |
| SPAC22H10.02 | -0.073447019 |
| SPBC1604.20c | -0.073381456 |
| SPAC926.03 | -0.073376879 |
| SPBC1198.12 | -0.073083718 |
| SPAC2F7.04 | -0.073018174 |
| SPAC2G11.10c | -0.072952266 |
| SPAC25B8.13c | -0.07293358 |
| SPAC1834.03c | -0.072598774 |
| SPAC664.02c | -0.072507061 |
| SPBPB10D8.07c | -0.072343286 |
| SPAC29B12.05c | -0.072253671 |
| SPAC186.01 | -0.071943614 |
| SPBC16C6.10 | -0.071938447 |
| SPBC83.19c | -0.071826479 |
| SPBC4.02c | -0.071491545 |
| SPBC14F5.10c | -0.071352444 |
| SPAC12G12.10 | -0.071218159 |
| SPBC839.14c | -0.071195099 |
| SPAC12G12.03 | -0.071141821 |
| SPACUNK4.15 | -0.071120032 |
| SPAC977.17 | -0.071041001 |
| SPAPJ696.01c | -0.070835421 |
| SPAC17H9.13c | -0.070782604 |
| SPAC18B11.03c | -0.070696521 |
| SPAC4H3.13 | -0.070597059 |
| SPAC19G12.13c | -0.070516934 |
| SPCC16C4.12 | -0.070316237 |
| SPBC30D10.04 | -0.070187279 |
| SPAC19G12.16c | -0.070171393 |
| SPBC18A7.02c | -0.070097216 |
| SPBC21B10.02 | -0.069837408 |
| SPBC646.09c | -0.069834411 |
| SPAC1006.04c | -0.069728708 |
| SPBC28E12.04 | -0.06972615 |
| SPCC285.11 | -0.069706066 |
| SPBC6B1.04 | -0.069681878 |
| SPAC683.02c | -0.069572942 |
| SPBC4.06 | -0.069200227 |

| | |
|---|---|
| SPCC965.05c | -0.068921989 |
| SPCC757.11c | -0.06858957 |
| SPAC1687.05 | -0.068565529 |
| SPAC1D4.06c | -0.068334598 |
| SPBC651.03c | -0.067920569 |
| SPAC1071.06 | -0.0678725 |
| SPAC3F10.09 | -0.067608718 |
| SPCC330.03c | -0.067550265 |
| SPBC119.12 | -0.06724021 |
| SPCC1020.05 | -0.067061628 |
| SPBC32F12.02 | -0.066883038 |
| SPBC24C6.09c | -0.066724289 |
| SPAC22F3.07c | -0.066649027 |
| SPAC12B10.09 | -0.06647499 |
| SPAC12G12.07c | -0.066251959 |
| SPBC29A3.03c | -0.066135147 |
| SPAC19A8.03 | -0.066058597 |
| SPAC1687.07 | -0.065993476 |
| SPBC13G1.03c | -0.065910175 |
| SPBC23G7.16 | -0.065814433 |
| SPAC9G1.12 | -0.065748473 |
| SPAC3A12.03c | -0.065687154 |
| SPCC1840.02c | -0.065618969 |
| SPAC1F8.01 | -0.065516875 |
| SPCC622.11 | -0.064785015 |
| SPBC19G7.01c | -0.064691235 |
| SPCC18.02 | -0.06452443 |
| SPAC222.08c | -0.064465216 |
| SPAC2F7.06c | -0.064437846 |
| SPAC823.03 | -0.064381393 |
| SPCC1840.05c | -0.064370357 |
| SPAC3G6.06c | -0.064345656 |
| SPBC115.03 | -0.064297448 |
| SPBC16E9.18 | -0.064289598 |
| SPAC24C9.05c | -0.064096581 |
| SPAC8C9.17c | -0.063796353 |
| SPAPJ696.02 | -0.063422668 |
| SPCC4G3.15c | -0.063301964 |
| SPBC32F12.05c | -0.063179093 |
| SPBC32C12.03c | -0.0630525 |
| SPAC6C3.08 | -0.063026691 |
| SPAC9.11 | -0.062997889 |
| SPAC3C7.01c | -0.062725604 |
| SPAC6G10.03c | -0.062724168 |
| SPBC1734.06 | -0.062714072 |
| SPBC1348.02 | -0.062690279 |
| SPBC27.04 | -0.062653969 |
| SPAC3H8.05c | -0.062633221 |

| | |
|---|---|
| SPAC24H6.04 | -0.062542701 |
| SPAC13F5.01c | -0.062486074 |
| SPAC22H10.03c | -0.062397371 |
| SPBC32H8.03 | -0.06235457 |
| SPBC713.05 | -0.062278532 |
| SPBC1347.02 | -0.062208698 |
| SPBC17D11.08 | -0.062048211 |
| SPAC25H1.04 | -0.061822419 |
| SPBC16G5.07c | -0.061672566 |
| SPBC3B9.06c | -0.061610245 |
| SPBC3B9.04 | -0.061556533 |
| SPBC2G2.02 | -0.0615055 |
| SPBC16H5.07c | -0.061483188 |
| SPAC6C3.02c | -0.061440955 |
| SPBC13E7.11 | -0.061347888 |
| SPBC56F2.12 | -0.060856551 |
| SPAC22A12.06c | -0.060855046 |
| SPAC24C9.02c | -0.06056366 |
| SPBC15D4.03 | -0.060525086 |
| SPCC74.06 | -0.060185292 |
| SPCC895.09c | -0.059716384 |
| SPCC4G3.12c | -0.059574708 |
| SPBC14F5.03c | -0.059554971 |
| SPBC2D10.15c | -0.059344384 |
| SPBPB2B2.13 | -0.059270729 |
| SPAC23A1.03 | -0.058709606 |
| SPCC663.15c | -0.058622986 |
| SPAC9E9.15 | -0.058370138 |
| SPBC32H8.05 | -0.058287833 |
| SPBC1703.06 | -0.058246822 |
| SPBC28F2.11 | -0.058242863 |
| SPBC3H7.05c | -0.058136743 |
| SPAC2E1P5.03 | -0.057989104 |
| SPAC1805.05 | -0.057947075 |
| SPBC27B12.05 | -0.057853996 |
| SPBC9B6.11c | -0.057774338 |
| SPBC1348.07 | -0.057763995 |
| SPAC23G3.03 | -0.0576699 |
| SPAC664.04c | -0.057146425 |
| SPBC19F8.02 | -0.057104616 |
| SPAC1296.06 | -0.057103726 |
| SPBC3F6.05 | -0.056951847 |
| SPAC521.05 | -0.056841602 |
| SPCC63.14 | -0.056766885 |
| SPBP8B7.09c | -0.056697995 |
| SPAC11E3.11c | -0.056675479 |
| SPAC23G3.05c | -0.056596303 |
| SPBC29A10.13 | -0.056527938 |

| | |
|---|---|
| SPAC18B11.09c | -0.056437893 |
| SPAC24C9.08 | -0.056431969 |
| SPBC365.20c | -0.056414799 |
| SPAC9G1.05 | -0.05640802 |
| SPAC821.05 | -0.05629577 |
| SPCC4G3.11 | -0.056290703 |
| SPAC1039.03 | -0.056287764 |
| SPCC1442.07c | -0.056192107 |
| SPAPB2B4.07 | -0.056006515 |
| SPAC24H6.13 | -0.055910181 |
| SPCPB1C11.03 | -0.05578822 |
| SPBC56F2.04 | -0.05545218 |
| SPAP8A3.12c | -0.055444566 |
| SPCC1223.09 | -0.055403358 |
| SPBC3B8.05 | -0.05537219 |
| SPAPB8E5.10 | -0.054602117 |
| SPBC31F10.15c | -0.054526722 |
| SPAC3H1.13 | -0.054466412 |
| SPAC23C11.08 | -0.054318122 |
| SPCC162.06c | -0.054236583 |
| SPBCPT2R1.02 | -0.054015628 |
| SPAC16.03c | -0.05393273 |
| SPBC1105.12 | -0.053829038 |
| SPAC15E1.06 | -0.053750037 |
| SPBC9B6.09c | -0.053677953 |
| SPBC2D10.11c | -0.053476685 |
| SPBC16H5.04 | -0.053394746 |
| SPCC594.04c | -0.053063009 |
| SPBC1105.11c | -0.053033429 |
| SPAPB2B4.02 | -0.052945264 |
| SPBC2A9.03 | -0.052888902 |
| SPBC2A9.13 | -0.052810573 |
| SPBC1861.07 | -0.05264259 |
| SPAC1A6.08c | -0.052552974 |
| SPAC9.05 | -0.052508441 |
| SPBC947.10 | -0.052433188 |
| SPAC22H12.01c | -0.052324872 |
| SPAC17C9.16c | -0.052269868 |
| SPAC30.04c | -0.052188846 |
| SPBC106.01 | -0.052140093 |
| SPAC11H11.01 | -0.052110753 |
| SPAC227.11c | -0.052076213 |
| SPAC824.08 | -0.051996292 |
| SPBC649.04 | -0.05189176 |
| SPBC1683.06c | -0.051667296 |
| SPBC1289.16c | -0.051649433 |
| SPBC14F5.07 | -0.051543529 |
| SPAC12G12.11c | -0.051476362 |

| | |
|---|---|
| SPBC337.03 | -0.051396773 |
| SPAC19G12.08 | -0.051385265 |
| SPBC29A10.08 | -0.051295927 |
| SPBC577.03c | -0.051221514 |
| SPAC521.02 | -0.05110653 |
| SPBC1215.01 | -0.051048467 |
| SPAPJ695.01c | -0.050724149 |
| SPCC338.08 | -0.050697294 |
| SPAC26A3.11 | -0.050486735 |
| SPBC947.06c | -0.05044064 |
| SPBC337.16 | -0.050231678 |
| SPBC902.05c | -0.050068977 |
| SPBC8E4.01c | -0.050038488 |
| SPAC14C4.05c | -0.049747149 |
| SPBC16G5.17 | -0.049725924 |
| SPCC1235.13 | -0.049653053 |
| SPCC188.02 | -0.049363831 |
| SPBC8D2.03c | -0.049183563 |
| SPAC6C3.06c | -0.049111538 |
| SPBP4H10.07 | -0.048630552 |
| SPAC1296.04 | -0.048437941 |
| SPAC328.06 | -0.048435239 |
| SPAC1296.03c | -0.048352336 |
| SPAC24H6.03 | -0.048203304 |
| SPAC1565.07c | -0.048190259 |
| SPAC31G5.18c | -0.047892825 |
| SPAC13C5.04 | -0.047891235 |
| SPBC16A3.13 | -0.047675918 |
| SPCPJ732.01 | -0.047542194 |
| SPAC222.13c | -0.047321288 |
| SPAC26H5.09c | -0.047288674 |
| SPBC3H7.14 | -0.047271783 |
| SPAC1F5.07c | -0.047217441 |
| SPAC1786.04 | -0.046873713 |
| SPBC1685.01 | -0.046827704 |
| SPBC36B7.05c | -0.046593779 |
| SPBC2G5.06c | -0.046525815 |
| SPBC18E5.11c | -0.046521543 |
| SPAC3C7.07c | -0.046422149 |
| SPAC664.12c | -0.046395148 |
| SPBC29B5.02c | -0.0463789 |
| SPAC10F6.11c | -0.046343805 |
| SPAC13G6.08 | -0.046341123 |
| SPAC29B12.14c | -0.045992237 |
| SPBC28E12.03 | -0.045986728 |
| SPBC543.07 | -0.045927439 |
| SPAC25H1.05 | -0.045862874 |
| SPAP14E8.05c | -0.045790914 |

| | |
|---|---|
| SPBC32H8.11 | -0.045776756 |
| SPCC962.01 | -0.045436812 |
| SPCC736.04c | -0.04533931 |
| SPBC1E8.02 | -0.044844044 |
| SPBC8D2.10c | -0.044755235 |
| SPBC800.03 | -0.044751289 |
| SPAC15A10.09c | -0.044748716 |
| SPAC8E11.10 | -0.044554824 |
| SPCC364.07 | -0.044439169 |
| SPAPB1A10.05 | -0.04438006 |
| SPCC1183.09c | -0.04435658 |
| SPAC5H10.06c | -0.044345457 |
| SPAC13G6.04 | -0.044259826 |
| SPAC959.07 | -0.044219495 |
| SPBC16C6.06 | -0.04420113 |
| SPBC1734.05c | -0.044151389 |
| SPAPB24D3.07c | -0.04407475 |
| SPAC9E9.12c | -0.044022053 |
| SPCC330.02 | -0.043808259 |
| SPBC776.15c | -0.043807088 |
| SPBC1685.15c | -0.043789283 |
| SPBC1271.06c | -0.043532099 |
| SPBC2G2.15c | -0.043362037 |
| SPBC1921.04c | -0.043352035 |
| SPAC3C7.09 | -0.043322842 |
| SPBC2G2.08 | -0.04296985 |
| SPAC1093.03 | -0.042903309 |
| SPBC725.02 | -0.042843602 |
| SPBC18H10.15 | -0.042814911 |
| SPAC227.15 | -0.042717994 |
| SPBC30B4.01c | -0.042657635 |
| SPBC29A10.09c | -0.042643068 |
| SPAC16E8.12c | -0.042610202 |
| SPBC1539.08 | -0.042561802 |
| SPAC6F6.11c | -0.042502136 |
| SPCC74.09 | -0.042474069 |
| SPAC25B8.05 | -0.042311918 |
| SPAC17C9.05c | -0.042049441 |
| SPBC27.03 | -0.042005148 |
| SPBC25B2.04c | -0.041944036 |
| SPCC320.06 | -0.041767004 |
| SPBC713.11c | -0.041765089 |
| SPBC1A4.02c | -0.041622714 |
| SPAC26F1.01 | -0.041511467 |
| SPBC365.07c | -0.041268626 |
| SPCC569.06 | -0.041204904 |
| SPAC30D11.02c | -0.041180296 |
| SPBC16E9.03c | -0.041109154 |

| | |
|---|---|
| SPBC29A3.05 | -0.041081118 |
| SPBC1709.18 | -0.041003054 |
| SPCC965.14c | -0.040992891 |
| SPAC1250.05 | -0.040861192 |
| SPBC16A3.10 | -0.040839511 |
| SPBC25B2.06c | -0.0404982 |
| SPCC1919.11 | -0.040381132 |
| SPCC306.07c | -0.040354483 |
| SPAC6F6.04c | -0.040327055 |
| SPCC1020.03 | -0.040237898 |
| SPAC6B12.06c | -0.040216898 |
| SPAC19G12.11 | -0.040202953 |
| SPAC4D7.11 | -0.040192009 |
| SPAC3H1.03 | -0.040147145 |
| SPAC56F8.09 | -0.040079352 |
| SPCC1827.03c | -0.039963855 |
| SPBC839.03c | -0.0398674 |
| SPBC1685.07c | -0.039846698 |
| SPAC607.08c | -0.039812234 |
| SPBC21C3.17c | -0.039811378 |
| SPAC926.02 | -0.03973918 |
| SPAC1F12.05 | -0.039517255 |
| SPAC513.02 | -0.039453628 |
| SPAC4F10.04 | -0.03935714 |
| SPBC685.02 | -0.039180422 |
| SPBC83.04 | -0.039033952 |
| SPAC1039.10 | -0.038984679 |
| SPAC1486.10 | -0.038962074 |
| SPCC330.11 | -0.038934002 |
| SPAP27G11.08c | -0.038881541 |
| SPAC25B8.07c | -0.038815255 |
| SPBC1683.03c | -0.038806028 |
| SPAC19B12.06c | -0.038802169 |
| SPAC664.15 | -0.038388502 |
| SPAC23H4.01c | -0.038330468 |
| SPAC1A6.10 | -0.038058243 |
| SPAC458.06 | -0.037915694 |
| SPBC1778.06c | -0.037771825 |
| SPAC6G9.08 | -0.037711681 |
| SPBC609.05 | -0.037626142 |
| SPAC17A5.18c | -0.037464567 |
| SPBC21B10.06c | -0.03737376 |
| SPAC1399.05c | -0.03735186 |
| SPBC3H7.06c | -0.037306256 |
| SPBC1685.13 | -0.037179752 |
| SPBC30B4.02c | -0.037019804 |
| SPAC17A5.11 | -0.037017258 |
| SPBC30D10.16 | -0.037004093 |

| | |
|---|---|
| SPAC9.13c | -0.036954731 |
| SPBC25H2.05 | -0.036825618 |
| SPBC25B2.02c | -0.036819212 |
| SPAC2G11.13 | -0.036784096 |
| SPAC3A11.06 | -0.036736052 |
| SPAC1952.03 | -0.036555195 |
| SPBC2G2.01c | -0.036429427 |
| SPBC14C8.09c | -0.036363883 |
| SPAC637.06 | -0.036345403 |
| SPAC1B3.10c | -0.036318678 |
| SPAC16E8.01 | -0.03631067 |
| SPCC1183.02 | -0.036303833 |
| SPBC1778.01c | -0.036282225 |
| SPAP27G11.10c | -0.036231435 |
| SPCC1020.07 | -0.036195831 |
| SPAC4G8.10 | -0.036166394 |
| SPAC4F10.15c | -0.036138246 |
| SPBC25B2.07c | -0.036092515 |
| SPAC922.07c | -0.036086868 |
| SPBC2A9.04c | -0.036047756 |
| SPCC1442.05c | -0.035967621 |
| SPBC31F10.13c | -0.035917775 |
| SPAC4F10.11 | -0.035855781 |
| SPAC17A5.10 | -0.035848666 |
| SPAC3H5.04 | -0.035846844 |
| SPBC1703.12 | -0.035822134 |
| SPAC977.16c | -0.035797126 |
| SPBC1709.14 | -0.035352421 |
| SPAC22F3.02 | -0.035220081 |
| SPAC22F3.03c | -0.035162627 |
| SPBP4H10.19c | -0.035149866 |
| SPAP8A3.03 | -0.035134919 |
| SPAC29B12.12 | -0.035028553 |
| SPBC428.17c | -0.034933948 |
| SPAC1071.03c | -0.034842853 |
| SPBC56F2.01 | -0.03469636 |
| SPCC594.05c | -0.034617584 |
| SPBC1711.14 | -0.034587049 |
| SPAC8C9.04 | -0.034514937 |
| SPBC337.15c | -0.034412795 |
| SPAC6B12.12 | -0.034398992 |
| SPBC1711.06 | -0.034397053 |
| SPCC794.15 | -0.034353798 |
| SPBC25H2.10c | -0.03434587 |
| SPBC1711.09c | -0.034108229 |
| SPBP16F5.04 | -0.034005545 |
| SPBC2A9.05c | -0.034002642 |
| SPCC191.03c | -0.033997038 |

| | |
|---|---|
| SPBC1539.04 | -0.033915596 |
| SPAC25H1.03 | -0.033823771 |
| SPAC767.01c | -0.033806362 |
| SPBC16C6.03c | -0.033759437 |
| SPBC56F2.08c | -0.03361708 |
| SPBC21C3.06 | -0.033612745 |
| SPCC1235.08c | -0.033585288 |
| SPCC306.11 | -0.03343764 |
| SPBC16E9.06c | -0.033400034 |
| SPCC794.03 | -0.033357905 |
| SPCC297.05 | -0.033317116 |
| SPCC736.13 | -0.033191485 |
| SPBC29A3.10c | -0.033168534 |
| SPAC890.05 | -0.033148773 |
| SPAC1687.06c | -0.032930352 |
| SPBC12D12.02c | -0.032898289 |
| SPAC19D5.01 | -0.032801687 |
| SPAC1782.04 | -0.032699692 |
| SPAC12B10.15c | -0.03269586 |
| SPBP35G2.14 | -0.032663681 |
| SPAC17A2.12 | -0.032512164 |
| SPAC3G6.09c | -0.032429016 |
| SPAC1805.04 | -0.032292291 |
| SPBC530.01 | -0.032156685 |
| SPAC1A6.05c | -0.032097128 |
| SPAC890.02c | -0.032065841 |
| SPBC3E7.09 | -0.032054366 |
| SPCC794.02 | -0.031959447 |
| SPBC725.10 | -0.031774749 |
| SPAC23H3.12c | -0.031749189 |
| SPCC4G3.02 | -0.031720768 |
| SPAC4H3.01 | -0.031712902 |
| SPBC16G5.15c | -0.031695563 |
| SPCC548.06c | -0.03168762 |
| SPAC13A11.01c | -0.031684865 |
| SPBC19C7.11 | -0.031629627 |
| SPCC794.10 | -0.031598602 |
| SPAPB24D3.09c | -0.03138656 |
| SPAC1805.07c | -0.031219259 |
| SPAC23H4.14 | -0.031137686 |
| SPBC947.05c | -0.031072806 |
| SPBPB2B2.19c | -0.031031044 |
| SPBC21D10.11c | -0.031012678 |
| SPAC19G12.12 | -0.030953918 |
| SPBC651.06 | -0.030890734 |
| SPAC27E2.02 | -0.030866526 |
| SPBC336.03 | -0.03086474 |
| SPAC25H1.06 | -0.030833263 |

| | |
|---|---|
| SPBC530.07c | -0.030820081 |
| SPAPB1A10.09 | -0.030740134 |
| SPCC548.07c | -0.030714723 |
| SPAC1F5.08c | -0.030684156 |
| SPAC3A11.11c | -0.030597269 |
| SPBC1706.03 | -0.030595922 |
| SPAC24C9.07c | -0.030558132 |
| SPAC1A6.07 | -0.03047505 |
| SPBC11C11.01 | -0.030341123 |
| SPAPB24D3.01 | -0.030180576 |
| SPAC30C2.08 | -0.030091789 |
| SPAC3H5.05c | -0.030009319 |
| SPAPB17E12.08 | -0.029739206 |
| SPAC31G5.07 | -0.029737397 |
| SPAC1F3.03 | -0.029652423 |
| SPAC1A6.06c | -0.02954359 |
| SPBC1703.14c | -0.029521503 |
| SPAC29A4.19c | -0.029509743 |
| SPBC1347.06c | -0.029506701 |
| SPCC777.17c | -0.029430164 |
| SPBC83.01 | -0.029408105 |
| SPBC29A3.17 | -0.02936021 |
| SPAC977.15 | -0.02921553 |
| SPCC790.03 | -0.029203804 |
| SPAC2F7.09c | -0.029203721 |
| SPAC343.12 | -0.029145487 |
| SPCP20C8.02c | -0.029039045 |
| SPBC405.03c | -0.028812437 |
| SPAC24H6.10c | -0.028808394 |
| SPAC23C11.01 | -0.02879704 |
| SPBC15C4.06c | -0.028762689 |
| SPAC589.12 | -0.02873427 |
| SPBC428.11 | -0.028647461 |
| SPBC3E7.16c | -0.028637394 |
| SPAC186.07c | -0.02857425 |
| SPBC36.01c | -0.028486114 |
| SPCC330.12c | -0.028413106 |
| SPAC6C3.05 | -0.028313392 |
| SPAC1039.07c | -0.028261866 |
| SPBC577.08c | -0.028224959 |
| SPBC28F2.03 | -0.02818731 |
| SPBC1921.03c | -0.028117707 |
| SPBC20F10.02c | -0.028098428 |
| SPBC2G5.04c | -0.028070071 |
| SPAC823.10c | -0.028019626 |
| SPBC3E7.07c | -0.027902799 |
| SPAC3A11.03 | -0.027802711 |
| SPBC32H8.01c | -0.027655196 |

| | |
|---|---|
| SPAC22G7.11c | -0.02762198 |
| SPBC317.01 | -0.027612344 |
| SPAC1782.08c | -0.027365809 |
| SPAC664.07c | -0.027358977 |
| SPAC2G11.07c | -0.027259969 |
| SPAC15A10.08 | -0.027203104 |
| SPCC16C4.17 | -0.027202589 |
| SPAC644.06c | -0.027087929 |
| SPBC19G7.03c | -0.027047266 |
| SPAC12B10.11 | -0.026910735 |
| SPBC29A3.09c | -0.026844788 |
| SPAC823.15 | -0.026814 |
| SPCC306.08c | -0.026796385 |
| SPCC4G3.09c | -0.026706287 |
| SPCC285.16c | -0.026706104 |
| SPCC13B11.04c | -0.026692041 |
| SPAC57A7.07c | -0.026672824 |
| SPBC1778.09 | -0.026654887 |
| SPCC1827.07c | -0.026476212 |
| SPAC22E12.03c | -0.026199503 |
| SPCC613.01 | -0.026137967 |
| SPAC19B12.12c | -0.026022331 |
| SPAC4F10.07c | -0.026010978 |
| SPCC1259.05c | -0.025959246 |
| SPBC609.04 | -0.025950998 |
| SPAC25G10.05c | -0.025930545 |
| SPBC646.13 | -0.025883 |
| SPAC32A11.01 | -0.025821152 |
| SPBC365.14c | -0.025804876 |
| SPAC1F7.01c | -0.025791538 |
| SPCC1442.15c | -0.025706254 |
| SPCC364.06 | -0.025678237 |
| SPACUNK4.09 | -0.025650806 |
| SPAC328.09 | -0.025596104 |
| SPBC577.13 | -0.025589289 |
| SPBP22H7.05c | -0.02554612 |
| SPAC26A3.02 | -0.02553487 |
| SPCC1884.02 | -0.025519417 |
| SPBC106.02c | -0.025365793 |
| SPAC17A5.05c | -0.025251596 |
| SPBC1921.07c | -0.025211772 |
| SPCC364.01 | -0.025182733 |
| SPAC140.04 | -0.025163794 |
| SPAC20G8.04c | -0.025000766 |
| SPBC13G1.10c | -0.024937936 |
| SPCC320.05 | -0.024912809 |
| SPAC1805.15c | -0.024900724 |
| SPBC19C2.14 | -0.024855178 |

| | |
|---|---|
| SPAC1783.01 | -0.024503226 |
| SPBC1711.13 | -0.02445712 |
| SPBC359.05 | -0.024365071 |
| SPBC4F6.05c | -0.024340803 |
| SPBC1347.11 | -0.024238691 |
| SPAC17G6.15c | -0.024233024 |
| SPCC757.02c | -0.024125282 |
| SPAC57A10.06 | -0.024062163 |
| SPAC4F10.18 | -0.024004117 |
| SPAC1F12.06c | -0.023891817 |
| SPAC1B3.08 | -0.023788755 |
| SPAC13C5.01c | -0.023764737 |
| SPCC126.04c | -0.023747751 |
| SPBPB10D8.01 | -0.023690103 |
| SPAC23C4.12 | -0.023644132 |
| SPAC2C4.07c | -0.02355666 |
| SPAC6B12.07c | -0.02352134 |
| SPAC16E8.13 | -0.023502509 |
| SPCC830.08c | -0.023401032 |
| SPAC2E1P5.02c | -0.023369899 |
| SPAC869.10c | -0.023337968 |
| SPAC29B12.11c | -0.023297735 |
| SPAC5D6.10c | -0.023258077 |
| SPBC23G7.14 | -0.023191279 |
| SPAC343.15 | -0.023167131 |
| SPCC613.11c | -0.023063529 |
| SPBC1711.12 | -0.023045412 |
| SPAC27F1.06c | -0.02302377 |
| SPAC3A11.14c | -0.02290133 |
| SPBC17G9.09 | -0.022855909 |
| SPBC3H7.09 | -0.022814465 |
| SPBC30D10.18c | -0.022759022 |
| SPCC1840.08c | -0.022706088 |
| SPBC887.08 | -0.022685314 |
| SPAC4C5.01 | -0.022673936 |
| SPAC19B12.11c | -0.022611167 |
| SPBC902.04 | -0.022554589 |
| SPAC13G6.02c | -0.02243294 |
| SPAPB8E5.03 | -0.022280445 |
| SPAC1296.01c | -0.022273072 |
| SPBC29A10.14 | -0.022235891 |
| SPAC1610.04 | -0.022225484 |
| SPAC2G11.15c | -0.022189242 |
| SPBC713.06 | -0.022166449 |
| SPCC794.09c | -0.022163583 |
| SPCC417.12 | -0.022146302 |
| SPBC29A10.05 | -0.021977968 |
| SPCC70.02c | -0.021971922 |

| | |
|---|---|
| SPAC13C5.05c | -0.021956016 |
| SPBC1778.07 | -0.021883142 |
| SPBC11B10.07c | -0.021878145 |
| SPAC1250.03 | -0.02187269 |
| SPBC31F10.03 | -0.021809857 |
| SPAC1142.06 | -0.021739451 |
| SPAC9E9.08 | -0.021696274 |
| SPBC725.15 | -0.021665254 |
| SPBC17D1.07c | -0.021645043 |
| SPCC1235.06 | -0.021618927 |
| SPBC16A3.12c | -0.021611685 |
| SPBC31F10.02 | -0.021572313 |
| SPBC3B8.08 | -0.021521256 |
| SPAC139.03 | -0.021342517 |
| SPBC2D10.05 | -0.021169758 |
| SPCC1827.04 | -0.021104042 |
| SPAC23C4.07 | -0.02093629 |
| SPBC19G7.02 | -0.020924193 |
| SPBC17D1.02 | -0.020909535 |
| SPAC823.13c | -0.020867102 |
| SPCC16C4.04 | -0.020825156 |
| SPBC119.14 | -0.020579448 |
| SPBC16E9.14c | -0.02052171 |
| SPAC13A11.06 | -0.02048178 |
| SPBC800.02 | -0.020288115 |
| SPCC794.08 | -0.020274669 |
| SPCC18.15 | -0.020187012 |
| SPAC15E1.04 | -0.020168057 |
| SPAC637.13c | -0.020056791 |
| SPBC1709.09 | -0.020013932 |
| SPAP7G5.05 | -0.019951273 |
| SPCC1620.08 | -0.019867409 |
| SPBC16H5.05c | -0.019822311 |
| SPAC694.03 | -0.019757042 |
| SPCC4F11.02 | -0.019685059 |
| SPBC4C3.06 | -0.019661469 |
| SPBP23A10.16 | -0.019569552 |
| SPAC4F8.08 | -0.019567369 |
| SPCC1739.01 | -0.019444225 |
| SPAC11H11.02c | -0.019438874 |
| SPAC2E1P3.04 | -0.019398629 |
| SPBC646.02 | -0.019346869 |
| SPBC18H10.09 | -0.019345098 |
| SPAC1952.15c | -0.019092717 |
| SPCC70.10 | -0.019090489 |
| SPBC1683.08 | -0.019030306 |
| SPAC458.04c | -0.018798471 |
| SPAC23H4.02 | -0.018776543 |

| | |
|---|---|
| SPAC1D4.09c | -0.018686563 |
| SPAC1805.16c | -0.018677771 |
| SPAC23C4.09c | -0.018667452 |
| SPAC3H1.04c | -0.01860608 |
| SPAC4H3.14c | -0.018572379 |
| SPCC1682.15 | -0.018526154 |
| SPAC140.03 | -0.018410066 |
| SPBC36.04 | -0.018243908 |
| SPAC8F11.05c | -0.018239192 |
| SPAC821.09 | -0.018201505 |
| SPBC651.12c | -0.018111713 |
| SPCC569.02c | -0.018091153 |
| SPBC1198.03c | -0.018028247 |
| SPCC757.05c | -0.018019917 |
| SPBC16A3.07c | -0.017900836 |
| SPAC1399.02 | -0.0178817 |
| SPAP27G11.14c | -0.017874032 |
| SPBC2G2.06c | -0.017836825 |
| SPBC11C11.08 | -0.017820694 |
| SPCC613.03 | -0.017764299 |
| SPAC144.11 | -0.01775139 |
| SPAC3C7.06c | -0.017658461 |
| SPCC285.14 | -0.017587174 |
| SPBC16E9.12c | -0.01757629 |
| SPBC1271.03c | -0.017562354 |
| SPAC19A8.08 | -0.017540333 |
| SPAC13F5.05 | -0.017511733 |
| SPAC1D4.05c | -0.017502413 |
| SPBC26H8.09c | -0.017256335 |
| SPBC16D10.05 | -0.017235132 |
| SPAC644.07 | -0.017229214 |
| SPAC14C4.09 | -0.017139078 |
| SPAC30D11.01c | -0.017119758 |
| SPCC1223.11 | -0.016983197 |
| SPAPB1A10.03 | -0.016962254 |
| SPBC354.09c | -0.016946961 |
| SPCC1739.08c | -0.016935717 |
| SPBP22H7.08 | -0.016827749 |
| SPAC1039.05c | -0.016788189 |
| SPBC2F12.05c | -0.016780678 |
| SPAC17G6.04c | -0.016689942 |
| SPCC663.08c | -0.016680218 |
| SPBC365.08c | -0.016645102 |
| SPBC685.06 | -0.016544024 |
| SPCC1020.09 | -0.016484612 |
| SPCC1739.13 | -0.016434123 |
| SPAC14C4.16 | -0.016387364 |
| SPCC5E4.05c | -0.016387333 |

| | |
|---|---|
| SPAC1399.01c | -0.016358615 |
| SPBC2D10.06 | -0.016246907 |
| SPBC365.11 | -0.016245223 |
| SPBP4H10.04 | -0.016134582 |
| SPBC20F10.06 | -0.016123584 |
| SPBC1683.02 | -0.016107977 |
| SPCC1020.12c | -0.016098714 |
| SPAC1002.12c | -0.015962112 |
| SPCP31B10.07 | -0.015902939 |
| SPBC1652.01 | -0.015875496 |
| SPBC8E4.02c | -0.015826354 |
| SPBC21C3.03 | -0.015821306 |
| SPCC965.12 | -0.015792968 |
| SPAC22E12.01 | -0.01574766 |
| SPAC4G9.20c | -0.015743407 |
| SPAC1071.02 | -0.015694637 |
| SPBC1347.13c | -0.015639577 |
| SPBC8D2.02c | -0.015634676 |
| SPBC409.10 | -0.01539561 |
| SPCC364.02c | -0.01523255 |
| SPAC1486.01 | -0.015175117 |
| SPBC1271.07c | -0.015174712 |
| SPAPB24D3.03 | -0.015133312 |
| SPAC18G6.10 | -0.015026932 |
| SPAC26H5.10c | -0.014859124 |
| SPAC17C9.07 | -0.014808964 |
| SPBC1773.02c | -0.014754517 |
| SPBC1683.09c | -0.014672074 |
| SPAC16E8.06c | -0.014591418 |
| SPAC139.02c | -0.014476999 |
| SPBC26H8.01 | -0.014435406 |
| SPBC19C2.06c | -0.014413569 |
| SPBC800.12c | -0.014284667 |
| SPAC23H3.11c | -0.01427886 |
| SPAPB2B4.06 | -0.014268813 |
| SPAC2F3.08 | -0.014229381 |
| SPBC428.07 | -0.014069222 |
| SPAC105.03c | -0.014044898 |
| SPBC1861.03 | -0.013938346 |
| SPBC405.05 | -0.013901986 |
| SPBC725.07 | -0.013869988 |
| SPAC26A3.04 | -0.013854202 |
| SPBP18G5.03 | -0.013803352 |
| SPAC14C4.13 | -0.013727656 |
| SPAC23H3.08c | -0.013567969 |
| SPBC365.12c | -0.01352007 |
| SPBC1105.02c | -0.013514847 |
| SPCC16C4.11 | -0.01351342 |

| | |
|---|---|
| SPCC1020.10 | -0.01346655 |
| SPCP1E11.09c | -0.013326704 |
| SPBC3B8.07c | -0.013263935 |
| SPAC3C7.03c | -0.01325149 |
| SPAC26A3.17c | -0.013247016 |
| SPAC343.06c | -0.013202073 |
| SPBP26C9.03c | -0.013003788 |
| SPAPYUG7.04c | -0.012946754 |
| SPAC12B10.16c | -0.01289113 |
| SPAP27G11.02 | -0.012869577 |
| SPAC186.03 | -0.01278276 |
| SPBC1683.10c | -0.012769263 |
| SPAC16A10.04 | -0.012736799 |
| SPBP35G2.02 | -0.012735421 |
| SPAC23C4.03 | -0.01270261 |
| SPCC1840.10 | -0.012680213 |
| SPAC222.05c | -0.012663595 |
| SPBC839.02 | -0.012652307 |
| SPBC15D4.05 | -0.012629805 |
| SPAC16A10.03c | -0.012561691 |
| SPAC1687.19c | -0.012513544 |
| SPBC530.02 | -0.012398155 |
| SPAC630.07c | -0.012392559 |
| SPBC83.13 | -0.012264009 |
| SPCC757.03c | -0.01224878 |
| SPAC19A8.14 | -0.012199454 |
| SPBC16H5.13 | -0.012137035 |
| SPAC6G10.11c | -0.012051203 |
| SPBC839.04 | -0.012033332 |
| SPCC1919.13c | -0.012023008 |
| SPAC1F7.12 | -0.01198942 |
| SPBC3E7.05c | -0.011937623 |
| SPCP31B10.02 | -0.011901326 |
| SPAC57A10.04 | -0.011839969 |
| SPAC1D4.01 | -0.011838477 |
| SPBC1347.07 | -0.011824816 |
| SPBPB10D8.04c | -0.011751206 |
| SPAC21E11.05c | -0.011710014 |
| SPAC1093.01 | -0.011700222 |
| SPCC1450.09c | -0.011619907 |
| SPCC24B10.04 | -0.011619813 |
| SPAC1B3.02c | -0.011615957 |
| SPAC19D5.06c | -0.011582768 |
| SPAC2G11.06 | -0.011563299 |
| SPAC4H3.07c | -0.011551596 |
| SPBP4H10.12 | -0.011389375 |
| SPAC13F5.04c | -0.011339045 |
| SPAC3G9.03 | -0.011275353 |

| | |
|---|---|
| SPBC902.02c | -0.011209027 |
| SPBC18H10.08c | -0.011198653 |
| SPAC22F8.09 | -0.011141428 |
| SPAPJ691.02 | -0.011141263 |
| SPBC1105.18c | -0.011000008 |
| SPAC1952.17c | -0.010998296 |
| SPAC29A4.16 | -0.010890506 |
| SPBC577.06c | -0.010885847 |
| SPBC947.11c | -0.010877538 |
| SPBC15C4.04c | -0.01086828 |
| SPBC1703.07 | -0.010855689 |
| SPAPB1A10.13 | -0.01075493 |
| SPAC821.04c | -0.010437159 |
| SPAC3G6.05 | -0.01032945 |
| SPBC21C3.18 | -0.010326694 |
| SPBC28E12.02 | -0.01031045 |
| SPBC3B8.06 | -0.0102602 |
| SPCC18.09c | -0.010239016 |
| SPBC3B8.04c | -0.010080004 |
| SPBC1289.01c | -0.010054233 |
| SPBC11B10.08 | -0.009998667 |
| SPAC2G11.04 | -0.009997615 |
| SPBC24C6.10c | -0.00992306 |
| SPBC11G11.05 | -0.009862902 |
| SPAPB8E5.04c | -0.009788946 |
| SPAC24C9.16c | -0.009778481 |
| SPAPB1A10.08 | -0.009751708 |
| SPCC569.04 | -0.009547355 |
| SPAC20G8.02 | -0.009513056 |
| SPCC1682.11c | -0.009466408 |
| SPCC962.05 | -0.009448223 |
| SPBC31E1.01c | -0.009379798 |
| SPBC17A3.08 | -0.00936079 |
| SPAP7G5.04c | -0.009324363 |
| SPCC1442.04c | -0.009285711 |
| SPAC3H1.06c | -0.009274953 |
| SPAC2E1P3.02c | -0.009228634 |
| SPAC19D5.02c | -0.009218601 |
| SPAC1002.02 | -0.009209345 |
| SPBC337.13c | -0.009178881 |
| SPCC965.10 | -0.009173295 |
| SPAC6B12.09 | -0.009157695 |
| SPBC30D10.10c | -0.009142252 |
| SPBC1198.07c | -0.009103666 |
| SPBC1198.08 | -0.009098621 |
| SPBP4G3.02 | -0.009096845 |
| SPAC30.01c | -0.009089486 |
| SPBC25H2.16c | -0.008978408 |

| | |
|---|---|
| SPAC607.07c | -0.008962923 |
| SPBC1711.15c | -0.008820046 |
| SPAC1556.01c | -0.008629892 |
| SPAC3A11.07 | -0.008622294 |
| SPAC1399.03 | -0.008608077 |
| SPBC23E6.01c | -0.008555408 |
| SPBP8B7.27 | -0.00826562 |
| SPAC26F1.08c | -0.008213639 |
| SPAC14C4.07 | -0.008208296 |
| SPBC14C8.15 | -0.008198481 |
| SPAC1783.06c | -0.00817744 |
| SPAC27E2.09 | -0.008169314 |
| SPAC14C4.03 | -0.008107722 |
| SPAC1805.11c | -0.007978266 |
| SPAC6G9.03c | -0.007955217 |
| SPBC1683.11c | -0.00795493 |
| SPBC146.02 | -0.007944096 |
| SPBC409.06 | -0.00790689 |
| SPAC23A1.16c | -0.007880576 |
| SPCC1529.01 | -0.007835472 |
| SPAC3H1.07 | -0.007831604 |
| SPBC947.03c | -0.007644893 |
| SPBC13E7.07 | -0.007629511 |
| SPBC15D4.15 | -0.007531752 |
| SPAC12B10.03 | -0.007523537 |
| SPBC30D10.05c | -0.007356949 |
| SPBC12C2.04 | -0.007259356 |
| SPAC56F8.14c | -0.007240701 |
| SPAC644.08 | -0.007187255 |
| SPBC1703.11 | -0.007151822 |
| SPCC18.01c | -0.007011553 |
| SPAC27D7.03c | -0.006980194 |
| SPAC17A2.14 | -0.006938159 |
| SPAC688.13 | -0.006931413 |
| SPAC6G9.01c | -0.00686209 |
| SPCC1840.07c | -0.006850622 |
| SPBC18H10.04c | -0.006815354 |
| SPBC354.07c | -0.006808037 |
| SPAC10F6.12c | -0.006795072 |
| SPAC1805.03c | -0.006774559 |
| SPAC343.11c | -0.006744292 |
| SPCC4E9.02 | -0.006488132 |
| SPBC83.10 | -0.006398786 |
| SPAC22G7.01c | -0.006386885 |
| SPAC31G5.04 | -0.006356614 |
| SPBC23G7.11 | -0.006221626 |
| SPBC1773.05c | -0.00608589 |
| SPAC57A10.09c | -0.005987569 |

| | |
|---|---|
| SPBC13A2.04c | -0.005967999 |
| SPBP35G2.03c | -0.005756953 |
| SPAC7D4.14c | -0.005654013 |
| SPAC1834.08 | -0.005513107 |
| SPCC70.08c | -0.005468963 |
| SPBC902.03 | -0.005420373 |
| SPBC13G1.04c | -0.005396249 |
| SPBC1703.04 | -0.005384119 |
| SPAC23H3.13c | -0.005166052 |
| SPAC1782.06c | -0.005148811 |
| SPAC19D5.07 | -0.005134825 |
| SPAC13A11.04c | -0.004932404 |
| SPAC328.05 | -0.004796023 |
| SPAC4G8.06c | -0.004724601 |
| SPAC22H10.08 | -0.004723286 |
| SPAC1B1.02c | -0.004623674 |
| SPBC543.08 | -0.004579045 |
| SPBC16D10.02 | -0.004549654 |
| SPCC569.01c | -0.004489129 |
| SPCC645.07 | -0.004414818 |
| SPBC713.02c | -0.004401955 |
| SPBC16A3.03c | -0.004397661 |
| SPBC8D2.12c | -0.004351342 |
| SPBC16H5.08c | -0.004302771 |
| SPAC343.18 | -0.00428874 |
| SPCC191.11 | -0.004243493 |
| SPAC9E9.05 | -0.004231504 |
| SPAC323.01c | -0.004188744 |
| SPCC757.12 | -0.004178195 |
| SPAC57A10.10c | -0.004157841 |
| SPAC3A12.17c | -0.004142139 |
| SPAC22A12.01c | -0.00405836 |
| SPBC25D12.05 | -0.004050212 |
| SPAC31G5.15 | -0.004026367 |
| SPBC530.09c | -0.003930145 |
| SPBC359.03c | -0.003882968 |
| SPBC1773.15 | -0.003836657 |
| SPBC530.08 | -0.003824073 |
| SPAC3A11.13 | -0.003796704 |
| SPBC18H10.13 | -0.00377717 |
| SPBC19C7.04c | -0.003761526 |
| SPAC323.03c | -0.003731822 |
| SPAC1420.03 | -0.00371279 |
| SPAC664.13 | -0.003634216 |
| SPBC1815.01 | -0.003598318 |
| SPBC428.14 | -0.003587031 |
| SPAC607.06c | -0.003564398 |
| SPCC569.08c | -0.003516972 |

| | |
|---|---|
| SPBC19C7.01 | -0.003466848 |
| SPCC1919.12c | -0.003442045 |
| SPAC23H4.10c | -0.003436242 |
| SPAC4A8.07c | -0.003389408 |
| SPAC513.07 | -0.00325938 |
| SPBC13A2.02 | -0.003173532 |
| SPAC869.02c | -0.00316799 |
| SPAC869.09 | -0.003083649 |
| SPAC22F8.05 | -0.003060183 |
| SPAC17A5.09c | -0.003038244 |
| SPCC306.02c | -0.002999659 |
| SPAC16E8.17c | -0.002978676 |
| SPBC36B7.06c | -0.002973046 |
| SPBC16D10.07c | -0.002929502 |
| SPBC3D6.10 | -0.002916003 |
| SPBC947.04 | -0.002870467 |
| SPAC13A11.03 | -0.002754944 |
| SPAC4F8.10c | -0.002692059 |
| SPAC926.05c | -0.002621 |
| SPAC27F1.08 | -0.002606978 |
| SPAC821.07c | -0.002586823 |
| SPBC16E9.13 | -0.002490729 |
| SPAC25A8.02 | -0.002376013 |
| SPBPB2B2.08 | -0.002335806 |
| SPAP8A3.04c | -0.00232109 |
| SPCC4B3.06c | -0.002255774 |
| SPBC29A3.14c | -0.002219596 |
| SPBC28E12.06c | -0.002077741 |
| SPBC354.04 | -0.00203911 |
| SPCC4G3.05c | -0.001915968 |
| SPAC9G1.08c | -0.001906355 |
| SPAC186.08c | -0.001622583 |
| SPBPB10D8.05c | -0.001550127 |
| SPAC11H11.04 | -0.001527688 |
| SPAC23A1.02c | -0.00152587 |
| SPCC320.03 | -0.001464859 |
| SPAC6F6.02c | -0.001383341 |
| SPCC1494.01 | -0.001305635 |
| SPBC1271.01c | -0.001290564 |
| SPAC105.01c | -0.001278899 |
| SPAC17G6.17 | -0.001215514 |
| SPBC21C3.15c | -0.001150564 |
| SPAC17G8.08c | -0.00112934 |
| SPAC227.07c | -0.000978784 |
| SPAC688.06c | -0.000968592 |
| SPAC16.04 | -0.000950683 |
| SPBC1198.06c | -0.000913081 |
| SPBC17D11.03c | -0.00089858 |

| | |
|---|---|
| SPBC6B1.08c | -0.000880443 |
| SPAC3C7.14c | -0.000857173 |
| SPAC1687.21 | -0.000825326 |
| SPAC22G7.02 | -0.000806605 |
| SPBC6B1.10 | -0.000779541 |
| SPAC3A11.04 | -0.000779238 |
| SPBP35G2.05c | -0.00069609 |
| SPCC663.14c | -0.000612573 |
| SPAC167.06c | -0.000575342 |
| SPBC21H7.07c | -0.00052938 |
| SPBC1734.09 | -0.000402171 |
| SPAC8C9.09c | -0.000330482 |
| SPCC594.07c | -0.000204728 |
| SPAC1B3.01c | -0.000159737 |
| SPBC1711.11 | -0.000136997 |
| SPBC83.18c | -9.94E-05 |
| SPBC30D10.14 | -8.96E-05 |
| SPBC25B2.10 | -2.23E-05 |
| SPAC13D6.04c | 3.40E-05 |
| SPAC9G1.04 | 4.96E-05 |
| SPBC4B4.02c | 0.000113934 |
| SPCC24B10.12 | 0.000139015 |
| SPAPYUK71.03c | 0.000191317 |
| SPCC5E4.10c | 0.000244118 |
| SPBPB2B2.01 | 0.000290796 |
| SPAC20G4.01 | 0.000488771 |
| SPCC1235.02 | 0.000509203 |
| SPBC3B8.10c | 0.00054935 |
| SPAC3H8.04 | 0.00063808 |
| SPBC1773.12 | 0.000732983 |
| SPAC513.04 | 0.00076258 |
| SPCP1E11.11 | 0.00078512 |
| SPBC428.02c | 0.000810739 |
| SPAC1F12.04c | 0.000813739 |
| SPAC1834.07 | 0.000864226 |
| SPCC965.07c | 0.000873437 |
| SPAC17A2.11 | 0.000920938 |
| SPBC2D10.18 | 0.001095768 |
| SPAC23H3.04 | 0.001165152 |
| SPAC20H4.03c | 0.001167911 |
| SPAC637.10c | 0.001169393 |
| SPAC1B3.07c | 0.001226761 |
| SPAC824.03c | 0.001244294 |
| SPCC736.02 | 0.001252138 |
| SPAC18G6.13 | 0.001260762 |
| SPBC1685.05 | 0.001266396 |
| SPAC3F10.06c | 0.001366267 |
| SPBPB2B2.14c | 0.001404286 |

| | |
|---|---|
| SPAC23G3.12c | 0.001439935 |
| SPAC20H4.08 | 0.001458856 |
| SPACUNK4.10 | 0.001722644 |
| SPCC364.05 | 0.002087387 |
| SPCC191.10 | 0.002123005 |
| SPAC23C11.07 | 0.002150428 |
| SPBC1778.03c | 0.002154788 |
| SPAC2C4.15c | 0.002199794 |
| SPBC18E5.01 | 0.00222613 |
| SPAC821.03c | 0.002234681 |
| SPCC737.09c | 0.002257256 |
| SPCC569.03 | 0.002258041 |
| SPCC1672.12c | 0.002276937 |
| SPBC1198.14c | 0.002314477 |
| SPBC725.05c | 0.002337987 |
| SPBC13E7.08c | 0.002348321 |
| SPAC10F6.06 | 0.002380311 |
| SPCC663.10 | 0.002386754 |
| SPAC3G9.11c | 0.002471808 |
| SPAC17C9.08 | 0.002473957 |
| SPAC11G7.06c | 0.002503853 |
| SPBC16D10.11c | 0.002570839 |
| SPBC19C2.10 | 0.002588453 |
| SPAC17C9.10 | 0.002590439 |
| SPCC622.14 | 0.002730421 |
| SPCC553.12c | 0.002740384 |
| SPAC694.04c | 0.002745584 |
| SPBC56F2.05c | 0.00277768 |
| SPCC1183.04c | 0.002791588 |
| SPBP8B7.04 | 0.00291952 |
| SPAC27D7.04 | 0.002932929 |
| SPBC2G2.03c | 0.003048099 |
| SPAC20G4.08 | 0.003070651 |
| SPCPB1C11.01 | 0.003084644 |
| SPCC1494.09c | 0.003137143 |
| SPCC1919.03c | 0.003154805 |
| SPAC222.15 | 0.003290421 |
| SPBPB7E8.02 | 0.003301473 |
| SPAC27D7.08c | 0.003318731 |
| SPAC4G9.16c | 0.003323098 |
| SPBC16G5.06 | 0.003354868 |
| SPAC26H5.04 | 0.003364502 |
| SPBC1921.05 | 0.003368936 |
| SPBC1604.07 | 0.003369368 |
| SPBC13E7.03c | 0.003381843 |
| SPBC2D10.03c | 0.003430365 |
| SPAC4H3.04c | 0.003480904 |
| SPAC12B10.12c | 0.003501788 |

| | |
|---|---|
| SPBC6B1.03c | 0.00355354 |
| SPBC1773.17c | 0.00371739 |
| SPAC23H4.09 | 0.003743505 |
| SPAC29B12.02c | 0.003777123 |
| SPBC1289.13c | 0.003797602 |
| SPCPJ732.02c | 0.003833448 |
| SPBC19G7.18c | 0.003898021 |
| SPBC11B10.05c | 0.003910753 |
| SPBC2D10.19c | 0.003976391 |
| SPAC30C2.02 | 0.004094728 |
| SPBC16E9.17c | 0.004165675 |
| SPAC6F6.09 | 0.004191477 |
| SPAC1F7.10 | 0.004197891 |
| SPAC6F12.04 | 0.004255728 |
| SPBC146.10 | 0.00428865 |
| SPAC1687.08 | 0.004329761 |
| SPCC162.10 | 0.004367021 |
| SPAC12B10.10 | 0.004397807 |
| SPBC16D10.01c | 0.004482344 |
| SPBC56F2.14 | 0.004488542 |
| SPAC3F10.04 | 0.00449231 |
| SPBC18H10.10c | 0.004500127 |
| SPAC10F6.15 | 0.004521891 |
| SPAC18G6.01c | 0.004541994 |
| SPAC19G12.04 | 0.004547391 |
| SPBC8D2.18c | 0.004574502 |
| SPBC18E5.10 | 0.004595641 |
| SPAC15F9.02 | 0.00460618 |
| SPAC20G8.07c | 0.004617185 |
| SPBC28F2.05c | 0.004665178 |
| SPBC8D2.16c | 0.004716875 |
| SPBPB2B2.09c | 0.004741908 |
| SPAC227.18 | 0.004827383 |
| SPBC1921.01c | 0.004869671 |
| SPAC694.06c | 0.005034473 |
| SPBC12C2.05c | 0.005069741 |
| SPBC11B10.06 | 0.005093048 |
| SPAC1834.04 | 0.005123164 |
| SPBC14F5.13c | 0.005158704 |
| SPBC36B7.02 | 0.005195814 |
| SPAC2E12.03c | 0.005205509 |
| SPBC8E4.05c | 0.005305997 |
| SPAC1805.01c | 0.005424066 |
| SPAC1565.03 | 0.005543885 |
| SPAC19G12.03 | 0.005594802 |
| SPAC24B11.12c | 0.005610499 |
| SPBPB21E7.01c | 0.005614474 |
| SPCC1620.13 | 0.005669019 |

| | |
|---|---|
| SPCC70.09c | 0.005679955 |
| SPAC26F1.04c | 0.005924326 |
| SPAC589.03c | 0.005959414 |
| SPAC29B12.08 | 0.006001571 |
| SPAC5D6.04 | 0.006024007 |
| SPAC56F8.06c | 0.006026951 |
| SPBC3H7.13 | 0.006031394 |
| SPAC1805.06c | 0.006122785 |
| SPAC7D4.12c | 0.006170168 |
| SPBC660.14 | 0.006204537 |
| SPBC18H10.07 | 0.006242173 |
| SPAC212.04c | 0.006281492 |
| SPBC691.03c | 0.006287226 |
| SPBC3B8.03 | 0.006346614 |
| SPBC776.04 | 0.006352898 |
| SPBC18H10.05 | 0.006380336 |
| SPAC589.08c | 0.006380848 |
| SPBC21D10.10 | 0.006396827 |
| SPAC1002.18 | 0.006429085 |
| SPAC589.07c | 0.006434421 |
| SPAC1805.14 | 0.006455594 |
| SPAC6C3.03c | 0.006467 |
| SPAC1F7.08 | 0.006482506 |
| SPAC6B12.14c | 0.006563544 |
| SPAC823.16c | 0.006634083 |
| SPAC644.11c | 0.006669407 |
| SPBC1539.02 | 0.006740942 |
| SPBC8D2.17 | 0.006823311 |
| SPAC30C2.04 | 0.006856108 |
| SPBC12D12.06 | 0.006860426 |
| SPBC3H7.12 | 0.006890278 |
| SPAC11H11.05c | 0.006944949 |
| SPCC1840.09 | 0.006980847 |
| SPBC1773.13 | 0.007005162 |
| SPAC20H4.04 | 0.007015294 |
| SPAC11E3.01c | 0.007023552 |
| SPACUNK4.08 | 0.007036841 |
| SPBC32F12.03c | 0.00710055 |
| SPBC1348.01 | 0.007177166 |
| SPBC19F8.03c | 0.00722582 |
| SPAC16C9.02c | 0.007367577 |
| SPBP22H7.06 | 0.00740276 |
| SPBC29A3.08 | 0.007426897 |
| SPAC14C4.12c | 0.007531837 |
| SPAC6G9.12 | 0.00754009 |
| SPBC902.06 | 0.007571345 |
| SPBC2G2.05 | 0.007572199 |
| SPAC57A7.05 | 0.007617438 |

| | |
|---|---|
| SPAC227.03c | 0.007622067 |
| SPBPB2B2.02 | 0.00768428 |
| SPAC3C7.10 | 0.00769392 |
| SPAC17C9.14 | 0.007697348 |
| SPBC25H2.09 | 0.007703178 |
| SPAC10F6.07c | 0.007757115 |
| SPAC15F9.01c | 0.00777608 |
| SPAC22A12.17c | 0.00782548 |
| SPAC1039.06 | 0.007959959 |
| SPAC8C9.16c | 0.007985471 |
| SPBC354.01 | 0.008013661 |
| SPCC576.14 | 0.008019956 |
| SPCC31H12.04c | 0.008074456 |
| SPAC521.03 | 0.008102605 |
| SPCC1442.13c | 0.00811947 |
| SPBC887.11 | 0.008130641 |
| SPAC3A11.09 | 0.008180857 |
| SPBC359.01 | 0.008206674 |
| SPBC11G11.01 | 0.008220025 |
| SPBC23G7.07c | 0.008287722 |
| SPBC119.08 | 0.008291769 |
| SPAC14C4.11 | 0.008490729 |
| SPBP4H10.10 | 0.00853761 |
| SPAC4A8.06c | 0.008544073 |
| SPAC4F10.08 | 0.008558271 |
| SPCC285.15c | 0.008598343 |
| SPCC126.10 | 0.00870346 |
| SPAC20H4.06c | 0.008764825 |
| SPBC21C3.19 | 0.008776455 |
| SPCC188.09c | 0.008974158 |
| SPAC688.03c | 0.008988593 |
| SPBC3D6.13c | 0.009039115 |
| SPBC2G5.01 | 0.009042546 |
| SPBPB2B2.07c | 0.009129758 |
| SPBPB2B2.11 | 0.009221027 |
| SPAC2G11.12 | 0.009292273 |
| SPBP8B7.02 | 0.009417329 |
| SPAC6G9.16c | 0.009498668 |
| SPBC1105.14 | 0.009638249 |
| SPAC8C9.05 | 0.009638251 |
| SPBC839.17c | 0.009765353 |
| SPBC418.01c | 0.009786561 |
| SPBC557.05 | 0.009834472 |
| SPBC36.11 | 0.009838297 |
| SPAC6F12.12 | 0.009919764 |
| SPAC6F6.17 | 0.010204454 |
| SPAC1687.09 | 0.01030348 |
| SPAC343.16 | 0.010304849 |

| | |
|---|---|
| SPBC23G7.10c | 0.010372038 |
| SPBC3D6.05 | 0.010385975 |
| SPBC1773.01 | 0.010411112 |
| SPAPB17E12.03 | 0.010413963 |
| SPAC6B12.05c | 0.010513893 |
| SPBC19G7.06 | 0.010540429 |
| SPBC17D11.01 | 0.010570995 |
| SPBC1685.06 | 0.010601993 |
| SPAPYUG7.02c | 0.010656536 |
| SPCC1235.03 | 0.01066155 |
| SPAC25B8.09 | 0.010662981 |
| SPBP8B7.07c | 0.010702555 |
| SPBC887.15c | 0.010705377 |
| SPBPJ4664.05 | 0.010757647 |
| SPAC323.07c | 0.010777828 |
| SPCC417.07c | 0.010943553 |
| SPAPB8E5.08 | 0.010963854 |
| SPBC18H10.18c | 0.010971407 |
| SPAC869.07c | 0.011077411 |
| SPCC1902.01 | 0.011089967 |
| SPCC306.04c | 0.011134535 |
| SPBC16C6.01c | 0.011152961 |
| SPAC1399.04c | 0.011222294 |
| SPCC74.04 | 0.011243553 |
| SPAC19B12.09 | 0.011253453 |
| SPCC1494.03 | 0.011398323 |
| SPBC19C7.08c | 0.01140558 |
| SPAPB2B4.03 | 0.011549921 |
| SPBC1347.08c | 0.011564216 |
| SPBC409.11 | 0.011573293 |
| SPBC1539.03c | 0.011656223 |
| SPBC336.10c | 0.011811991 |
| SPBC25B2.08 | 0.011842592 |
| SPBC3B8.02 | 0.011966072 |
| SPAC1142.03c | 0.011987358 |
| SPBC530.06c | 0.011989262 |
| SPAC22H10.13 | 0.012007124 |
| SPAP8A3.13c | 0.012037777 |
| SPBC1685.04 | 0.012088465 |
| SPBC83.17 | 0.012113974 |
| SPAPB8E5.05 | 0.012134758 |
| SPBC2F12.03c | 0.012150218 |
| SPAC1D4.03c | 0.012227388 |
| SPCC777.15 | 0.012234702 |
| SPAC637.03 | 0.012306911 |
| SPCC1795.03 | 0.01232431 |
| SPAP7G5.03 | 0.012335512 |
| SPAC4G9.14 | 0.012380125 |

| | |
|---|---|
| SPAC3G6.04 | 0.012399615 |
| SPAC4A8.14 | 0.012480439 |
| SPBC21C3.12c | 0.012536279 |
| SPAC1B9.02c | 0.012573999 |
| SPBC16H5.11c | 0.012584055 |
| SPAC4D7.07c | 0.012704304 |
| SPAC30D11.06c | 0.012779913 |
| SPBC651.04 | 0.012871991 |
| SPAC26A3.01 | 0.012879876 |
| SPAC890.06 | 0.012916231 |
| SPCC162.04c | 0.01296541 |
| SPAC6F6.06c | 0.012966212 |
| SPBC3H7.15 | 0.013029569 |
| SPAC6C3.07 | 0.013038422 |
| SPBC216.02 | 0.013046835 |
| SPBC887.01 | 0.01320975 |
| SPCC1672.03c | 0.013301091 |
| SPCC1919.01 | 0.013402739 |
| SPCPB16A4.04c | 0.013446487 |
| SPAC6C3.04 | 0.013552094 |
| SPBP35G2.13c | 0.013564792 |
| SPAC1805.02c | 0.013666101 |
| SPAC13G6.09 | 0.013676337 |
| SPAC2F3.05c | 0.013711315 |
| SPBC11G11.03 | 0.013730411 |
| SPBC21B10.09 | 0.01375056 |
| SPBC106.03 | 0.013858578 |
| SPAC227.04 | 0.013950389 |
| SPBC1711.04 | 0.013996582 |
| SPAC1142.08 | 0.014033997 |
| SPCC1450.08c | 0.014041946 |
| SPAC15E1.02c | 0.014110277 |
| SPAC32A11.02c | 0.014121581 |
| SPAC4D7.02c | 0.014181484 |
| SPBP35G2.12 | 0.014327154 |
| SPAC10F6.17c | 0.014336548 |
| SPCC1906.02c | 0.014342536 |
| SPAC23G3.04 | 0.014368384 |
| SPBC146.06c | 0.014431065 |
| SPBC530.14c | 0.01463646 |
| SPAC3F10.10c | 0.014640555 |
| SPAC11E3.09 | 0.014688787 |
| SPBC1A4.05 | 0.014722226 |
| SPCC1902.02 | 0.014801261 |
| SPAC31G5.14 | 0.014926092 |
| SPAC13C5.03 | 0.014949593 |
| SPBC29A10.07 | 0.014955707 |
| SPCC1840.04 | 0.015126189 |

| | |
|---|---|
| SPBC25H2.03 | 0.015141647 |
| SPBP4H10.17c | 0.015184301 |
| SPAC3H1.12c | 0.015363237 |
| SPBP23A10.10 | 0.01538619 |
| SPAC3A11.02 | 0.015393171 |
| SPAC6B12.03c | 0.015554491 |
| SPCC965.09 | 0.015555679 |
| SPAC15A10.07 | 0.01556036 |
| SPAC589.05c | 0.015574927 |
| SPAC12B10.14c | 0.015649644 |
| SPAC1002.14 | 0.015713586 |
| SPAC513.01c | 0.015730987 |
| SPAC1687.16c | 0.015762495 |
| SPCC126.08c | 0.015882472 |
| SPAC212.02 | 0.015908378 |
| SPAC19E9.02 | 0.015964214 |
| SPAC19G12.09 | 0.016018945 |
| SPBC1271.08c | 0.016024627 |
| SPAC823.09c | 0.016203231 |
| SPAC57A10.07 | 0.016222641 |
| SPAC4A8.02c | 0.016227667 |
| SPBC14C8.11c | 0.016236261 |
| SPAC27D7.06 | 0.016306957 |
| SPBC16G5.03 | 0.016397714 |
| SPBC30D10.03c | 0.016510974 |
| SPAC22A12.10 | 0.016525779 |
| SPBC19C2.09 | 0.016559493 |
| SPBC8D2.11 | 0.016736493 |
| SPBC16H5.06 | 0.016773932 |
| SPBC4C3.09 | 0.016799535 |
| SPBC428.12c | 0.016804106 |
| SPCC4G3.08 | 0.016844475 |
| SPAC21E11.04 | 0.016941831 |
| SPCC1840.12 | 0.017102456 |
| SPAC23H3.05c | 0.017178873 |
| SPAC7D4.05 | 0.01721025 |
| SPAC2E1P3.05c | 0.017211486 |
| SPBC2F12.09c | 0.017297869 |
| SPAC1002.19 | 0.017387988 |
| SPAP27G11.15 | 0.0174138 |
| SPBC11B10.10c | 0.017439762 |
| SPAC16C9.01c | 0.017843382 |
| SPAC1782.09c | 0.017951865 |
| SPAC56F8.12 | 0.017985701 |
| SPAPYUG7.06 | 0.01808983 |
| SPCC13B11.03c | 0.018112366 |
| SPAC1F7.06 | 0.018172368 |
| SPAC4F8.15 | 0.018219274 |

| | |
|---|---|
| SPBC839.05c | 0.018273708 |
| SPAC22F3.11c | 0.018363436 |
| SPBC2G5.03 | 0.01838535 |
| SPAC22A12.16 | 0.018409908 |
| SPBPB2B2.10c | 0.018410005 |
| SPBC29A3.01 | 0.018424234 |
| SPAC8F11.10c | 0.018501345 |
| SPBC1709.01 | 0.018540306 |
| SPAC17A5.04c | 0.018568186 |
| SPBC1711.08 | 0.018568497 |
| SPAC1039.04 | 0.018570361 |
| SPCC1919.04 | 0.018894922 |
| SPCC1906.04 | 0.018925115 |
| SPCC663.03 | 0.018952527 |
| SPAC22G7.08 | 0.018972643 |
| SPBC14C8.03 | 0.019029195 |
| SPAC644.13c | 0.019129244 |
| SPAC1952.08c | 0.019157919 |
| SPBP35G2.11c | 0.01916365 |
| SPCC777.03c | 0.019298593 |
| SPAC1142.01 | 0.019334283 |
| SPAC11E3.13c | 0.019347712 |
| SPBC336.14c | 0.01944907 |
| SPCC970.02 | 0.019482783 |
| SPAC25A8.01c | 0.019490159 |
| SPBC776.11 | 0.019507603 |
| SPBC2D10.09 | 0.019694394 |
| SPBC530.03c | 0.019727561 |
| SPBC13G1.14c | 0.019754485 |
| SPBC56F2.09c | 0.019801226 |
| SPAC22F3.09c | 0.019975331 |
| SPAC644.15 | 0.020222859 |
| SPAC13D6.01 | 0.020288926 |
| SPCC320.07c | 0.020295591 |
| SPAC22F3.10c | 0.020295631 |
| SPAC1786.01c | 0.020320439 |
| SPCC4G3.19 | 0.02054125 |
| SPBC1709.19c | 0.020725769 |
| SPBC530.15c | 0.020741589 |
| SPBC1683.12 | 0.020743551 |
| SPAC3H8.02 | 0.020799558 |
| SPBC11C11.06c | 0.020827769 |
| SPAC26A3.09c | 0.020915184 |
| SPAC23C11.13c | 0.020920536 |
| SPAC144.17c | 0.021014831 |
| SPAC18B11.02c | 0.021061149 |
| SPCC18B5.05c | 0.021063063 |
| SPAC25H1.09 | 0.021085392 |

| | |
|---|---|
| SPBC23G7.04c | 0.02122539 |
| SPBC27B12.03c | 0.021396735 |
| SPAC4F10.06 | 0.02148992 |
| SPAC7D4.02c | 0.021495546 |
| SPAPB18E9.04c | 0.021506329 |
| SPAC23D3.13c | 0.021521528 |
| SPBC15C4.01c | 0.021535071 |
| SPAC17H9.12c | 0.02163798 |
| SPBPB2B2.05 | 0.021691145 |
| SPCC970.05 | 0.021777742 |
| SPACUNK4.19 | 0.021790432 |
| SPAC22A12.02c | 0.021792573 |
| SPAC24H6.02c | 0.021847112 |
| SPCC1739.05 | 0.021901126 |
| SPAC8E11.03c | 0.021927394 |
| SPAC12B10.13 | 0.022171443 |
| SPAC589.10c | 0.022181192 |
| SPAC144.01 | 0.022196019 |
| SPAC12G12.01c | 0.022224427 |
| SPBC887.06c | 0.02222565 |
| SPBC20F10.03 | 0.022410628 |
| SPAP27G11.12 | 0.022462808 |
| SPAC17A5.16 | 0.022489529 |
| SPBC15D4.12c | 0.022507425 |
| SPBC725.06c | 0.022615242 |
| SPCP1E11.07c | 0.022618558 |
| SPBC342.05 | 0.022666336 |
| SPAC20G4.03c | 0.022808775 |
| SPAC29E6.05c | 0.022830592 |
| SPAC959.05c | 0.022850977 |
| SPAC57A7.13 | 0.02288877 |
| SPBC1347.09 | 0.022924972 |
| SPBC365.16 | 0.023003837 |
| SPAC8E11.04c | 0.023015051 |
| SPBP4H10.05c | 0.023050704 |
| SPAC1952.12c | 0.023078096 |
| SPBC713.09 | 0.023093197 |
| SPBPJ4664.06 | 0.023109711 |
| SPBC543.10 | 0.023152885 |
| SPBC660.11 | 0.023192025 |
| SPAC664.10 | 0.023296671 |
| SPBC1773.14 | 0.023310416 |
| SPBC17G9.12c | 0.02331991 |
| SPBC216.03 | 0.023322147 |
| SPAC694.02 | 0.023332406 |
| SPAC9E9.14 | 0.023336614 |
| SPCC1672.09 | 0.023351569 |
| SPAC9E9.10c | 0.023353281 |

| | |
|---|---|
| SPAC27D7.12c | 0.023367488 |
| SPBC12C2.09c | 0.023422082 |
| SPBP35G2.10 | 0.023519799 |
| SPAC1420.01c | 0.023649234 |
| SPAC17C9.15c | 0.023654115 |
| SPBC4C3.08 | 0.023687949 |
| SPAC27D7.05c | 0.023700383 |
| SPAPB1A11.04c | 0.02376187 |
| SPAC30D11.09 | 0.023810311 |
| SPBC776.16 | 0.023828224 |
| SPBC947.01 | 0.023940069 |
| SPAC664.14 | 0.024024043 |
| SPAC15A10.15 | 0.024098497 |
| SPBC30B4.06c | 0.024113846 |
| SPBC146.11c | 0.024159651 |
| SPAC22E12.04 | 0.024184541 |
| SPCC285.17 | 0.024347409 |
| SPCC1739.06c | 0.024387952 |
| SPAC1805.10 | 0.024421924 |
| SPAC12B10.05 | 0.024699548 |
| SPBC13E7.06 | 0.024893847 |
| SPBC1773.08c | 0.024896098 |
| SPBC776.14 | 0.02494299 |
| SPAC869.03c | 0.025018619 |
| SPBC27.02c | 0.025087554 |
| SPBC776.05 | 0.025336245 |
| SPAP14E8.04 | 0.025343705 |
| SPBC106.19 | 0.025454317 |
| SPBC725.09c | 0.025566572 |
| SPAC3C7.02c | 0.025567481 |
| SPCC330.19c | 0.025681592 |
| SPCC1442.16c | 0.025698987 |
| SPAC27F1.10 | 0.025719401 |
| SPBC530.11c | 0.025722265 |
| SPAC5D6.13 | 0.02575321 |
| SPAC1610.01 | 0.025756045 |
| SPAC343.19 | 0.025767377 |
| SPAC4A8.09c | 0.025771043 |
| SPBC1271.05c | 0.025891963 |
| SPCC18.10 | 0.025912363 |
| SPBC11C11.11c | 0.025914887 |
| SPBC8E4.03 | 0.025941662 |
| SPAC869.06c | 0.026101772 |
| SPBC16G5.09 | 0.02616306 |
| SPAC9.08c | 0.026177206 |
| SPBC23E6.10c | 0.02623349 |
| SPCC569.05c | 0.026300442 |
| SPBC4.05 | 0.026351337 |

| | |
|---|---|
| SPCC4G3.13c | 0.026669437 |
| SPBC1773.04 | 0.026858088 |
| SPBC1289.14 | 0.026886392 |
| SPAC24H6.09 | 0.026942562 |
| SPAC23C4.02 | 0.027017288 |
| SPBC29A3.02c | 0.027130134 |
| SPAC1565.01 | 0.027149233 |
| SPBC3F6.01c | 0.027154513 |
| SPAC57A7.09 | 0.027246486 |
| SPCC1442.11c | 0.027365612 |
| SPAC17C9.11c | 0.027477781 |
| SPAC1952.09c | 0.027571409 |
| SPAC14C4.08 | 0.027668237 |
| SPAC17G6.03 | 0.027722317 |
| SPBC1105.09 | 0.027775883 |
| SPBC19F8.04c | 0.027787508 |
| SPAC977.05c | 0.02783655 |
| SPAC27D7.11c | 0.02784934 |
| SPBP8B7.21 | 0.027908154 |
| SPBC19C7.05 | 0.027919908 |
| SPAC1B3.15c | 0.027925597 |
| SPAC15E1.09 | 0.027941625 |
| SPAC23C11.04c | 0.027947072 |
| SPAC22E12.18 | 0.028070519 |
| SPCC18B5.03 | 0.028078731 |
| SPAC1527.03 | 0.028095277 |
| SPAC23D3.03c | 0.028147392 |
| SPBC14C8.05c | 0.028285801 |
| SPAP8A3.02c | 0.028307894 |
| SPBC56F2.06 | 0.028618886 |
| SPBC582.06c | 0.028713936 |
| SPBC1271.12 | 0.028774852 |
| SPAC23G3.07c | 0.028829113 |
| SPAC12B10.04 | 0.028863653 |
| SPCC285.10c | 0.028907002 |
| SPCC553.07c | 0.028979116 |
| SPBC13G1.12 | 0.029049883 |
| SPCC594.01 | 0.029199408 |
| SPAC26H5.08c | 0.029264176 |
| SPBC8D2.19 | 0.029288319 |
| SPCC18B5.06 | 0.029387542 |
| SPAC29A4.18 | 0.029484256 |
| SPCC777.04 | 0.029526783 |
| SPBC16C6.05 | 0.029653233 |
| SPBC660.09 | 0.029684546 |
| SPBC20F10.05 | 0.029690018 |
| SPCC126.02c | 0.029713922 |
| SPBC418.02 | 0.029955033 |

| | |
|---|---|
| SPCC663.09c | 0.029976855 |
| SPBC354.08c | 0.030004124 |
| SPCC18.06c | 0.030050268 |
| SPBC16E9.11c | 0.030086785 |
| SPAC57A7.12 | 0.030182517 |
| SPAC29A4.09 | 0.030195572 |
| SPBC16G5.11c | 0.030203504 |
| SPAPB8E5.06c | 0.030265604 |
| SPBC14F5.09c | 0.030267481 |
| SPAC11E3.14 | 0.030285845 |
| SPBC3B9.11c | 0.030354097 |
| SPAC922.06 | 0.030379968 |
| SPAC1952.10c | 0.030450286 |
| SPBC428.10 | 0.030505505 |
| SPAC17H9.06c | 0.030506799 |
| SPCC1020.11c | 0.030550571 |
| SPCC162.03 | 0.030574548 |
| SPAC186.06 | 0.030608407 |
| SPBC29A10.11c | 0.03070962 |
| SPBC1685.02c | 0.030744244 |
| SPBC106.05c | 0.030752765 |
| SPCC417.03 | 0.030878667 |
| SPAC3A12.10 | 0.031150347 |
| SPBP4H10.16c | 0.031192553 |
| SPCC417.05c | 0.031335015 |
| SPCC553.03 | 0.031433453 |
| SPCC4F11.03c | 0.031437139 |
| SPBC1271.11 | 0.031580427 |
| SPAC8F11.09c | 0.031644829 |
| SPBC215.10 | 0.031680637 |
| SPAC1610.03c | 0.031731237 |
| SPBC106.11c | 0.031841071 |
| SPCC962.04 | 0.031870617 |
| SPAC10F6.04 | 0.031874281 |
| SPBC23E6.03c | 0.031881586 |
| SPCC285.04 | 0.031947815 |
| SPAC6G10.08 | 0.031957442 |
| SPBC21H7.03c | 0.031993471 |
| SPBC32H8.08c | 0.032047573 |
| SPAC1F12.10c | 0.032157243 |
| SPBP8B7.08c | 0.032160634 |
| SPCC737.07c | 0.032200871 |
| SPAC7D4.06c | 0.032242348 |
| SPAC1B3.06c | 0.032285723 |
| SPCC18B5.11c | 0.032372661 |
| SPBC1105.05 | 0.032403613 |
| SPCC191.09c | 0.032435075 |
| SPAC6F6.13c | 0.032551788 |

| | |
|---|---|
| SPAC4C5.02c | 0.032573312 |
| SPBC1734.07c | 0.032672973 |
| SPCC16A11.03c | 0.032677121 |
| SPBC26H8.11c | 0.03270376 |
| SPAC3H1.14 | 0.032722184 |
| SPCC14G10.04 | 0.032823021 |
| SPAC14C4.04 | 0.032831577 |
| SPAC22G7.03 | 0.033044423 |
| SPBC1778.04 | 0.033097506 |
| SPCPB1C11.02 | 0.033104188 |
| SPAC22F8.02c | 0.033113311 |
| SPCC18.13 | 0.033142885 |
| SPBC21.03c | 0.033274373 |
| SPCC18.03 | 0.033303892 |
| SPCC576.04 | 0.033339751 |
| SPAC688.12c | 0.033348216 |
| SPAC17G6.02c | 0.033550709 |
| SPBC31E1.02c | 0.033603838 |
| SPAC1783.04c | 0.033608905 |
| SPAC13F5.03c | 0.033623511 |
| SPAC1687.23c | 0.03370402 |
| SPBC146.04 | 0.033763035 |
| SPAC3A12.13c | 0.033819775 |
| SPAC19D5.03 | 0.033884748 |
| SPAC3G9.01 | 0.034095966 |
| SPAC2H10.01 | 0.03415189 |
| SPAC29A4.05 | 0.034217656 |
| SPBC2A9.07c | 0.03423163 |
| SPAC3G6.13c | 0.034257843 |
| SPAC26H5.07c | 0.034284544 |
| SPCP31B10.05 | 0.034306048 |
| SPAC24C9.12c | 0.034320665 |
| SPAC30.03c | 0.034323321 |
| SPAC9E9.11 | 0.034395603 |
| SPBC947.15c | 0.034537056 |
| SPCC126.03 | 0.034607521 |
| SPAC922.04 | 0.034667901 |
| SPAC1556.04c | 0.034682636 |
| SPAC1782.01 | 0.034736676 |
| SPBC15D4.01c | 0.034850028 |
| SPCC1235.12c | 0.034868384 |
| SPCC1450.07c | 0.03487168 |
| SPAC20H4.09 | 0.034903949 |
| SPAC11E3.08c | 0.03494227 |
| SPAPB1E7.04c | 0.03494752 |
| SPAC1B3.03c | 0.034963176 |
| SPBC21D10.12 | 0.034967259 |
| SPAC19A8.02 | 0.035112256 |

| | |
|---|---|
| SPBC16E9.16c | 0.035262379 |
| SPBC1604.19c | 0.035265866 |
| SPBC4B4.12c | 0.035288718 |
| SPBC725.04 | 0.035344714 |
| SPBC1604.04 | 0.035493663 |
| SPAC57A10.08c | 0.035496191 |
| SPBC29A3.13 | 0.03561398 |
| SPAC29B12.06c | 0.035646174 |
| SPAC977.06 | 0.035908985 |
| SPAC25B8.01 | 0.035938905 |
| SPCC794.12c | 0.035947252 |
| SPAC1002.07c | 0.035973222 |
| SPBC685.04c | 0.036298044 |
| SPAC2F3.01 | 0.036362782 |
| SPBC2D10.04 | 0.036424914 |
| SPCC16A11.08 | 0.036511109 |
| SPBC660.12c | 0.036562121 |
| SPBC3E7.06c | 0.036652613 |
| SPCC830.04c | 0.03677663 |
| SPAC23H4.16c | 0.036850316 |
| SPAC17H9.11 | 0.036859813 |
| SPBC1778.02 | 0.03686515 |
| SPCC663.13c | 0.036938465 |
| SPAC20G4.02c | 0.03710148 |
| SPAC22F3.08c | 0.037177756 |
| SPACUNK4.14 | 0.037187459 |
| SPBC1105.08 | 0.037245178 |
| SPAC1687.17c | 0.037377125 |
| SPBC660.10 | 0.037397026 |
| SPAC869.01 | 0.037401236 |
| SPBC211.06 | 0.03740535 |
| SPAC6F12.03c | 0.03748229 |
| SPBC8D2.01 | 0.037691133 |
| SPAC5D6.09c | 0.037770912 |
| SPAC23H3.15c | 0.037817344 |
| SPAC14C4.15c | 0.03784081 |
| SPAC29A4.14c | 0.038243129 |
| SPAC3A12.06c | 0.03848756 |
| SPBC17D1.05 | 0.038521848 |
| SPAPB1E7.07 | 0.038522366 |
| SPCC613.02 | 0.038753836 |
| SPBC1271.09 | 0.038878714 |
| SPCC16C4.03 | 0.038935464 |
| SPBC16A3.14 | 0.039024335 |
| SPCC297.06c | 0.039039915 |
| SPCC191.05c | 0.039055168 |
| SPBPB8B6.04c | 0.039098536 |
| SPAC1071.09c | 0.039239432 |

| | |
|---|---|
| SPAC11G7.04 | 0.039241306 |
| SPBC4B4.04 | 0.039415534 |
| SPAC17A2.02c | 0.039447702 |
| SPAC5D6.02c | 0.039466868 |
| SPBC21D10.08c | 0.039508804 |
| SPBC1604.03c | 0.039577995 |
| SPBC18A7.01 | 0.039690971 |
| SPAC25B8.15c | 0.039810818 |
| SPBPB21E7.09 | 0.03987014 |
| SPAC6G10.06 | 0.039875889 |
| SPCC16C4.10 | 0.03988485 |
| SPAC890.07c | 0.040003325 |
| SPAC1039.02 | 0.040062528 |
| SPAC29A4.11 | 0.040219113 |
| SPAC4G8.05 | 0.040265369 |
| SPCC18B5.01c | 0.040336202 |
| SPBP16F5.08c | 0.040400596 |
| SPAC23C4.08 | 0.040435955 |
| SPCC126.07c | 0.040671244 |
| SPCC18B5.09c | 0.040786105 |
| SPBC1778.10c | 0.040854452 |
| SPBC119.04 | 0.040875439 |
| SPAC4F8.03 | 0.040894081 |
| SPBC31F10.08 | 0.040928835 |
| SPAC23C11.02c | 0.040973731 |
| SPAC1F7.09c | 0.041017 |
| SPCC1620.03 | 0.041041875 |
| SPAC27D7.13c | 0.041166051 |
| SPBC146.13c | 0.041304149 |
| SPAP27G11.16 | 0.041324888 |
| SPAP32A8.03c | 0.041425985 |
| SPAC6G9.14 | 0.041426479 |
| SPAC23H3.14 | 0.041496095 |
| SPAC1A6.03c | 0.041509278 |
| SPBC216.04c | 0.04170162 |
| SPBC13G1.02 | 0.041737111 |
| SPCPB16A4.02c | 0.041827239 |
| SPBP8B7.25 | 0.041856732 |
| SPACUNK4.17 | 0.041869992 |
| SPAC2F3.02 | 0.041929388 |
| SPBC25D12.02c | 0.042017581 |
| SPCC16A11.16c | 0.042048309 |
| SPCC663.12 | 0.042099607 |
| SPBC1604.01 | 0.042284183 |
| SPCC663.11 | 0.042295634 |
| SPBC27B12.11c | 0.042338015 |
| SPBCPT2R1.01c | 0.042486716 |
| SPBC17G9.08c | 0.042488456 |

| | |
|---|---|
| SPAC3H8.03 | 0.04249002 |
| SPBC17F3.01c | 0.042631388 |
| SPAC1687.10 | 0.042746631 |
| SPCC737.03c | 0.04275718 |
| SPBC1347.03 | 0.042763731 |
| SPAC29E6.09 | 0.042834065 |
| SPAC12B10.06c | 0.042865314 |
| SPCC16A11.15c | 0.042912718 |
| SPCC191.06 | 0.043015353 |
| SPAPB1A11.02 | 0.043075544 |
| SPAPB24D3.02c | 0.043157695 |
| SPAC4D7.03 | 0.043214129 |
| SPBC18H10.20c | 0.043316692 |
| SPCC31H12.03c | 0.043330139 |
| SPAC140.02 | 0.04349697 |
| SPCC663.06c | 0.043564143 |
| SPAC4A8.10 | 0.043672105 |
| SPAC3C7.05c | 0.04371321 |
| SPAC9E9.03 | 0.043763947 |
| SPCC417.11c | 0.043797005 |
| SPAC6F12.06 | 0.043890926 |
| SPBC409.17c | 0.043907524 |
| SPCP1E11.02 | 0.043949259 |
| SPAC25G10.02 | 0.04396459 |
| SPBC119.16c | 0.044177352 |
| SPBC215.13 | 0.044357469 |
| SPAC959.08 | 0.044417374 |
| SPBC19C2.04c | 0.044424586 |
| SPBC83.11 | 0.044569196 |
| SPCC1442.03 | 0.044596057 |
| SPAC31G5.11 | 0.044624916 |
| SPAC4G9.06c | 0.044637591 |
| SPBC646.06c | 0.044734429 |
| SPBC16C6.04 | 0.044790091 |
| SPAC19B12.08 | 0.0448318 |
| SPBC83.05 | 0.044897582 |
| SPBC1289.11 | 0.044926784 |
| SPBC1709.06 | 0.044985851 |
| SPAC139.05 | 0.045086889 |
| SPBC215.02 | 0.045113937 |
| SPBC16E9.02c | 0.045226383 |
| SPBP35G2.04c | 0.04523762 |
| SPAC21E11.03c | 0.045267717 |
| SPAC22H12.05c | 0.045332007 |
| SPCC364.04c | 0.045416797 |
| SPAC4A8.03c | 0.045461847 |
| SPBC800.05c | 0.045475018 |
| SPAC22E12.06c | 0.045654379 |

| | |
|---|---|
| SPAC1F3.06c | 0.045697095 |
| SPAC29A4.20 | 0.045803989 |
| SPAC8E11.01c | 0.046075346 |
| SPAPB2C8.01 | 0.046188591 |
| SPBC32F12.08c | 0.046246278 |
| SPBC83.16c | 0.046393926 |
| SPBC405.02c | 0.046510301 |
| SPBC15D4.09c | 0.046854563 |
| SPBC1778.05c | 0.046972403 |
| SPCC1742.01 | 0.047104129 |
| SPAC22E12.14c | 0.047329799 |
| SPBC557.02c | 0.047437639 |
| SPAC11G7.01 | 0.047455122 |
| SPBC29A10.03c | 0.047518739 |
| SPAC56F8.05c | 0.04753797 |
| SPBC691.05c | 0.047559533 |
| SPBC19F8.06c | 0.047602661 |
| SPBP4H10.14c | 0.047941211 |
| SPCC31H12.06 | 0.048059975 |
| SPCC1223.02 | 0.04812323 |
| SPBC8D2.04 | 0.048269591 |
| SPAPB2B4.04c | 0.048316919 |
| SPBC354.10 | 0.048332283 |
| SPAC30C2.07 | 0.04839284 |
| SPBP8B7.30c | 0.048397336 |
| SPBC29A10.10c | 0.048551185 |
| SPAC17A2.07c | 0.048650336 |
| SPAC6G9.15c | 0.04866856 |
| SPBC15D4.07c | 0.048701751 |
| SPBC342.03 | 0.048703739 |
| SPBC12D12.09 | 0.048880764 |
| SPBPB10D8.02c | 0.048930558 |
| SPAC22A12.14c | 0.04903037 |
| SPCC70.04c | 0.049204787 |
| SPAC2C4.17c | 0.049465094 |
| SPBC12C2.03c | 0.049520401 |
| SPBC32H8.07 | 0.049644029 |
| SPAC20H4.11c | 0.049797852 |
| SPAC4G8.03c | 0.049854877 |
| SPCC1827.08c | 0.049921925 |
| SPAC343.09 | 0.049953017 |
| SPAC31G5.10 | 0.050253673 |
| SPBC21C3.14c | 0.05028044 |
| SPBC32F12.12c | 0.050518389 |
| SPCP31B10.04 | 0.050578657 |
| SPCC736.09c | 0.050580076 |
| SPAC25B8.10 | 0.050717978 |
| SPBC31A8.01c | 0.050725473 |

| | |
|---|---|
| SPBC1718.02 | 0.050820578 |
| SPAC607.10 | 0.050907201 |
| SPAP27G11.07c | 0.05090965 |
| SPCC191.01 | 0.051135787 |
| SPCC569.07 | 0.051300355 |
| SPCC737.05 | 0.051301991 |
| SPAC2C4.05 | 0.05136549 |
| SPBC3B9.05 | 0.05138339 |
| SPBC32H8.06 | 0.051488373 |
| SPBC1709.13c | 0.051501287 |
| SPAC26F1.07 | 0.051591275 |
| SPAC4A8.05c | 0.05165558 |
| SPAC2C4.14c | 0.051663004 |
| SPAC15A10.10 | 0.051765431 |
| SPBC216.01c | 0.05207901 |
| SPAC8C9.11 | 0.05211407 |
| SPCC63.03 | 0.052211713 |
| SPAC3H5.09c | 0.052242033 |
| SPAC16E8.05c | 0.052463719 |
| SPAC1002.06c | 0.052510552 |
| SPAC3H8.07c | 0.052594337 |
| SPAC1834.05 | 0.052601921 |
| SPAC4G8.04 | 0.052745051 |
| SPBC543.05c | 0.052759662 |
| SPAC1486.02c | 0.052841909 |
| SPCC1020.08 | 0.05291617 |
| SPCC24B10.20 | 0.053020284 |
| SPAC4F8.11 | 0.053112645 |
| SPCC70.03c | 0.053173242 |
| SPAC12G12.16c | 0.053181699 |
| SPCC16A11.01 | 0.053244846 |
| SPAC4F10.17 | 0.05330173 |
| SPCC576.17c | 0.053450312 |
| SPAC20G8.10c | 0.053492786 |
| SPCPB16A4.05c | 0.05370054 |
| SPCC285.13c | 0.053823949 |
| SPAC688.14 | 0.054424738 |
| SPAC14C4.01c | 0.054526743 |
| SPAC4H3.02c | 0.054539106 |
| SPAC9E9.09c | 0.054575463 |
| SPBC19F5.01c | 0.054609827 |
| SPBC839.15c | 0.05498616 |
| SPAC1002.01 | 0.054998993 |
| SPBC215.11c | 0.055063976 |
| SPAC9.06c | 0.055368339 |
| SPBP19A11.02c | 0.055486224 |
| SPAC1B1.04c | 0.055565767 |
| SPCC1223.01 | 0.055804223 |

| | |
|---|---|
| SPAC167.01 | 0.055821613 |
| SPBC582.08 | 0.055842889 |
| SPAC56F8.16 | 0.055851593 |
| SPBC839.13c | 0.056155074 |
| SPBC409.20c | 0.056201201 |
| SPAC6F12.09 | 0.056264985 |
| SPCC1393.09c | 0.056365536 |
| SPAPJ760.02c | 0.056366169 |
| SPBC25H2.08c | 0.056473923 |
| SPBC1683.04 | 0.056539718 |
| SPCC1393.07c | 0.057201222 |
| SPBC1105.01 | 0.057371655 |
| SPAC7D4.08 | 0.057640211 |
| SPBC557.04 | 0.057875512 |
| SPAC17G8.14c | 0.057921949 |
| SPAC9.12c | 0.058104382 |
| SPCC16C4.20c | 0.05843572 |
| SPBC3D6.08c | 0.05868429 |
| SPBC725.03 | 0.058914965 |
| SPCC757.13 | 0.058980894 |
| SPAC630.04c | 0.059002433 |
| SPBC30B4.08 | 0.059097071 |
| SPBC21D10.09c | 0.059241466 |
| SPBC16A3.19 | 0.05930605 |
| SPBC16C6.08c | 0.059315207 |
| SPAC20G4.05c | 0.059458209 |
| SPBC17A3.02 | 0.059550353 |
| SPAC222.14c | 0.059610404 |
| SPBC3E7.08c | 0.060151523 |
| SPBC20F10.07 | 0.060207744 |
| SPBC1683.01 | 0.060504421 |
| SPAC3F10.15c | 0.060508504 |
| SPAC29B12.10c | 0.060671914 |
| SPBC29B5.04c | 0.060877966 |
| SPBC1685.14c | 0.060940453 |
| SPCC16A11.07 | 0.060982292 |
| SPAC17G6.13 | 0.061105774 |
| SPBC2G2.14 | 0.061108529 |
| SPAC23H3.09c | 0.061191192 |
| SPAC688.10 | 0.061265543 |
| SPCC162.02c | 0.061295946 |
| SPBC354.15 | 0.061388675 |
| SPBC1773.03c | 0.061532566 |
| SPAPB24D3.04c | 0.061650401 |
| SPAC30D11.14c | 0.061679187 |
| SPAC1F7.11c | 0.061710465 |
| SPAC3G9.04 | 0.061875365 |
| SPBC27B12.14 | 0.062048491 |

| | |
|---|---|
| SPAC6G9.10c | 0.062119458 |
| SPBC16A3.06 | 0.062167571 |
| SPCC622.08c | 0.062182008 |
| SPBC3H7.10 | 0.062225221 |
| SPAC23A1.06c | 0.062405488 |
| SPBC215.06c | 0.062514006 |
| SPAC4F10.13c | 0.062583591 |
| SPCC965.08c | 0.062658374 |
| SPAC4F10.19c | 0.062729731 |
| SPAC23C4.05c | 0.062734478 |
| SPAC20G4.04c | 0.062970562 |
| SPAC4G9.11c | 0.063061999 |
| SPCC1739.04c | 0.063285566 |
| SPAC10F6.14c | 0.063293329 |
| SPCC63.04 | 0.063375746 |
| SPCP31B10.06 | 0.063448232 |
| SPAC19A8.10 | 0.063502732 |
| SPAC12B10.01c | 0.063528072 |
| SPAC25G10.09c | 0.063788328 |
| SPCC965.13 | 0.064207839 |
| SPAC821.06 | 0.064328633 |
| SPCC553.01c | 0.064452309 |
| SPCC576.01c | 0.064517788 |
| SPAC26F1.10c | 0.064866329 |
| SPAC869.05c | 0.064939844 |
| SPAC25H1.07 | 0.064952748 |
| SPAC16E8.14c | 0.064976255 |
| SPBC887.04c | 0.06507008 |
| SPCC63.13 | 0.065809609 |
| SPCC1020.06c | 0.065973652 |
| SPAC3G6.11 | 0.066060343 |
| SPAC186.02c | 0.066071726 |
| SPBC16A3.02c | 0.06615383 |
| SPAC1556.02c | 0.066177366 |
| SPAC17H9.01 | 0.066265305 |
| SPBC660.07 | 0.066430624 |
| SPAC1834.09 | 0.066515648 |
| SPBC3E7.15c | 0.066764244 |
| SPBC16E9.09c | 0.066793992 |
| SPAPB1A11.01 | 0.066932413 |
| SPAC3F10.05c | 0.066968016 |
| SPCC1739.10 | 0.067098002 |
| SPBC36B7.08c | 0.067267335 |
| SPBC31F10.17c | 0.067340096 |
| SPAC56F8.02 | 0.067547715 |
| SPCC737.04 | 0.067661922 |
| SPBC17G9.05 | 0.067786003 |
| SPBC29B5.03c | 0.067832649 |

| | |
|---|---|
| SPBC359.04c | 0.067886024 |
| SPBC3E7.11c | 0.067947616 |
| SPCC306.05c | 0.068121401 |
| SPBC25B2.11 | 0.068424268 |
| SPAC186.09 | 0.068625767 |
| SPBC21C3.20c | 0.068703825 |
| SPCC11E10.03 | 0.068762324 |
| SPBC3D6.06c | 0.06892087 |
| SPBP8B7.31 | 0.06897259 |
| SPBC31F10.12 | 0.068986226 |
| SPBC1734.15 | 0.069095206 |
| SPAC4G9.19 | 0.069113993 |
| SPAC8C9.12c | 0.069391508 |
| SPBC337.04 | 0.069603177 |
| SPCC1393.13 | 0.069782878 |
| SPAC607.02c | 0.069904411 |
| SPBC337.07c | 0.070237951 |
| SPCC330.07c | 0.070295546 |
| SPBC609.03 | 0.070493105 |
| SPAC821.10c | 0.070775932 |
| SPACUNK4.07c | 0.070843591 |
| SPAC1B3.04c | 0.070858961 |
| SPCC18.17c | 0.071005248 |
| SPBC428.03c | 0.07133021 |
| SPCC1919.05 | 0.071370934 |
| SPBC2F12.12c | 0.071380245 |
| SPAC31F12.01 | 0.071394564 |
| SPBC18E5.05c | 0.071426462 |
| SPBC354.14c | 0.071624302 |
| SPAC3F10.13 | 0.072333476 |
| SPBC18H10.16 | 0.0723685 |
| SPCC16C4.06c | 0.072473509 |
| SPBP4G3.03 | 0.072517039 |
| SPBC12C2.07c | 0.072672811 |
| SPAC1952.06c | 0.072677219 |
| SPAC1002.20 | 0.07267808 |
| SPCC63.06 | 0.072714075 |
| SPAC23C11.06c | 0.07345834 |
| SPAC3G6.01 | 0.073458746 |
| SPBC530.05 | 0.073551432 |
| SPBC36B7.04 | 0.073583683 |
| SPAC513.06c | 0.073606128 |
| SPBP8B7.24c | 0.073766092 |
| SPCC794.06 | 0.07386329 |
| SPBC3H7.11 | 0.073882906 |
| SPAC144.14 | 0.073980505 |
| SPAC22F8.04 | 0.074264397 |
| SPBC1289.10c | 0.07437462 |

| | |
|---|---|
| SPAC1002.17c | 0.074404851 |
| SPAC29A4.02c | 0.074808709 |
| SPAC12G12.12 | 0.074875055 |
| SPBC9B6.03 | 0.075014122 |
| SPAC20H4.07 | 0.07502962 |
| SPAC926.07c | 0.075610039 |
| SPAC5D6.07c | 0.075702921 |
| SPCC1393.05 | 0.07575545 |
| SPCC965.06 | 0.075764504 |
| SPCC777.12c | 0.075825308 |
| SPBC20F10.10 | 0.076043469 |
| SPAC7D4.03c | 0.076397197 |
| SPBC1773.06c | 0.076583866 |
| SPBC25H2.14 | 0.076859217 |
| SPAC4G8.07c | 0.076898666 |
| SPAPB24D3.10c | 0.0770504 |
| SPBC29A10.02 | 0.077060312 |
| SPAC3H8.10 | 0.077074879 |
| SPBC577.14c | 0.077117974 |
| SPBP4H10.20 | 0.077176581 |
| SPAC922.05c | 0.077213707 |
| SPCC188.12 | 0.07735391 |
| SPAC1565.02c | 0.077521098 |
| SPBC24C6.08c | 0.077808683 |
| SPBC543.02c | 0.078082497 |
| SPCC830.06 | 0.078153814 |
| SPBC428.04 | 0.078602819 |
| SPBC12C2.08 | 0.078838719 |
| SPAC22A12.04c | 0.078967905 |
| SPBC19G7.08c | 0.079146975 |
| SPAC2F3.16 | 0.079394783 |
| SPCC965.11c | 0.079444711 |
| SPAC13G7.06 | 0.079672218 |
| SPBC26H8.03 | 0.079766673 |
| SPBC18E5.07 | 0.079891463 |
| SPBC32F12.09 | 0.080141132 |
| SPAC25B8.08 | 0.080383526 |
| SPBC725.12 | 0.080476967 |
| SPAPB1A10.15 | 0.081174511 |
| SPAC23D3.11 | 0.081326081 |
| SPAC8F11.03 | 0.081382913 |
| SPBC13E7.09 | 0.08139944 |
| SPBC19C7.09c | 0.081490601 |
| SPAC23D3.10c | 0.081503108 |
| SPCC2H8.05c | 0.081530072 |
| SPCC74.03c | 0.0819026 |
| SPAC1834.10c | 0.082092732 |
| SPCC306.09c | 0.082188012 |

| | |
|---|---|
| SPBC1861.02 | 0.082294399 |
| SPAC17H9.19c | 0.082676027 |
| SPAC922.03 | 0.082742677 |
| SPBC365.01 | 0.083117357 |
| SPAC23G3.10c | 0.083313708 |
| SPCC777.07 | 0.083553138 |
| SPAC29B12.13 | 0.083725763 |
| SPBC543.03c | 0.08385366 |
| SPCC1753.05 | 0.0838877 |
| SPBC649.02 | 0.084048051 |
| SPCC970.06 | 0.084174127 |
| SPAC18G6.04c | 0.08419379 |
| SPCC11E10.08 | 0.084561481 |
| SPACUNK4.16c | 0.084889132 |
| SPBC4F6.15c | 0.084912464 |
| SPAC806.03c | 0.084940242 |
| SPBC725.11c | 0.08496944 |
| SPBC215.07c | 0.084973588 |
| SPBC336.05c | 0.085472764 |
| SPBC1709.12 | 0.085501333 |
| SPBC1773.09c | 0.085608515 |
| SPAC3H8.08c | 0.08577537 |
| SPBC1D7.01 | 0.085982427 |
| SPAC823.05c | 0.086114811 |
| SPAC30D11.11 | 0.086144973 |
| SPBC1773.16c | 0.086313004 |
| SPCC825.04c | 0.086567747 |
| SPCC24B10.13 | 0.086890427 |
| SPAC1071.04c | 0.086894841 |
| SPAPJ760.03c | 0.087093202 |
| SPAPB24D3.08c | 0.087215173 |
| SPBC336.06c | 0.087233015 |
| SPAC26F1.12c | 0.087366033 |
| SPACUNK4.13c | 0.087598593 |
| SPAC22H10.11c | 0.088126747 |
| SPAC1952.07 | 0.088965713 |
| SPBC31F10.14c | 0.08904615 |
| SPBC17G9.10 | 0.08906611 |
| SPAC19B12.07c | 0.089091718 |
| SPAC167.07c | 0.089272739 |
| SPBC1709.04c | 0.089307591 |
| SPAC9G1.06c | 0.089482181 |
| SPACUNK4.11c | 0.089585927 |
| SPBC11G11.02c | 0.089655716 |
| SPCC584.13 | 0.089930729 |
| SPAC139.01c | 0.089974507 |
| SPBC6B1.02 | 0.090186071 |
| SPAC4H3.06 | 0.090244433 |

| | |
|---|---|
| SPCC4B3.07 | 0.090538401 |
| SPAC4D7.06c | 0.090635942 |
| SPAC13G7.13c | 0.090877636 |
| SPAC6F6.12 | 0.091001733 |
| SPCC622.19 | 0.091038467 |
| SPCC338.11c | 0.091179486 |
| SPBC3B9.15c | 0.091191725 |
| SPBC839.06 | 0.091283362 |
| SPAC13G7.05 | 0.091377119 |
| SPBC2F12.04 | 0.091695088 |
| SPAC18B11.07c | 0.091916417 |
| SPBC1921.06c | 0.091930454 |
| SPAC890.03 | 0.092000614 |
| SPCC584.11c | 0.092473072 |
| SPAC2F7.08c | 0.092595903 |
| SPCC1620.12c | 0.092620298 |
| SPBC4C3.04c | 0.092971799 |
| SPBC577.11 | 0.093086352 |
| SPBC30B4.03c | 0.093156443 |
| SPBC21.07c | 0.093392966 |
| SPAC186.05c | 0.093677011 |
| SPCC777.02 | 0.093757101 |
| SPCC24B10.16c | 0.093910386 |
| SPCC1235.11 | 0.094102466 |
| SPCC4G3.10c | 0.09421224 |
| SPAC11E3.12 | 0.094343113 |
| SPAC22A12.11 | 0.094465418 |
| SPBC1271.10c | 0.094520831 |
| SPAC926.06c | 0.094684031 |
| SPBC776.03 | 0.094946718 |
| SPCC61.03 | 0.095233792 |
| SPBC27.08c | 0.095307164 |
| SPAC19B12.04 | 0.095313424 |
| SPAC23C4.17 | 0.095393531 |
| SPAC24C9.15c | 0.095619496 |
| SPBC1734.12c | 0.096109561 |
| SPCC24B10.15 | 0.09612104 |
| SPBC409.08 | 0.096316799 |
| SPBC12C2.12c | 0.096541193 |
| SPAC22F8.03c | 0.096756223 |
| SPCC1442.17c | 0.097182927 |
| SPCC584.16c | 0.097581395 |
| SPCC622.15c | 0.097595998 |
| SPAC13G7.11 | 0.097779276 |
| SPCC1223.03c | 0.097946702 |
| SPAC343.10 | 0.098079537 |
| SPAC23H4.08 | 0.098091195 |
| SPCC4G3.03 | 0.098225651 |

| | |
|---|---|
| SPBP35G2.08c | 0.09849211 |
| SPBC2D10.07c | 0.098692852 |
| SPBC839.07 | 0.099004631 |
| SPAC6G10.02c | 0.099459858 |
| SPAC13G7.04c | 0.099515647 |
| SPAC227.10 | 0.099908908 |
| SPAC17G6.06 | 0.099947544 |
| SPBC2D10.12 | 0.100002236 |
| SPAC732.02c | 0.100197502 |
| SPAC750.06c | 0.100408809 |
| SPCC24B10.03 | 0.10158595 |
| SPCC16C4.09 | 0.101704566 |
| SPCC188.07 | 0.101981038 |
| SPAC1F3.05 | 0.102118864 |
| SPBC17G9.02c | 0.102886985 |
| SPBC354.05c | 0.102936613 |
| SPAC926.09c | 0.10308096 |
| SPAC3F10.11c | 0.103214547 |
| SPCC126.11c | 0.103326591 |
| SPAC4C5.03 | 0.103375756 |
| SPCC1281.04 | 0.103597342 |
| SPBC2F12.11c | 0.103642899 |
| SPAC1805.09c | 0.103828837 |
| SPAC3A11.05c | 0.104061407 |
| SPBC2F12.13 | 0.104085244 |
| SPAC22A12.03c | 0.104679593 |
| SPCC24B10.10c | 0.104720468 |
| SPBC6B1.05c | 0.104886917 |
| SPAC12B10.02c | 0.105174099 |
| SPBC106.20 | 0.105294833 |
| SPAC2E1P5.01c | 0.105438893 |
| SPAC3H1.05 | 0.105716709 |
| SPAC20H4.05c | 0.105959863 |
| SPCC11E10.09c | 0.10634678 |
| SPCC338.06c | 0.106360449 |
| SPCC417.09c | 0.106521332 |
| SPBC18E5.14c | 0.106532759 |
| SPAC9G1.11c | 0.106682127 |
| SPAC1250.02 | 0.107789657 |
| SPAC6G9.05 | 0.107856263 |
| SPCP1E11.04c | 0.108156858 |
| SPBC21B10.08c | 0.108172899 |
| SPBC12D12.04c | 0.108333273 |
| SPCPB16A4.06c | 0.108831404 |
| SPCC18B5.07c | 0.108846034 |
| SPAPB1A10.12c | 0.109118934 |
| SPAC22A12.07c | 0.10931702 |
| SPBC1706.01 | 0.109742473 |

| | |
|---|---|
| SPAC1B3.05 | 0.109816477 |
| SPCC24B10.11c | 0.109874704 |
| SPAC17A2.10c | 0.110703351 |
| SPBC30D10.09c | 0.111294594 |
| SPAC27F1.03c | 0.111560719 |
| SPAPB17E12.14c | 0.111596903 |
| SPAC17G8.07 | 0.112115323 |
| SPAC23D3.04c | 0.112473203 |
| SPBC2D10.14c | 0.112488223 |
| SPBC405.04c | 0.112807461 |
| SPCC16A11.10c | 0.113252557 |
| SPAC1071.07c | 0.113259251 |
| SPAC222.12c | 0.114055391 |
| SPAC1002.03c | 0.114414239 |
| SPCC1223.04c | 0.114887418 |
| SPBC409.16c | 0.115099499 |
| SPCC11E10.05c | 0.115108182 |
| SPAC1952.05 | 0.115563139 |
| SPAPB8E5.02c | 0.115934253 |
| SPBC839.11c | 0.116160976 |
| SPAC22F8.07c | 0.116513279 |
| SPCC594.06c | 0.116780322 |
| SPAC630.14c | 0.116788326 |
| SPBC582.09 | 0.116828872 |
| SPBC16G5.02c | 0.116990138 |
| SPBC21B10.03c | 0.118327159 |
| SPAC18G6.12c | 0.118436345 |
| SPBC3H7.07c | 0.11969544 |
| SPCC4B3.12 | 0.120485928 |
| SPBC2D10.20 | 0.121034735 |
| SPAC13A11.05 | 0.121043312 |
| SPBC365.13c | 0.121309842 |
| SPCC550.01c | 0.121950604 |
| SPAC637.09 | 0.122175003 |
| SPBC3E7.02c | 0.122373853 |
| SPAC20H4.10 | 0.122494611 |
| SPAC23G3.08c | 0.122558756 |
| SPBC4F6.12 | 0.12277368 |
| SPBC725.01 | 0.123207967 |
| SPAC227.14 | 0.123738259 |
| SPAC6B12.04c | 0.124202146 |
| SPAC1952.02 | 0.124488851 |
| SPAC144.04c | 0.125298995 |
| SPCC550.12 | 0.125869248 |
| SPAC17G8.09 | 0.125979556 |
| SPCC548.05c | 0.126066041 |
| SPCC576.13 | 0.126334696 |
| SPAC630.10 | 0.126467077 |

| | |
|---|---|
| SPAC3H1.10 | 0.126782403 |
| SPCC1450.16c | 0.126929625 |
| SPBC405.07 | 0.1272121 |
| SPBC23E6.05 | 0.127588563 |
| SPAC29B12.04 | 0.127827172 |
| SPAC19A8.04 | 0.127877438 |
| SPAC3H5.12c | 0.128925143 |
| SPAC20G8.08c | 0.129430717 |
| SPAC7D4.04 | 0.129595421 |
| SPAC31G5.17c | 0.129826623 |
| SPCC1235.01 | 0.130462084 |
| SPBC24C6.05 | 0.130825819 |
| SPAC56E4.03 | 0.131112452 |
| SPAP32A8.02 | 0.13141799 |
| SPCC1259.10 | 0.131808448 |
| SPAC17G8.10c | 0.13207315 |
| SPBC17A3.10 | 0.132159785 |
| SPAPB17E12.05 | 0.132513592 |
| SPAPJ691.03 | 0.132554053 |
| SPAC22E12.11c | 0.132631358 |
| SPAC1786.02 | 0.132740602 |
| SPBC800.11 | 0.133134743 |
| SPAC25G10.01 | 0.13338151 |
| SPCC4B3.02c | 0.133490894 |
| SPBC713.03 | 0.133774706 |
| SPAC227.05 | 0.133954685 |
| SPBC1861.01c | 0.134010944 |
| SPAC2F3.18c | 0.134323668 |
| SPAC2F7.02c | 0.134456568 |
| SPAC4F10.16c | 0.134607536 |
| SPBC651.05c | 0.134687732 |
| SPBC530.13 | 0.134814715 |
| SPCC188.08c | 0.13487198 |
| SPAC11E3.05 | 0.134961815 |
| SPAC6F6.01 | 0.135237932 |
| SPCC895.07 | 0.135516443 |
| SPAC1142.02c | 0.135876633 |
| SPAC16A10.05c | 0.136307703 |
| SPAPB1E7.11c | 0.136702048 |
| SPAC16E8.18 | 0.136857358 |
| SPCC1259.14c | 0.136995225 |
| SPAC22E12.05c | 0.137061705 |
| SPBC1348.14c | 0.137166283 |
| SPBC776.01 | 0.137579235 |
| SPAC144.06 | 0.13802984 |
| SPAC57A7.08 | 0.138246436 |
| SPBC215.01 | 0.138354704 |
| SPAC29B12.03 | 0.138447936 |

| | |
|---|---|
| SPCC4B3.03c | 0.138573815 |
| SPCC1795.01c | 0.138858585 |
| SPAC144.05 | 0.139074044 |
| SPBC1347.12 | 0.139915153 |
| SPCC24B10.18 | 0.140042201 |
| SPAPYUG7.03c | 0.140169232 |
| SPAC1952.16 | 0.140927023 |
| SPBC342.06c | 0.141095074 |
| SPAC23D3.12 | 0.141725907 |
| SPAC6F12.02 | 0.141946578 |
| SPAC3A12.12 | 0.141993185 |
| SPAC17D4.01 | 0.142142021 |
| SPBC56F2.03 | 0.142255504 |
| SPAC8E11.06 | 0.142361907 |
| SPCC1259.09c | 0.142394113 |
| SPAC25B8.11 | 0.142462585 |
| SPBPB7E8.01 | 0.142869679 |
| SPAC328.07c | 0.142916861 |
| SPAC2G11.09 | 0.143144833 |
| SPAC328.01c | 0.144168298 |
| SPCC622.16c | 0.144190232 |
| SPBC16H5.14c | 0.14451048 |
| SPAC631.01c | 0.144542731 |
| SPAC1783.02c | 0.145484354 |
| SPAPB17E12.02 | 0.145714704 |
| SPCC4B3.17 | 0.14572489 |
| SPBC428.15 | 0.145908844 |
| SPBC337.10c | 0.14594241 |
| SPAC17H9.04c | 0.146375644 |
| SPAC1002.05c | 0.146472016 |
| SPAC23A1.07 | 0.146758683 |
| SPAC4F8.01 | 0.147825112 |
| SPBC16G5.16 | 0.147975169 |
| SPBC11B10.02c | 0.148249771 |
| SPAC23A1.04c | 0.148461794 |
| SPCC777.06c | 0.14918376 |
| SPBC691.04 | 0.149234584 |
| SPCC4B3.05c | 0.149583611 |
| SPBC3E7.01 | 0.149611556 |
| SPAC1952.11c | 0.149695195 |
| SPBC2A9.02 | 0.149920072 |
| SPAC1006.06 | 0.149965418 |
| SPCC417.06c | 0.150438291 |
| SPAC31A2.02 | 0.150986373 |
| SPAC25A8.03c | 0.151645616 |
| SPCC16C4.01 | 0.15207222 |
| SPBC19F8.01c | 0.152676541 |
| SPAC823.14 | 0.153125051 |

| | |
|---|---|
| SPCC24B10.09 | 0.153313126 |
| SPBC23E6.02 | 0.153370612 |
| SPAC1751.01c | 0.15346049 |
| SPCC757.09c | 0.154207458 |
| SPBC16H5.12c | 0.156763273 |
| SPAC9.07c | 0.156850403 |
| SPBC17D11.04c | 0.157614629 |
| SPCC330.14c | 0.157621438 |
| SPBC1347.01c | 0.158258012 |
| SPBC1861.06c | 0.159084551 |
| SPBC15D4.13c | 0.159201633 |
| SPAC11E3.10 | 0.159202314 |
| SPAC31G5.03 | 0.159468214 |
| SPAC1A6.04c | 0.159808916 |
| SPBC1734.13 | 0.160024341 |
| SPCC4B3.10c | 0.160811048 |
| SPCC24B10.08c | 0.160855435 |
| SPAC4G9.02 | 0.16102282 |
| SPBC6B1.09c | 0.16105548 |
| SPAC23E2.01 | 0.161297083 |
| SPAC3H5.07 | 0.161549038 |
| SPBC337.09 | 0.161812768 |
| SPBC582.10c | 0.161845264 |
| SPCC1450.02 | 0.162103778 |
| SPAC30D11.05 | 0.162117174 |
| SPAC17D4.04 | 0.162457354 |
| SPAC26F1.14c | 0.16325844 |
| SPAC6B12.08 | 0.163523197 |
| SPBC11C11.10 | 0.163559486 |
| SPAC17G8.11c | 0.163840866 |
| SPBC21B10.10 | 0.163898315 |
| SPAC17A2.05 | 0.164491198 |
| SPAC30D11.12 | 0.165427156 |
| SPAC29A4.13 | 0.165658325 |
| SPCC584.01c | 0.166076205 |
| SPAC16.01 | 0.166130475 |
| SPBC19G7.17 | 0.166313763 |
| SPAC1006.01 | 0.166426602 |
| SPAC688.04c | 0.166437417 |
| SPBP8B7.26 | 0.166801297 |
| SPAC23G3.02c | 0.166905043 |
| SPAC631.02 | 0.167018675 |
| SPAC869.04 | 0.168691721 |
| SPCC777.08c | 0.170341436 |
| SPAC637.11 | 0.170401574 |
| SPAPB21F2.02 | 0.171069496 |
| SPBP16F5.03c | 0.171333636 |
| SPCC16A11.04 | 0.17156725 |

| | |
|---|---|
| SPBPB21E7.05 | 0.171990056 |
| SPCC61.05 | 0.172110721 |
| SPAC23H4.17c | 0.172584814 |
| SPAC1687.14c | 0.172853924 |
| SPAC2C4.08 | 0.173649626 |
| SPAC3F10.18c | 0.174126878 |
| SPCC1450.05c | 0.174350569 |
| SPCC1795.10c | 0.17451387 |
| SPBC1734.08 | 0.174618426 |
| SPAC13G7.12c | 0.174632376 |
| SPBC8E4.04 | 0.174926639 |
| SPAC18G6.15 | 0.178088658 |
| SPBC36.10 | 0.179460473 |
| SPBC17D1.06 | 0.18074448 |
| SPBC1271.14 | 0.180783164 |
| SPAC9.10 | 0.181025338 |
| SPAC1687.15 | 0.181063025 |
| SPAC2C4.06c | 0.181512632 |
| SPCC4G3.17 | 0.181833469 |
| SPBC1604.18c | 0.182285537 |
| SPCC1682.07 | 0.182444175 |
| SPBC31F10.16 | 0.182588071 |
| SPBC56F2.11 | 0.182851261 |
| SPAC22F3.06c | 0.186545545 |
| SPCC777.13 | 0.187668922 |
| SPCC1223.12c | 0.188678957 |
| SPCC4B3.11c | 0.188948206 |
| SPAP11E10.01 | 0.189432979 |
| SPAC8C9.06c | 0.189974441 |
| SPBC1703.08c | 0.190451469 |
| SPAC23A1.14c | 0.190680921 |
| SPAC17A2.06c | 0.192679964 |
| SPAC17A5.02c | 0.192723805 |
| SPAC644.09 | 0.193143625 |
| SPAC9E9.13 | 0.193418604 |
| SPCC23B6.02c | 0.194678016 |
| SPAC630.11 | 0.19598258 |
| SPAC17H9.03c | 0.196078083 |
| SPBC15D4.10c | 0.197074677 |
| SPCC126.01c | 0.197716156 |
| SPAC29E6.07 | 0.197834325 |
| SPAC29E6.01 | 0.198453865 |
| SPAC4G9.10 | 0.199722048 |
| SPBC36.06c | 0.199748889 |
| SPCC613.06 | 0.199823428 |
| SPBC1D7.03 | 0.200223557 |
| SPAC2E1P3.01 | 0.200624393 |
| SPBC15D4.02 | 0.201315041 |

| | |
|---|---|
| SPAC4H3.05 | 0.201319324 |
| SPCC622.18 | 0.201557684 |
| SPBC354.03 | 0.202810206 |
| SPAC57A10.14 | 0.205088852 |
| SPAC824.02 | 0.205152506 |
| SPBC3B9.09 | 0.205358752 |
| SPAC17H9.08 | 0.206982624 |
| SPCC1450.12 | 0.207088643 |
| SPAC589.02c | 0.207211836 |
| SPBC359.06 | 0.207558084 |
| SPAC9G1.07 | 0.207683401 |
| SPAC13G6.12c | 0.208286678 |
| SPCC550.03c | 0.208582421 |
| SPAC1327.01c | 0.208842966 |
| SPAC2H10.02c | 0.208950806 |
| SPBC29A10.01 | 0.20906604 |
| SPCC550.07 | 0.209505788 |
| SPBC1539.10 | 0.210022803 |
| SPAC22E12.19 | 0.21079881 |
| SPCC895.08c | 0.210833354 |
| SPAC3H1.09c | 0.211137078 |
| SPAC1527.02 | 0.211349942 |
| SPBP23A10.12 | 0.211826481 |
| SPAC18G6.05c | 0.212619858 |
| SPCC4B3.14 | 0.213182825 |
| SPCC1259.08 | 0.215149434 |
| SPAC3G9.08 | 0.217693245 |
| SPCC550.10 | 0.218298691 |
| SPAC1556.03 | 0.222027714 |
| SPCC4B3.13 | 0.223047595 |
| SPCC1795.09 | 0.223668979 |
| SPCC645.13 | 0.225907713 |
| SPCC23B6.03c | 0.226561704 |
| SPCC1753.02c | 0.227626821 |
| SPBP8B7.18c | 0.228563411 |
| SPAC30D11.10 | 0.228699766 |
| SPAC869.08 | 0.229179301 |
| SPCC1682.13 | 0.22930907 |
| SPAC1D4.11c | 0.229826863 |
| SPCC550.14 | 0.23011446 |
| SPAC3A12.08 | 0.230194836 |
| SPCC1393.08 | 0.230214815 |
| SPCC1259.13 | 0.230265628 |
| SPAC1805.08 | 0.231127828 |
| SPBC56F2.10c | 0.231639025 |
| SPCC24B10.17 | 0.231698856 |
| SPAC27E2.01 | 0.232366855 |
| SPBC32H8.02c | 0.232785406 |

| | |
|---|---|
| SPCC550.15c | 0.232894297 |
| SPCC1739.15 | 0.234118846 |
| SPAC3C7.12 | 0.237572687 |
| SPCC1259.03 | 0.237830384 |
| SPAC3A11.10c | 0.238858836 |
| SPCC970.01 | 0.241287377 |
| SPBC25H2.15 | 0.242222331 |
| SPBC21C3.01c | 0.242793876 |
| SPAC2F7.07c | 0.243736218 |
| SPCC74.05 | 0.244892674 |
| SPAC27E2.03c | 0.245072224 |
| SPBC4B4.11 | 0.245679394 |
| SPAC8C9.03 | 0.246511601 |
| SPCC645.12c | 0.248814102 |
| SPCC338.05c | 0.250114234 |
| SPAC13C5.07 | 0.250504176 |
| SPBC530.04 | 0.251060554 |
| SPBC16E9.19 | 0.251155636 |
| SPAC2F7.11 | 0.251905541 |
| SPBC1198.09 | 0.25272369 |
| SPAC824.09c | 0.255098307 |
| SPAP8A3.05 | 0.256420442 |
| SPBC1604.11 | 0.257223304 |
| SPCC1393.03 | 0.257681908 |
| SPAC17G6.05c | 0.259443957 |
| SPBC17A3.03c | 0.260038222 |
| SPCC1393.02c | 0.260572172 |
| SPCC1020.13c | 0.261015685 |
| SPBC4B4.03 | 0.262784112 |
| SPAC19G12.02c | 0.264611938 |
| SPCC11E10.06c | 0.264781255 |
| SPCC645.11c | 0.264930417 |
| SPCPB16A4.03c | 0.264963837 |
| SPAC139.04c | 0.265164839 |
| SPBC2A9.11c | 0.266886013 |
| SPCC285.05 | 0.266938104 |
| SPBP8B7.05c | 0.267972286 |
| SPAC24C9.14 | 0.268834933 |
| SPBC215.05 | 0.269670122 |
| SPCP1E11.03 | 0.27277295 |
| SPCC895.06 | 0.273046292 |
| SPAC13G7.09c | 0.274396189 |
| SPAC16A10.02 | 0.274442507 |
| SPAC25B8.17 | 0.274505746 |
| SPAC13C5.06c | 0.277030783 |
| SPAC644.14c | 0.277792567 |
| SPBC106.10 | 0.278530711 |
| SPBC646.12c | 0.279890326 |

| | |
|---|---|
| SPCC4E9.01c | 0.280237566 |
| SPCC622.01c | 0.280768219 |
| SPCC1183.06 | 0.285034084 |
| SPCC4B3.15 | 0.285206213 |
| SPBC4B4.06 | 0.287201182 |
| SPBC215.04 | 0.289918926 |
| SPAC26H5.03 | 0.28996352 |
| SPBC3B9.08c | 0.290695314 |
| SPCC550.09 | 0.294778792 |
| SPCC126.12 | 0.299686396 |
| SPAC1039.01 | 0.300499597 |
| SPBC27B12.08 | 0.300725071 |
| SPBC25B2.01 | 0.300969393 |
| SPCC1620.02 | 0.302276124 |
| SPCC1235.15 | 0.303275184 |
| SPAC694.05c | 0.303644443 |
| SPCC825.05c | 0.305344276 |
| SPBC106.04 | 0.30659062 |
| SPBC29A3.12 | 0.307608744 |
| SPAC3A12.09c | 0.307781613 |
| SPCC1739.07 | 0.307880049 |
| SPCC24B10.19c | 0.309734735 |
| SPCC1259.11c | 0.31037865 |
| SPCC1450.06c | 0.313234203 |
| SPBC29A10.16c | 0.317277957 |
| SPBC582.05c | 0.318421647 |
| SPCC1442.02 | 0.318610786 |
| SPAC19G12.15c | 0.319220374 |
| SPBC1711.03 | 0.322626377 |
| SPBC106.08c | 0.324298839 |
| SPBC1539.07c | 0.32524981 |
| SPCC4B3.01 | 0.328716343 |
| SPBC577.05c | 0.329988593 |
| SPAC1851.03 | 0.333754422 |
| SPCC550.08 | 0.336057034 |
| SPAC16.05c | 0.336378564 |
| SPAC1527.01 | 0.34168376 |
| SPAC1635.01 | 0.342546899 |
| SPAC513.05 | 0.34367171 |
| SPCC1281.07c | 0.3515056 |
| SPAC10F6.08c | 0.353346703 |
| SPBC336.01 | 0.35656296 |
| SPCC1393.12 | 0.35702724 |
| SPBC26H8.05c | 0.36436712 |
| SPBP8B7.13 | 0.36744357 |
| SPBC16A3.08c | 0.371439654 |
| SPCC2H8.02 | 0.373479542 |
| SPCC613.12c | 0.375377367 |

| | |
|---|---|
| SPCC1223.10c | 0.378106428 |
| SPCC18B5.10c | 0.381031717 |
| SPAPB18E9.01 | 0.394433466 |
| SPCC645.08c | 0.405480424 |
| SPAC1639.02c | 0.406239824 |
| SPAC26A3.06 | 0.40652419 |
| SPCC1322.01 | 0.417463821 |
| SPCP25A2.02c | 0.422520554 |
| SPAC5H10.08c | 0.437833099 |
| SPBC83.12 | 0.450962246 |
| SPAC15E1.07c | 0.455053525 |
| SPAC31G5.19 | 0.459129838 |
| SPAC140.01 | 0.462146946 |
| SPAC4G9.05 | 0.462801378 |
| SPCC23B6.04c | 0.474791423 |
| SPCC1183.10 | 0.479374911 |
| SPAC144.03 | 0.48489707 |
| SPBC577.04 | 0.485505874 |
| SPAC1F5.05c | 0.485902528 |
| SPAC13D6.02c | 0.486675818 |
| SPBC2G2.17c | 0.494836757 |
| SPAPB1E7.12 | 0.506197108 |
| SPCC825.02 | 0.514791352 |
| SPCC550.11 | 0.516916038 |
| SPCC338.04 | 0.521291237 |
| SPBC1703.03c | 0.529402885 |
| SPAC4D7.01c | 0.539908446 |
| SPAC222.04c | 0.55507034 |
| SPAC2C4.10c | 0.596311915 |
| SPBC1A4.03c | 0.637231921 |
| SPCC645.14c | 0.639551455 |
| SPBC27B12.07 | 0.661938682 |
| SPAC1F7.07c | 0.677038069 |
| SPBC19C7.10 | 0.680254574 |
| SPAC23H4.12 | 0.681919741 |
| SPBC1198.11c | 0.70563314 |
| SPCC338.12 | 0.766789643 |
| SPAC31A2.14 | 0.889454053 |
